# Anti-inflammatory role of curcumin in Lipopolysaccharide treated A549 cells at global proteome level and on mycobacterial infection

**DOI:** 10.1101/721100

**Authors:** Suchita Singh, Rakesh Arya, Rhishikesh R Bargaje, Mrinal Kumar Das, Subia Akram, Hossain Md. Faruquee, Rajendra Kumar Behera, Ranjan Kumar Nanda, Anurag Agrawal

## Abstract

A diet derived agent Curcumin (Diferuloylmethane), demonstrated its clinical application in inflammation, infection and cancer conditions. Nevertheless, its impact on the proteome of epithelial cells of non-small cell lung carcinoma (NSCLC) is yet to be explored. We employed a stable isotope labeling method for cell culture (SILAC) based relative quantitative proteomics and informatics analysis to comprehend global proteome change in A549 cells treated with curcumin and/or Lipopolysaccharide (LPS). Pretreated A549 cells were infected with *Mycobacterium tuberculosis* H37Rv strain to monitor bacterial load. With exposure to curcumin and LPS, out of the 1492 identified proteins, 305 and 346 proteins showed deregulation respectively. The expression of BID and AIFM1 mitochondrial proteins which play critical role in apoptotic pathway were deregulated in curcumin treated cells. Higher mitochondria intensity was observed in curcumin treated A549 cells than LPS treatment. Simultaneous treatment of curcumin and LPS neutralized the effect of LPS. Curcumin and/or LPS pretreated A549 cells infected with H37Rv, showed successful bacterial internalization. LPS treated A549 cells after infection showed increased bacterial load than curcumin compared to non-treated infected cells. However, simultaneous treatment of curcumin and LPS neutralized the effect of LPS. This study generated molecular evidence to deepen our understanding of the anti-inflammatory role of curcumin and may be useful to identify molecular targets for drug discovery.

## Introduction

Curcumin, a naturally occurring polyphenolic yellow pigmented dietary compound is found mainly in the rhizomes of *Curcuma longa*. The therapeutic and protective properties of curcumin have been extensively studied and reported to have anti-inflammatory, anti-oxidant and anti-carcinogenic activities (1, 2). Curcumin, is a potent inhibitor of NF-κB activation in response to various inflammatory stimuli (3, 4).

It inhibits the 12-O-tetradecanoylphorbol-13-acetate (TPA)-induced NF-κB activation by preventing the degradation of the inhibitory protein IκB; and the subsequent translocation of the p65 subunit in cultured human pro-myelocytic leukemia (HL-60) cells (5). Curcumin is reported to inhibit NO synthase (NOS) in RAW 264.7 cells activated with lipopolysaccharide (LPS) (6). Curcumin ameliorates inflammatory responses associated with asthma by activating Nrf2/HO-1 and down-regulate the expression of pro-inflammatory cytokines like TNF-α, IL-1β and IL-6 (7). In mouse model, curcumin relieve the psoriasis-like inflammation by decreasing the levels of IL-17A, IL-17F, IL-22, IL-1β,IL-6 and TNF-α cytokines (8). In high-fat diet (HFD) induced insulin resistance and type 2 diabetes, rats treated with oral curcumin showed an anti-hyperglycemic effect and improved insulin sensitivity, attributed by its anti-inflammatory properties as evident by attenuated TNF-α levels (9). Curcumin effectively inhibited prostate cancer proliferation, invasion, and tumorigenesis in both in vitro and in vivo models (10).

LPS present in the outer membranes of Gram-negative bacteria causes pro-inflammatory, pro-apoptotic, and anti-apoptotic pathways upon binding to macrophage TLR4 (11). LPS exposed macrophages show activation, altered morphology and response to various infection (12). Pathogens breach alveolar epithelium leading to internalization and colonization, causing infection. A549, human pulmonary adenocarcinoma cells, is a well-studied cellular model system for respiratory infection (13). Non-small cell lung carcinoma (NSCLC) develops inherent resistance to chemotherapy and therefore, discovering effective bioactive agents is the need of the hour. Curcumin has been reported to arrest cell-cycle followed by apoptosis in A549 cells (14).

In this study, we aim to explore the role of curcumin on LPS exposed A549 cells. LPS elicit a strong immune response in animals through Toll-like receptors (TLRs) mainly TLR4, that triggers the pro-inflammatory response to eradicate the bacteria (15). Establishing the mechanism of curcumin action on LPS treated and *Mycobacterium tuberculosis* (H37Rv) infected A549 cells might be useful to identify novel targets to reduce LPS induced damage to treat different disease conditions. Schematic experimental plan adopted in this study is presented in Fig. 1.

**Figure 1.**
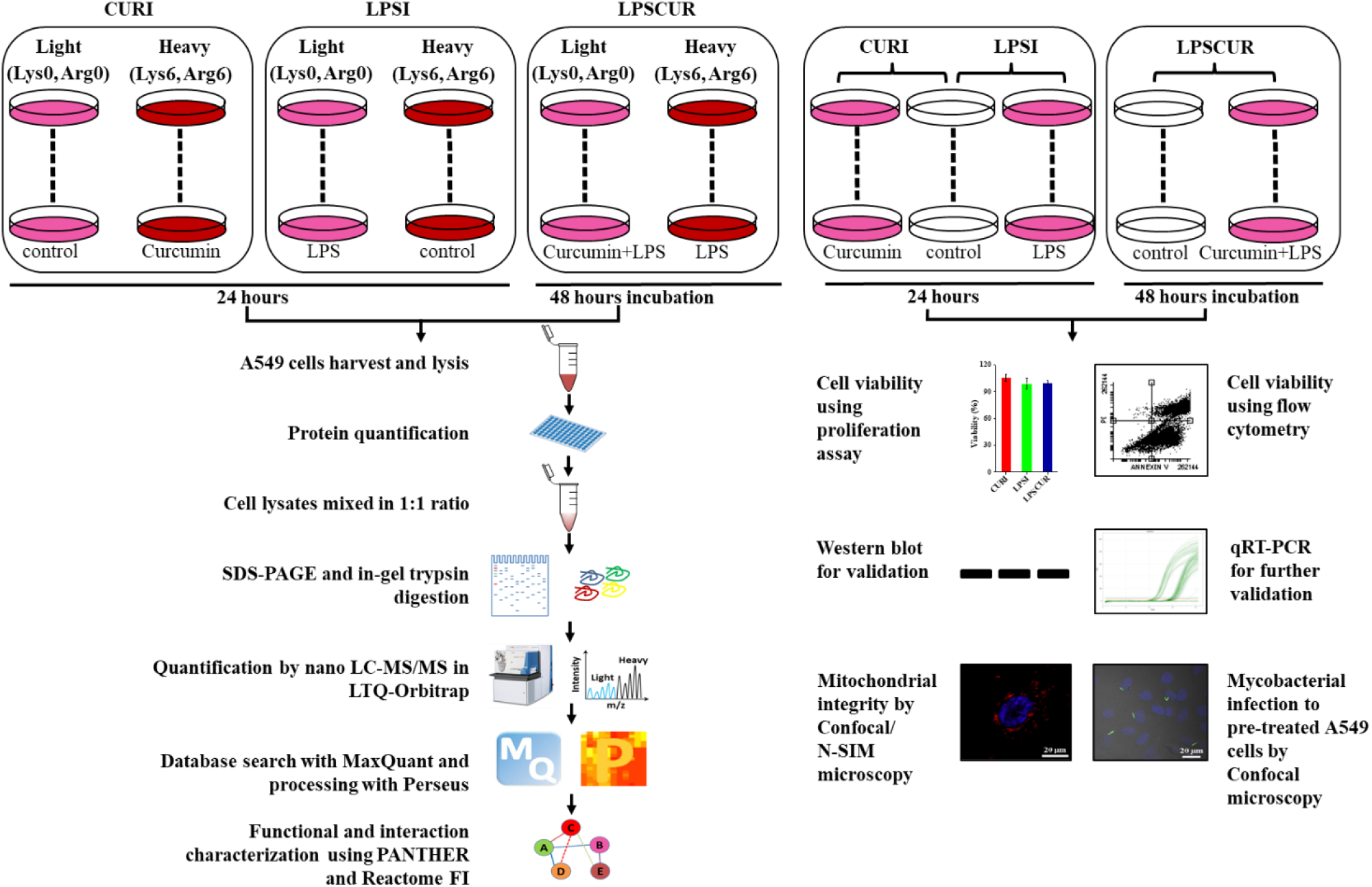
Experimental strategy adopted in this study to understand the anti-inflammatory role of curcumin in A549 cells treated with LPS. Experimental design used to analyze altered proteome in A549 cells treated with curcumin (10 μM) and LPS (1 μg/ml) in SILAC experiments and other function assays used in this study including Mycobacteria infection.

## Materials and Methods

### Cell Culture

Human lung adenocarcinoma cells (A549) procured from National Centre for Cell Science (NCCS), Pune were grown in SILAC DMEM-Flex Media without lysine and arginine (MS10030 Invitrogen, Oslo, Norway). The media was supplemented with dialyzed fetal bovine serum (10 % v/v, Invitrogen, Carlsbad, CA), L-glutamine (2 mM), glucose (8 mM, Sigma-Aldrich, St. Louis, MO), penicillin (50 U/ml), streptomycin (50 µg/ml, Invitrogen, Carlsbad, CA), light labeled (L-arginine-^12^C_6_, L-lysine-HCl-^12^C_6_) or heavy labeled (L-arginine-^13^C_6_, L-lysine-HCl-^13^C_6_) amino acids (Sigma-Aldrich, St. Louis, MO) and cultured at 37 °C in a 5% CO_2_ environment. After 6th passage, cells were checked for % incorporation of heavy amino acid by performing mass spectrometry before starting the drug treatment experiments.

### Treatment Method of Cultured Cells

A schematic detail of the experimental setups used in this study is presented in Figure 1. In the first experimental set up (CURI), cells were grown in presence of heavy labeled amino acids with curcumin (10 µM) and light labeled amino acids for 24 hours at 37 °C with 5% CO_2_. In the second experiment (LPSI), cells were grown in light labeled amino acids with LPS (1 µg/ml) and heavy labeled amino acids for 24 hours at 37 °C. In the third experiment (LPSCUR), cells were grown with light labeled amino acids with both LPS (1 µg/ml) and curcumin (10 µM, Sigma-Aldrich, St. Louis, MO) and heavy labeled amino acids with LPS (1 µg/ml) cultured for extended period of 48 hours at 37 °C.

Cell viability was estimated using the CellTiter 96® AQueous One Solution Cell Proliferation Assay (Promega, Madison, USA). Briefly, 10 µL of reagent was added per 100 µL of culture media in each well, the plates were incubated at 37 °C for 1 hour, and the absorbance was read at 490 nm using a microplate reader (SpectraMax, Molecular Devices Co., Sunnyvale, CA, USA).

### Cell apoptosis measurement

A549 cells were seeded (6×10^5^ per well) on six-well plates and treated with either curcumin or LPS or both for 24/48- hour duration. Adherent cells were scrapped and harvested by centrifuging at 371 *× g* for 5 minutes at 4 °C. The cells were washed with phosphate buffer saline (PBS) and then binding buffer and stained with annexin V-FITC and Propidium iodide (50 µg/ml; Sigma-Aldrich, St. Louis, MO) for 15 min. The stained cells were analyzed immediately using flow cytometry (BD FACSCantoII) with FITC (Em: 525 nm) and PerCP (Em: 675 nm) filters. Flowing Software (ver. 2.5.1, University of Turku, Finland) was used for the data analysis.

### Sample Preparation for Cell Proteome Analysis

After the mentioned incubation periods, cells were harvested by centrifugation (15,000 *×g* for 20 minutes at 4 °C), and washed twice with chilled PBS. Cells were incubated with lysis buffer (4 % SDS, 0.1 M DTT, and 0.1 M Tris-HCl pH 7.5) at 95 °C for 5 minutes. After sonication (40 % amplitude, 10 sec ON/OFF pulse for 2 minutes), cell lysates were centrifuged (15,000 *×g* for 20 minutes at 4 °C) to collect the supernatant. Protein quantification was carried out using BCA (Bicinchoninic Acid, Sigma-Aldrich, St. Louis, MO) method taking bovine serum albumin (BSA) as standard.

Equal amount of proteins (100 µg) from each experimental sets were pooled together (100 µg of light and heavy samples) and separated on a 10% SDS-PAGE gel and in-gel trypsin digestion was carried out. Briefly, after staining the gel with colloidal commassie Blue, each gel lane was cut into 8/10 slices in different tubes. Colloidal commassie Blue was removed by washing with ammonium bicarbonate (50 mM) and acetonitrile solution (50 %). Excised gel pieces were treated with acetonitrile (100 %) for 5 minutes and dried in a speed-vac. Proteins were reduced with 150 µl of DTT (10 mM) in ammonium bicarbonate buffer (100 mM) for 30 minutes at 56 °C and alkylated with 100 µl of iodoacetamide (50 mM) in 100 mM ammonium bicarbonate for 30 minutes at room temperature in dark. Gel pieces were again washed and dehydrated using ethanol at room temperature. Trypsinisation was carried out by incubating the gel pieces overnight at 37 °C in 50 µl of trypsin (1: 25:: trypsin: protein, 20 µg/ml, sequencing grade trypsin V5111, Promega, Madison, WI) prepared in ammonium bicarbonate (50 mM). Tryptic peptides were extracted using acetonitrile (60 %) and formic acid (0.1 %) for three times (150 μl). The extracted tryptic peptides were dried at 37 °C for 3-4 hours. The dried samples were finally resuspended in 20 µl of formic acid (5 %) in ammonium bicarbonate (100 mM) just before proceeding with LC-MS/MS analysis.

### NanoLC-MS/MS Analysis

Tryptic peptides were analyzed using an LTQ-Orbitrap XL (Thermo Scientific, Berman, Germany) equipped with a nano-LC system (EASY-nLC; Proxeon Biosystems). Tryptic peptides (5 µg) were loaded onto a 0.3 x 5 mm capillary trap column and then separated by reverse-phase chromatography on 15 cm long column (inner diameter 75 µm, 3 µm Reprosil-Pur C18-AQ medium) at a flow rate of 500 nL/min. Buffer A consists of formic acid (0.1%) and buffer B acetonitrile with 0.1% formic acid. The gradient was 0 to 108 min of 3-35% buffer B, 108-110 min of 35-100% buffer B and last 10 min of buffer B with a total run time of 120 min was used throughout the study. Mass spectra were acquired in the positive ion mode applying a data dependent automatic (DDA) switch between survey scan and MS/MS acquisition. Mass spectra from survey full scans (m/z 350-2000) were acquired on the Orbitrap. The resolution of the instrument was set to 60,000 with a target value of 1E6 charges in the LTQ mass analyzer. The six most-intense precursor ions were selected from the data dependent MS scan for subsequent collision-induced dissociation (CID). Fragmentation in the LTQ was performed with a target value of 5,000 ions, normalized collision energy of 35% and ion selection threshold of 1,000 counts. Charge state screening was enabled for +2, +3, +4, and above ions. The parameters that were used for dynamic exclusion were as follows: repeat count-1, repeat duration-20, list size-500, exclusion duration-40. A total of 8 LC-MS/MS data files for CURI, 10 each for LPSI and LPSCUR were generated.

### Data Analysis for Cell Proteome and Phosphoproteome using MaxQuant and Perseus

All raw MS files from each experiment were processed for peak detection and quantification by using MaxQuant (1.5.2.8). Peptide identification was performed by using the Andromeda search engine and Swiss-Prot database (89,796 human protein entries as on 10^th^ January 2015). The experimental design included all three experiments (CURI, LPSI and LPSCUR) following the steps as described earlier. Search criteria used in this study were trypsin specificity, fixed modification of carbamidomethyl (cysteine), variable modifications of oxidation (methionine), and allowed for up to two missed cleavages. A minimum of six amino acids in the peptide length were required for identification. By using a decoy database strategy, peptide identification was accepted based on the posterior error probability with a false discovery rate (FDR) of 1%. The precursor mass tolerance was set at 6 ppm and the fragments ion tolerance was 20 ppm. The “match between runs” option in MaxQuant was enabled within a time window of 2 minute, to match the identifications across adjacent fractions. Quantification of SILAC pairs was performed by MaxQuant with a minimum ratio count of 2. All the statistical analysis of the MaxQuant output tables were performed with the Perseus software (1.5.1.6). In Perseus, the light and heavy labeled peptide intensities were loaded for analysis. The potent contaminants (trypsin and keratin) were filtered out and fold change in protein abundance were expressed in log_2_ scale. Proteins with at least a change of 2 fold higher or lower abundance (log_2_ fold change ≥ ±1) were considered as significant and selected for further analysis **(Table S1).** The results obtained from MaxQuant were further evaluated by Perseus and plots were generated using OriginPro 8.0 software.

Phosphopeptides were identified using an additional variable modification of phosphorylation (Serine, Threonine and Tyrosine: STY). The precursor mass tolerance was set at 4.5 ppm and the fragments ion tolerance was 20 ppm. The minimum peptide length was set to 7 with matching time window of 0.7 min. For identification of phosphorylation sites, the maximum FDR was set to 0.01. Phsophorylation sites were considered as “class I” if the localization probability was at least 0.75 (75%) and the localization score difference is greater than 5 **(Table S2A-C)** (43, 44). The number of phosphosites, distribution of charge states and number of phosphogroups per peptide in each CURI, LPSI and LPSCUR experiments were calculated using the phospho (STY) sites file obtained as output from MaxQuant **(Table S3)**.

### Data Availability

All the MS/MS proteomics data files (.raw and .mgf), phosphoproteome data and output tables of MaxQuant analysis have been deposited in the ProteomeXchange Consortium (PXD012652) (45).

### Analysis of protein-protein interactions and functional characterization

The deregulated protein lists from CURI, LPSI and LPSCUR were converted to gene list by employing Uniprot “Retrieve ID mapping” option. The gene list was used as input to Reactome Functional Interaction (*ver. 4.2.0 beta*) application in Cytoscape (*ver. 3.4.0*) to construct network with 2015 algorithm for “gene sets” using “Fetch FI annotations” and “use linker genes”. Genes were clustered by the spectral partitioning-based network clustering algorithm using the clustering FI network option (46). The FDR for the analysis of GO Biological processes was set to 0.05. Further network functions were analyzed for “Gene Ontology Biological Processes” of all clusters. To get molecular function information, Protein ANalysis THrough Evolutionary Relationships (PANTHER) database (ver. 11.0) was used and the deregulated genes in CURI, LPSI and LPSCUR experiments were used as input parameters.

### Monitoring Important proteins using Western Blot Experiments

Important mitochondrial proteins (log_2_ fold change ≥ ±1) identified in any of these experiments, were monitored in a separate set of experimental protein samples extracted from control and treated cells. Isolated proteins were separated on 12% SDS-PAGE and transferred to a PVDF membrane (FluoroTrans, Pall Life Sciences, USA). After blocking with BSA (3 %) in TBST, blots were incubated with primary antibodies BID (1:1000, sc-373939, Santa Cruz, CA, USA), AIFM1 (1:1000, sc-55519, Santa Cruz, CA, USA) and β-actin (1:1000, sc-47778, Santa Cruz, CA, USA) overnight at 4 °C. Blots were then washed and incubated with goat anti-mouse secondary antibody labeled with HRP (sc-2060, Santa Cruz, CA, USA) and protein bands were visualized using the SuperSignal West Pico PLUS Substrate kit (Thermo Fisher Scientific, Waltham, MA). Relative band intensities were calculated using ImageJ software (Bethesda, MD, USA) with respect to β-actin which was used as loading control.

### qRT-PCR

To monitor expression of BID and AIFM1 gene at transcript level, qRT-PCR was performed from a separate set of treated samples. β-actin mRNA was used as an endogenous control. Total RNA from A549 cells (1 × 10^6^ cells) was extracted using miRNeasy Mini kit (Qiagen, Denmark) according to the manufacturer’s instructions and eluted in 60 µl of nuclease free water. The purity and concentration of the isolated RNA was measured by Nanodrop 2000 spectrophotometer (Thermo Fisher Scientific, Waltham, MA). Total RNA (1 µg) from each sample sets (n=3) was taken for cDNA preparation using iScript™ cDNA Synthesis Kit (Bio-Rad, USA). Real-Time qPCR was performed using PowerUp SYBR Green Master Mix (Thermo Fisher Scientific, Waltham, MA). qPCR was performed in a total reaction volume of 10 µl, including 5 µl of Master Mix, 1 µl each of forward and reverse qPCR Primers (5 mM), 2 µl of cDNA (10 ng) and 1 µl double-distilled water. Reactions were performed and analysed using a CFX96 Real-Time PCR Detection System (Bio-Rad, USA). Primers used for β-actin, 5’-ATTGCCGACAGGATGCAGAA- 3’ (forward) and 5’- GCTGATCCACATCTGCTGGAA-3’ (reverse), for BID 5’- GACTGTGAGGTCAACAACGG-3’ (forward) and 5’-GCTTTGGAGGAAGCCAAACA-3’ (reverse) and for AIFM1 were 5’-TAGGGCTGACACCAGAACAG-3’ (forward) and 5’- GTGCCTCCACCAATTAGCAG-3’ (reverse). qPCR reactions were performed for each cDNA sample in triplicate for each gene with non-template control in duplicates and repeated at least three times. Cycling parameters used for qPCR was as follows: UDG activation for 2 min at 50 °C, Dual-Lock DNA polymerase step for 2 min at 90 °C, and an initial denaturation step of 3 sec at 95 °C with 40 cycles and annealing/extension of 30 sec at 62 °C. The relative quantity of mRNAs was determined using the CFX Manager software (ver. 3.1).

### Monitoring Mitochondrial distribution in treated A549 cells using Confocal Microscopy

A549 cells were grown overnight on 18 mm coverslip in twelve well plates at 37 °C with 5% CO_2_ and incubated with curcumin/LPS/both for appropriate durations. Cells were washed 3 times with PBS and stained with MitoTracker® red (200 nM, Thermo Fisher Scientific, Waltham, MA) for 30 minutes at 37 °C at 5% CO_2_. After washing with PBS, cells were fixed using 4% paraformaldehyde for 15 minutes at room temperature in dark. Cell nucleus was stained using DAPI (1 μg/mL, D9542, Sigma Aldrich, USA) and after washing with PBS/MilliQ water mounted onto a glass slide using ProLong Gold antifade reagent (Life Technologies, USA). Cells were visualized using 4′,6-diamidino-2-phenylindole (DAPI; Ex/Em-358/461 nm) or Tetramethylrhodamine (TRITC; Ex/Em-551/567 nm) and TD (Bright field) filters in a Nikon confocal microscope system A1R (Nikon USA, Melville, NY). ImageJ software (Bethesda, MD, USA) was used to calculate mean intensity of three focal areas of each slide in triplicates.

### Infection experiments

A549 cells cultured in DMEM/F-12 supplemented with 10% FBS media, were pretreated with curcumin, LPS or both, for the stipulated period before incubating with GFP-labeled *Mycobacterium tuberculosis* strain, H37Rv, at a multiplicity of infection (MOI) of 1:5 for 6 hours at 37 °C with 5% CO_2_. All these infection experiments were carried out at Tuberculosis Aerosol Challenge Facility at ICGEB, New Delhi following Biosafety Label-III guidelines. After removing extracellular bacteria by washing the infected cells with PBS and supplemented with fresh media, cells were incubated for 24 hours at 37 °C with 5% CO_2_. Post incubation, cells were processed further to perform confocal microscopy and colony forming unit (CFU) experiments in triplicates. For confocal microscopy experiments, infected cells grown on glass cover slips were washed with PBS (1x) and fixed using freshly prepared paraformaldehyde (4%) for 15 min at room temperature in dark. After washing with PBS, the fixed cells were incubated with 4′,6-diamidino-2-phenylindole (DAPI, 1 µg/ml) for 15 min at room temperature in dark. Cells were then washed with PBS and mounted onto a glass slide using ProLong Gold antifade reagent (Life Technologies, USA). Image were acquired by NIS-Elements AR analysis software (ver 4.0, Life Technologies, USA) using the Nikon A1R Laser Scanning Confocal Microscope equipped with a Nikon Plan Apo 60 X objective lens with 1.40 numerical aperture (NA) oil; immersion objective lens and were visualized using DAPI (Ex/Em-358/461 nm), PE/AlexaFluor488 (Ex/Em-490/525 nm), TRITC (Ex/Em-551/567 nm) and TD (Bright field) filters. For CFU, infected cells were washed with PBS (1x) thrice and then lysed with 0.06% SDS. Diluted cell lysates were plated onto 7H11 plates in duplicates and incubated at 37 °C. CFU count was performed after 15 days of incubation and expressed in log_10_(CFU/ml). The expression level of BID and AIFM1 in the Mtb infected samples were monitored by Western blot analysis using Glyceraldehyde 3-phosphate dehydrogenase (GAPDH) as loading control.

## Results

Effect of curcumin and/or LPS on A549 cell viability as shown in Fig. 2A, treatment with curcumin (10 μM: CURI) or LPS (1 μg/ml: LPSI) for 24 hours did not show any significant reduction in cell viability. Cells treated with both LPS and curcumin (LPSCUR) showed a 23.3% reduction in viability. DNA contents of A549 cells in CURI, LPSI and LPSCUR showed curcumin or LPS inhibits cell cycle arrest (Fig. 2B, **Fig. S1).** The Annexin V-positive cells were significantly increased as a result of cells undergoing early-apoptotic at 85.1%, 86.1% and 72.1% in CURI, LPSI and LPSCUR respectively (Fig. 2C). The apoptotic cells in LPSCUR were lowest with comparison to either curcumin or LPS treatment.

**Figure 2.**
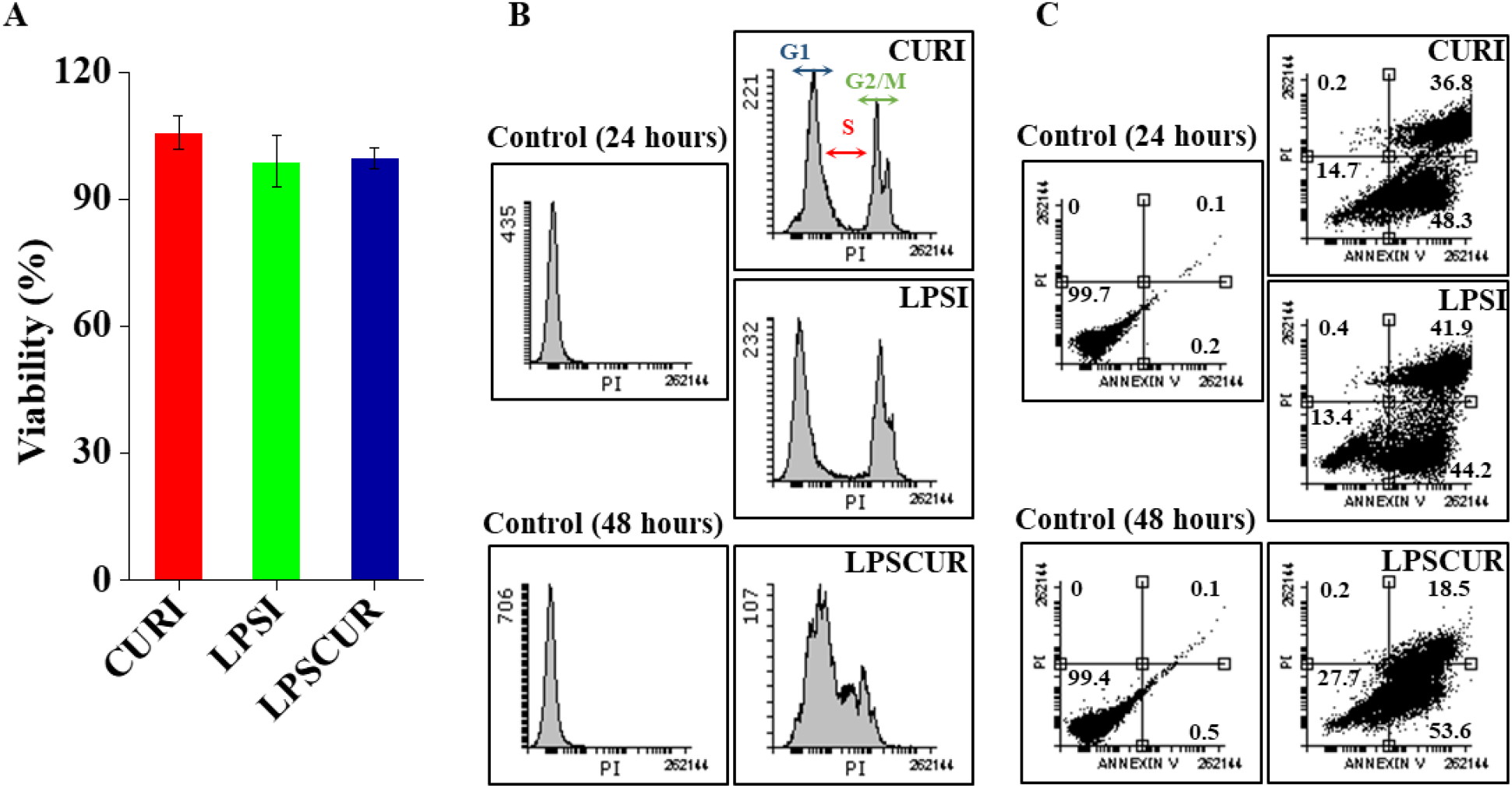
Effect of curcumin and/or LPS on cell viability and apoptosis of A549 cells. A549 cells were treated with curcumin (10 µM) and/or LPS (1 µg/ml) for different time. (A) Cell viability was monitored by CellTiter 96® AQueous One Solution Cell Proliferation Assay. Data is presented as mean ± Standard error of mean of six data points. (B) Cells were fixed and then stained with propidium iodide (50 µg/ml) and Annexin V-FITC. The DNA contents were analyzed by flow cytometry. Representative result from three independent experiments is presented. x- and y-axis present DNA content and cell number, respectively. (C) Cells (in %) undergoing early-apoptotic or late-apoptotic phase were calculated using Flowing software.

To characterize the temporal proteome changes in A549 cells in response to curcumin and/or LPS treatment, a comparative quantitative proteomics approach was adopted. We cultured the cells in heavy isotope labeled media for over 10 doubling period and confirmed the incorporation of heavy isotope to be over 99 % by using mass spectrometry. In the CURI, LPSI and LPSCUR experiments, a total of 1109, 1135 and 1155 proteins respectively were identified at 1% false discovery rate (FDR) at both protein and peptide levels (Fig. 3A). Altogether we identified a total 1492 distinct proteins at least one of these 3 experimental set **(Table S1)**. A set of 81, 204 and 78 unique proteins were identified in CURI, LPSI and LPSCUR treated cells respectively and 778 proteins were common. Functional interaction network of common proteins resulted in 696 original nodes with 73 linker genes. These genes were clustered in 7 modules with a modularity of 0.53 and contribute to different gene ontology (GO) biological processes viz. translation, protein folding, respiratory electron transport chain, RNA splicing and TCA cycle (Fig. 3B). A total of 804, 789 and 783 proteins showed similar abundance in cells treated with CURI, LPSI and LPSCUR respectively (Fig. 3C). Common 427 proteins with similar abundance mostly represent house-keeping genes, forms a network with 8 modules (385 nodes with 47 linker genes) and 0.53 modularity. The identified common proteins are involved in major GO biological processes like translation, protein folding and apoptosis process (Fig. 3D). Venn diagram representing the total number of deregulated proteins (log_2_ fold change ≥ ± 1) in CURI, LPSI and LPSCUR presented in **Fig. S2**.

**Figure 3.**
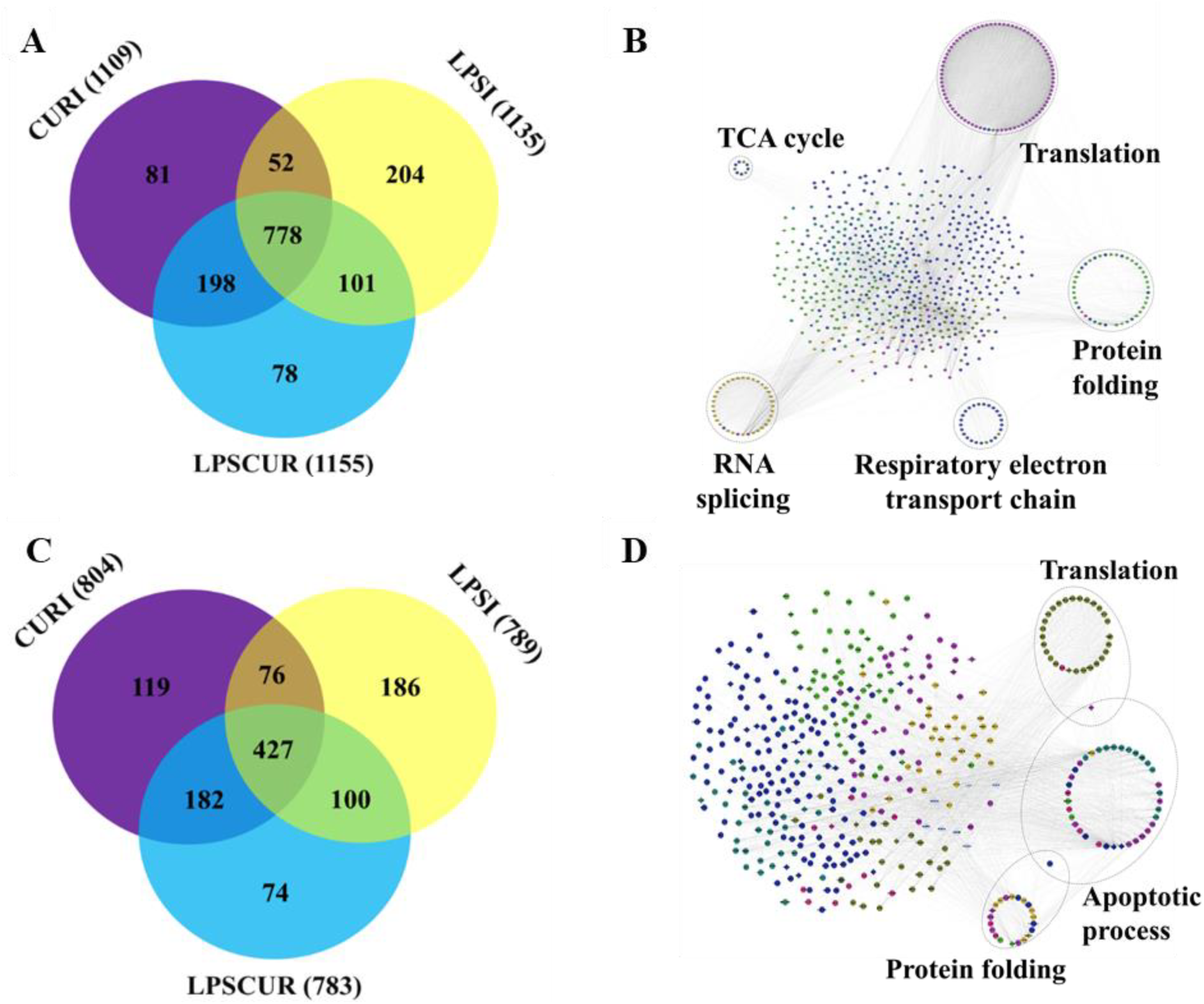
Global A549 proteome and common proteins identified in curcumin and/or LPS treated cells. (A) Venn diagram representing the total number of proteins identified in CURI, LPSI and LPSCUR experiments. (B) Subnetwork of 778 common genes (696 were mapped to the database with 65 linker genes) to all 3 experiments (CURI, LPSI and LPSCUR). (C) Venn diagram representing the proteins identified in CURI, LPSI and LPSCUR experiments with similar abundance in expression. (D) Subnetwork of 427 common genes (385 genes were mapped to database with 47 linker genes) with similar abundance in all the 3 experiments are shown. Color coded modules in which the interaction of the genes are highest based on the modularity. The functions of the genes are predicted by Reactome FI plugin in Cytoscape 3.4.0. The circled maps of the subnetworks involve different biological processes. Linker genes are shown by diamond shaped nodes.

In CURI, out of 1109, 3 proteins were up-regulated; while 302 were down-regulated. In LPSI, 333 proteins were up-regulated and 13 proteins were found to be down-regulated (Fig. 4A). Out of the 305 deregulated proteins in CURI, 257 nodes were mapped with 75 linker genes resulting 13 modules **(Fig. S3A)**. In case of 346 deregulated proteins in LPSI, 265 nodes with 65 linkers formed a network resulting 12 modules **(Fig. S3B)**. Modularity of CURI and LPSI were calculated to be 0.45, 0.65 respectively. These proteins are involved in diverse molecular functions viz. translation regulator activity, binding, receptor activity, structural molecule activity, catalytic activity, antioxidant activity and transporter activity. Molecular functions like binding and catalytic activity were found to be higher in LPSI and reversely regulated in CURI (Fig. 4B). Functional network analysis of 90 common deregulated proteins between CURI and LPSI experiments resulted a network with 35 nodes. The genes involved in translation process account for ~31.4 %, RNA splicing ~20 % and other linked genes ~48.6 % **(Fig. S3C)**.

**Figure 4.**
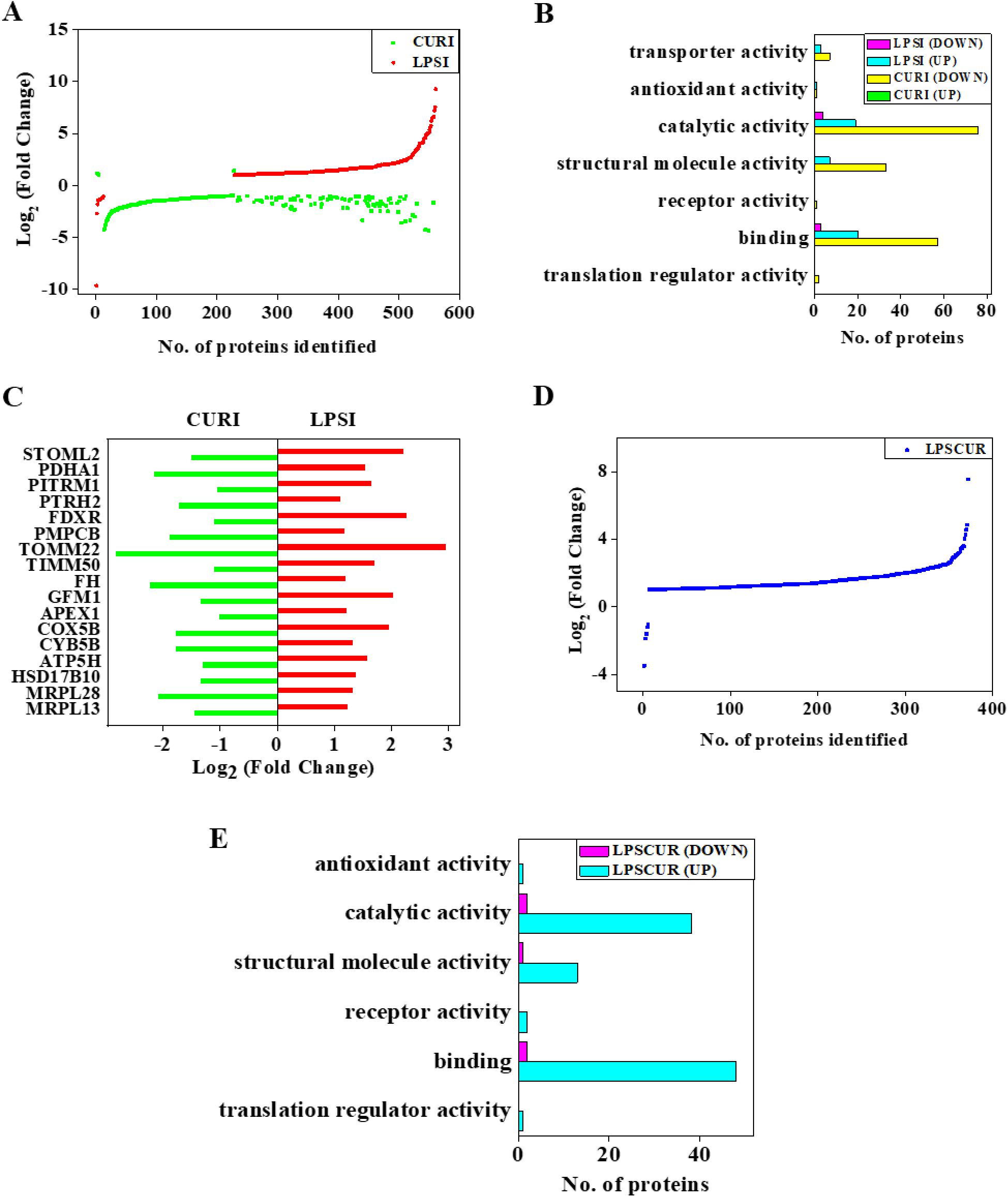
Deregulated global proteome in the A549 cells treated with curcumin and/or LPS. (A) Treatment of curcumin (10 µM) or LPS (1 µg/ml) for 24 hours alters proteome levels in A549 cell lines. Subnetwork in (B) Functional categorization and distribution of deregulated proteins in CURI (305) and LPSI (346) experiments. (C) Mitochondrial proteins showing inverse relationship in both CURI and LPSI experiments. (D) Simultaneous treatment of curcumin (10 µM) and LPS (1 µg/ml) for 48 hours induced proteome change in A549 cell lines. (E) Functional categorization and the distribution of LPSCUR (372) differentially expressed proteins to extract the information of their known functions.

Interestingly, a set of mitochondrial proteins (n=17, STOML2, PDHA1, PITRM1, PTRH2, FDXR, PMPCB, TOMM22, TIMM50, FH, GFM1, APEX1, COX5B, CYB5B, ATP5H, HSD17B10, MRPL28 and MRPL13) showed significant inverse relationship in protein abundance between LPS and CUR treated cells (Fig. 4C). BID protein showed significant down regulation on curcumin treatment. AIFM1 showed similar abundance on curcumin treatment and could be an important target of curcumin in apoptotic pathway.

In LPSCUR experiment, 366 proteins were found to be up-regulated and 6 were down-regulated (Fig. 4D). From the 372 deregulated proteins, 317 nodes with 76 linker genes generated 9 modules with 0.54 modularity **(Fig. S4)**. Majority of these proteins explained GO biological processes like translation, protein folding, protein targeting to mitochondrion, RNA splicing and respiratory electron transport chain (Fig. 4E). Majority of the up- regulated proteins in LPSCUR involves in binding and catalytic activities (Fig. 4E).

In fact, 878 proteins were found to be common between LPSI and LPSCUR. A set of proteins (8.77%) were up-regulated in LPSI and did not show any change in both LPS and curcumin treated cells **(Fig. S5)**. These variations may be due to the different incubation periods in LPSI (24 hours) and LPSCUR (48 hours). BID and AIFM1 proteins up-regulated on simultaneous treatment with respect to controls.

From the identified protein sets, 11 (5/6: up/down) and 17 (12/5: up/down) deregulated phosphoproteins were observed in CURI and LPSI respectively, whereas 5 deregulated phosphoproteins were observed in LPSCUR condition. Majority of these phosphosites were found to be contributed from Serine residues in CURI and LPSI experiments, but none in LPSCUR experiment. Most of these phosphoproteins in CURI, LPSI and LPSCUR were singly or doubly phosphorylated **(Fig. S6A-C)**.

Two major proteins from the apoptotic pathway, BH3-interacting domain death agonist (BID) and apoptosis-inducing factor protein 1 (AIFM1-1) were qualified at least two-fold up or down regulations in treated condition and identified as important molecules. Relative expression of BID and AIFM1 with respect to β-actin were probed using Western blot experiments using an independent sample set not used in discovery. AIFM1 and BID expression showed similar fold changes in western blot experiment and supported Mass Spectrometry data **(Fig. S7)**. In LPSCUR experiments, expression of these important molecules showed similar trend in both western blot and mass spectrometry fold change values (**Fig. S7)**. Additional blots and corresponding silver stained gels for loading controls are presented in **Fig. S7**.

From the confocal/N-SIM microscopy experiments, we observed the mean intensity of mitochondria in CURI was significantly higher than control and remain unchanged in LPS exposed cells **(Fig. S8A-B)**. Low mean mitochondrial intensity was observed in LPSCUR with respect to 48 hours control with statistically insignificant change. At the transcript levels of BID and AIFM1, we did not find significant variation in their expression in these experiments (**Fig. S8C)**.

Pretreated A549 cells, with CURI, LPS or LPSCUR, when infected with virulent laboratory strain *Mycobacterium tuberculosis* (H37Rv) showed successful internalization and varied bacterial load (Fig. 6A-B, **Fig. S9A-C)**. We observed higher bacterial load in cells pre-incubated with LPS, however curcumin pre-exposed cells (CURI) showed similar bacterial load compared to controls. Whereas at 48 hours post infection cells exposed to both LPS and curcumin (LPSCUR) showed lower but statistically insignificant difference in bacterial load with respect to controls.

The changes in the expression of BID and AIFM1 was analysed after treating the A549 cells with curcumin and/or LPS and then infecting with *Mtb* (Fig. 6C). The expression level of BID was further reduced in CURI (log_2_FC= −7.87) experiment and concordant with the mass spectrometry data (log_2_FC= −1.31). Also, the AIFM1 protein showed upregulation in LPSCUR (log_2_FC= 2.55) experiment and matches with the mass spectrometry data (log_2_FC= 1.32).

## Discussion

Curcumin has been used as a dietary compound and by recent evidence demonstrated its chemopreventive and chemo-protective role as suggested on its tumour suppressive and apoptosis inducing activities (20). Here we report that curcumin has anti-inflammatory role and impact cell proteome changes that negates the damage caused by LPS on A549 cells.

Curcumin and/or LPS treatment, at the concentration and time duration used in this study, did not affect growth of the A549 cells. One of the major checkpoints in cell cycle is G2/M and defective checkpoint function leads to tumorigenesis. Therefore, the regulation of checkpoint signalling is important and it can alter tumor cells sensitivity to drugs (21). However, the major strategy of eliminating tumor cells is by arresting the cell-cycle (22). Curcumin and LPS treatment results in cell cycle arrest in A549 cells at G2/M phase. Our data corroborated the earlier finding of curcumin on human melanoma cells (23). But upon simultaneous treatment with LPS and curcumin, there is a decrease in the G2/M phase cells and increase in S phase cells suggesting chemo protective role of curcumin in presence of LPS.

Our global proteomics experiment on A549 cells treated with curcumin and/or LPS generated important data. Curcumin and/or LPS altered global A549 proteome. Curcumin induces apoptosis through perturbing mitochondria molecular pathways which are reported to be activated during pharmacological deregulation of proteins and proteasome function. This results into the deregulation of several proteins involved in the biogenesis, transport and activity of mitochondria on curcumin treatment. Interestingly, 17 mitochondrial proteins inversely expressed on curcumin or LPS treatment which suggests that curcumin reduces abundance of majority of proteins involved in apoptotic pathway. How such changes in presence of curcumin disturbs the cellular functions is not yet clear? But it could do so by binding and/or regulating some of the important proteins required for the gene expression (24, 25). These findings demonstrated the antioxidant activity of curcumin (26). Bacterial LPS or endotoxins are responsible for the activation of the innate immune response and antigen-specific immune response by binding to various cell surface receptors (27). We analyzed the anti-inflammatory effect and mechanism of action of curcumin in regulating the TLR-4 signaling pathway in A549 cells. It has been reported that curcumin is involved in the inhibition of many important processes such as homodimerization of TLR-4 (28). We identified an Interferon-induced, double-stranded RNA-activated protein kinase (PKR) in both CURI and LPSI which showed similar expression level (data not shown). PKR plays a key role in the innate immune response to viral infection and is also involved in the regulation of signal transduction, apoptosis, cell proliferation and differentiation.

Apoptosis is the cellular mechanism of undergoing programmed cell death by the means of many apoptotic proteins (Fig. 5). Apoptosis occurs via a mitochondrial-dependent intrinsic pathway or a death receptor-mediated extrinsic pathway (29, 30, 31). Curcumin causes dose-dependent apoptosis and DNA fragmentation of Caki cells, which is preceded by the sequential dephosphorylation of Akt, down-regulation of the anti-apoptotic Bcl-2, Bcl-xL and IAP proteins, release of cytochrome c and activation of caspase 3 (20).

**Figure 5.**
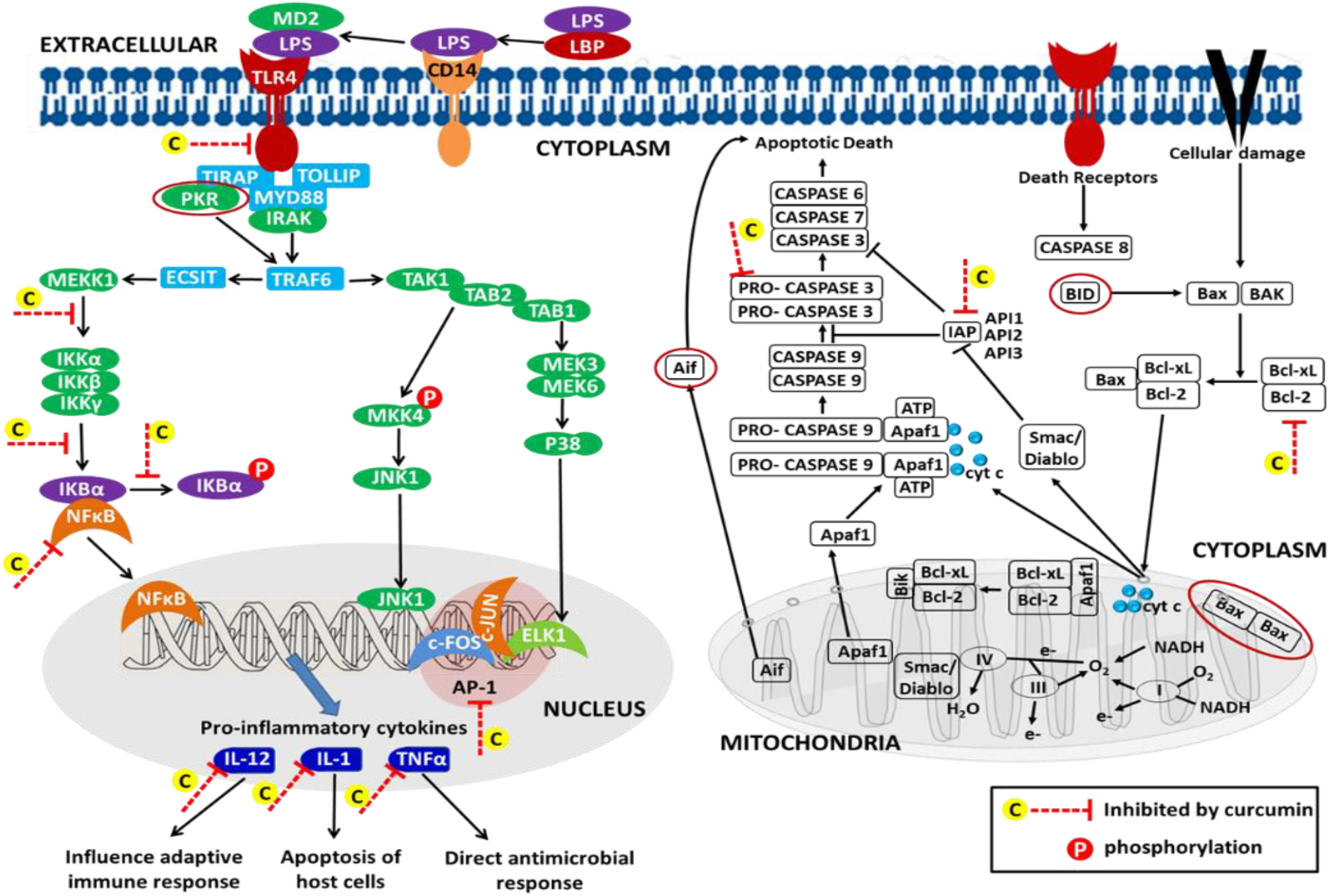
Schematic diagram showing probable mechanism of curcumin action on LPS treated A549 cells. Curcumin (C) inhibits activation of several proteins and the ones marked with dark red oval are identified in this study.

**Figure 6.**
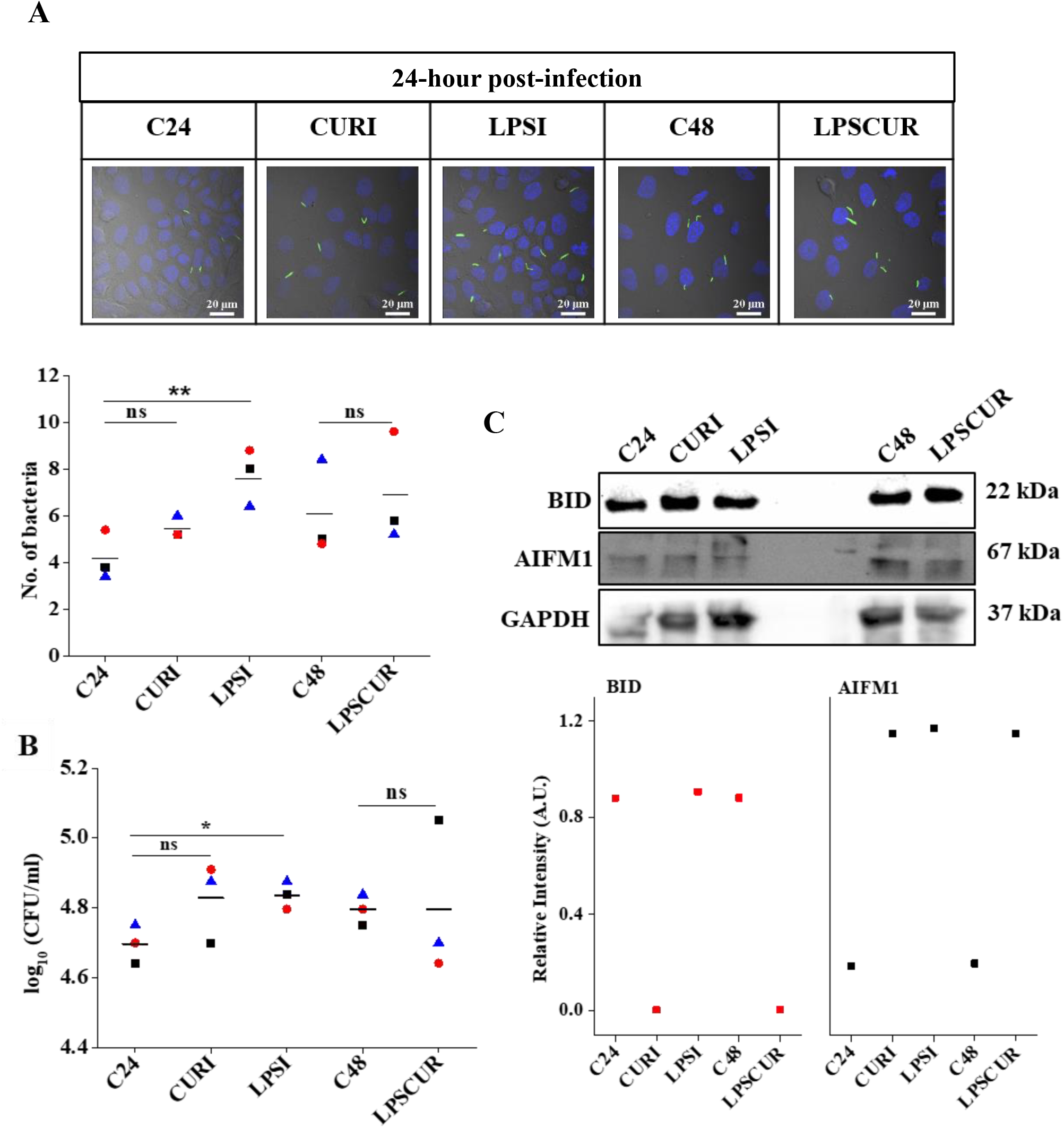
H37Rv-GFP infected A549 cells, treated with curcumin and/or LPS, for 24 hours at multiple of infectivity (1:5) showed successful Mtb internalization and varied bacterial load. (A) Representative confocal microscopy images of cells stained with DAPI (blue, 1 µg/ml) to stain nucleus. Scale: 20 µm and calculation from average of 5 different focal points in triplicates for each experiment. Additional images are available in Figure S9A, S9B and S9C. (B) Colony forming unit (CFU) counts at different conditions. * at 95 % confidence and ** at 99 % confidence. (C) Expression of BID and AIFM1 proteins as probed using Western blot analysis. GAPDH was used as loading control and relative band intensity are presented.

Curcumin treatment down regulated BH3-interacting domain death agonist (BID) suggesting that BID deactivate the anti-apoptotic function of Bcl-2/Bcl-xL and allow the formation of pores on the outer mitochondrial membrane and cytochrome c release from mitochondria (32). In this study, we identified two proteins from the apoptotic pathway. BID, which is a member of the Bcl-2 family of proteins that regulate the permeabilization of the outer mitochondrial membrane (OMM), a critical event during apoptosis showed significant down regulation (log_2_ fold change = −1.31) with curcumin exposure. The other Apoptosis-inducing factor 1(AIFM1-1) which showed up regulation upon simultaneous treatment with LPS and curcumin (log_2_ fold change = 1.32), functions as regulator of apoptosis in a caspase-independent pathway. Diablo homolog identified in curcumin treated cells is reported to release to the cytosol from mitochondrial intermembrane space. It promotes apoptosis by activating caspases in the cytochrome c/Apaf-1/caspase-9 pathway and acts by affecting the inhibitory activity of IAP (inhibitor of apoptosis proteins) (33). Bax was identified in all three experiments with similar abundance (data not shown). Indeed, curcumin has been reported to promote Bax oligomerization, via a caspase-independent mechanism, suggesting that Bax activation and oligomerization are early events during curcumin-induced caspase activation.

Apoptosis-inducing factor is a mitochondrial flavoprotein, which normally resides in the inner mitochondrial membrane, and it possesses an NADH oxidase activity (34). We observed upregulation of AIFM1 level in LPS treated A549 cells that acts as a scavenger of ROS, particularly peroxides (35). Translocation of AIF from the mitochondria to the cytosol as well as to the nucleus exerts caspase-independent apoptosis in a number of model systems (36, 37). AIF can trigger nuclear apoptosis in a caspase-independent manner (38). Curcumin induces AIF translocation from mitochondria to the cytosol and the nucleus (39).

We observed high mitochondrial intensity in A549 cells treated with curcumin. Earlier reports on SHSY5Y cells treated with curcumin (66.3 µM) for 24 hours demonstrated enhanced mitochondrial function (40). These modifications in mitochondrial function could be due to increased mitochondrial ATP, cytochrome oxidase activity, and reduced free radicals and oxidative stress (44). Curcumin treated MCF-7 breast cancer cells showed suppressed cellular oxygen consumption rate, cell mitochondrial membrane potential and ROS generation, and shifting of metabolism from mitochondrial respiration to glycolytic flux (41). These findings strongly suggest that curcumin boosts mitochondrial function to promote cell longevity and its anticancer action. Mice (db/db) treated with curcumin demonstrated diminished mitochondrial dysfunction in mitochondria isolated from the liver and kidneys of these mice (42). Mitochondria of curcumin treated A549 cells might demonstrate enhanced function and needs further investigation.

We extracted the phosphoproteome profile from the identified protein sets in A549 cells treated with curcumin and/or LPS. Different sets of phosphorylated proteins were observed in these experimental samples however; we did not probe the intensity of these molecules in independent samples using antibodies. It is expected that a perturbation, like treatment of curcumin and/or LPS, will induce phosphorylation in cell lines and may show time dependent variations. Interesting findings on LPS treated phosphoproteome analysis in primary macrophages has been demonstrated and a similar study in presence of curcumin would be informative (43). The qualitative data on number of phosphosites that showed certain selectivity to each of the amino acids Serine (S), Threonine (T), Tyrosine (Y) still need further verification and could generate additional information about signalling events that are controlled by phosphorylation of intermediate molecules with curcumin treatment.

We observed successful internalization of H37Rv Mtb strains in all curcumin and/or LPS pre-treated A549 cells by confocal microscopy. Curcumin pretreated A549 cells showed similar bacterial load as controls whereas LPS pretreated cells showed significantly higher Mtb load. Interestingly, A549 cells treated with LPS and CUR for 48 hours, Mtb bacterial load was found to be lower than the control group. It indicates that curcumin contributes in countering the effect of LPS on the A549 cells. Pretreatment of cells with LPS contributes a change in phenotype compromising the clearance of intracellular pathogens. Macrophages chronically exposed to LPS reported to show compromised TNFα production. In response to secondary LPS challenges and treatment with murayl dipeptide partially restores TNFα production, and antimicrobial activity in these cells (44). Animal experiments might provide further useful information on the chemoprotective role of Curcumin. In H37Rv infected mice model, introduction of Curcumin nanoparticles by intraperitoneal route to mice at 100 mg/kg body weight with tuberculosis drugs, improved Mtb clearance by promoting antitubercular immunity (45). In fact, the team also demonstrated that animals receiving curcumin were attaining sterile immunity and protected from reactivation of TB. In a separate study, curcumin inhibited intracellular growth of clinical MDR and laboratory Mtb H37Rv strains in Raw 264.7 cell line (46). It highlights the chemopreventive roles of curcumin in tuberculosis infection.

The alteration in the protein expression levels of BID and AIFM1 is induced by the infection with Mtb. Since, AIFM1 functions as pro-apoptotic factor in a caspase-independent pathway in response to apoptotic stimuli when it is released from the mitochondrion to the nucleus. During infection the increased level of AIFM1 indicates that simultaneous LPS and curcumin treatment is inducing apoptosis in the A549 cells faster than the uninfected cells. BID is a pro-apoptotic protein of the Bcl-2 family that functions for death receptor-mediated apoptosis. BID decreased level in curcumin treated cells and after infection indicate that curcumin is protecting the A549 cell undergoing apoptosis during infection with Mtb.

Our results provide a detailed proteomics insight into the pathways mediated by curcumin to counteract the effect of LPS in A549 cells. We observed several deregulated proteins in curcumin and/or LPS exposed cells involved in TLR4 and apoptotic signaling pathways. BID protein from the apoptotic signaling pathway was inhibited with curcumin treatment. AIFM1 protein, from the apoptotic signaling pathway, showed upregulation with curcumin and LPS treatment. Therefore, BID and AIFM1 proteins can be used as potential targets of curcumin. LPS pre-exposed cells upon infection show higher Mycobacterial load whereas on simultaneous exposure to curcumin and LPS demonstrated lower bacterial load then control. These findings could relate to the mechanism of curcumin as a chemopreventive, anti-oxidative and anti-inflammatory agent. Thus, curcumin could be used as a potential therapeutic agent for the development of novel anti-inflammatory and anticancer drugs.

### Statistical Analysis

Relative intensities of western blot band and mean intensity of fluorescence of mitochondria was analyzed by ImageJ (NIH), Origin (OriginLab, Northampton, MA) and Microsoft Excel 2016 (Microsoft Corp., USA). All data points were plotted as means ± standard errors of the means (SEM). Student’s *t* test and paired t-test was carried out to find group specific differences between data sets. In all tests, P values of <0.05 were considered statistically significant. P values are indicated in figures as follows: *, P < 0.05; **, P < 0.01; and ***, P < 0.001.

## Abbreviations

NSCLC: non-small cell lung carcinoma
SILAC: stable isotope labeling method for cell culture
LPS: Lipopolysaccharide
Mtb: *Mycobacterium tuberculosis*
BSA: bovine serum albumin
PBS: phosphate buffer saline
BID: BH3-interacting domain death agonist
AIFM1: apoptosis-inducing factor protein 1
NCCS: National Centre for Cell Science
DAPI: 4′,6-diamidino-2-phenylindole
TLRs: Toll-like receptors
TNF-α: Tumor necrosis factor α
GAPDH: Glyceraldehyde 3-phosphate dehydrogenase

## Acknowledgements

Supports from core budget of Institute of Genomics and Integrative Biology, New Delhi and International Centre for Genetic Engineering and Biotechnology New Delhi Component are highly acknowledged. We thank Beena Pillai for contributing reagents and materials. Department of Biotechnology supported Tuberculosis Aerosol Challenge Facility at International Centre for Genetic Engineering and Biotechnology is kindly acknowledged.

## Author Contributions

S. Singh and R. Arya designed the project, performed research, contributed new reagents or analytic tools, analyzed data and wrote the paper; R.K. Nanda, R.R. Bargaje, M.K. Das, and R.K. Behera designed the project and prepared the manuscript; S. Akram, H.M. Faruquee performed research, analyzed data and wrote the manuscript; A. Agrawal and R.K. Nanda contributed new reagents, analytic tools and wrote the manuscript.

