## Supplemental Figures for "Anti-inflammatory role of curcumin in Lipopolysaccharide treated A549 cells at global proteome level and on mycobacterial infection"

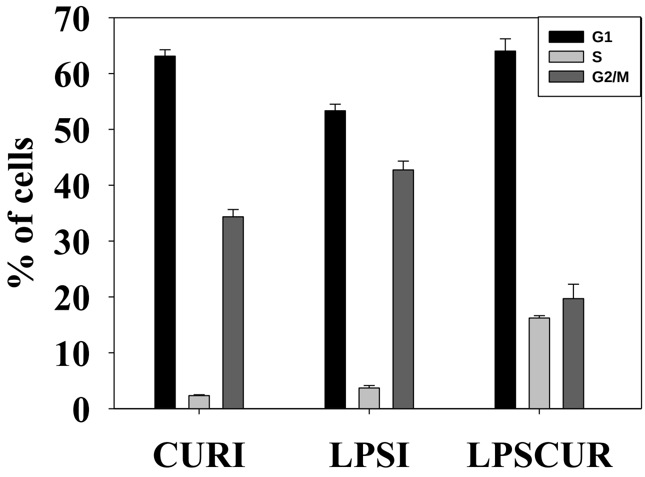


**Figure S1: Effect of curcumin and/or LPS treatment on A549 cell viability** A549 cells were treated with curcumin (10 µM) and/or LPS or 1 µg/ml for the indicated times and after fixation were stained with propidium iodide and Annexin V-FITC. The DNA contents were determined by flow cytometry to calculate percentage of cells present in each phase of the cell cycle (G1, S and G2/M) using Flowing analysis software.

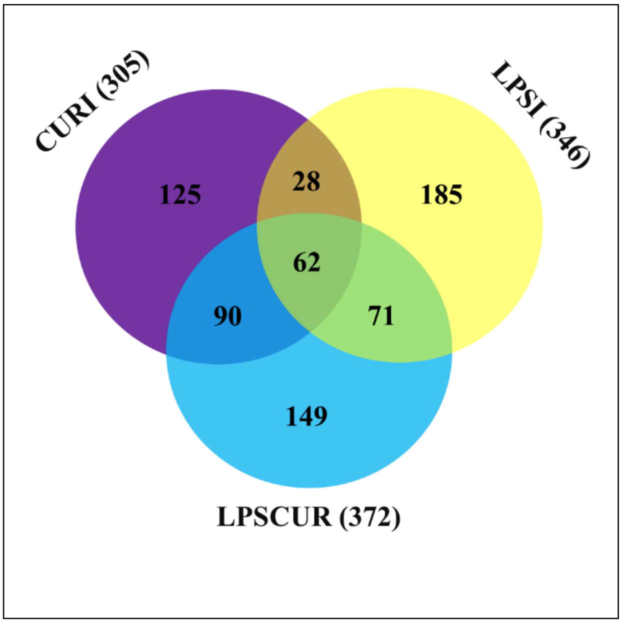


**Figure S2: Total proteins identified in all the three experiments and their distribution betwee curcumin and/or LPS treated conditions.** The proteins showing differential expressions ($\log_2$ fold change≥2) in these experiments were presented in the venn diagram and certain number of proteins are common in all three experiments.

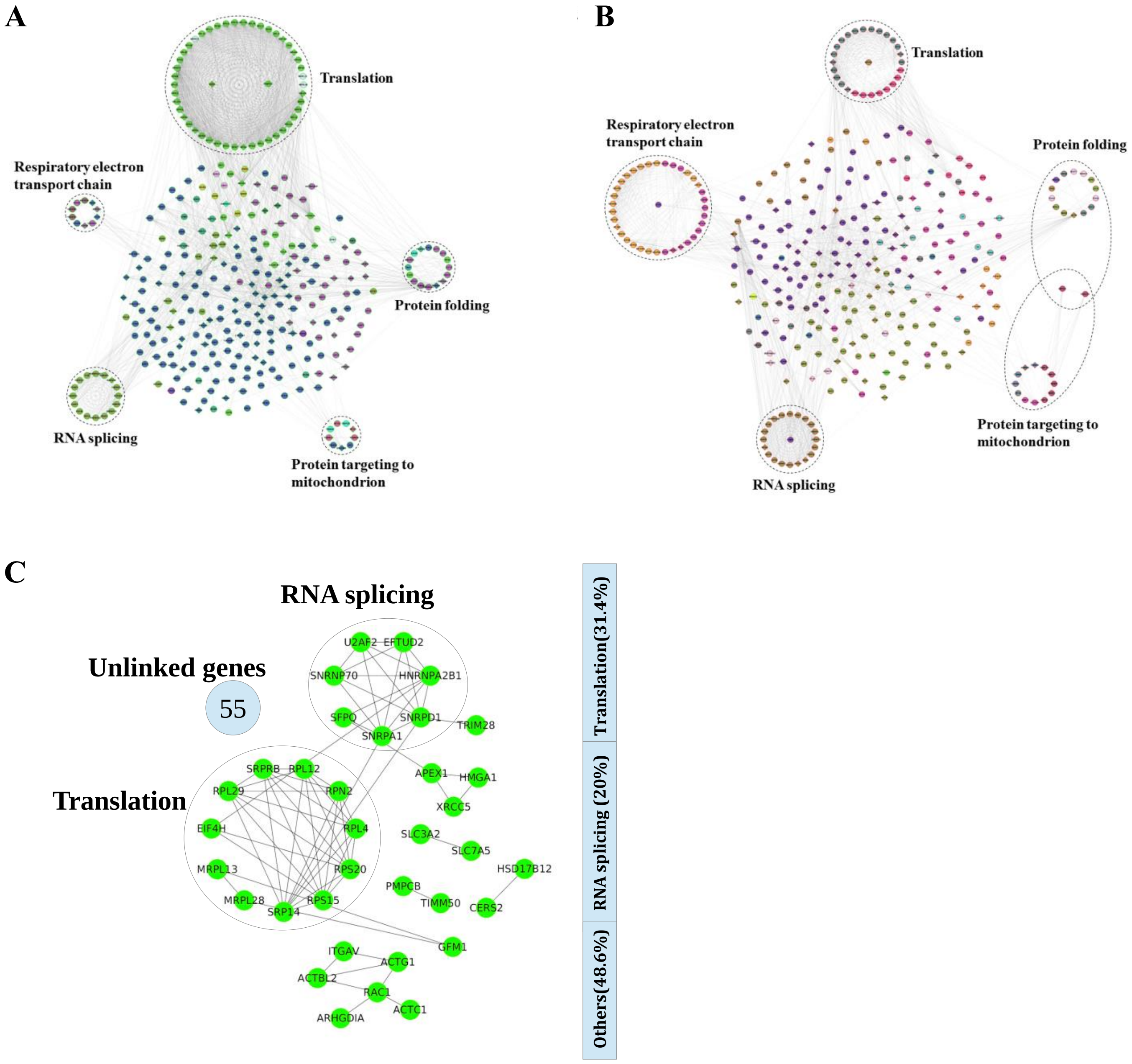


**Figure S3: Treatment of the A549 cells with curcumin and LPS show altered biological activity.** Subnetwork in (A) curcumin treated cells (out of 305 deregulated genes only 257 were mapped to the database with 75 linker genes) and (B) LPs treated cells (out of 346 deregulated genes only 265 were mapped to the database with 65 linker genes). Different colors shows different modules in which the interaction of the genes are highest based on the modularity. The circled maps of the subnetworks involves different biological processes. Linker genes are shown by diamond shaped nodes. (C) Reactome FI network of 90 common deregulated genes/proteins between curcumin and LPS treated experiments. The important biological processes like translation and RNA splicing accounts for 31.4% and 20% respectively. About 48.4% of genes which are not linked to other genes may be involved in some other biological processes.

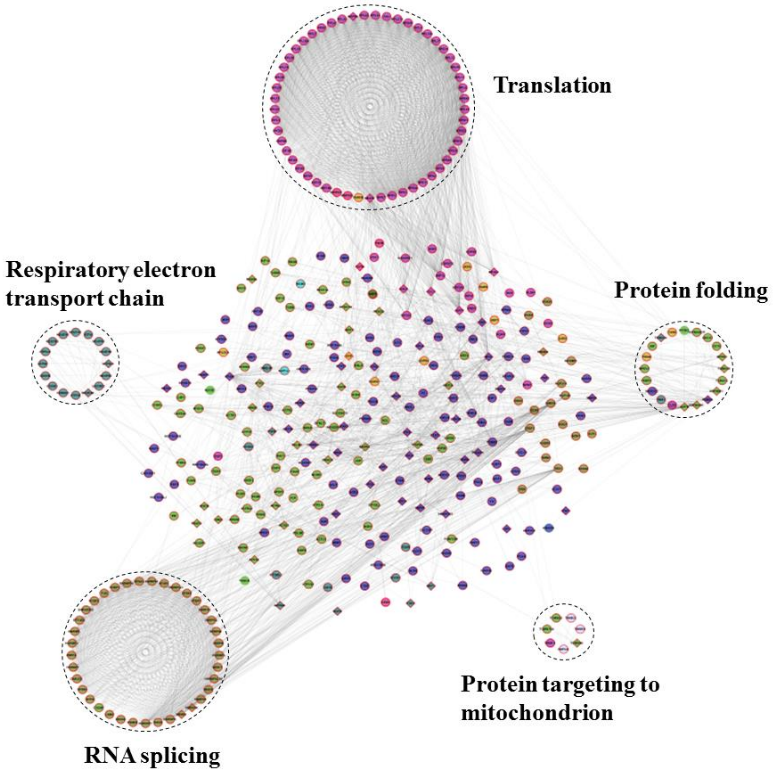


**Figure S4**: **Simultaneous treatment of A549 cells with curcumin and LPS and its effect on the biological pathways**. Subnetwork in LPSCUR (out of 372 deregulated genes only 317 were mapped to the database with 76 linker genes). Functional categorization and the distribution of LPSCUR differentially expressed proteins to extract the information of their known functions.

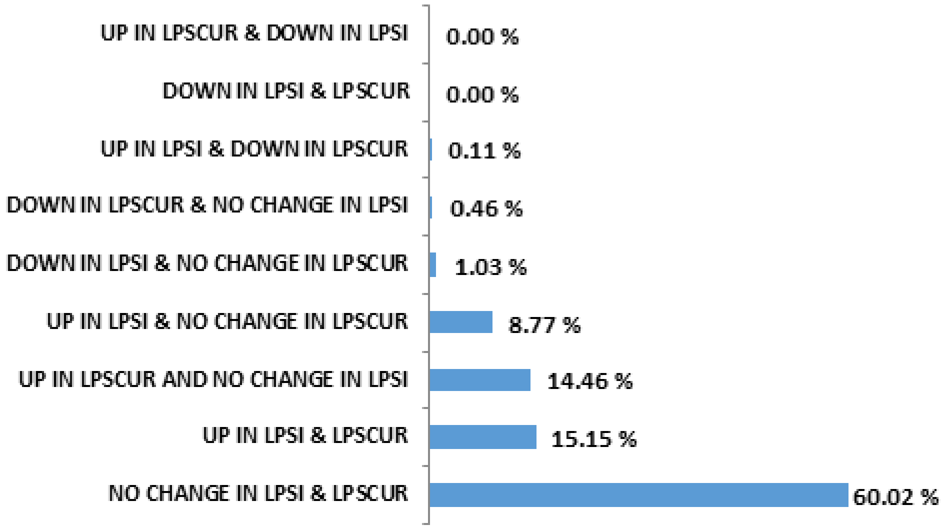


**Figure S5: Percentage distribution of the deregulated proteins in LPS treated and both LPS and curcumin treated cells.** A total of 878 proteins were common in these experiments. The proteins upregulated in LPS treated cells did not show change in both LPS and curcumin treated cells accounts for 8.77%. About 15.15% accounts for upregulation in both the experiments with 60% with no significant change in both experiments.

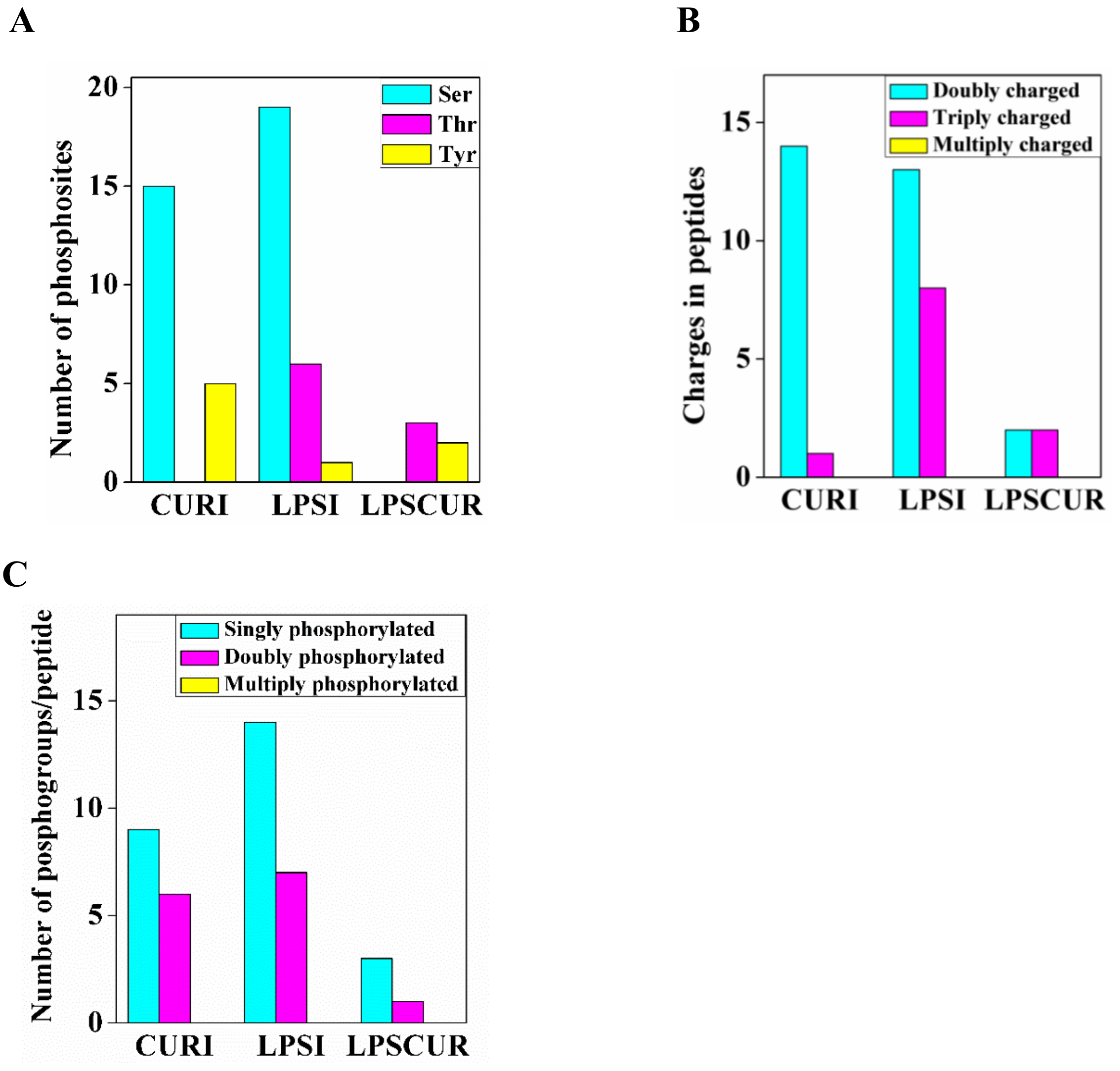


**Figure S6: Alteration in phosphoproteins in curcumin and/or LPS treated cells.** (A) Distribution of the phosphosite localization of class I identified phsphosites. (B) Distribution of the charges in peptides of phosphorylated proteins. (C) Distribution of no. of phosphogroups per peptide.

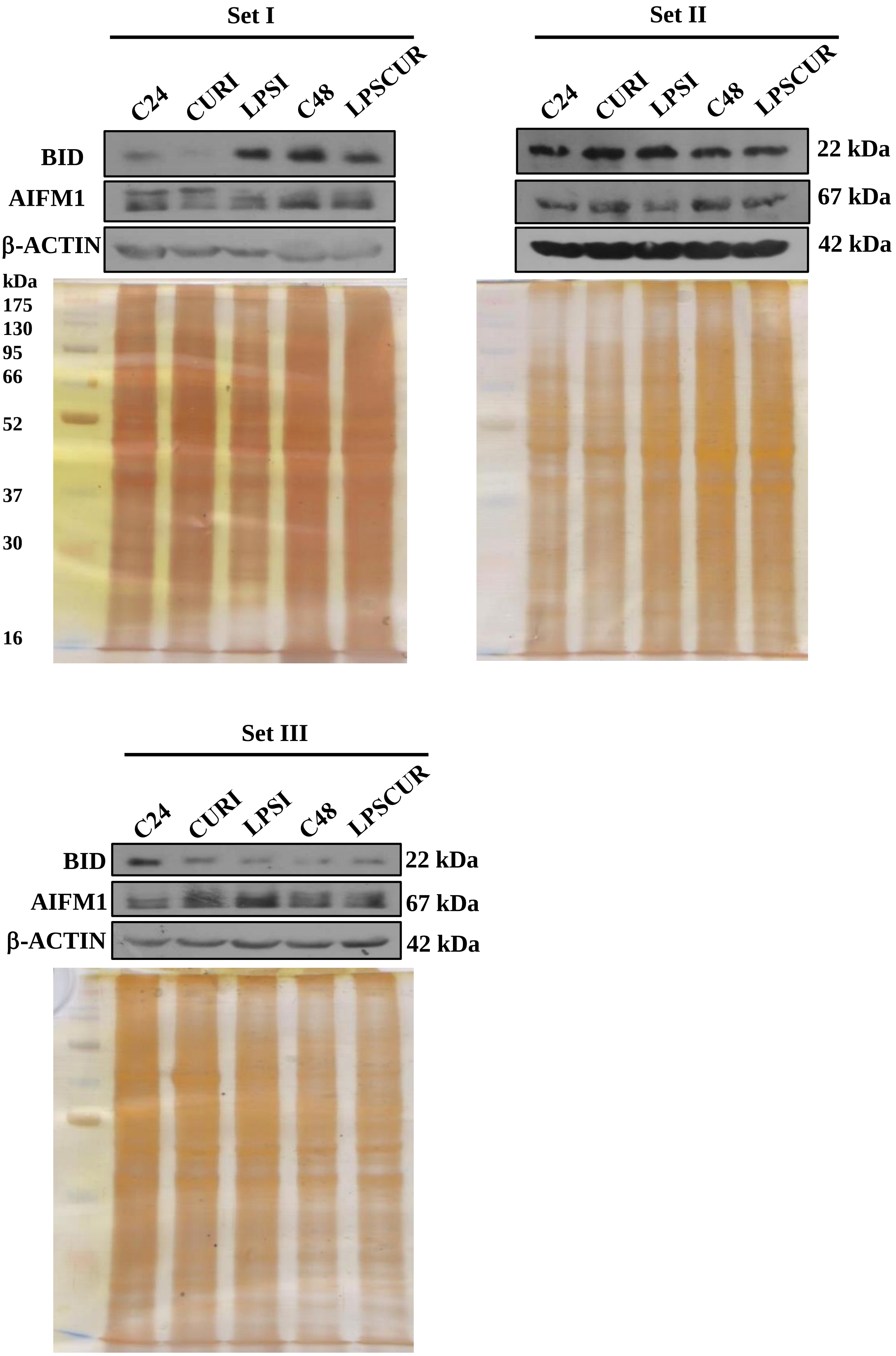


**Figure S7: Western blot analysis of BID and AIFM1 proteins isolated from A549 cells treated with LPS and/or curcumin in independent sample sets.** A549 cells were treated with curcumin (10 µM) or LPS (1 µg/ml) and both for 24 hours and 48 hours respectively showed BID and AIFM1 expression as analysed by Western blot. Silver staining gels are shown for loading control (6 µg). Proteins were transferred to membrane and the blot was cut into 3 parts for probing BID, AIFM1 and β-Actin. Actin was used as loading control and relative band intensity are presented.

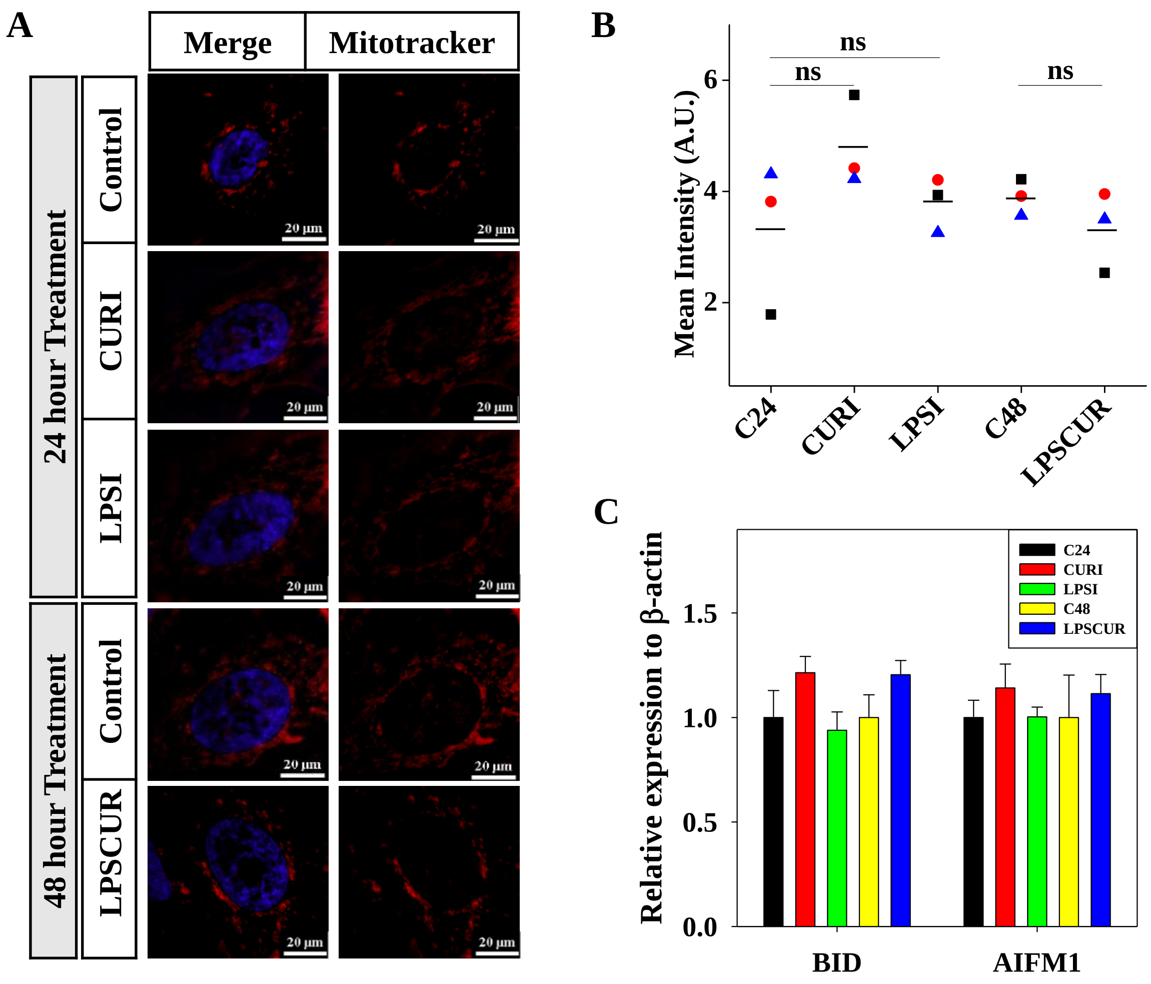


**Figure S8: A549 cells treated with Curcumin (CURI), Lipopolysaccharide (LPSI) or both (LPSCUR) for different incubation periods** (A, B) For N-SIM image analysis: cells were incubated with Mitotracker® red (red, 200 nM) and DAPI (blue, 1 µg/ml) to stain mitochondria and nucleus respectively. Scale- 20 µm; ns = not significant at 95 % confidence. Bars shows standard error of means. (C) Expression of BID and AIFM1 at transcript level using qRT-PCR showed insignificant variation.

### REPLICATE A

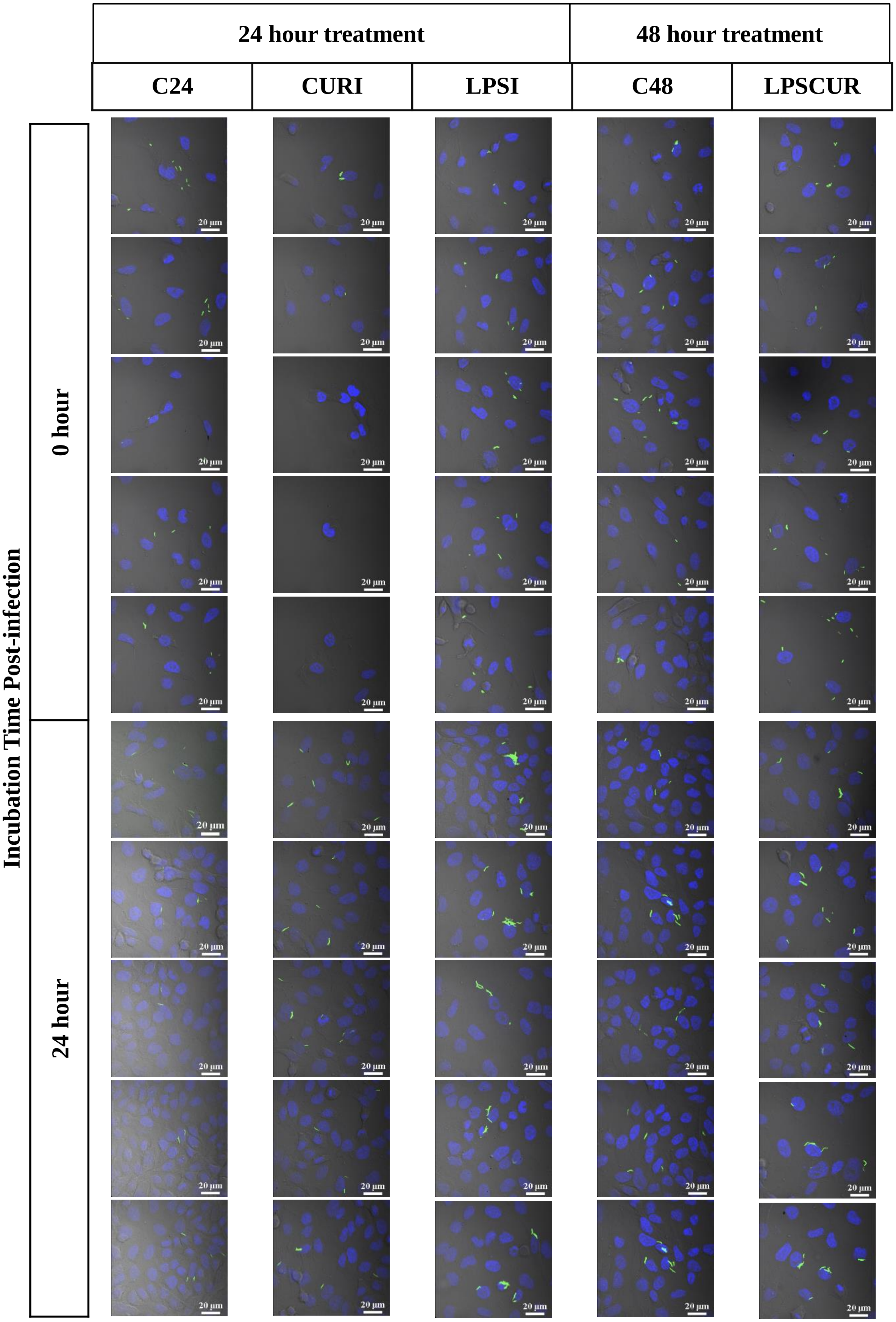


**Figure S9A: Confocal images of H37Rv-GFP infected A549 cells (at multiplicity of infection 1:5) treated with Curcumin (CURI), Lipopolysaccharide (LPSI) or both (LPSCUR).** Cells were incubated with DAPI (blue, 1 µg/ml) to stain nucleus. Scale : 20 µm

**REPLICATE B**

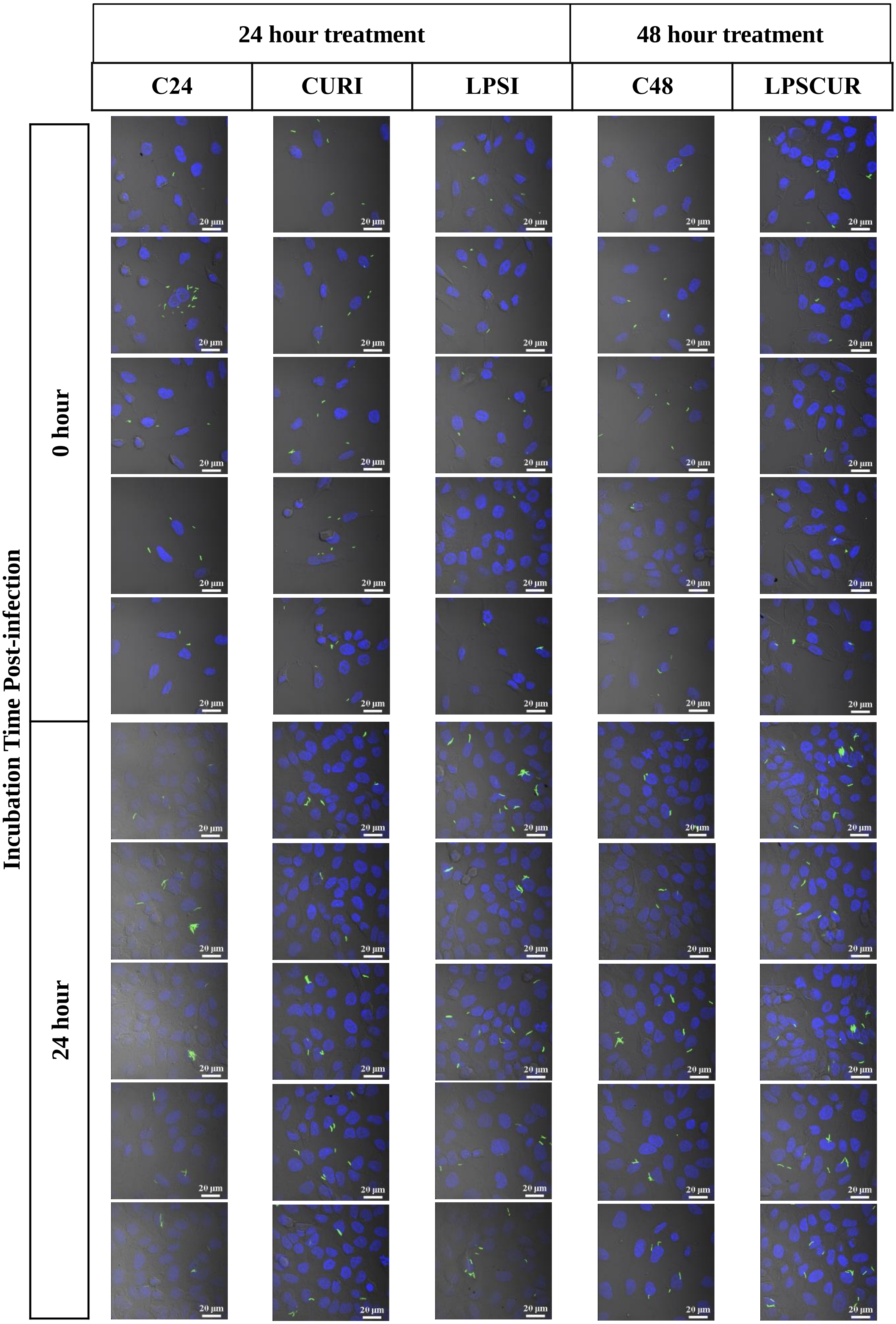


**Figure S9B: Confocal images of H37Rv-GFP infected A549 cells (at multiplicity of infection 1:5) treated with Curcumin (CURI), Lipopolysaccharide (LPSI) or both (LPSCUR).** Cells were incubated with DAPI (blue, 1 µg/ml) to stain nucleus. Scale : 20 µm

### REPLICATE C

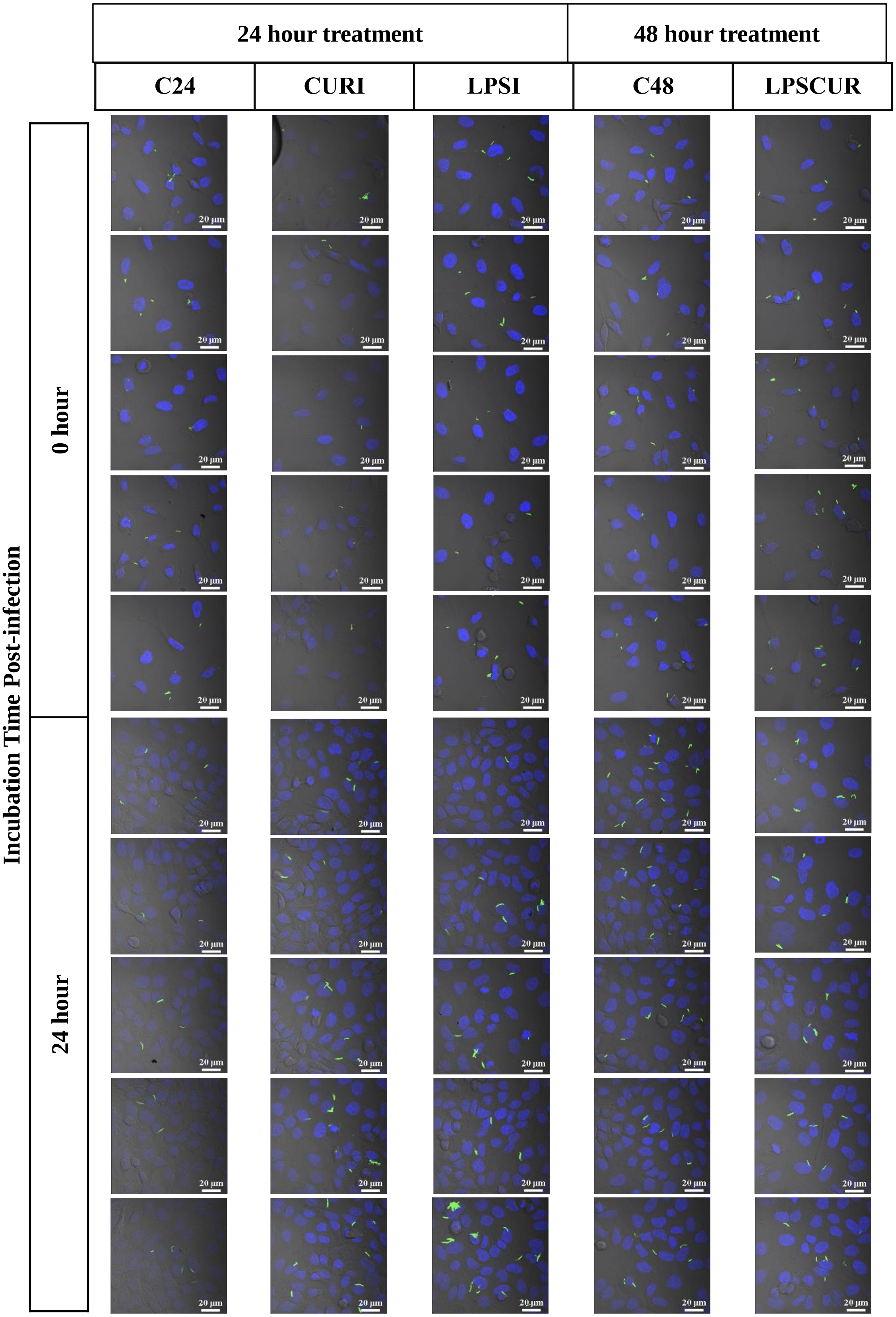


**Figure S9C: Confocal images of H37Rv-GFP infected A549 cells (at multiplicity of infection 1:5) treated with Curcumin (CURI), Lipopolysaccharide (LPSI) or both (LPSCUR).** Cells were incubated with DAPI (blue, 1 µg/ml) to stain nucleus. Scale : 20 µm
