## Supplemental Table S1 for "Anti-inflammatory role of curcumin in Lipopolysaccharide treated A549 cells at global proteome level and on mycobacterial infection"

**Table S1. List of all identified proteins from SILAC experiments in CURI, LPSI and LPSCUR**

| | **Experiments** | | | | | | | **Peptides** | | | **Sequence coverage (%)** | | |
|---|---|---|---|---|---|---|---|---|---|---|---|---|---|
| **S. No.** | **CURI** | **LPSI** | **LPSCUR** | **protein ID** | **Protein names** | **Length** | **Gene** | **CURI** | **LPSI** | **LPSCUR** | **CURI** | **LPSI** | **LPSCUR** |
| 1 | -4.32 | 5.04 | 7.56 | P81605 | Dermcidin (EC 3.4.-.-) (Preproteolysin) [Cleaved into: Survival-promoting peptide; DCD-1] | 110 | DCD | 2 | 5 | 2 | 25.5 | 35.5 | 12.7 |
| 2 | -4.28 | 4.55 | 2.47 | P35908 | Keratin, type II cytoskeletal 2 epidermal (Cytokeratin-2e) (CK-2e) (Epithelial keratin-2e) (Keratin-2 epidermis) (Keratin-2e) (K2e) (Type-II keratin Kb2) | 639 | KRT2 | 16 | 52 | 21 | 37.6 | 82.9 | 47.1 |
| 3 | -4.23 | 0.81 | 1.23 | K7ENV7 | Isochorismatase domain-containing protein 2, mitochondrial (Fragment) | 174 | ISOC2 | 1 | 1 | 1 | 19 | 9.2 | 9.2 |
| 4 | -3.92 | 0.00 | 0.00 | J3KT51 | Hematological and neurological-expressed 1 protein | 104 | HN1 | 1 | 0 | 0 | 24 | 0 | 0 |
| 5 | -3.80 | 0.00 | 3.47 | Q05639 | Elongation factor 1-alpha 2 (EF-1-alpha-2) (Eukaryotic elongation factor 1 A-2) (eEF1A-2) (Statin-S1) | 463 | EEF1A2 | 8 | 4 | 8 | 21 | 8.4 | 21 |

| | | | | | | | | | | | | | |
|---|---|---|---|---|---|---|---|---|---|---|---|---|---|
| 6 | -3.59 | 2.30 | 0.00 | P17096 | High mobility group protein HMG-I/HMG-Y (HMG-I(Y)) (High mobility group AT-hook protein 1) (High mobility group protein A1) (High mobility group protein R) | 107 | HMGA1 | 2 | 1 | 0 | 23.4 | 15 | 0 |
| 7 | -3.46 | 0.00 | 3.47 | A0A087 WY55 | Chromosome 6 open reading frame 55, isoform CRA_b (Vacuolar protein sorting-associated protein VTA1 homolog) | 280 | VTA1 | 1 | 0 | 1 | 5 | 0 | 5 |
| 8 | -3.43 | 2.43 | 4.28 | P08621 | U1 small nuclear ribonucleoprotein 70 kDa (U1 snRNP 70 kDa) (U1-70K) (snRNP70) | 437 | SNRNP70 | 1 | 1 | 2 | 2.5 | 2.5 | 5.5 |
| 9 | -3.39 | 0.00 | 3.13 | H7C3S9 | COP9 signalosome complex subunit 8 (Fragment) | 83 | COPS8 | 2 | 1 | 1 | 27.7 | 12 | 15.7 |
| 10 | -3.36 | 2.68 | 0.00 | P16402 | Histone H1.3 (Histone H1c) (Histone H1s-2) | 221 | HIST1H1D | 6 | 5 | 5 | 22.6 | 22.2 | 14.9 |
| 11 | -3.35 | 1.69 | 1.71 | Q6S8J3 | POTE ankyrin domain family member E (ANKRD26-like family C member 1A) (Prostate, ovary, testis-expressed protein on chromosome 2) (POTE-2) | 1075 | POTEE | 6 | 8 | 4 | 7.7 | 9.7 | 6.3 |
| 12 | -3.29 | 0.00 | 0.00 | P25325 | 3-mercaptopyruvate sulfurtransferase (MST) (EC 2.8.1.2) | 297 | MPST | 1 | 0 | 0 | 9.1 | 0 | 0 |
| 13 | -3.28 | -0.46 | 0.42 | Q99536 | Synaptic vesicle membrane protein VAT-1 homolog (EC 1.-.-.-) | 393 | VAT1 | 1 | 1 | 2 | 3.8 | 3.6 | 7.4 |
| 14 | -3.04 | 0.00 | 2.30 | P05026 | Sodium/potassium-transporting ATPase subunit beta-1 (Sodium/potassium-dependent ATPase subunit beta-1) | 303 | ATP1B1 | 1 | 0 | 1 | 4.7 | 0 | 8.3 |
| 15 | -3.01 | 3.17 | 3.20 | Q562R1 | Beta-actin-like protein 2 (Kappa-actin) | 376 | ACTBL2 | 4 | 4 | 4 | 14.1 | 14.1 | 14.1 |
| 16 | -2.83 | 0.64 | 0.00 | H3BQZ9 | Adenine phosphoribosyltransferase | 153 | APRT | 2 | 1 | 0 | 17 | 8.5 | 0 |

| | | | | | | | | | | | | | |
|---|---|---|---|---|---|---|---|---|---|---|---|---|---|
| 17 | -2.83 | 2.97 | 2.95 | Q9NS69 | Mitochondrial import receptor subunit TOM22 homolog (hTom22) (1C9-2) (Translocase of outer membrane 22 kDa subunit homolog) | 142 | TOMM22 | 3 | 3 | 3 | 43.7 | 39.4 | 43.7 |
| 18 | -2.81 | 0.00 | 1.54 | P43243 | Matrin-3 | 847 | MATR3 | 1 | 0 | 2 | 1.7 | 0 | 3.8 |
| 19 | -2.71 | 0.57 | 0.00 | J3QL05 | Serine/arginine-rich-splicing factor 2 (Fragment) | 130 | SRSF2 | 1 | 1 | 0 | 12.3 | 6.2 | 0 |
| 20 | -2.71 | 0.00 | 1.36 | Q53FA7 | Quinone oxidoreductase PIG3 (EC 1.-.-.-) (Tumor protein p53-inducible protein 3) (p53-induced gene 3 protein) | 332 | TP53I3 | 3 | 0 | 1 | 16.6 | 0 | 4.2 |
| 21 | -2.57 | 2.20 | 2.37 | Q16543 | Hsp90 co-chaperone Cdc37 (Hsp90 chaperone protein kinase-targeting subunit) (p50Cdc37) [Cleaved into: Hsp90 co-chaperone Cdc37, N-terminally processed] | 378 | CDC37 | 1 | 1 | 1 | 5.6 | 5.6 | 5.6 |
| 22 | -2.55 | 0.00 | 0.00 | H7C2I4 | Zinc phosphodiesterase ELAC protein 2 (Fragment) | 286 | ELAC2 | 1 | 0 | 0 | 7.3 | 0 | 0 |
| 23 | -2.54 | 0.00 | 0.00 | Q9Y584 | Mitochondrial import inner membrane translocase subunit Tim22 (Testis-expressed sequence 4) | 194 | TIMM22 | 1 | 0 | 0 | 7.7 | 0 | 0 |
| 24 | -2.49 | 2.03 | 2.04 | P47914 | 60S ribosomal protein L29 (Cell surface heparin-binding protein HIP) | 159 | RPL29 | 2 | 1 | 2 | 14.5 | 9.4 | 14.5 |
| 25 | -2.46 | 0.00 | 1.65 | P00374 | Dihydrofolate reductase (EC 1.5.1.3) | 187 | DHFR | 1 | 0 | 1 | 11.9 | 0 | 11.9 |
| 26 | -2.46 | 0.00 | 2.02 | Q00796 | Sorbitol dehydrogenase (EC 1.1.1.14) (L-iditol 2-dehydrogenase) | 357 | SORD | 2 | 0 | 1 | 35 | 0 | 20 |
| 27 | -2.46 | 1.75 | 0.00 | Q9NRF9 | DNA polymerase epsilon subunit 3 (EC 2.7.7.7) (Arsenic-transactivated protein) (AsTP) (Chromatin accessibility complex 17 kDa protein) (CHRAC-17) (HuCHRAC17) (DNA polymerase II subunit 3) (DNA polymerase epsilon subunit p17) | 147 | POLE3 | 1 | 1 | 0 | 10.9 | 10.9 | 0 |

| | | | | | | | | | | | | | |
|---|---|---|---|---|---|---|---|---|---|---|---|---|---|
| 28 | -2.44 | 1.94 | 1.97 | K7EM56 | 40S ribosomal protein S15 | 112 | RPS15 | 4 | 1 | 3 | 52.7 | 10.7 | 28.6 |
| 29 | -2.43 | 0.97 | 2.02 | Q16531 | DNA damage-binding protein 1 (DDB p127 subunit) (DNA damage-binding protein a) (DDBa) (Damage-specific DNA-binding protein 1) (HBV X-associated protein 1) (XAP-1) (UV-damaged DNA-binding factor) (UV-damaged DNA-binding protein 1) (UV-DDB 1) (XPE-binding factor) (XPE-BF) (Xeroderma pigmentosum group E-complementing protein) (XPCe) | 1140 | DDB1 | 1 | 2 | 2 | 0.9 | 2 | 3 |
| 30 | -2.41 | 0.84 | 1.28 | P24666 | Low molecular weight phosphotyrosine protein phosphatase (LMW-PTP) (LMW-PTPase) (EC 3.1.3.48) (Adipocyte acid phosphatase) (Low molecular weight cytosolic acid phosphatase) (EC 3.1.3.2) (Red cell acid phosphatase 1) | 158 | ACP1 | 1 | 4 | 1 | 11.4 | 41.8 | 8.2 |
| 31 | -2.40 | -0.04 | 1.58 | Q00839 | Heterogeneous nuclear ribonucleoprotein U (hnRNP U) (Scaffold attachment factor A) (SAF-A) (p120) (pp120) | 825 | HNRNPU | 7 | 3 | 7 | 11.3 | 3.8 | 9.2 |
| 32 | -2.38 | 0.00 | 2.20 | Q8NBX0 | Saccharopine dehydrogenase-like oxidoreductase (EC 1.-.-.-) | 429 | SCCPDH | 1 | 0 | 1 | 3.3 | 0 | 3.3 |
| 33 | -2.36 | 0.97 | -0.18 | P33991 | DNA replication licensing factor MCM4 (EC 3.6.4.12) (CDC21 homolog) (P1-CDC21) | 863 | MCM4 | 2 | 3 | 1 | 4.5 | 5.7 | 1.5 |
| 34 | -2.34 | -0.07 | 0.86 | Q9UMY4 | Sorting nexin-12 | 172 | SNX12 | 2 | 4 | 1 | 24.7 | 20.4 | 4.3 |
| 35 | -2.34 | 1.21 | 2.03 | J3KTF8 | Rho GDP-dissociation inhibitor 1 (Fragment) | 193 | ARHGDIA | 3 | 1 | 1 | 31.6 | 7.8 | 16.6 |
| 36 | -2.28 | 0.00 | 0.47 | Q6PUV4 | Complexin-2 (Complexin II) (CPX II) (Synaphin-1) | 134 | CPLX2 | 1 | 0 | 1 | 8.2 | 0 | 4.5 |

| | | | | | | | | | | | | | |
|---|---|---|---|---|---|---|---|---|---|---|---|---|---|
| 37 | -2.27 | 0.00 | 0.00 | Q16537 | Serine/threonine-protein phosphatase 2A 56 kDa regulatory subunit epsilon isoform (PP2A B subunit isoform B'-epsilon) (PP2A B subunit isoform B56-epsilon) (PP2A B subunit isoform PR61-epsilon) (PP2A B subunit isoform R5-epsilon) | 467 | PPP2R5E | 1 | 0 | 0 | 3.2 | 0 | 0 |
| 38 | -2.25 | 0.93 | 2.29 | E9PES6 | High mobility group protein B3 (Fragment) | 153 | HMGB3 | 1 | 2 | 3 | 9.2 | 19 | 28.1 |
| 39 | -2.24 | 1.20 | 1.13 | P07954 | Fumarate hydratase, mitochondrial (Fumarase) (EC 4.2.1.2) | 510 | FH | 3 | 3 | 3 | 9.9 | 7.5 | 10.1 |
| 40 | -2.22 | 0.00 | 1.97 | C9IZ80 | Basic leucine zipper and W2 domain-containing protein 1 (Fragment) | 294 | BZW1 | 1 | 0 | 2 | 8.8 | 0 | 12.9 |
| 41 | -2.18 | 0.31 | 1.85 | F8VSD4 | Ubiquitin-conjugating enzyme E2 N | 105 | UBE2N | 6 | 4 | 4 | 63.8 | 42.9 | 38.1 |
| 42 | -2.17 | 1.52 | 1.57 | P63000 | Ras-related C3 botulinum toxin substrate 1 (Cell migration-inducing gene 5 protein) (Ras-like protein TC25) (p21-Rac1) | 192 | RAC1 | 4 | 2 | 3 | 25.5 | 13 | 19.8 |
| 43 | -2.17 | 0.00 | 1.95 | P40818 | Ubiquitin carboxyl-terminal hydrolase 8 (EC 3.4.19.12) (Deubiquitinating enzyme 8) (Ubiquitin isopeptidase Y) (hUBPy) (Ubiquitin thioesterase 8) (Ubiquitin-specific-processing protease 8) | 1118 | USP8 | 1 | 0 | 1 | 1 | 0 | 1 |
| 44 | -2.17 | 1.54 | 0.00 | P08559 | Pyruvate dehydrogenase E1 component subunit alpha, somatic form, mitochondrial (EC 1.2.4.1) (PDHE1-A type I) | 390 | PDHA1 | 1 | 2 | 0 | 3.6 | 6.7 | 0 |
| 45 | -2.14 | -0.79 | 1.80 | P60983 | Glia maturation factor beta (GMF-beta) | 142 | GMFB | 1 | 1 | 1 | 16.2 | 6.3 | 16.2 |

| | | | | | | | | | | | | | |
|---|---|---|---|---|---|---|---|---|---|---|---|---|---|
| 46 | -2.14 | 0.00 | 0.44 | O60762 | Dolichol-phosphate mannosyltransferase subunit 1 (EC 2.4.1.83) (Dolichol-phosphate mannose synthase subunit 1) (DPM synthase subunit 1) (Dolichyl-phosphate beta-D-mannosyltransferase subunit 1) (Mannose-P-dolichol synthase subunit 1) (MPD synthase subunit 1) | 260 | DPM1 | 2 | 0 | 1 | 8.8 | 0 | 4.2 |
| 47 | -2.13 | 0.00 | 0.00 | H7C2Y1 | E3 ubiquitin-protein ligase TRIP12 (Fragment) | 187 | TRIP12 | 1 | 0 | 0 | 5.9 | 0 | 0 |
| 48 | -2.11 | 0.29 | 0.00 | B1AH58 | Intraflagellar transport protein 27 homolog (Fragment) | 116 | IFT27 | 1 | 1 | 0 | 18.1 | 6.9 | 0 |
| 49 | -2.10 | 1.34 | 0.00 | A2IDC6 | 39S ribosomal protein L28, mitochondrial (Fragment) | 240 | MRPL28 | 1 | 1 | 0 | 5 | 2.9 | 0 |
| 50 | -2.09 | 1.20 | 1.28 | Q9Y320 | Thioredoxin-related transmembrane protein 2 (Cell proliferation-inducing gene 26 protein) (Thioredoxin domain-containing protein 14) | 296 | TMX2 | 1 | 2 | 1 | 5.8 | 10.5 | 4.7 |
| 51 | -2.09 | 0.00 | 0.00 | Q9NX08 | COMM domain-containing protein 8 | 183 | COMMD8 | 1 | 0 | 0 | 6.6 | 0 | 0 |
| 52 | -2.07 | 0.00 | 0.00 | C9J306 | 2-hydroxyacyl-CoA lyase 1 (Fragment) | 244 | HACL1 | 1 | 0 | 0 | 4.1 | 0 | 0 |
| 53 | -2.07 | 0.00 | 0.98 | D6R9A6 | High mobility group protein B2 (Fragment) | 134 | HMGB2 | 2 | 0 | 2 | 23.1 | 0 | 17.2 |
| 54 | -2.05 | 0.00 | 0.56 | Q9UHX1 | Poly(U)-binding-splicing factor PUF60 (60 kDa poly(U)-binding-splicing factor) (FUSE-binding protein-interacting repressor) (FBP-interacting repressor) (Ro-binding protein 1) (RoBP1) (Siah-binding protein 1) (Siah-BP1) | 559 | PUF60 | 2 | 0 | 2 | 8.6 | 0 | 6.8 |
| 55 | -2.04 | 0.32 | 0.70 | P18669 | Phosphoglycerate mutase 1 (EC 3.1.3.13) (EC 5.4.2.11) (EC 5.4.2.4) (BPG-dependent PGAM 1) (Phosphoglycerate mutase isozyme B) (PGAM-B) | 254 | PGAM1 | 10 | 5 | 6 | 50 | 31.1 | 34.6 |

| | | | | | | | | | | | | | |
|---|---|---|---|---|---|---|---|---|---|---|---|---|---|
| 56 | -2.02 | 0.00 | 0.00 | C9JQQ5 | Atlastin-2 (Fragment) | 175 | ATL2 | 1 | 0 | 0 | 8.6 | 0 | 0 |
| 57 | -2.02 | 0.00 | 0.00 | Q9UN86 | Ras GTPase-activating protein-binding protein 2 (G3BP-2) (GAP SH3 domain-binding protein 2) | 482 | G3BP2 | 1 | 0 | 0 | 3.8 | 0 | 0 |
| 58 | -2.00 | 0.00 | 2.05 | P53618 | Coatomer subunit beta (Beta-coat protein) (Beta-COP) | 953 | COPB1 | 4 | 0 | 5 | 9.2 | 0 | 9.8 |
| 59 | -1.98 | 1.80 | 0.00 | C9JAZ1 | Metaxin-2 (Fragment) | 229 | MTX2 | 1 | 2 | 0 | 9.6 | 16.6 | 0 |
| 60 | -1.97 | 1.93 | 1.12 | Q15056 | Eukaryotic translation initiation factor 4H (eIF-4H) (Williams-Beuren syndrome chromosomal region 1 protein) | 248 | EIF4H | 1 | 2 | 3 | 9.2 | 21.9 | 28.1 |
| 61 | -1.97 | 0.00 | 2.14 | A0A087 WZX2 | NADH dehydrogenase [ubiquinone] 1 beta subcomplex subunit 6 | 97 | NDUFB6 | 1 | 0 | 1 | 18.6 | 0 | 18.6 |
| 62 | -1.96 | 1.73 | 2.17 | B3KUB4 | Carbonic anhydrase 12 (Carbonic anhydrase XII, isoform CRA_d) (cDNA FLJ39526 fis, clone PUAEN2003018, highly similar to CARBONIC ANHYDRASE XII (EC 4.2.1.1)) | 283 | CA12 | 4 | 2 | 3 | 23.3 | 9.5 | 13.4 |
| 63 | -1.96 | 0.00 | 0.00 | P52907 | F-actin-capping protein subunit alpha-1 (CapZ alpha-1) | 286 | CAPZA1 | 1 | 0 | 0 | 5.2 | 0 | 0 |
| 64 | -1.95 | 1.99 | 1.94 | Q14764 | Major vault protein (MVP) (Lung resistance-related protein) | 893 | MVP | 3 | 1 | 3 | 4.3 | 1.8 | 4.8 |
| 65 | -1.94 | 1.16 | 1.47 | P22626 | Heterogeneous nuclear ribonucleoproteins A2/B1 (hnRNP A2/B1) | 353 | HNRNPA2B 1 | 4 | 5 | 6 | 19.3 | 16.7 | 14.7 |
| 66 | -1.92 | 0.00 | 1.79 | A0A087 WTP3 | Far upstream element-binding protein 2 | 711 | KHSRP | 2 | 0 | 1 | 6 | 0 | 1.5 |
| 67 | -1.91 | 0.00 | 1.45 | O00625 | Pirin (EC 1.13.11.24) (Probable quercetin 2,3-dioxygenase PIR) (Probable quercetinase) | 290 | PIR | 2 | 0 | 4 | 11 | 0 | 18.3 |
| 68 | -1.90 | 1.18 | 1.83 | O75439 | Mitochondrial-processing peptidase subunit beta (EC 3.4.24.64) (Beta-MPP) (P-52) | 489 | PMPCB | 1 | 1 | 1 | 2.2 | 2.2 | 2.2 |

| | | | | | | | | | | | | | |
|---|---|---|---|---|---|---|---|---|---|---|---|---|---|
| 69 | -1.90 | 0.00 | 0.47 | P49915 | GMP synthase [glutamine-hydrolyzing] (EC 6.3.5.2) (GMP synthetase) (Glutamine amidotransferase) | 693 | GMPS | 2 | 0 | 3 | 3.9 | 0 | 5.7 |
| 70 | -1.90 | 0.78 | 1.08 | P52895 | Aldo-keto reductase family 1 member C2 (EC 1.-.-.-) (3-alpha-HSD3) (Chlordecone reductase homolog HAKRD) (Dihydrodiol dehydrogenase 2) (DD-2) (DD2) (Dihydrodiol dehydrogenase/bile acid-binding protein) (DD/BABP) (Trans-1,2-dihydrobenzene-1,2-diol dehydrogenase) (EC 1.3.1.20) (Type III 3-alpha-hydroxysteroid dehydrogenase) (EC 1.1.1.357) | 323 | AKR1C2 | 8 | 10 | 10 | 39.6 | 49.2 | 49.2 |
| 71 | -1.89 | 0.00 | 2.47 | P11498 | Pyruvate carboxylase, mitochondrial (EC 6.4.1.1) (Pyruvic carboxylase) (PCB) | 1178 | PC | 2 | 0 | 1 | 2.2 | 0 | 1.1 |
| 72 | -1.88 | 0.00 | 0.00 | C9JYQ9 | 60S ribosomal protein L22-like 1 | 121 | RPL22L1 | 1 | 0 | 0 | 19.8 | 0 | 0 |
| 73 | -1.88 | 0.00 | 1.25 | O00159 | Unconventional myosin-Ic (Myosin I beta) (MMI-beta) (MMIb) | 1063 | MYO1C | 1 | 0 | 2 | 1.8 | 0 | 2.9 |
| 74 | -1.87 | 0.23 | 0.39 | Q01105 | Protein SET (HLA-DR-associated protein II) (Inhibitor of granzyme A-activated DNase) (IGAAD) (PHAPII) (Phosphatase 2A inhibitor I2PP2A) (I-2PP2A) (Template-activating factor I) (TAF-I) | 290 | SET | 2 | 5 | 2 | 7.9 | 30.2 | 7.9 |
| 75 | -1.87 | 1.28 | 1.97 | Q9BSJ8 | Extended synaptotagmin-1 (E-Syt1) (Membrane-bound C2 domain-containing protein) | 1104 | ESYT1 | 2 | 4 | 1 | 2.9 | 4.9 | 1.4 |

| | | | | | | | | | | | | | |
|---|---|---|---|---|---|---|---|---|---|---|---|---|---|
| 76 | -1.87 | 0.56 | 1.34 | P30041 | Peroxiredoxin-6 (EC 1.11.1.15) (1-Cys peroxiredoxin) (1-Cys PRX) (24 kDa protein) (Acidic calcium-independent phospholipase A2) (aiPLA2) (EC 3.1.1.-) (Antioxidant protein 2) (Liver 2D page spot 40) (Non-selenium glutathione peroxidase) (NSGPx) (EC 1.11.1.9) (Red blood cells page spot 12) | 224 | PRDX6 | 8 | 9 | 9 | 41.5 | 51.3 | 33.9 |
| 77 | -1.86 | 0.70 | 2.05 | P62861 | 40S ribosomal protein S30 | 59 | FAU | 1 | 1 | 1 | 16.9 | 16.9 | 16.9 |
| 78 | -1.86 | 1.77 | 1.64 | Q14696 | LDLR chaperone MESD (Mesoderm development candidate 2) (Mesoderm development protein) (Renal carcinoma antigen NY-REN-61) | 234 | MESDC2 | 1 | 2 | 1 | 4.3 | 9 | 4.7 |
| 79 | -1.85 | 0.00 | 0.00 | H0YN84 | Serine/threonine-protein phosphatase 2A 56 kDa regulatory subunit gamma isoform (Fragment) | 89 | PPP2R5C | 1 | 0 | 0 | 12.4 | 0 | 0 |
| 80 | -1.85 | 0.00 | 2.33 | O76031 | ATP-dependent Clp protease ATP-binding subunit clpX-like, mitochondrial | 633 | CLPX | 1 | 0 | 1 | 2.1 | 0 | 2.1 |
| 81 | -1.85 | 0.00 | 0.00 | O60884 | DnaJ homolog subfamily A member 2 (Cell cycle progression restoration gene 3 protein) (Dnj3) (Dj3) (HIRA-interacting protein 4) (Renal carcinoma antigen NY-REN-14) | 412 | DNAJA2 | 1 | 0 | 0 | 3.6 | 0 | 0 |
| 82 | -1.82 | 1.97 | 1.71 | P51970 | NADH dehydrogenase [ubiquinone] 1 alpha subcomplex subunit 8 (Complex I-19kD) (CI-19kD) (Complex I-PGIV) (CI-PGIV) (NADH-ubiquinone oxidoreductase 19 kDa subunit) | 172 | NDUFA8 | 2 | 3 | 1 | 22.7 | 27.3 | 10.5 |
| 83 | -1.82 | 0.72 | 1.11 | Q8TEX9 | Importin-4 (Imp4) (Importin-4b) (Imp4b) (Ran-binding protein 4) (RanBP4) | 1081 | IPO4 | 4 | 3 | 2 | 5.5 | 3.1 | 2 |
| 84 | -1.80 | 1.89 | 1.70 | Q5SZE2 | Ceramide synthase 2 (Fragment) | 128 | CERS2 | 1 | 1 | 1 | 12.5 | 12.5 | 12.5 |

| | | | | | | | | | | | | | |
|---|---|---|---|---|---|---|---|---|---|---|---|---|---|
| 85 | -1.80 | 1.34 | 2.08 | J3KNF8 | Cytochrome b5 type B (Cytochrome b5 type B (Outer mitochondrial membrane), isoform CRA_a) | 150 | CYB5B | 2 | 4 | 2 | 35.3 | 53.3 | 35.3 |
| 86 | -1.79 | 1.30 | 2.00 | P07099 | Epoxide hydrolase 1 (EC 3.3.2.9) (Epoxide hydratase) (Microsomal epoxide hydrolase) | 455 | EPHX1 | 6 | 5 | 9 | 24 | 14.7 | 35.2 |
| 87 | -1.79 | 1.96 | 2.08 | P10606 | Cytochrome c oxidase subunit 5B, mitochondrial (Cytochrome c oxidase polypeptide Vb) | 129 | COX5B | 2 | 4 | 2 | 18.6 | 24.8 | 17.8 |
| 88 | -1.79 | 0.00 | 1.13 | F5GZ49 | Glycolipid transfer protein (Glycolipid transfer protein isoform 1) | 67 | GLTP | 1 | 0 | 1 | 22.4 | 0 | 22.4 |
| 89 | -1.76 | 0.00 | 1.32 | Q9Y2V2 | Calcium-regulated heat stable protein 1 (Calcium-regulated heat-stable protein of 24 kDa) (CRHSP-24) | 147 | CARHSP1 | 2 | 0 | 2 | 35.4 | 0 | 35.4 |
| 90 | -1.76 | -0.05 | 1.04 | P62899 | 60S ribosomal protein L31 | 125 | RPL31 | 3 | 4 | 1 | 24.8 | 31.2 | 6.4 |
| 91 | -1.75 | 0.00 | 2.30 | Q5VWN6 | Protein FAM208B | 2430 | FAM208B | 1 | 0 | 1 | 0.4 | 0 | 0.4 |
| 92 | -1.75 | 3.47 | 1.29 | K7ESE8 | Bleomycin hydrolase (Fragment) | 231 | BLMH | 1 | 1 | 1 | 4.8 | 4.8 | 4.8 |
| 93 | -1.74 | 1.12 | 1.35 | Q9Y3E5 | Peptidyl-tRNA hydrolase 2, mitochondrial (PTH 2) (EC 3.1.1.29) (Bcl-2 inhibitor of transcription 1) | 179 | PTRH2 | 3 | 5 | 2 | 31.8 | 44.7 | 21.2 |
| 94 | -1.73 | 1.73 | 0.00 | Q8N766 | ER membrane protein complex subunit 1 | 993 | EMC1 | 1 | 2 | 0 | 1.6 | 3.1 | 0 |
| 95 | -1.72 | 2.20 | 1.71 | P62314 | Small nuclear ribonucleoprotein Sm D1 (Sm-D1) (Sm-D autoantigen) (snRNP core protein D1) | 119 | SNRPD1 | 2 | 2 | 2 | 26.1 | 26.1 | 26.1 |
| 96 | -1.70 | 2.07 | 1.64 | Q09666 | Neuroblast differentiation-associated protein AHNAK (Desmoyokin) | 5890 | AHNAK | 7 | 22 | 17 | 4.2 | 15.3 | 10.6 |

| | | | | | | | | | | | | | |
|---|---|---|---|---|---|---|---|---|---|---|---|---|---|
| 97 | -1.68 | 0.75 | 1.12 | P35268 | 60S ribosomal protein L22 (EBER-associated protein) (EAP) (Epstein-Barr virus small RNA-associated protein) (Heparin-binding protein HBp15) | 128 | RPL22 | 1 | 1 | 1 | 20.3 | 8.6 | 8.6 |
| 98 | -1.68 | 0.13 | 1.48 | E1CEI4 | Glutamate--cysteine ligase catalytic subunit (Glutamate-cysteine ligase delta4 alternative splicing variant) | 599 | GCLC | 1 | 1 | 2 | 1.5 | 1.5 | 4.5 |
| 99 | -1.68 | 0.00 | 1.27 | Q15233 | Non-POU domain-containing octamer-binding protein (NonO protein) (54 kDa nuclear RNA- and DNA-binding protein) (55 kDa nuclear protein) (DNA-binding p52/p100 complex, 52 kDa subunit) (NMT55) (p54(nrb)) (p54nrb) | 471 | NONO | 2 | 0 | 3 | 7.9 | 0 | 11.5 |
| 100 | -1.68 | 6.59 | 0.69 | P21399 | Cytoplasmic aconitate hydratase (Aconitase) (EC 4.2.1.3) (Citrate hydro-lyase) (Ferritin repressor protein) (Iron regulatory protein 1) (IRP1) (Iron-responsive element-binding protein 1) (IRE-BP 1) | 889 | ACO1 | 2 | 1 | 2 | 4.3 | 0.9 | 3.7 |
| 101 | -1.67 | 0.44 | 1.96 | E9PLL6 | 60S ribosomal protein L27a | 108 | RPL27A | 2 | 2 | 1 | 23.1 | 18.5 | 11.1 |
| 102 | -1.67 | 0.00 | 2.52 | Q6PI78 | Transmembrane protein 65 | 240 | TMEM65 | 1 | 0 | 1 | 4.6 | 0 | 4.6 |
| 103 | -1.66 | 1.65 | 0.00 | Q9Y2B0 | Protein canopy homolog 2 (MIR-interacting saposin-like protein) (Putative secreted protein Zsig9) (Transmembrane protein 4) | 182 | CNPY2 | 1 | 4 | 0 | 8.8 | 31.3 | 0 |
| 104 | -1.66 | 0.00 | 0.00 | B1ANR0 | Polyadenylate-binding protein (PABP) | 615 | PABPC4 | 4 | 2 | 4 | 8.5 | 4.4 | 8.8 |
| 105 | -1.65 | 0.00 | 0.43 | P08243 | Asparagine synthetase [glutamine-hydrolyzing] (EC 6.3.5.4) (Cell cycle control protein TS11) (Glutamine-dependent asparagine synthetase) | 561 | ASNS | 2 | 0 | 1 | 4.6 | 0 | 2 |
| 106 | -1.64 | -0.54 | 0.75 | A0A087WSW9 | Thioredoxin reductase 1, cytoplasmic | 548 | TXNRD1 | 9 | 1 | 10 | 23.2 | 1.8 | 19.3 |

| | | | | | | | | | | | | | |
|---|---|---|---|---|---|---|---|---|---|---|---|---|---|
| 107 | -1.64 | 0.17 | 1.57 | P61353 | 60S ribosomal protein L27 | 136 | RPL27 | 1 | 1 | 2 | 6.6 | 6.6 | 22.1 |
| 108 | -1.63 | 0.00 | 0.83 | E9PNF3 | L-aminoadipate-semialdehyde dehydrogenase-phosphopantetheinyl transferase (Fragment) | 160 | AASDHPPT | 2 | 0 | 1 | 17.5 | 0 | 10.6 |
| 109 | -1.61 | 0.00 | 0.32 | D6RD69 | GTP-binding protein SAR1b (Fragment) | 170 | SAR1B | 3 | 2 | 4 | 27.6 | 11.2 | 34.1 |
| 110 | -1.61 | 1.06 | 2.06 | Q99880 | Histone H2B type 1-L (Histone H2B.c) (H2B/c) | 126 | HIST1H2BL | 2 | 4 | 3 | 15.1 | 23.8 | 27 |
| 111 | -1.61 | 0.92 | 2.62 | Q5T7C4 | High mobility group protein B1 | 158 | HMGB1 | 5 | 5 | 2 | 32.9 | 34.8 | 14.6 |
| 112 | -1.58 | 0.76 | 0.00 | E9PGT1 | Translin | 223 | TSN | 1 | 1 | 0 | 7.2 | 7.2 | 0 |
| 113 | -1.58 | 0.77 | 1.63 | Q08257 | Quinone oxidoreductase (EC 1.6.5.5) (NADPH:quinone reductase) (Zeta-crystallin) | 329 | CRYZ | 4 | 5 | 3 | 21.7 | 24.1 | 17.3 |
| 114 | -1.57 | 1.09 | 1.12 | P68032 | Actin, alpha cardiac muscle 1 (Alpha-cardiac actin) | 377 | ACTC1 | 7 | 10 | 7 | 22.3 | 30.5 | 22.3 |
| 115 | -1.55 | 1.94 | 0.00 | Q9BYN0 | Sulfiredoxin-1 (EC 1.8.98.2) | 137 | SRXN1 | 1 | 2 | 0 | 10.2 | 14.6 | 0 |
| 116 | -1.54 | 0.13 | 1.69 | M0R3D6 | 60S ribosomal protein L18a (Fragment) | 141 | RPL18A | 2 | 4 | 1 | 13.5 | 29.1 | 7.1 |
| 117 | -1.54 | 0.56 | 0.70 | P11586 | C-1-tetrahydrofolate synthase, cytoplasmic (C1-THF synthase) [Cleaved into: C-1-tetrahydrofolate synthase, cytoplasmic, N-terminally processed] [Includes: Methylenetetrahydrofolate dehydrogenase (EC 1.5.1.5); Methenyltetrahydrofolate cyclohydrolase (EC 3.5.4.9); Formyltetrahydrofolate synthetase (EC 6.3.4.3)] | 935 | MTHFD1 | 5 | 4 | 6 | 7.4 | 6.8 | 10.9 |
| 118 | -1.54 | 0.00 | 1.57 | J3QSB5 | 60S ribosomal protein L36 | 94 | RPL36 | 1 | 1 | 1 | 9.6 | 9.6 | 9.6 |
| 119 | -1.53 | 0.00 | 0.00 | Q9NUQ9 | Protein FAM49B (L1) | 324 | FAM49B | 1 | 0 | 0 | 10.7 | 0 | 0 |
| 120 | -1.53 | 0.00 | 1.85 | F5H6T1 | ARP2 actin-related protein 2 homolog (Yeast), isoform CRA_d (Actin-related protein 2) | 339 | ACTR2 | 2 | 0 | 2 | 8 | 0 | 9.4 |
| 121 | -1.53 | 0.00 | 2.15 | P62277 | 40S ribosomal protein S13 | 151 | RPS13 | 4 | 1 | 3 | 27.8 | 7.3 | 25.2 |

| | | | | | | | | | | | | | |
|---|---|---|---|---|---|---|---|---|---|---|---|---|---|
| 122 | -1.53 | 0.00 | 0.00 | F8WBT4 | Ribulose-phosphate 3-epimerase | 60 | RPE | 1 | 0 | 0 | 31.7 | 0 | 0 |
| 123 | -1.52 | 2.22 | 1.06 | Q9UJZ1 | Stomatin-like protein 2, mitochondrial (SLP-2) (EPB72-like protein 2) (Paraprotein target 7) (Paratarg-7) | 356 | STOML2 | 2 | 3 | 6 | 10.3 | 15.4 | 39.5 |
| 124 | -1.51 | 0.00 | 0.00 | B0QZK9 | Heterochromatin protein 1-binding protein 3 (Fragment) | 75 | HP1BP3 | 1 | 0 | 0 | 20 | 0 | 0 |
| 125 | -1.50 | 0.48 | 0.30 | Q9NTK5 | Obg-like ATPase 1 (DNA damage-regulated overexpressed in cancer 45) (DOC45) (GTP-binding protein 9) | 396 | OLA1 | 4 | 1 | 5 | 12.9 | 3.8 | 14.6 |
| 126 | -1.50 | 0.38 | 2.18 | P62854 | 40S ribosomal protein S26 | 115 | RPS26 | 1 | 1 | 1 | 13 | 13 | 13 |
| 127 | -1.49 | 0.51 | 1.87 | P62081 | 40S ribosomal protein S7 | 194 | RPS7 | 6 | 6 | 7 | 34.5 | 30.4 | 45.9 |
| 128 | -1.49 | 0.46 | 0.09 | Q15691 | Microtubule-associated protein RP/EB family member 1 (APC-binding protein EB1) (End-binding protein 1) (EB1) | 268 | MAPRE1 | 3 | 7 | 2 | 15.7 | 27.2 | 11.2 |
| 129 | -1.49 | 0.00 | 1.15 | H0YL99 | 28S ribosomal protein S11, mitochondrial | 118 | MRPS11 | 2 | 0 | 1 | 22 | 0 | 9.3 |
| 130 | -1.48 | 0.95 | 1.09 | P23396 | 40S ribosomal protein S3 (EC 4.2.99.18) | 243 | RPS3 | 9 | 10 | 8 | 45.3 | 47.7 | 30 |
| 131 | -1.47 | 0.00 | 0.00 | Q9UIM3 | FK506-binding protein-like (WAF-1/CIP1 stabilizing protein 39) (WISp39) | 349 | FKBPL | 1 | 0 | 0 | 2.3 | 0 | 0 |
| 132 | -1.47 | 0.00 | 0.00 | O00764 | Pyridoxal kinase (EC 2.7.1.35) (Pyridoxine kinase) | 312 | PDXK | 1 | 0 | 0 | 8.8 | 0 | 0 |
| 133 | -1.47 | 1.71 | 2.32 | Q9H061 | Transmembrane protein 126A | 195 | TMEM126A | 1 | 1 | 1 | 10.3 | 4.6 | 10.3 |
| 134 | -1.47 | 0.76 | 0.90 | P62263 | 40S ribosomal protein S14 | 151 | RPS14 | 5 | 2 | 2 | 30.5 | 13.9 | 7.9 |
| 135 | -1.47 | 0.92 | 0.67 | O96008 | Mitochondrial import receptor subunit TOM40 homolog (Protein Haymaker) (Translocase of outer membrane 40 kDa subunit homolog) (p38.5) | 361 | TOMM40 | 3 | 3 | 2 | 11.9 | 7.2 | 4.7 |
| 136 | -1.46 | 0.00 | 0.00 | P58557 | Putative ribonuclease | 167 | YBEY | 1 | 0 | 0 | 9.6 | 0 | 0 |

| | | | | | | | | | | | | | |
|---|---|---|---|---|---|---|---|---|---|---|---|---|---|
| 137 | -1.46 | 0.00 | 1.19 | P09972 | Fructose-bisphosphate aldolase C (EC 4.1.2.13) (Brain-type aldolase) | 364 | ALDOC | 1 | 1 | 1 | 6.3 | 1.9 | 6.3 |
| 138 | -1.46 | 1.23 | 0.00 | E5RJI7 | 39S ribosomal protein L13, mitochondrial (Fragment) | 147 | MRPL13 | 1 | 1 | 0 | 10.2 | 10.2 | 0 |
| 139 | -1.45 | 1.08 | 1.73 | Q9Y295 | Developmentally-regulated GTP-binding protein 1 (DRG-1) (Neural precursor cell expressed developmentally down-regulated protein 3) (NEDD-3) | 367 | DRG1 | 1 | 2 | 2 | 3 | 6.8 | 6.8 |
| 140 | -1.45 | 0.71 | 0.40 | Q9HC38 | Glyoxalase domain-containing protein 4 | 313 | GLOD4 | 4 | 2 | 4 | 13.8 | 8.1 | 13.1 |
| 141 | -1.45 | -0.09 | 0.39 | Q9UKK9 | ADP-sugar pyrophosphatase (EC 3.6.1.13) (8-oxo-dGDP phosphatase) (EC 3.6.1.58) (Nucleoside diphosphate-linked moiety X motif 5) (Nudix motif 5) (YSA1H) | 219 | NUDT5 | 4 | 3 | 5 | 26.5 | 16.4 | 26 |
| 142 | -1.44 | 0.67 | 1.06 | H0Y2V1 | Microtubule-associated protein (Fragment) | 463 | MAP4 | 1 | 1 | 1 | 3.7 | 3.7 | 3.7 |
| 143 | -1.44 | 1.58 | 0.84 | P04844 | Dolichyl-diphosphooligosaccharide--protein glycosyltransferase subunit 2 (EC 2.4.99.18) (Dolichyl-diphosphooligosaccharide--protein glycosyltransferase 63 kDa subunit) (RIBIIR) (Ribophorin II) (RPN-II) (Ribophorin-2) | 631 | RPN2 | 3 | 6 | 4 | 8.3 | 19 | 13.2 |
| 144 | -1.44 | 0.92 | 1.68 | M0R3H0 | 40S ribosomal protein S16 | 100 | RPS16 | 2 | 3 | 2 | 18 | 29 | 18 |
| 145 | -1.44 | 1.01 | 1.76 | K7ENG2 | Splicing factor U2AF 65 kDa subunit | 307 | U2AF2 | 3 | 1 | 2 | 8.5 | 5.2 | 9.1 |

| | | | | | | | | | | | | | |
|---|---|---|---|---|---|---|---|---|---|---|---|---|---|
| 146 | -1.43 | 0.79 | 1.11 | P23141 | Liver carboxylesterase 1 (Acyl-coenzyme A:cholesterol acyltransferase) (ACAT) (Brain carboxylesterase hBr1) (Carboxylesterase 1) (CE-1) (hCE-1) (EC 3.1.1.1) (Cocaine carboxylesterase) (Egasyn) (HMSE) (Methylumbelliferyl-acetate deacetylase 1) (EC 3.1.1.56) (Monocyte/macrophage serine esterase) (Retinyl ester hydrolase) (REH) (Serine esterase 1) (Triacylglycerol hydrolase) (TGH) | 567 | CES1 | 5 | 5 | 3 | 13.6 | 13.8 | 7.8 |
| 147 | -1.43 | 0.00 | 0.00 | Q9NR09 | Baculoviral IAP repeat-containing protein 6 (EC 6.3.2.-) (BIR repeat-containing ubiquitin-conjugating enzyme) (BRUCE) (Ubiquitin-conjugating BIR domain enzyme apollon) (APOLLON) | 4857 | BIRC6 | 1 | 0 | 0 | 0.2 | 0 | 0 |
| 148 | -1.43 | 0.00 | 0.92 | P29144 | Tripeptidyl-peptidase 2 (TPP-2) (EC 3.4.14.10) (Tripeptidyl aminopeptidase) (Tripeptidyl-peptidase II) (TPP-II) | 1249 | TPP2 | 2 | 0 | 1 | 2.2 | 0 | 0.8 |
| 149 | -1.43 | 0.63 | 0.59 | Q99832 | T-complex protein 1 subunit eta (TCP-1-eta) (CCT-eta) (HIV-1 Nef-interacting protein) | 543 | CCT7 | 10 | 4 | 10 | 20.1 | 10.9 | 22.5 |
| 150 | -1.43 | 0.00 | 1.03 | H0YKU1 | Tropomodulin-3 (Fragment) | 187 | TMOD3 | 1 | 0 | 1 | 7.5 | 0 | 7.5 |
| 151 | -1.42 | 1.13 | 1.83 | P30050 | 60S ribosomal protein L12 | 165 | RPL12 | 3 | 4 | 3 | 24.2 | 35.8 | 26.1 |
| 152 | -1.42 | 0.00 | 0.00 | F8VUA7 | Oxysterol-binding protein (Fragment) | 694 | OSBPL8 | 1 | 0 | 0 | 1.6 | 0 | 0 |
| 153 | -1.42 | -0.60 | 0.59 | P00491 | Purine nucleoside phosphorylase (PNP) (EC 2.4.2.1) (Inosine phosphorylase) (Inosine-guanosine phosphorylase) | 289 | PNP | 2 | 2 | 6 | 8.7 | 7.3 | 29.8 |
| 154 | -1.42 | 1.07 | 1.05 | Q9Y5M8 | Signal recognition particle receptor subunit beta (SR-beta) (Protein APMCF1) | 271 | SRPRB | 1 | 3 | 1 | 7 | 17 | 5.9 |

| | | | | | | | | | | | | | |
|---|---|---|---|---|---|---|---|---|---|---|---|---|---|
| 155 | -1.41 | 0.05 | 0.39 | Q14566 | DNA replication licensing factor MCM6 (EC 3.6.4.12) (p105MCM) | 821 | MCM6 | 5 | 1 | 3 | 10.6 | 2.1 | 4.9 |
| 156 | -1.41 | 0.89 | 0.99 | J3QRD1 | Fatty aldehyde dehydrogenase | 393 | ALDH3A2 | 7 | 4 | 6 | 23.7 | 13.2 | 20.4 |
| 157 | -1.40 | 0.48 | 1.23 | Q02878 | 60S ribosomal protein L6 (Neoplasm-related protein C140) (Tax-responsive enhancer element-binding protein 107) (TaxREB107) | 288 | RPL6 | 4 | 2 | 4 | 16.3 | 8.7 | 14.2 |
| 158 | -1.40 | 1.01 | 1.45 | P41250 | Glycine--tRNA ligase (EC 6.1.1.14) (Diadenosine tetraphosphate synthetase) (AP-4-A synthetase) (Glycyl-tRNA synthetase) (GlyRS) | 739 | GARS | 5 | 1 | 5 | 9.3 | 1.5 | 11 |
| 159 | -1.39 | 0.65 | 1.52 | H0YEN5 | 40S ribosomal protein S2 (Fragment) | 195 | RPS2 | 4 | 5 | 5 | 24.1 | 28.2 | 26.7 |
| 160 | -1.39 | 0.61 | 1.25 | Q5JP53 | Tubulin beta chain | 426 | TUBB | 6 | 10 | 9 | 20.9 | 34.3 | 27.2 |
| 161 | -1.39 | 0.63 | 0.61 | P30153 | Serine/threonine-protein phosphatase 2A 65 kDa regulatory subunit A alpha isoform (Medium tumor antigen-associated 61 kDa protein) (PP2A subunit A isoform PR65-alpha) (PP2A subunit A isoform R1-alpha) | 589 | PPP2R1A | 4 | 2 | 2 | 9.7 | 5.3 | 5.1 |
| 162 | -1.39 | 1.20 | 0.90 | B4DJV2 | Citrate synthase | 453 | CS | 5 | 2 | 5 | 9.7 | 4.4 | 9.5 |
| 163 | -1.38 | -0.02 | 0.78 | P11413 | Glucose-6-phosphate 1-dehydrogenase (G6PD) (EC 1.1.1.49) | 515 | G6PD | 11 | 6 | 12 | 27 | 12.4 | 34.6 |
| 164 | -1.38 | 1.21 | 2.04 | Q9HDC9 | Adipocyte plasma membrane-associated protein (Protein BSCv) | 416 | APMAP | 3 | 4 | 2 | 14.2 | 19.7 | 11.4 |
| 165 | -1.38 | -0.24 | 0.90 | P46781 | 40S ribosomal protein S9 | 194 | RPS9 | 3 | 3 | 3 | 15.5 | 11.9 | 11.9 |
| 166 | -1.38 | 0.00 | 0.00 | H7C286 | N-acetyl-D-glucosamine kinase | 196 | NAGK | 1 | 0 | 0 | 8.7 | 0 | 0 |

| | | | | | | | | | | | | | |
|---|---|---|---|---|---|---|---|---|---|---|---|---|---|
| 167 | -1.37 | 0.54 | 1.78 | Q9NZI8 | Insulin-like growth factor 2 mRNA-binding protein 1 (IGF2 mRNA-binding protein 1) (IMP-1) (IMP1) (Coding region determinant-binding protein) (CRD-BP) (IGF-II mRNA-binding protein 1) (VICKZ family member 1) (Zipcode-binding protein 1) (ZBP-1) | 577 | IGF2BP1 | 1 | 1 | 1 | 2.4 | 2.4 | 2.4 |
| 168 | -1.37 | 0.74 | 1.39 | P62913 | 60S ribosomal protein L11 (CLL-associated antigen KW-12) | 178 | RPL11 | 4 | 3 | 4 | 18.1 | 16.9 | 18.1 |
| 169 | -1.36 | 1.53 | 2.42 | P68036 | Ubiquitin-conjugating enzyme E2 L3 (EC 6.3.2.19) (L-UBC) (UbcH7) (Ubiquitin carrier protein L3) (Ubiquitin-conjugating enzyme E2-F1) (Ubiquitin-protein ligase L3) | 154 | UBE2L3 | 1 | 2 | 2 | 7.4 | 19.7 | 19.7 |
| 170 | -1.36 | 1.38 | 1.53 | Q99714 | 3-hydroxyacyl-CoA dehydrogenase type-2 (EC 1.1.1.35) (17-beta-hydroxysteroid dehydrogenase 10) (17-beta-HSD 10) (EC 1.1.1.51) (3-hydroxy-2-methylbutyryl-CoA dehydrogenase) (EC 1.1.1.178) (3-hydroxyacyl-CoA dehydrogenase type II) (Endoplasmic reticulum-associated amyloid beta-peptide-binding protein) (Mitochondrial ribonuclease P protein 2) (Mitochondrial RNase P protein 2) (Short chain dehydrogenase/reductase family 5C member 1) (Short-chain type dehydrogenase/reductase XH98G2) (Type II HADH) | 261 | HSD17B10 | 5 | 7 | 9 | 33.7 | 51.3 | 51.7 |
| 171 | -1.36 | 2.04 | 1.27 | F8WAU4 | Elongation factor G, mitochondrial | 453 | GFM1 | 1 | 1 | 1 | 2.4 | 2.4 | 2.4 |
| 172 | -1.36 | 0.00 | 0.00 | Q6PIU2 | Neutral cholesterol ester hydrolase 1 (NCEH) (EC 3.1.1.-) (Arylacetamide deacetylase-like 1) | 408 | NCEH1 | 1 | 0 | 0 | 5.8 | 0 | 0 |

| | | | | | | | | | | | | | |
|---|---|---|---|---|---|---|---|---|---|---|---|---|---|
| 173 | -1.35 | -0.56 | 0.00 | Q9H8S9 | MOB kinase activator 1A (Mob1 alpha) (Mob1A) (Mob1 homolog 1B) (Mps one binder kinase activator-like 1B) | 216 | MOB1A | 1 | 1 | 1 | 5.1 | 5.6 | 5.1 |
| 174 | -1.35 | 1.74 | 1.34 | Q15084 | Protein disulfide-isomerase A6 (EC 5.3.4.1) (Endoplasmic reticulum protein 5) (ER protein 5) (ERp5) (Protein disulfide isomerase P5) (Thioredoxin domain-containing protein 7) | 440 | PDIA6 | 6 | 6 | 7 | 21.5 | 20.4 | 25.4 |
| 175 | -1.35 | 1.20 | 0.46 | Q53GQ0 | Very-long-chain 3-oxoacyl-CoA reductase (EC 1.1.1.330) (17-beta-hydroxysteroid dehydrogenase 12) (17-beta-HSD 12) (3-ketoacyl-CoA reductase) (KAR) (Estradiol 17-beta-dehydrogenase 12) (EC 1.1.1.62) (Short chain dehydrogenase/reductase family 12C member 1) | 312 | HSD17B12 | 2 | 5 | 2 | 9.3 | 17 | 9.6 |
| 176 | -1.34 | 0.42 | 1.69 | H0YA96 | Heterogeneous nuclear ribonucleoprotein D0 (Fragment) | 210 | HNRNPD | 3 | 3 | 3 | 15.2 | 12.4 | 15.2 |
| 177 | -1.33 | 0.00 | -0.08 | G3V158 | 2-deoxyribose-5-phosphate aldolase homolog (C. elegans), isoform CRA_a (Deoxyribose-phosphate aldolase) | 230 | DERA | 1 | 0 | 1 | 6.5 | 0 | 5.2 |
| 178 | -1.33 | 0.23 | 0.83 | P62750 | 60S ribosomal protein L23a | 156 | RPL23A | 4 | 4 | 4 | 20.5 | 19.2 | 18.6 |
| 179 | -1.33 | 0.63 | 1.11 | Q58FF8 | Putative heat shock protein HSP 90-beta 2 (Heat shock protein 90-beta b) (Heat shock protein 90Bb) | 381 | HSP90AB2P | 8 | 4 | 8 | 19.7 | 11.3 | 19.7 |
| 180 | -1.32 | 0.07 | 0.61 | P62937 | Peptidyl-prolyl cis-trans isomerase A (PPIase A) (EC 5.2.1.8) (Cyclophilin A) (Cyclosporin A-binding protein) (Rotamase A) [Cleaved into: Peptidyl-prolyl cis-trans isomerase A, N-terminally processed] | 165 | PPIA | 9 | 9 | 7 | 43 | 38.8 | 33.9 |

| | | | | | | | | | | | | | |
|---|---|---|---|---|---|---|---|---|---|---|---|---|---|
| 181 | -1.32 | 0.31 | 1.12 | P54136 | Arginine--tRNA ligase, cytoplasmic (EC 6.1.1.19) (Arginyl-tRNA synthetase) (ArgRS) | 660 | RARS | 5 | 4 | 4 | 10.4 | 8 | 8.2 |
| 182 | -1.32 | -0.05 | 0.76 | Q07020 | 60S ribosomal protein L18 | 188 | RPL18 | 2 | 2 | 1 | 13.2 | 15.1 | 6.9 |
| 183 | -1.32 | 2.41 | 0.00 | K9J7I2 | Uncharacterized protein | 107 | | 1 | 2 | 0 | 17.8 | 19.6 | 0 |
| 184 | -1.32 | 1.21 | 0.00 | C9J6B1 | Ras-related protein Ral-B (Fragment) | 167 | RALB | 1 | 1 | 0 | 9.6 | 9.6 | 0 |
| 185 | -1.32 | 1.59 | 1.74 | O75947 | ATP synthase subunit d, mitochondrial (ATPase subunit d) | 161 | ATP5H | 2 | 7 | 4 | 19.3 | 41 | 32.3 |
| 186 | -1.31 | 0.00 | 0.00 | P55957 | BH3-interacting domain death agonist (p22 BID) (BID) [Cleaved into: BH3-interacting domain death agonist p15 (p15 BID); BH3-interacting domain death agonist p13 (p13 BID); BH3-interacting domain death agonist p11 (p11 BID)] | 195 | BID | 1 | 1 | 0 | 15.2 | 15.2 | 0 |
| 187 | -1.31 | 1.42 | 1.30 | Q15029 | 116 kDa U5 small nuclear ribonucleoprotein component (Elongation factor Tu GTP-binding domain-containing protein 2) (SNU114 homolog) (hSNU114) (U5 snRNP-specific protein, 116 kDa) (U5-116 kDa) | 972 | EFTUD2 | 3 | 3 | 5 | 4.3 | 3.9 | 7.3 |
| 188 | -1.30 | -0.30 | 0.81 | P00390 | Glutathione reductase, mitochondrial (GR) (GRase) (EC 1.8.1.7) | 522 | GSR | 6 | 1 | 8 | 20.7 | 2.3 | 21.9 |
| 189 | -1.30 | 1.52 | 1.90 | Q9NR30 | Nucleolar RNA helicase 2 (EC 3.6.4.13) (DEAD box protein 21) (Gu-alpha) (Nucleolar RNA helicase Gu) (Nucleolar RNA helicase II) (RH II/Gu) | 783 | DDX21 | 3 | 1 | 4 | 4.6 | 1.5 | 6.2 |
| 190 | -1.30 | 0.03 | -0.97 | P47897 | Glutamine--tRNA ligase (EC 6.1.1.18) (Glutaminyl-tRNA synthetase) (GlnRS) | 775 | QARS | 5 | 1 | 1 | 7.7 | 2.1 | 2.1 |
| 191 | -1.30 | 0.65 | 0.52 | P49368 | T-complex protein 1 subunit gamma (TCP-1-gamma) (CCT-gamma) (hTRiC5) | 545 | CCT3 | 9 | 3 | 5 | 17.6 | 6.2 | 8.4 |

| | | | | | | | | | | | | | |
|---|---|---|---|---|---|---|---|---|---|---|---|---|---|
| 192 | -1.30 | 2.06 | 1.36 | P23246 | Splicing factor, proline- and glutamine-rich (100 kDa DNA-pairing protein) (hPOMp100) (DNA-binding p52/p100 complex, 100 kDa subunit) (Polypyrimidine tract-binding protein-associated-splicing factor) (PSF) (PTB-associated-splicing factor) | 707 | SFPQ | 1 | 4 | 2 | 2.4 | 9.6 | 5.1 |
| 193 | -1.29 | 0.84 | 1.29 | Q9HB71 | Calcyclin-binding protein (CacyBP) (hCacyBP) (S100A6-binding protein) (Siah-interacting protein) | 228 | CACYBP | 6 | 3 | 6 | 40.4 | 17.1 | 36 |
| 194 | -1.29 | -0.36 | -0.11 | Q7Z6Z7 | E3 ubiquitin-protein ligase HUWE1 (EC 6.3.2.-) (ARF-binding protein 1) (ARF-BP1) (HECT, UBA and WWE domain-containing protein 1) (Homologous to E6AP carboxyl terminus homologous protein 9) (HectH9) (Large structure of UREB1) (LASU1) (Mcl-1 ubiquitin ligase E3) (Mule) (Upstream regulatory element-binding protein 1) (URE-B1) (URE-binding protein 1) | 4374 | HUWE1 | 2 | 1 | 1 | 0.8 | 0.3 | 0.3 |
| 195 | -1.29 | -0.29 | 0.94 | Q99497 | Protein deglycase DJ-1 (DJ-1) (EC 3.1.2.-) (EC 3.5.1.-) (Oncogene DJ1) (Parkinson disease protein 7) | 189 | PARK7 | 10 | 6 | 9 | 68.8 | 36 | 50.8 |
| 196 | -1.29 | 1.34 | 0.87 | M0R3D4 | Prenylated Rab acceptor protein 1 (Rab acceptor 1 (Prenylated), isoform CRA_a) | 151 | RABAC1 | 1 | 1 | 1 | 9.9 | 9.9 | 9.9 |
| 197 | -1.29 | 0.00 | 2.91 | P35637 | RNA-binding protein FUS (75 kDa DNA-pairing protein) (Oncogene FUS) (Oncogene TLS) (POMp75) (Translocated in liposarcoma protein) | 526 | FUS | 2 | 0 | 1 | 5.3 | 0 | 3 |

| | | | | | | | | | | | | | |
|---|---|---|---|---|---|---|---|---|---|---|---|---|---|
| 198 | -1.29 | 0.00 | 0.62 | P20042 | Eukaryotic translation initiation factor 2 subunit 2 (Eukaryotic translation initiation factor 2 subunit beta) (eIF-2-beta) | 333 | EIF2S2 | 1 | 0 | 1 | 4.5 | 0 | 4.5 |
| 199 | -1.29 | 0.24 | 0.77 | P41567 | Eukaryotic translation initiation factor 1 (eIF1) (A121) (Protein translation factor SUI1 homolog) (Sui1iso1) | 113 | EIF1 | 1 | 4 | 1 | 15 | 53.1 | 14.2 |
| 200 | -1.29 | 0.90 | 0.79 | Q9UQ80 | Proliferation-associated protein 2G4 (Cell cycle protein p38-2G4 homolog) (hG4-1) (ErbB3-binding protein 1) | 394 | PA2G4 | 4 | 6 | 5 | 11.9 | 21.6 | 16.2 |
| 201 | -1.29 | 0.03 | 0.91 | Q04828 | Aldo-keto reductase family 1 member C1 (EC 1.1.1.-) (20-alpha-hydroxysteroid dehydrogenase) (20-alpha-HSD) (EC 1.1.1.149) (Chlordecone reductase homolog HAKRC) (Dihydrodiol dehydrogenase 1/2) (DD1/DD2) (High-affinity hepatic bile acid-binding protein) (HBAB) (Indanol dehydrogenase) (EC 1.1.1.112) (Trans-1,2-dihydrobenzene-1,2-diol dehydrogenase) (EC 1.3.1.20) | 323 | AKR1C1 | 8 | 9 | 10 | 39.6 | 43.3 | 49.2 |
| 202 | -1.29 | 0.00 | 0.00 | A2A2U4 | Ribosyldihydronicotinamide dehydrogenase [quinone] (Fragment) | 63 | NQO2 | 1 | 0 | 0 | 17.5 | 0 | 0 |
| 203 | -1.28 | 1.13 | 0.00 | Q8NC51 | Plasminogen activator inhibitor 1 RNA-binding protein (PAI1 RNA-binding protein 1) (PAI-RBP1) (SERPINE1 mRNA-binding protein 1) | 408 | SERBP1 | 1 | 1 | 0 | 4.1 | 2.3 | 0 |
| 204 | -1.27 | 0.41 | 0.91 | P56537 | Eukaryotic translation initiation factor 6 (eIF-6) (B(2)GCN homolog) (B4 integrin interactor) (CAB) (p27(BBP)) | 245 | EIF6 | 4 | 5 | 5 | 31.4 | 33.5 | 30.6 |

| | | | | | | | | | | | | | |
|---|---|---|---|---|---|---|---|---|---|---|---|---|---|
| 205 | -1.27 | -0.12 | 0.97 | Q9ULC4 | Malignant T-cell-amplified sequence 1 (MCT-1) (Multiple copies T-cell malignancies) | 181 | MCTS1 | 2 | 2 | 1 | 19.9 | 14.9 | 9.4 |
| 206 | -1.25 | 0.00 | 0.00 | A8MXH2 | Nucleosome assembly protein 1-like 4 (Fragment) | 156 | NAP1L4 | 1 | 0 | 0 | 11.5 | 0 | 0 |
| 207 | -1.23 | 0.00 | 1.78 | E9PR16 | Nuclear pore complex protein Nup160 (Fragment) | 1123 | NUP160 | 1 | 0 | 1 | 1.1 | 0 | 1.1 |
| 208 | -1.23 | 0.44 | 1.58 | P41252 | Isoleucine--tRNA ligase, cytoplasmic (EC 6.1.1.5) (Isoleucyl-tRNA synthetase) (IRS) (IleRS) | 1262 | IARS | 1 | 5 | 1 | 1 | 4.8 | 1 |
| 209 | -1.22 | 0.47 | 1.05 | Q96Q11 | CCA tRNA nucleotidyltransferase 1, mitochondrial (EC 2.7.7.72) (Mitochondrial tRNA nucleotidyl transferase, CCA-adding) (mt CCA-adding enzyme) (mt tRNA CCA-diphosphorylase) (mt tRNA CCA-pyrophosphorylase) (mt tRNA adenylyltransferase) | 434 | TRNT1 | 1 | 1 | 1 | 2.4 | 2.4 | 2.4 |
| 210 | -1.21 | 1.21 | 1.47 | P60866 | 40S ribosomal protein S20 | 119 | RPS20 | 2 | 2 | 1 | 22.7 | 19.3 | 12.6 |
| 211 | -1.21 | -0.03 | 0.04 | Q9NQR4 | Omega-amidase NIT2 (EC 3.5.1.3) (Nitrilase homolog 2) | 276 | NIT2 | 4 | 4 | 2 | 19.9 | 23.9 | 10.1 |
| 212 | -1.21 | 0.17 | 0.00 | Q13283 | Ras GTPase-activating protein-binding protein 1 (G3BP-1) (EC 3.6.4.12) (EC 3.6.4.13) (ATP-dependent DNA helicase VIII) (hDH VIII) (GAP SH3 domain-binding protein 1) | 466 | G3BP1 | 2 | 1 | 0 | 7.1 | 3.6 | 0 |
| 213 | -1.21 | -0.17 | 0.00 | Q14691 | DNA replication complex GINS protein PSF1 (GINS complex subunit 1) | 196 | GINS1 | 1 | 1 | 0 | 7.7 | 7.1 | 0 |

| | | | | | | | | | | | | | |
|---|---|---|---|---|---|---|---|---|---|---|---|---|---|
| 214 | -1.21 | 1.45 | 2.14 | P07858 | Cathepsin B (EC 3.4.22.1) (APP secretase) (APPS) (Cathepsin B1) [Cleaved into: Cathepsin B light chain; Cathepsin B heavy chain] | 339 | CTSB | 2 | 5 | 2 | 10.3 | 20.6 | 8.3 |
| 215 | -1.21 | 0.60 | 1.13 | Q02790 | Peptidyl-prolyl cis-trans isomerase FKBP4 (PPIase FKBP4) (EC 5.2.1.8) (51 kDa FK506-binding protein) (FKBP51) (52 kDa FK506-binding protein) (52 kDa FKBP) (FKBP-52) (59 kDa immunophilin) (p59) (FK506-binding protein 4) (FKBP-4) (FKBP59) (HSP-binding immunophilin) (HBI) (Immunophilin FKBP52) (Rotamase) [Cleaved into: Peptidyl-prolyl cis-trans isomerase FKBP4, N-terminally processed] | 459 | FKBP4 | 2 | 2 | 6 | 6.1 | 5.7 | 20 |
| 216 | -1.20 | 0.20 | 0.78 | P09211 | Glutathione S-transferase P (EC 2.5.1.18) (GST class-pi) (GSTP1-1) | 210 | GSTP1 | 1 | 4 | 1 | 4.8 | 26.7 | 4.8 |
| 217 | -1.20 | 0.13 | 0.00 | A6NDF3 | Protein PBDC1 | 232 | PBDC1 | 1 | 2 | 0 | 6.5 | 11.2 | 0 |
| 218 | -1.19 | 1.12 | 0.00 | P00387 | NADH-cytochrome b5 reductase 3 (B5R) (Cytochrome b5 reductase) (EC 1.6.2.2) (Diaphorase-1) [Cleaved into: NADH-cytochrome b5 reductase 3 membrane-bound form; NADH-cytochrome b5 reductase 3 soluble form] | 301 | CYB5R3 | 2 | 2 | 0 | 8.6 | 8.3 | 0 |
| 219 | -1.19 | 0.43 | 0.93 | P09960 | Leukotriene A-4 hydrolase (LTA-4 hydrolase) (EC 3.3.2.6) (Leukotriene A(4) hydrolase) | 611 | LTA4H | 5 | 1 | 3 | 9.7 | 1.9 | 5.6 |
| 220 | -1.18 | 0.98 | 1.18 | H0YHC3 | Nucleosome assembly protein 1-like 1 (Fragment) | 198 | NAP1L1 | 3 | 2 | 4 | 20.7 | 14.1 | 29.3 |
| 221 | -1.18 | 0.20 | 1.24 | D3YTB1 | 60S ribosomal protein L32 (Fragment) | 133 | RPL32 | 6 | 2 | 5 | 30.1 | 20.3 | 36.8 |
| 222 | -1.18 | 0.96 | 1.37 | P62318 | Small nuclear ribonucleoprotein Sm D3 (Sm-D3) (snRNP core protein D3) | 126 | SNRPD3 | 2 | 2 | 2 | 13.3 | 13.3 | 13.3 |

| | | | | | | | | | | | | | |
|---|---|---|---|---|---|---|---|---|---|---|---|---|---|
| 223 | -1.17 | 1.15 | 0.98 | P08195 | 4F2 cell-surface antigen heavy chain (4F2hc) (4F2 heavy chain antigen) (Lymphocyte activation antigen 4F2 large subunit) (Solute carrier family 3 member 2) (CD antigen CD98) | 630 | SLC3A2 | 13 | 14 | 12 | 32.4 | 31.2 | 27.1 |
| 224 | -1.17 | 1.11 | 1.09 | P13010 | X-ray repair cross-complementing protein 5 (EC 3.6.4.-) (86 kDa subunit of Ku antigen) (ATP-dependent DNA helicase 2 subunit 2) (ATP-dependent DNA helicase II 80 kDa subunit) (CTC box-binding factor 85 kDa subunit) (CTC85) (CTCBF) (DNA repair protein XRCC5) (Ku80) (Ku86) (Lupus Ku autoantigen protein p86) (Nuclear factor IV) (Thyroid-lupus autoantigen) (TLAA) (X-ray repair complementing defective repair in Chinese hamster cells 5 (double-strand-break rejoining)) | 732 | XRCC5 | 8 | 7 | 9 | 14.6 | 12.2 | 13.5 |
| 225 | -1.17 | 1.14 | 2.25 | Q15758 | Neutral amino acid transporter B(0) (ATB(0)) (Baboon M7 virus receptor) (RD114/simian type D retrovirus receptor) (Sodium-dependent neutral amino acid transporter type 2) (Solute carrier family 1 member 5) | 541 | SLC1A5 | 2 | 1 | 3 | 4.3 | 2.4 | 7.9 |
| 226 | -1.16 | 1.34 | 1.32 | P63261 | Actin, cytoplasmic 2 (Gamma-actin) [Cleaved into: Actin, cytoplasmic 2, N-terminally processed] | 375 | ACTG1 | 13 | 18 | 12 | 51.2 | 67.2 | 47.5 |
| 227 | -1.16 | 1.75 | 1.74 | P36578 | 60S ribosomal protein L4 (60S ribosomal protein L1) | 427 | RPL4 | 6 | 1 | 7 | 16.6 | 3.7 | 21.1 |
| 228 | -1.15 | 0.00 | 0.00 | Q2TAA2 | Isoamyl acetate-hydrolyzing esterase 1 homolog (EC 3.1.-.-) | 248 | IAH1 | 1 | 0 | 0 | 12.6 | 0 | 0 |

| | | | | | | | | | | | | | |
|---|---|---|---|---|---|---|---|---|---|---|---|---|---|
| 229 | -1.15 | -0.13 | 0.28 | P06744 | Glucose-6-phosphate isomerase (GPI) (EC 5.3.1.9) (Autocrine motility factor) (AMF) (Neuroleukin) (NLK) (Phosphoglucose isomerase) (PGI) (Phosphohexose isomerase) (PHI) (Sperm antigen 36) (SA-36) | 558 | GPI | 7 | 5 | 4 | 15.9 | 11.5 | 8.4 |
| 230 | -1.15 | 0.00 | 0.00 | Q13618 | Cullin-3 (CUL-3) | 768 | CUL3 | 1 | 0 | 1 | 1.3 | 0 | 1.5 |
| 231 | -1.15 | 0.00 | 0.00 | C9J255 | Mannose-1-phosphate guanyltransferase alpha (Fragment) | 179 | GMPPA | 1 | 0 | 0 | 5.6 | 0 | 0 |
| 232 | -1.15 | 0.00 | 0.96 | C9IZG3 | Ubiquitin fusion degradation protein 1 homolog (Fragment) | 190 | UFD1L | 2 | 0 | 1 | 12.1 | 0 | 5.3 |
| 233 | -1.15 | 0.09 | 0.41 | Q14847 | LIM and SH3 domain protein 1 (LASP-1) (Metastatic lymph node gene 50 protein) (MLN 50) | 261 | LASP1 | 1 | 5 | 1 | 5 | 19.9 | 3.4 |
| 234 | -1.14 | -0.16 | -0.30 | A2A2D0 | Stathmin (Fragment) | 85 | STMN1 | 3 | 3 | 1 | 42.4 | 35.3 | 10.6 |
| 235 | -1.14 | 0.58 | 1.04 | P05388 | 60S acidic ribosomal protein P0 (60S ribosomal protein L10E) | 317 | RPLP0 | 10 | 8 | 11 | 45.4 | 32.8 | 47.3 |
| 236 | -1.14 | -0.29 | 1.15 | Q5W0S5 | UV excision repair protein RAD23 homolog B (Fragment) | 146 | RAD23B | 1 | 2 | 1 | 7.5 | 13 | 7.5 |
| 237 | -1.14 | 1.75 | 0.00 | Q01650 | Large neutral amino acids transporter small subunit 1 (4F2 light chain) (4F2 LC) (4F2LC) (CD98 light chain) (Integral membrane protein E16) (L-type amino acid transporter 1) (hLAT1) (Solute carrier family 7 member 5) (y+ system cationic amino acid transporter) | 507 | SLC7A5 | 1 | 3 | 1 | 3.6 | 7.1 | 1.8 |
| 238 | -1.14 | 0.00 | 0.98 | A6NJA2 | Ubiquitin carboxyl-terminal hydrolase (EC 3.4.19.12) | 448 | USP14 | 2 | 0 | 1 | 5.4 | 0 | 2.5 |
| 239 | -1.14 | 0.00 | 1.06 | C9JVE2 | DCN1-like protein (Defective in cullin neddylation protein 1-like protein) | 244 | DCUN1D1 | 2 | 0 | 1 | 12.3 | 0 | 5.3 |

| | | | | | | | | | | | | | |
|---|---|---|---|---|---|---|---|---|---|---|---|---|---|
| 240 | -1.14 | 0.00 | 0.00 | P23381 | Tryptophan--tRNA ligase, cytoplasmic (EC 6.1.1.2) (Interferon-induced protein 53) (IFP53) (Tryptophanyl-tRNA synthetase) (TrpRS) (hWRS) [Cleaved into: T1-TrpRS; T2-TrpRS] | 471 | WARS | 2 | 0 | 0 | 8.3 | 0 | 0 |
| 241 | -1.13 | 0.00 | 0.81 | A0A087 WW66 | 26S proteasome non-ATPase regulatory subunit 1 | 953 | PSMD1 | 5 | 0 | 5 | 8 | 0 | 7 |
| 242 | -1.13 | 0.94 | 1.65 | P19338 | Nucleolin (Protein C23) | 710 | NCL | 12 | 10 | 10 | 18.5 | 17.2 | 16.8 |
| 243 | -1.13 | 0.76 | 1.35 | Q15365 | Poly(rC)-binding protein 1 (Alpha-CP1) (Heterogeneous nuclear ribonucleoprotein E1) (hnRNP E1) (Nucleic acid-binding protein SUB2.3) | 356 | PCBP1 | 1 | 3 | 2 | 7 | 10.7 | 9.3 |
| 244 | -1.12 | 1.72 | 1.74 | Q3ZCQ8 | Mitochondrial import inner membrane translocase subunit TIM50 | 353 | TIMM50 | 3 | 3 | 3 | 12.2 | 12.2 | 12.2 |
| 245 | -1.12 | 0.53 | 0.94 | P31939 | Bifunctional purine biosynthesis protein PURH [Includes: Phosphoribosylaminoimidazolecarboxamide formyltransferase (EC 2.1.2.3) (5-aminoimidazole-4-carboxamide ribonucleotide formyltransferase) (AICAR transformylase); IMP cyclohydrolase (EC 3.5.4.10) (ATIC) (IMP synthase) (Inosinicase)] | 592 | ATIC | 13 | 4 | 10 | 27 | 10.5 | 21.3 |
| 246 | -1.12 | 2.27 | 1.57 | A0A0A0 MTN9 | NADPH:adrenodoxin oxidoreductase, mitochondrial (EC 1.18.1.6) | 439 | FDXR | 1 | 2 | 5 | 3.2 | 5.7 | 16.9 |
| 247 | -1.12 | 0.00 | 0.00 | D6R918 | OCIA domain-containing protein 1 (Fragment) | 73 | OCIAD1 | 1 | 0 | 0 | 24.7 | 0 | 0 |
| 248 | -1.12 | 1.01 | 1.07 | P17812 | CTP synthase 1 (EC 6.3.4.2) (CTP synthetase 1) (UTP--ammonia ligase 1) | 591 | CTPS1 | 2 | 1 | 2 | 3.4 | 2.7 | 5.1 |
| 249 | -1.12 | 1.42 | 1.13 | J3QS48 | Mannose-P-dolichol utilization defect 1 protein | 101 | MPDU1 | 2 | 2 | 2 | 23.8 | 23.8 | 23.8 |

| | | | | | | | | | | | | | |
|---|---|---|---|---|---|---|---|---|---|---|---|---|---|
| 250 | -1.11 | 0.99 | 1.05 | Q9H3N1 | Thioredoxin-related transmembrane protein 1 (Thioredoxin domain-containing protein 1) (Transmembrane Trx-related protein) | 280 | TMX1 | 2 | 6 | 3 | 8.2 | 20.7 | 11.4 |
| 251 | -1.11 | -0.06 | 0.79 | P46778 | 60S ribosomal protein L21 | 160 | RPL21 | 2 | 2 | 1 | 16.2 | 13.8 | 9.4 |
| 252 | -1.11 | -0.25 | 0.24 | O60493 | Sorting nexin-3 (Protein SDP3) | 162 | SNX3 | 3 | 3 | 2 | 16 | 10.5 | 10.5 |
| 253 | -1.10 | 0.00 | 0.00 | P61966 | AP-1 complex subunit sigma-1A (Adaptor protein complex AP-1 subunit sigma-1A) (Adaptor-related protein complex 1 subunit sigma-1A) (Clathrin assembly protein complex 1 sigma-1A small chain) (Clathrin coat assembly protein AP19) (Golgi adaptor HA1/AP1 adaptin sigma-1A subunit) (HA1 19 kDa subunit) (Sigma 1a subunit of AP-1 clathrin) (Sigma-adaptin 1A) (Sigma1A-adaptin) | 158 | AP1S1 | 1 | 0 | 0 | 10.1 | 0 | 0 |
| 254 | -1.10 | 1.64 | 0.99 | Q9NYL4 | Peptidyl-prolyl cis-trans isomerase FKBP11 (PPIase FKBP11) (EC 5.2.1.8) (19 kDa FK506-binding protein) (19 kDa FKBP) (FKBP-19) (FK506-binding protein 11) (FKBP-11) (Rotamase) | 201 | FKBP11 | 1 | 2 | 1 | 7.5 | 14.4 | 6.8 |
| 255 | -1.10 | -0.20 | 0.14 | H0YJG7 | Activator of 90 kDa heat shock protein ATPase homolog 1 (Fragment) | 216 | AHSA1 | 2 | 1 | 1 | 15.3 | 5.1 | 5.1 |
| 256 | -1.10 | 0.73 | 1.41 | H0Y8E6 | DNA replication licensing factor MCM2 (Fragment) | 836 | MCM2 | 4 | 3 | 1 | 6.1 | 4.4 | 1.4 |
| 257 | -1.09 | 0.58 | 1.21 | P55145 | Mesencephalic astrocyte-derived neurotrophic factor (Arginine-rich protein) (Protein ARMET) | 182 | MANF | 2 | 4 | 3 | 18.7 | 19.8 | 24.7 |

| | | | | | | | | | | | | | |
|---|---|---|---|---|---|---|---|---|---|---|---|---|---|
| 258 | -1.09 | 0.46 | 0.79 | Q14204 | Cytoplasmic dynein 1 heavy chain 1 (Cytoplasmic dynein heavy chain 1) (Dynein heavy chain, cytosolic) | 4646 | DYNC1H1 | 6 | 30 | 29 | 2.3 | 8.3 | 8.1 |
| 259 | -1.09 | 1.66 | 1.18 | P05023 | Sodium/potassium-transporting ATPase subunit alpha-1 (Na(+)/K(+) ATPase alpha-1 subunit) (EC 3.6.3.9) (Sodium pump subunit alpha-1) | 1023 | ATP1A1 | 4 | 4 | 6 | 6 | 5.2 | 7.4 |
| 260 | -1.09 | 0.00 | 0.70 | P04075 | Fructose-bisphosphate aldolase A (EC 4.1.2.13) (Lung cancer antigen NY-LU-1) (Muscle-type aldolase) | 364 | ALDOA | 7 | 9 | 7 | 28.8 | 30.5 | 31.3 |
| 261 | -1.08 | 0.00 | 0.00 | P55263 | Adenosine kinase (AK) (EC 2.7.1.20) (Adenosine 5'-phosphotransferase) | 362 | ADK | 1 | 0 | 0 | 4.9 | 0 | 0 |
| 262 | -1.08 | 0.00 | 1.09 | I3L4U9 | Nuclear protein localization protein 4 homolog | 135 | NPLOC4 | 1 | 0 | 1 | 13.3 | 0 | 13.3 |
| 263 | -1.08 | 0.00 | 3.58 | Q96AE4 | Far upstream element-binding protein 1 (FBP) (FUSE-binding protein 1) (DNA helicase V) (hDH V) | 644 | FUBP1 | 1 | 0 | 1 | 1.9 | 0 | 4 |
| 264 | -1.08 | 1.74 | 1.78 | H0YLR3 | U2 small nuclear ribonucleoprotein A' (Fragment) | 89 | SNRPA1 | 1 | 1 | 1 | 12.4 | 13.5 | 13.5 |
| 265 | -1.08 | 0.75 | 0.71 | Q9Y277 | Voltage-dependent anion-selective channel protein 3 (VDAC-3) (hVDAC3) (Outer mitochondrial membrane protein porin 3) | 283 | VDAC3 | 3 | 4 | 3 | 8.8 | 16.3 | 17 |
| 266 | -1.07 | 1.27 | 0.00 | H0YLA2 | Signal recognition particle 14 kDa protein | 115 | SRP14 | 1 | 1 | 0 | 11.3 | 11.3 | 0 |
| 267 | -1.07 | 0.39 | 0.76 | P07900 | Heat shock protein HSP 90-alpha (Heat shock 86 kDa) (HSP 86) (HSP86) (Lipopolysaccharide-associated protein 2) (LAP-2) (LPS-associated protein 2) (Renal carcinoma antigen NY-REN-38) | 732 | HSP90AA1 | 29 | 14 | 25 | 37.3 | 24.2 | 32.1 |

| | | | | | | | | | | | | | |
|---|---|---|---|---|---|---|---|---|---|---|---|---|---|
| 268 | -1.06 | 0.51 | 0.92 | P48507 | Glutamate--cysteine ligase regulatory subunit (GCS light chain) (Gamma-ECS regulatory subunit) (Gamma-glutamylcysteine synthetase regulatory subunit) (Glutamate--cysteine ligase modifier subunit) | 274 | GCLM | 3 | 2 | 1 | 15.3 | 10.6 | 5.8 |
| 269 | -1.06 | 0.79 | -0.13 | O15498 | Synaptobrevin homolog YKT6 (EC 2.3.1.-) | 198 | YKT6 | 1 | 2 | 1 | 10.4 | 18.3 | 5.5 |
| 270 | -1.06 | 0.79 | 0.84 | Q8NFH3 | Nucleoporin Nup43 (Nup107-160 subcomplex subunit Nup43) (p42) | 380 | NUP43 | 1 | 1 | 1 | 3.7 | 3.7 | 3.7 |
| 271 | -1.06 | 0.00 | 1.20 | O75607 | Nucleoplasmin-3 | 178 | NPM3 | 1 | 0 | 1 | 8.4 | 0 | 8.4 |
| 272 | -1.06 | 1.66 | 0.81 | Q5JRX3 | Presequence protease, mitochondrial (hPreP) (EC 3.4.24.-) (Pitrilysin metalloproteinase 1) (Metalloprotease 1) (hMP1) | 1037 | PITRM1 | 1 | 2 | 1 | 1.4 | 3.8 | 1.4 |
| 273 | -1.06 | 0.40 | 0.53 | P20290 | Transcription factor BTF3 (Nascent polypeptide-associated complex subunit beta) (NAC-beta) (RNA polymerase B transcription factor 3) | 206 | BTF3 | 4 | 3 | 4 | 30.2 | 29 | 38.3 |
| 274 | -1.06 | 0.57 | 0.83 | P28072 | Proteasome subunit beta type-6 (EC 3.4.25.1) (Macropain delta chain) (Multicatalytic endopeptidase complex delta chain) (Proteasome delta chain) (Proteasome subunit Y) | 239 | PSMB6 | 1 | 1 | 1 | 4.2 | 4.2 | 4.2 |
| 275 | -1.05 | 0.62 | 0.82 | P46777 | 60S ribosomal protein L5 | 297 | RPL5 | 3 | 1 | 6 | 13.1 | 4.7 | 22.6 |
| 276 | -1.05 | 0.11 | 1.16 | P62701 | 40S ribosomal protein S4, X isoform (SCR10) (Single copy abundant mRNA protein) | 263 | RPS4X | 6 | 3 | 5 | 20.9 | 9.5 | 19 |
| 277 | -1.05 | 0.00 | 1.05 | O00487 | 26S proteasome non-ATPase regulatory subunit 14 (EC 3.4.19.-) (26S proteasome regulatory subunit RPN11) (26S proteasome-associated PAD1 homolog 1) | 310 | PSMD14 | 1 | 0 | 1 | 4.2 | 0 | 4.2 |

| | | | | | | | | | | | | | |
|---|---|---|---|---|---|---|---|---|---|---|---|---|---|
| 278 | -1.04 | 1.41 | 0.86 | P37235 | Hippocalcin-like protein 1 (Calcium-binding protein BDR-1) (HLP2) (Visinin-like protein 3) (VILIP-3) | 193 | HPCAL1 | 1 | 3 | 1 | 6.2 | 19.2 | 6.2 |
| 279 | -1.04 | 0.86 | 1.15 | P22695 | Cytochrome b-c1 complex subunit 2, mitochondrial (Complex III subunit 2) (Core protein II) (Ubiquinol-cytochrome-c reductase complex core protein 2) | 453 | UQCRC2 | 8 | 4 | 5 | 29.1 | 12.6 | 17.9 |
| 280 | -1.04 | -0.11 | 1.74 | Q92499 | ATP-dependent RNA helicase DDX1 (EC 3.6.4.13) (DEAD box protein 1) (DEAD box protein retinoblastoma) (DBP-RB) | 740 | DDX1 | 1 | 1 | 1 | 2.9 | 2.9 | 2.9 |
| 281 | -1.04 | 0.14 | 0.97 | K7EK18 | Septin-9 (Fragment) | 195 | sept9 | 2 | 1 | 1 | 14.4 | 5.1 | 5.1 |
| 282 | -1.04 | 0.00 | -0.19 | Q9NRX4 | 14 kDa phosphohistidine phosphatase (EC 3.1.3.-) (Phosphohistidine phosphatase 1) (Protein janus-A homolog) | 125 | PHPT1 | 2 | 0 | 1 | 32.8 | 0 | 16 |
| 283 | -1.03 | 0.40 | 0.90 | P22314 | Ubiquitin-like modifier-activating enzyme 1 (Protein A1S9) (Ubiquitin-activating enzyme E1) | 1058 | UBA1 | 9 | 12 | 8 | 12.9 | 17.3 | 10.6 |
| 284 | -1.03 | 1.04 | 1.55 | P06756 | Integrin alpha-V (Vitronectin receptor subunit alpha) (CD antigen CD51) [Cleaved into: Integrin alpha-V heavy chain; Integrin alpha-V light chain] | 1048 | ITGAV | 3 | 4 | 3 | 3.4 | 5.9 | 5.6 |
| 285 | -1.02 | 0.92 | 1.25 | F8WAS3 | NADH dehydrogenase [ubiquinone] 1 alpha subcomplex subunit 5 | 70 | NDUFA5 | 1 | 4 | 1 | 14.3 | 51.4 | 14.3 |
| 286 | -1.02 | -0.07 | 0.88 | A0A087WYR0 | Signal recognition particle 19 kDa protein | 120 | SRP19 | 2 | 2 | 2 | 20 | 22.5 | 20 |
| 287 | -1.02 | 0.24 | 0.32 | P62979 | Ubiquitin-40S ribosomal protein S27a (Ubiquitin carboxyl extension protein 80) [Cleaved into: Ubiquitin; 40S ribosomal protein S27a] | 156 | RPS27A | 1 | 3 | 2 | 10.3 | 24.4 | 11.5 |

| | | | | | | | | | | | | | |
|---|---|---|---|---|---|---|---|---|---|---|---|---|---|
| 288 | -1.02 | 2.43 | 1.46 | Q13263 | Transcription intermediary factor 1-beta (TIF1-beta) (E3 SUMO-protein ligase TRIM28) (EC 6.3.2.-) (KRAB-associated protein 1) (KAP-1) (KRAB-interacting protein 1) (KRIP-1) (Nuclear corepressor KAP-1) (RING finger protein 96) (Tripartite motif-containing protein 28) | 835 | TRIM28 | 5 | 3 | 2 | 10.4 | 8.2 | 3.3 |
| 289 | -1.02 | 0.30 | 0.55 | P50395 | Rab GDP dissociation inhibitor beta (Rab GDI beta) (Guanosine diphosphate dissociation inhibitor 2) (GDI-2) | 445 | GDI2 | 5 | 5 | 6 | 13 | 16 | 16.6 |
| 290 | -1.02 | 1.26 | 0.00 | P26885 | Peptidyl-prolyl cis-trans isomerase FKBP2 (PPIase FKBP2) (EC 5.2.1.8) (13 kDa FK506-binding protein) (13 kDa FKBP) (FKBP-13) (FK506-binding protein 2) (FKBP-2) (Immunophilin FKBP13) (Rotamase) | 142 | FKBP2 | 1 | 3 | 0 | 10.6 | 33.8 | 0 |
| 291 | -1.02 | 1.22 | 1.25 | P27695 | DNA-(apurinic or apyrimidinic site) lyase (EC 3.1.-.-) (EC 4.2.99.18) (APEX nuclease) (APEN) (Apurinic-apyrimidinic endonuclease 1) (AP endonuclease 1) (APE-1) (REF-1) (Redox factor-1) [Cleaved into: DNA-(apurinic or apyrimidinic site) lyase, mitochondrial] | 318 | APEX1 | 1 | 2 | 3 | 5.7 | 12.3 | 16.7 |
| 292 | -1.02 | 0.37 | 0.89 | K7EJR3 | 26S proteasome non-ATPase regulatory subunit 8 (Fragment) | 250 | PSMD8 | 1 | 2 | 3 | 3.2 | 6.8 | 13.2 |
| 293 | -1.02 | 0.50 | 1.40 | B8ZZJ0 | Small ubiquitin-related modifier 1 | 58 | SUMO1 | 1 | 1 | 1 | 20.7 | 20.7 | 20.7 |
| 294 | -1.01 | 0.21 | 0.00 | P62633 | Cellular nucleic acid-binding protein (CNBP) (Zinc finger protein 9) | 177 | CNBP | 1 | 3 | 0 | 8.8 | 23.5 | 0 |

| | | | | | | | | | | | | | |
|---|---|---|---|---|---|---|---|---|---|---|---|---|---|
| 295 | -1.01 | 0.70 | 0.48 | P50990 | T-complex protein 1 subunit theta (TCP-1-theta) (CCT-theta) (Renal carcinoma antigen NY-REN-15) | 548 | CCT8 | 11 | 4 | 14 | 23.9 | 10.8 | 30.5 |
| 296 | -1.01 | 0.00 | 0.00 | C9JYM0 | Ribonuclease P protein subunit p20 (Fragment) | 137 | POP7 | 1 | 0 | 0 | 10.9 | 0 | 0 |
| 297 | -1.01 | 0.83 | 1.12 | E9PKD5 | 26S protease regulatory subunit 6A (Fragment) | 311 | PSMC3 | 3 | 2 | 5 | 14.5 | 9.6 | 24.4 |
| 298 | -1.01 | 0.27 | 0.89 | Q13151 | Heterogeneous nuclear ribonucleoprotein A0 (hnRNP A0) | 305 | HNRNPA0 | 1 | 2 | 1 | 4.9 | 2.6 | 4.9 |
| 299 | -1.01 | 0.00 | 0.00 | H7C531 | 26S proteasome non-ATPase regulatory subunit 6 (Fragment) | 138 | PSMD6 | 1 | 0 | 0 | 10.9 | 0 | 0 |
| 300 | -1.00 | 0.54 | 0.72 | P61923 | Coatomer subunit zeta-1 (Zeta-1-coat protein) (Zeta-1 COP) | 177 | COPZ1 | 3 | 4 | 2 | 29.9 | 30.5 | 16.9 |
| 301 | -1.00 | 0.48 | 1.83 | Q16555 | Dihydropyrimidinase-related protein 2 (DRP-2) (Collapsin response mediator protein 2) (CRMP-2) (N2A3) (Unc-33-like phosphoprotein 2) (ULIP-2) | 572 | DPYSL2 | 4 | 2 | 8 | 9.5 | 4.5 | 23.9 |
| 302 | -1.00 | 0.00 | 0.59 | O75822 | Eukaryotic translation initiation factor 3 subunit J (eIF3j) (Eukaryotic translation initiation factor 3 subunit 1) (eIF-3-alpha) (eIF3 p35) | 258 | EIF3J | 1 | 0 | 2 | 6.2 | 0 | 14.8 |
| 303 | -1.00 | -0.21 | 0.37 | P60174 | Triosephosphate isomerase (TIM) (EC 5.3.1.1) (Triose-phosphate isomerase) | 286 | TPI1 | 12 | 14 | 11 | 66.3 | 69.1 | 57 |
| 304 | -1.00 | 1.01 | 1.04 | P07339 | Cathepsin D (EC 3.4.23.5) [Cleaved into: Cathepsin D light chain; Cathepsin D heavy chain] | 412 | CTSD | 2 | 9 | 2 | 5.1 | 23.8 | 6.8 |
| 305 | -1.00 | 0.50 | 0.00 | A0A087WVZ9 | DNA-directed RNA polymerases I, II, and III subunit RPABC1 | 184 | POLR2E | 2 | 2 | 0 | 16.8 | 16.8 | 0 |

| | | | | | | | | | | | | | |
|---|---|---|---|---|---|---|---|---|---|---|---|---|---|
| 306 | -1.00 | -0.10 | 0.79 | P06733 | Alpha-enolase (EC 4.2.1.11) (2-phospho-D-glycerate hydro-lyase) (C-myc promoter-binding protein) (Enolase 1) (MBP-1) (MPB-1) (Non-neural enolase) (NNE) (Phosphopyruvate hydratase) (Plasminogen-binding protein) | 434 | ENO1 | 28 | 15 | 24 | 71 | 41.9 | 55.3 |
| 307 | -0.99 | 0.13 | 1.39 | P05455 | Lupus La protein (La autoantigen) (La ribonucleoprotein) (Sjoegren syndrome type B antigen) (SS-B) | 408 | SSB | 6 | 4 | 5 | 20.1 | 12.3 | 18.6 |
| 308 | -0.99 | 0.75 | 1.09 | A0A087WXM6 | 60S ribosomal protein L17 (Fragment) | 169 | RPL17 | 6 | 4 | 3 | 30.8 | 29.6 | 16 |
| 309 | -0.99 | 1.49 | 1.06 | P24752 | Acetyl-CoA acetyltransferase, mitochondrial (EC 2.3.1.9) (Acetoacetyl-CoA thiolase) (T2) | 427 | ACAT1 | 3 | 2 | 2 | 8 | 8.7 | 6.3 |
| 310 | -0.99 | 1.01 | 0.99 | B4DLN1 | Uncharacterized protein (cDNA FLJ60124, highly similar to Mitochondrial dicarboxylate carrier) | 442 | | 3 | 6 | 3 | 5.2 | 14.5 | 5.2 |
| 311 | -0.99 | 0.18 | 0.82 | D6R9P3 | Heterogeneous nuclear ribonucleoprotein A/B | 280 | HNRNPAB | 2 | 1 | 1 | 7.9 | 2.9 | 2.9 |
| 312 | -0.99 | 0.45 | 0.36 | P78371 | T-complex protein 1 subunit beta (TCP-1-beta) (CCT-beta) | 535 | CCT2 | 12 | 5 | 13 | 27.7 | 14 | 29.5 |
| 313 | -0.99 | 0.53 | 1.46 | P61158 | Actin-related protein 3 (Actin-like protein 3) | 418 | ACTR3 | 2 | 2 | 5 | 4.3 | 5.7 | 19.9 |
| 314 | -0.98 | 0.13 | 0.48 | P35270 | Sepiapterin reductase (SPR) (EC 1.1.1.153) | 261 | SPR | 6 | 2 | 3 | 32.2 | 13 | 13.8 |
| 315 | -0.98 | 0.20 | 0.59 | H0Y3P2 | Eukaryotic translation initiation factor 4 gamma 2 | 869 | EIF4G2 | 1 | 1 | 1 | 1.2 | 1.2 | 1.2 |
| 316 | -0.98 | 0.12 | -3.49 | O60925 | Prefoldin subunit 1 | 122 | PFDN1 | 1 | 2 | 1 | 8.2 | 16.4 | 10.7 |
| 317 | -0.98 | 0.12 | 1.12 | Q01518 | Adenylyl cyclase-associated protein 1 (CAP 1) | 475 | CAP1 | 4 | 5 | 4 | 14.1 | 14.1 | 12 |

| | | | | | | | | | | | | | |
|---|---|---|---|---|---|---|---|---|---|---|---|---|---|
| 318 | -0.98 | 2.02 | 0.49 | P08574 | Cytochrome c1, heme protein, mitochondrial (Complex III subunit 4) (Complex III subunit IV) (Cytochrome b-c1 complex subunit 4) (Ubiquinol-cytochrome-c reductase complex cytochrome c1 subunit) (Cytochrome c-1) | 325 | CYC1 | 2 | 1 | 2 | 8.9 | 7.4 | 11.4 |
| 319 | -0.98 | 0.86 | 0.96 | F8W8A6 | 3-hydroxyisobutyryl-CoA hydrolase, mitochondrial (Fragment) | 128 | HIBCH | 1 | 1 | 1 | 9.4 | 9.4 | 9.4 |
| 320 | -0.98 | 0.04 | 0.35 | P14618 | Pyruvate kinase PKM (EC 2.7.1.40) (Cytosolic thyroid hormone-binding protein) (CTHBP) (Opa-interacting protein 3) (OIP-3) (Pyruvate kinase 2/3) (Pyruvate kinase muscle isozyme) (Thyroid hormone-binding protein 1) (THBP1) (Tumor M2-PK) (p58) | 531 | PKM | 23 | 14 | 20 | 52.5 | 31.1 | 49.7 |
| 321 | -0.98 | -0.12 | 0.21 | P16152 | Carbonyl reductase [NADPH] 1 (EC 1.1.1.184) (15-hydroxyprostaglandin dehydrogenase [NADP(+)]) (EC 1.1.1.197) (NADPH-dependent carbonyl reductase 1) (Prostaglandin 9-ketoreductase) (Prostaglandin-E(2) 9-reductase) (EC 1.1.1.189) (Short chain dehydrogenase/reductase family 21C member 1) | 277 | CBR1 | 4 | 6 | 5 | 21.7 | 26 | 22.7 |
| 322 | -0.98 | 1.00 | 0.79 | P11021 | 78 kDa glucose-regulated protein (GRP-78) (Endoplasmic reticulum lumenal Ca(2+)-binding protein grp78) (Heat shock 70 kDa protein 5) (Immunoglobulin heavy chain-binding protein) (BiP) | 654 | HSPA5 | 17 | 12 | 12 | 32.4 | 28.1 | 22.5 |
| 323 | -0.98 | 1.01 | 0.59 | Q9HAV7 | GrpE protein homolog 1, mitochondrial (HMGE) (Mt-GrpE#1) | 217 | GRPEL1 | 3 | 5 | 3 | 18.9 | 27.2 | 18.9 |

| | | | | | | | | | | | | | |
|---|---|---|---|---|---|---|---|---|---|---|---|---|---|
| 324 | -0.98 | 1.37 | 1.31 | P00367 | Glutamate dehydrogenase 1, mitochondrial (GDH 1) (EC 1.4.1.3) | 558 | GLUD1 | 1 | 3 | 5 | 2.8 | 8.9 | 14.1 |
| 325 | -0.98 | 0.21 | 0.49 | E7EQR4 | Ezrin | 586 | EZR | 12 | 8 | 10 | 19.1 | 17.1 | 20.1 |
| 326 | -0.97 | 0.00 | 0.00 | Q96CS3 | FAS-associated factor 2 (Protein ETEA) (UBX domain-containing protein 3B) (UBX domain-containing protein 8) | 445 | FAF2 | 1 | 0 | 0 | 4 | 0 | 0 |
| 327 | -0.97 | 0.00 | 0.00 | P68402 | Platelet-activating factor acetylhydrolase IB subunit beta (EC 3.1.1.47) (PAF acetylhydrolase 30 kDa subunit) (PAF-AH 30 kDa subunit) (PAF-AH subunit beta) (PAFAH subunit beta) | 229 | PAFAH1B2 | 1 | 0 | 0 | 3.9 | 0 | 0 |
| 328 | -0.97 | 0.33 | 2.03 | P51858 | Hepatoma-derived growth factor (HDGF) (High mobility group protein 1-like 2) (HMG-1L2) | 240 | HDGF | 1 | 2 | 3 | 4.2 | 11.2 | 20 |
| 329 | -0.97 | 0.00 | 0.00 | Q96RS6 | NudC domain-containing protein 1 (Chronic myelogenous leukemia tumor antigen 66) (Tumor antigen CML66) | 583 | NUDCD1 | 1 | 0 | 0 | 2.9 | 0 | 0 |
| 330 | -0.96 | 1.09 | 1.28 | E7ENZ3 | T-complex protein 1 subunit epsilon | 486 | CCT5 | 8 | 4 | 7 | 21.2 | 14 | 20.2 |
| 331 | -0.96 | 1.15 | 1.02 | P04843 | Dolichyl-diphosphooligosaccharide--protein glycosyltransferase subunit 1 (EC 2.4.99.18) (Dolichyl-diphosphooligosaccharide--protein glycosyltransferase 67 kDa subunit) (Ribophorin I) (RPN-I) (Ribophorin-1) | 607 | RPN1 | 7 | 4 | 5 | 16.8 | 9.2 | 13.2 |
| 332 | -0.96 | 1.22 | 1.52 | P84095 | Rho-related GTP-binding protein RhoG | 191 | RHOG | 2 | 4 | 1 | 13.1 | 29.8 | 7.3 |
| 333 | -0.96 | 1.19 | 0.91 | P38117 | Electron transfer flavoprotein subunit beta (Beta-ETF) | 255 | ETFB | 4 | 8 | 7 | 18 | 34.1 | 26.7 |
| 334 | -0.95 | 0.51 | 0.70 | P13489 | Ribonuclease inhibitor (Placental ribonuclease inhibitor) (Placental RNase inhibitor) | 461 | RNH1 | 1 | 1 | 2 | 2.4 | 3 | 4.3 |

| | | | | | | | | | | | | | |
|---|---|---|---|---|---|---|---|---|---|---|---|---|---|
| | | | | | (Ribonuclease/angiogenin inhibitor 1) (RAI) | | | | | | | | |
| 335 | -0.94 | 1.30 | 1.27 | Q9P035 | Very-long-chain (3R)-3-hydroxyacyl-CoA dehydratase 3 (EC 4.2.1.134) (3-hydroxyacyl-CoA dehydratase 3) (HACD3) (Butyrate-induced protein 1) (B-ind1) (hB-ind1) (Protein-tyrosine phosphatase-like A domain-containing protein 1) | 362 | HACD3 | 2 | 4 | 1 | 8.8 | 15.5 | 5.8 |
| 336 | -0.94 | 0.60 | 0.24 | C9J3L8 | Translocon-associated protein subunit alpha | 265 | SSR1 | 1 | 2 | 2 | 4.2 | 7.2 | 7.2 |
| 337 | -0.94 | 0.41 | 0.54 | Q13185 | Chromobox protein homolog 3 (HECH) (Heterochromatin protein 1 homolog gamma) (HP1 gamma) (Modifier 2 protein) | 183 | CBX3 | 3 | 1 | 3 | 16.4 | 7.7 | 16.4 |
| 338 | -0.94 | 0.69 | 0.00 | Q8NCW5 | NAD(P)H-hydrate epimerase (EC 5.1.99.6) (Apolipoprotein A-I-binding protein) (AI-BP) (NAD(P)HX epimerase) (YjeF N-terminal domain-containing protein 1) (YjeF_N1) | 288 | APOA1BP | 1 | 2 | 0 | 5.9 | 16.2 | 0 |
| 339 | -0.94 | 0.39 | 1.37 | P26641 | Elongation factor 1-gamma (EF-1-gamma) (eEF-1B gamma) | 437 | EEF1G | 7 | 9 | 8 | 18.3 | 27.5 | 17.2 |
| 340 | -0.94 | 0.99 | 1.42 | P42765 | 3-ketoacyl-CoA thiolase, mitochondrial (EC 2.3.1.16) (Acetyl-CoA acyltransferase) (Beta-ketothiolase) (Mitochondrial 3-oxoacyl-CoA thiolase) (T1) | 397 | ACAA2 | 1 | 1 | 1 | 2.8 | 2.8 | 2.8 |
| 341 | -0.94 | 0.62 | 1.16 | Q9NSD9 | Phenylalanine--tRNA ligase beta subunit (EC 6.1.1.20) (Phenylalanyl-tRNA synthetase beta subunit) (PheRS) | 589 | FARSB | 3 | 4 | 4 | 5.1 | 7 | 7 |
| 342 | -0.93 | 0.18 | 0.60 | P37802 | Transgelin-2 (Epididymis tissue protein Li 7e) (SM22-alpha homolog) | 199 | TAGLN2 | 6 | 11 | 7 | 32.7 | 63.3 | 37.2 |

| | | | | | | | | | | | | | |
|---|---|---|---|---|---|---|---|---|---|---|---|---|---|
| 343 | -0.93 | 0.05 | 1.10 | Q5T8U3 | 60S ribosomal protein L7a (Ribosomal protein L7a) (Fragment) | 191 | RPL7A | 6 | 3 | 4 | 31.4 | 17.3 | 17.8 |
| 344 | -0.93 | 0.79 | 0.54 | Q00610 | Clathrin heavy chain 1 (Clathrin heavy chain on chromosome 17) (CLH-17) | 1675 | CLTC | 18 | 23 | 22 | 15.9 | 21.6 | 19.3 |
| 345 | -0.92 | 1.71 | 1.80 | Q9BSD7 | Cancer-related nucleoside-triphosphatase (NTPase) (EC 3.6.1.15) (Nucleoside triphosphate phosphohydrolase) | 190 | NTPCR | 2 | 3 | 3 | 15.8 | 27.9 | 23.7 |
| 346 | -0.92 | 0.00 | 1.72 | B0QYA5 | Eukaryotic translation initiation factor 3 subunit D (Fragment) | 269 | EIF3D | 2 | 0 | 1 | 9.7 | 0 | 4.5 |
| 347 | -0.92 | -0.17 | 1.00 | P62269 | 40S ribosomal protein S18 (Ke-3) (Ke3) | 152 | RPS18 | 4 | 4 | 4 | 19.1 | 19.7 | 19.1 |
| 348 | -0.92 | 0.00 | 0.47 | K7EML9 | Thioredoxin-like protein 1 (Fragment) | 138 | TXNL1 | 2 | 0 | 1 | 18.8 | 0 | 10.9 |
| 349 | -0.92 | 0.00 | 0.00 | Q96JB2 | Conserved oligomeric Golgi complex subunit 3 (COG complex subunit 3) (Component of oligomeric Golgi complex 3) (Vesicle-docking protein SEC34 homolog) (p94) | 828 | COG3 | 1 | 0 | 0 | 3.4 | 0 | 0 |
| 350 | -0.92 | 0.45 | 0.80 | Q92598 | Heat shock protein 105 kDa (Antigen NY-CO-25) (Heat shock 110 kDa protein) | 858 | HSPH1 | 7 | 6 | 9 | 12.8 | 9.5 | 15.2 |
| 351 | -0.92 | 0.72 | 0.09 | P51148 | Ras-related protein Rab-5C (L1880) (RAB5L) | 216 | RAB5C | 5 | 4 | 2 | 28.2 | 23.6 | 11.6 |
| 352 | -0.92 | 0.00 | 0.00 | Q9Y3C6 | Peptidyl-prolyl cis-trans isomerase-like 1 (PPIase) (EC 5.2.1.8) (Rotamase PPIL1) | 166 | PPIL1 | 1 | 0 | 0 | 10.2 | 0 | 0 |
| 353 | -0.91 | 4.52 | 0.00 | Q9BQ61 | Uncharacterized protein C19orf43 | 176 | C19orf43 | 1 | 1 | 0 | 7.4 | 13.6 | 0 |
| 354 | -0.91 | 0.00 | 0.00 | F5GXQ0 | BRO1 domain-containing protein BROX | 338 | BROX | 1 | 0 | 0 | 5.6 | 0 | 0 |
| 355 | -0.91 | 0.30 | 0.94 | A0A087WTT1 | Polyadenylate-binding protein (PABP) | 522 | PABPC1 | 7 | 5 | 10 | 17.4 | 12.6 | 24.1 |
| 356 | -0.91 | 0.00 | 0.92 | Q6NZI2 | Polymerase I and transcript release factor (Cavin-1) | 390 | PTRF | 1 | 0 | 1 | 5 | 0 | 5 |

| | | | | | | | | | | | | | |
|---|---|---|---|---|---|---|---|---|---|---|---|---|---|
| 357 | -0.90 | 0.00 | 0.20 | P54577 | Tyrosine--tRNA ligase, cytoplasmic (EC 6.1.1.1) (Tyrosyl-tRNA synthetase) (TyrRS) [Cleaved into: Tyrosine--tRNA ligase, cytoplasmic, N-terminally processed] | 528 | YARS | 1 | 0 | 1 | 2.3 | 0 | 2.3 |
| 358 | -0.90 | 1.01 | 0.54 | Q15021 | Condensin complex subunit 1 (Chromosome condensation-related SMC-associated protein 1) (Chromosome-associated protein D2) (hCAP-D2) (Non-SMC condensin I complex subunit D2) (XCAP-D2 homolog) | 1401 | NCAPD2 | 3 | 2 | 1 | 2.4 | 1.6 | 0.9 |
| 359 | -0.90 | 0.00 | 0.90 | Q01085 | Nucleolysin TIAR (TIA-1-related protein) | 375 | TIAL1 | 1 | 0 | 1 | 3.5 | 0 | 8.3 |
| 360 | -0.90 | 0.73 | 0.28 | F8W1R7 | Myosin light polypeptide 6 | 145 | MYL6 | 2 | 2 | 4 | 17.2 | 17.2 | 34.5 |
| 361 | -0.89 | 1.15 | 1.24 | P06576 | ATP synthase subunit beta, mitochondrial (EC 3.6.3.14) | 529 | ATP5B | 10 | 12 | 9 | 25.5 | 29.7 | 22.5 |
| 362 | -0.89 | 0.10 | 1.12 | P62266 | 40S ribosomal protein S23 | 143 | RPS23 | 2 | 3 | 2 | 15.4 | 21 | 15.4 |
| 363 | -0.89 | 0.54 | 0.99 | P30837 | Aldehyde dehydrogenase X, mitochondrial (EC 1.2.1.3) (Aldehyde dehydrogenase 5) (Aldehyde dehydrogenase family 1 member B1) | 517 | ALDH1B1 | 5 | 2 | 3 | 12.4 | 4.4 | 6.2 |
| 364 | -0.89 | 0.00 | 1.09 | Q5T5U7 | Selenide, water dikinase 1 (Fragment) | 91 | SEPHS1 | 1 | 0 | 1 | 11 | 0 | 11 |
| 365 | -0.89 | 0.99 | 1.04 | P13804 | Electron transfer flavoprotein subunit alpha, mitochondrial (Alpha-ETF) | 333 | ETFA | 7 | 8 | 6 | 31.2 | 36.9 | 22.2 |
| 366 | -0.89 | 0.00 | 0.66 | F2Z3P2 | Uridine 5'-monophosphate synthase | 57 | UMPS | 1 | 0 | 1 | 29.8 | 0 | 29.8 |
| 367 | -0.89 | -0.15 | 0.20 | P09382 | Galectin-1 (Gal-1) (14 kDa laminin-binding protein) (HLBP14) (14 kDa lectin) (Beta-galactoside-binding lectin L-14-I) (Galaptin) (HBL) (HPL) (Lactose-binding lectin 1) (Lectin galactoside-binding soluble 1) (Putative MAPK-activating protein PM12) (S-Lac lectin 1) | 135 | LGALS1 | 5 | 6 | 4 | 48.1 | 57 | 34.8 |

| | | | | | | | | | | | | | |
|---|---|---|---|---|---|---|---|---|---|---|---|---|---|
| 368 | -0.88 | 0.00 | 0.00 | J3QRD6 | 60S ribosome subunit biogenesis protein NIP7 homolog (Fragment) | 70 | NIP7 | 1 | 0 | 0 | 20 | 0 | 0 |
| 369 | -0.88 | 0.73 | 1.06 | A0A0A0 MSS8 | Aldo-keto reductase family 1 member C3 | 323 | AKR1C3 | 7 | 6 | 7 | 33.4 | 22.9 | 26 |
| 370 | -0.88 | 0.70 | -0.67 | I3L4X2 | Multidrug resistance-associated protein 1 (Fragment) | 1440 | ABCC1 | 2 | 6 | 3 | 2 | 5.8 | 2.6 |
| 371 | -0.88 | 0.00 | 0.87 | P54886 | Delta-1-pyrroline-5-carboxylate synthase (P5CS) (Aldehyde dehydrogenase family 18 member A1) [Includes: Glutamate 5-kinase (GK) (EC 2.7.2.11) (Gamma-glutamyl kinase); Gamma-glutamyl phosphate reductase (GPR) (EC 1.2.1.41) (Glutamate-5-semialdehyde dehydrogenase) (Glutamyl-gamma-semialdehyde dehydrogenase)] | 795 | ALDH18A1 | 2 | 0 | 3 | 3.2 | 0 | 5.2 |
| 372 | -0.88 | 1.16 | 0.97 | P42704 | Leucine-rich PPR motif-containing protein, mitochondrial (130 kDa leucine-rich protein) (LRP 130) (GP130) | 1394 | LRPPRC | 36 | 31 | 35 | 32.4 | 27.9 | 31.3 |
| 373 | -0.88 | 0.93 | 0.98 | O43809 | Cleavage and polyadenylation specificity factor subunit 5 (Cleavage and polyadenylation specificity factor 25 kDa subunit) (CFIm25) (CPSF 25 kDa subunit) (Nucleoside diphosphate-linked moiety X motif 21) (Nudix motif 21) (Pre-mRNA cleavage factor Im 25 kDa subunit) | 227 | NUDT21 | 5 | 4 | 4 | 24.2 | 30 | 18.9 |
| 374 | -0.87 | 1.70 | 0.00 | P11441 | Ubiquitin-like protein 4A (Ubiquitin-like protein GDX) | 157 | UBL4A | 1 | 1 | 0 | 6.4 | 9.6 | 0 |
| 375 | -0.87 | 0.00 | 1.40 | P51991 | Heterogeneous nuclear ribonucleoprotein A3 (hnRNP A3) | 378 | HNRNPA3 | 2 | 0 | 3 | 7.9 | 0 | 10.7 |
| 376 | -0.87 | 0.40 | 1.94 | P62805 | Histone H4 | 103 | HIST1H4A | 3 | 3 | 3 | 22.3 | 28.2 | 22.3 |

| | | | | | | | | | | | | | |
|---|---|---|---|---|---|---|---|---|---|---|---|---|---|
| 377 | -0.87 | -0.07 | 1.19 | P62847 | 40S ribosomal protein S24 | 133 | RPS24 | 4 | 2 | 2 | 30 | 20.8 | 20.8 |
| 378 | -0.87 | 1.37 | 0.91 | P10809 | 60 kDa heat shock protein, mitochondrial (60 kDa chaperonin) (Chaperonin 60) (CPN60) (Heat shock protein 60) (HSP-60) (Hsp60) (HuCHA60) (Mitochondrial matrix protein P1) (P60 lymphocyte protein) | 573 | HSPD1 | 25 | 20 | 26 | 45.4 | 41.7 | 39.1 |
| 379 | -0.87 | 0.69 | 0.52 | P50991 | T-complex protein 1 subunit delta (TCP-1-delta) (CCT-delta) (Stimulator of TAR RNA-binding) | 539 | CCT4 | 8 | 6 | 5 | 20.6 | 14.3 | 11.2 |
| 380 | -0.87 | 0.63 | 0.87 | P46821 | Microtubule-associated protein 1B (MAP-1B) [Cleaved into: MAP1B heavy chain; MAP1 light chain LC1] | 2468 | MAP1B | 4 | 5 | 3 | 2.2 | 2.8 | 1.7 |
| 381 | -0.87 | 0.00 | 0.81 | P38919 | Eukaryotic initiation factor 4A-III (eIF-4A-III) (eIF4A-III) (EC 3.6.4.13) (ATP-dependent RNA helicase DDX48) (ATP-dependent RNA helicase eIF4A-3) (DEAD box protein 48) (Eukaryotic initiation factor 4A-like NUK-34) (Eukaryotic translation initiation factor 4A isoform 3) (Nuclear matrix protein 265) (NMP 265) (hNMP 265) [Cleaved into: Eukaryotic initiation factor 4A-III, N-terminally processed] | 411 | EIF4A3 | 2 | 2 | 1 | 5.6 | 5.8 | 2.2 |
| 382 | -0.86 | 1.69 | 0.73 | F8W1A4 | Adenylate kinase 2, mitochondrial (AK 2) (EC 2.7.4.3) (ATP-AMP transphosphorylase 2) (ATP:AMP phosphotransferase) (Adenylate monophosphate kinase) | 232 | AK2 | 6 | 6 | 5 | 37.9 | 39.2 | 30.2 |

| | | | | | | | | | | | | | |
|---|---|---|---|---|---|---|---|---|---|---|---|---|---|
| 383 | -0.86 | 0.55 | 0.58 | P12956 | X-ray repair cross-complementing protein 6 (EC 3.6.4.-) (EC 4.2.99.-) (5'-deoxyribose-5-phosphate lyase Ku70) (5'-dRP lyase Ku70) (70 kDa subunit of Ku antigen) (ATP-dependent DNA helicase 2 subunit 1) (ATP-dependent DNA helicase II 70 kDa subunit) (CTC box-binding factor 75 kDa subunit) (CTC75) (CTCBF) (DNA repair protein XRCC6) (Lupus Ku autoantigen protein p70) (Ku70) (Thyroid-lupus autoantigen) (TLAA) (X-ray repair complementing defective repair in Chinese hamster cells 6) | 609 | XRCC6 | 13 | 9 | 10 | 30.4 | 17.2 | 19.9 |
| 384 | -0.86 | 0.66 | 0.00 | Q5VVW2 | GTPase-activating Rap/Ran-GAP domain-like protein 3 | 1013 | GARNL3 | 1 | 1 | 0 | 2.6 | 2.6 | 0 |
| 385 | -0.86 | 0.67 | 0.62 | P55072 | Transitional endoplasmic reticulum ATPase (TER ATPase) (EC 3.6.4.6) (15S Mg(2+)-ATPase p97 subunit) (Valosin-containing protein) (VCP) | 806 | VCP | 12 | 15 | 17 | 19.7 | 26.3 | 27.3 |
| 386 | -0.86 | 0.00 | 0.00 | D6RDJ3 | DNA-directed RNA polymerases I and III subunit RPAC1 (Fragment) | 124 | POLR1C | 1 | 0 | 0 | 10.5 | 0 | 0 |
| 387 | -0.86 | 1.00 | 1.23 | P13073 | Cytochrome c oxidase subunit 4 isoform 1, mitochondrial (Cytochrome c oxidase polypeptide IV) (Cytochrome c oxidase subunit IV isoform 1) (COX IV-1) | 169 | COX4I1 | 3 | 5 | 2 | 13.6 | 30.2 | 12.4 |

| | | | | | | | | | | | | | |
|---|---|---|---|---|---|---|---|---|---|---|---|---|---|
| 388 | -0.86 | - 0.11 | 0.63 | P30086 | Phosphatidylethanolamine-binding protein 1 (PEBP-1) (HCNPpp) (Neuropolypeptide h3) (Prostatic-binding protein) (Raf kinase inhibitor protein) (RKIP) [Cleaved into: Hippocampal cholinergic neurostimulating peptide (HCNP)] | 187 | PEBP1 | 2 | 4 | 2 | 8.6 | 24.6 | 8.6 |
| 389 | -0.85 | 1.12 | 1.08 | O75489 | NADH dehydrogenase [ubiquinone] iron-sulfur protein 3, mitochondrial (EC 1.6.5.3) (EC 1.6.99.3) (Complex I-30kD) (CI-30kD) (NADH-ubiquinone oxidoreductase 30 kDa subunit) | 264 | NDUFS3 | 1 | 5 | 1 | 6.4 | 26.1 | 4.9 |
| 390 | -0.85 | 0.00 | 1.35 | Q9Y4Z0 | U6 snRNA-associated Sm-like protein LSm4 (Glycine-rich protein) (GRP) | 139 | LSM4 | 1 | 0 | 3 | 5.8 | 0 | 21.6 |
| 391 | -0.85 | 0.06 | 1.08 | P62280 | 40S ribosomal protein S11 | 158 | RPS11 | 4 | 6 | 3 | 17.1 | 35.4 | 19 |
| 392 | -0.85 | 0.00 | 0.00 | Q8N1F7 | Nuclear pore complex protein Nup93 (93 kDa nucleoporin) (Nucleoporin Nup93) | 819 | NUP93 | 2 | 0 | 0 | 3.3 | 0 | 0 |
| 393 | -0.84 | 0.76 | 0.60 | P61978 | Heterogeneous nuclear ribonucleoprotein K (hnRNP K) (Transformation up-regulated nuclear protein) (TUNP) | 463 | HNRNPK | 8 | 10 | 7 | 21.6 | 32.5 | 24.8 |
| 394 | -0.84 | 0.00 | 1.23 | A0A087X 1S2 | Nuclease-sensitive element-binding protein 1 | 314 | YBX1 | 1 | 0 | 3 | 4.8 | 0 | 16.2 |
| 395 | -0.84 | 0.36 | 0.03 | O00303 | Eukaryotic translation initiation factor 3 subunit F (eIF3f) (Deubiquitinating enzyme eIF3f) (EC 3.4.19.12) (Eukaryotic translation initiation factor 3 subunit 5) (eIF-3-epsilon) (eIF3 p47) | 357 | EIF3F | 4 | 1 | 3 | 13.7 | 3.1 | 11.8 |
| 396 | -0.83 | - 0.41 | 0.73 | P42766 | 60S ribosomal protein L35 | 123 | RPL35 | 1 | 2 | 1 | 8.1 | 15.4 | 8.1 |

| | | | | | | | | | | | | | |
|---|---|---|---|---|---|---|---|---|---|---|---|---|---|
| 397 | -0.83 | 0.93 | 0.80 | P39656 | Dolichyl-diphosphooligosaccharide--protein glycosyltransferase 48 kDa subunit (DDOST 48 kDa subunit) (Oligosaccharyl transferase 48 kDa subunit) (EC 2.4.99.18) | 456 | DDOST | 2 | 2 | 2 | 5 | 4.6 | 4.1 |
| 398 | -0.83 | 1.18 | 0.02 | C9JFR7 | Cytochrome c (Fragment) | 101 | CYCS | 1 | 3 | 2 | 14.9 | 22.8 | 21.8 |
| 399 | -0.83 | 0.53 | 1.19 | P62829 | 60S ribosomal protein L23 (60S ribosomal protein L17) | 140 | RPL23 | 4 | 3 | 2 | 37.9 | 27.1 | 20 |
| 400 | -0.83 | -0.57 | 1.25 | Q15435 | Protein phosphatase 1 regulatory subunit 7 (Protein phosphatase 1 regulatory subunit 22) | 360 | PPP1R7 | 1 | 1 | 1 | 8.1 | 8.1 | 8.1 |
| 401 | -0.83 | 0.39 | 0.33 | Q92841 | Probable ATP-dependent RNA helicase DDX17 (EC 3.6.4.13) (DEAD box protein 17) (DEAD box protein p72) (RNA-dependent helicase p72) | 729 | DDX17 | 4 | 1 | 3 | 6.6 | 2 | 4.6 |
| 402 | -0.83 | 0.18 | 1.46 | E7EPB3 | 60S ribosomal protein L14 | 124 | RPL14 | 2 | 2 | 2 | 17.7 | 17.7 | 17.7 |
| 403 | -0.83 | 0.00 | 0.00 | O00203 | AP-3 complex subunit beta-1 (Adaptor protein complex AP-3 subunit beta-1) (Adaptor-related protein complex 3 subunit beta-1) (Beta-3A-adaptin) (Clathrin assembly protein complex 3 beta-1 large chain) | 1094 | AP3B1 | 1 | 0 | 0 | 1.1 | 0 | 0 |
| 404 | -0.82 | 0.57 | 1.26 | Q7L014 | Probable ATP-dependent RNA helicase DDX46 (EC 3.6.4.13) (DEAD box protein 46) (PRP5 homolog) | 1031 | DDX46 | 2 | 2 | 2 | 2.2 | 1.8 | 2 |
| 405 | -0.82 | 0.00 | 0.28 | F5H7R9 | Parathymosin (Fragment) | 57 | PTMS | 1 | 0 | 1 | 19.3 | 0 | 19.3 |
| 406 | -0.82 | 0.00 | 1.23 | P41091 | Eukaryotic translation initiation factor 2 subunit 3 (Eukaryotic translation initiation factor 2 subunit gamma X) (eIF-2-gamma X) (eIF-2gX) | 472 | EIF2S3 | 1 | 0 | 3 | 3.4 | 0 | 9.3 |
| 407 | -0.82 | 1.09 | 0.94 | Q5QP23 | RNA-binding protein 39 (Fragment) | 231 | RBM39 | 1 | 1 | 1 | 6.5 | 6.5 | 6.5 |
| 408 | -0.82 | -0.39 | 0.34 | Q9Y281 | Cofilin-2 (Cofilin, muscle isoform) | 166 | CFL2 | 4 | 4 | 5 | 34.3 | 36.1 | 32.5 |

| | | | | | | | | | | | | | |
|---|---|---|---|---|---|---|---|---|---|---|---|---|---|
| 409 | -0.82 | 1.00 | 0.00 | Q9UMS0 | NFU1 iron-sulfur cluster scaffold homolog, mitochondrial (HIRA-interacting protein 5) | 254 | NFU1 | 1 | 3 | 0 | 4.3 | 17 | 0 |
| 410 | -0.82 | 0.85 | 0.69 | P45880 | Voltage-dependent anion-selective channel protein 2 (VDAC-2) (hVDAC2) (Outer mitochondrial membrane protein porin 2) | 294 | VDAC2 | 3 | 5 | 6 | 17.3 | 20.5 | 30.4 |
| 411 | -0.81 | 0.00 | 0.00 | G3V4D9 | Nuclear export mediator factor NEMF (Fragment) | 178 | NEMF | 1 | 0 | 0 | 5.6 | 0 | 0 |
| 412 | -0.81 | 1.66 | 0.00 | A0A087 WTZ2 | Ran-binding protein 17 | 518 | RANBP17 | 1 | 2 | 0 | 2.1 | 3.9 | 0 |
| 413 | -0.81 | 0.78 | 0.67 | Q9UIJ7 | GTP:AMP phosphotransferase AK3, mitochondrial (EC 2.7.4.10) (Adenylate kinase 3) (AK 3) (Adenylate kinase 3 alpha-like 1) | 227 | AK3 | 2 | 5 | 3 | 11 | 26 | 14.5 |
| 414 | -0.81 | - 0.09 | 0.06 | P13797 | Plastin-3 (T-plastin) | 630 | PLS3 | 9 | 5 | 6 | 17.6 | 11.4 | 9.5 |
| 415 | -0.81 | 0.57 | 0.41 | P11142 | Heat shock cognate 71 kDa protein (Heat shock 70 kDa protein 8) (Lipopolysaccharide-associated protein 1) (LAP-1) (LPS-associated protein 1) | 646 | HSPA8 | 18 | 15 | 15 | 34.8 | 31.6 | 28.3 |
| 416 | -0.80 | 1.10 | 0.57 | F5GY37 | Prohibitin-2 | 267 | PHB2 | 3 | 5 | 2 | 11.2 | 22.8 | 7.1 |
| 417 | -0.80 | 0.87 | 0.57 | Q60FE5 | Filamin A (Filamin-A) | 2620 | FLNA | 25 | 26 | 28 | 13.5 | 14.2 | 14.8 |
| 418 | -0.80 | 0.03 | 0.55 | A0A0A0 MTR7 | E3 ubiquitin-protein ligase RNF213 | 5207 | RNF213 | 2 | 1 | 4 | 0.5 | 0.3 | 0.9 |
| 419 | -0.80 | 0.48 | 0.13 | E7EQ69 | N-alpha-acetyltransferase 50 | 168 | NAA50 | 5 | 3 | 6 | 28.6 | 19.6 | 28.6 |
| 420 | -0.80 | 0.30 | 0.62 | Q92616 | Translational activator GCN1 (HsGCN1) (GCN1-like protein 1) | 2671 | GCN1L1 | 8 | 9 | 17 | 4.3 | 4.3 | 8.8 |
| 421 | -0.80 | - 0.05 | 0.64 | O00571 | ATP-dependent RNA helicase DDX3X (EC 3.6.4.13) (DEAD box protein 3, X-chromosomal) (DEAD box, X isoform) (Helicase-like protein 2) (HLP2) | 662 | DDX3X | 6 | 2 | 4 | 11 | 4.5 | 7.1 |

| | | | | | | | | | | | | | |
|---|---|---|---|---|---|---|---|---|---|---|---|---|---|
| 422 | -0.79 | 1.29 | 0.93 | O95292 | Vesicle-associated membrane protein-associated protein B/C (VAMP-B/VAMP-C) (VAMP-associated protein B/C) (VAP-B/VAP-C) | 243 | VAPB | 3 | 3 | 4 | 16.9 | 16 | 22.6 |
| 423 | -0.79 | -0.66 | 0.64 | Q9NR33 | DNA polymerase epsilon subunit 4 (EC 2.7.7.7) (DNA polymerase II subunit 4) (DNA polymerase epsilon subunit p12) | 117 | POLE4 | 1 | 1 | 1 | 9.4 | 9.4 | 9.4 |
| 424 | -0.79 | -0.27 | 0.00 | P17931 | Galectin-3 (Gal-3) (35 kDa lectin) (Carbohydrate-binding protein 35) (CBP 35) (Galactose-specific lectin 3) (Galactoside-binding protein) (GALBP) (IgE-binding protein) (L-31) (Laminin-binding protein) (Lectin L-29) (Mac-2 antigen) | 250 | LGALS3 | 2 | 1 | 0 | 8.8 | 4.4 | 0 |
| 425 | -0.79 | 0.00 | 0.00 | M0R253 | Deoxyhypusine synthase | 132 | DHPS | 1 | 0 | 0 | 9.1 | 0 | 0 |
| 426 | -0.79 | 0.00 | 0.81 | Q15404 | Ras suppressor protein 1 (RSP-1) (Rsu-1) | 277 | RSU1 | 2 | 0 | 3 | 8 | 0 | 16.1 |
| 427 | -0.79 | 1.54 | 1.47 | B4DY09 | Interleukin enhancer-binding factor 2 (cDNA FLJ51660, highly similar to Interleukin enhancer-binding factor 2) | 352 | ILF2 | 1 | 1 | 2 | 5.7 | 4.5 | 8.5 |
| 428 | -0.79 | 0.08 | 0.78 | P61970 | Nuclear transport factor 2 (NTF-2) (Placental protein 15) (PP15) | 127 | NUTF2 | 1 | 2 | 1 | 6.3 | 11.8 | 12.6 |
| 429 | -0.78 | 1.28 | 0.31 | P05387 | 60S acidic ribosomal protein P2 (Renal carcinoma antigen NY-REN-44) | 115 | RPLP2 | 2 | 4 | 1 | 39.1 | 62.6 | 10.4 |
| 430 | -0.78 | 0.00 | 0.39 | F8WEG8 | Interferon-inducible double-stranded RNA-dependent protein kinase activator A | 95 | PRKRA | 1 | 0 | 1 | 10.5 | 0 | 10.5 |
| 431 | -0.78 | 0.00 | 0.78 | Q12765 | Secernin-1 | 414 | SCRN1 | 1 | 0 | 1 | 3.8 | 0 | 3.8 |
| 432 | -0.78 | 0.00 | 0.00 | Q96S52 | GPI transamidase component PIG-S (Phosphatidylinositol-glycan biosynthesis class S protein) | 555 | PIGS | 1 | 0 | 0 | 2 | 0 | 0 |

| | | | | | | | | | | | | | |
|---|---|---|---|---|---|---|---|---|---|---|---|---|---|
| 433 | -0.78 | 0.68 | 0.52 | P08107 | Heat shock 70 kDa protein 1A/1B (Heat shock 70 kDa protein 1/2) (HSP70-1/HSP70-2) (HSP70.1/HSP70.2) | 641 | HSPA1A | 11 | 11 | 8 | 21.1 | 25.3 | 13.7 |
| 434 | -0.78 | 0.00 | 0.81 | B4DQI4 | Alpha/beta hydrolase domain-containing protein 14B (cDNA FLJ52723, highly similar to Abhydrolase domain-containing protein 14B) | 188 | ABHD14B | 1 | 0 | 1 | 6.9 | 0 | 6.9 |
| 435 | -0.78 | 0.00 | 0.11 | H0YF12 | Tyrosine-protein phosphatase non-receptor type 11 (Fragment) | 108 | PTPN11 | 1 | 0 | 1 | 13 | 0 | 13 |
| 436 | -0.77 | -0.02 | 0.36 | P42224 | Signal transducer and activator of transcription 1-alpha/beta (Transcription factor ISGF-3 components p91/p84) | 750 | STAT1 | 3 | 1 | 2 | 6.2 | 1.7 | 3.5 |
| 437 | -0.77 | 0.00 | 0.00 | Q08J23 | tRNA (cytosine(34)-C(5))-methyltransferase (EC 2.1.1.203) (Myc-induced SUN domain-containing protein) (Misu) (NOL1/NOP2/Sun domain family member 2) (Substrate of AIM1/Aurora kinase B) (tRNA (cytosine-5-)-methyltransferase) (tRNA methyltransferase 4 homolog) (hTrm4) | 767 | NSUN2 | 1 | 0 | 0 | 2.1 | 0 | 0 |
| 438 | -0.77 | 0.15 | 0.27 | P60228 | Eukaryotic translation initiation factor 3 subunit E (eIF3e) (Eukaryotic translation initiation factor 3 subunit 6) (Viral integration site protein INT-6 homolog) (eIF-3 p48) | 445 | EIF3E | 5 | 2 | 3 | 11.7 | 4.7 | 6.3 |
| 439 | -0.77 | 0.00 | 0.85 | A0A087 WUT6 | Eukaryotic translation initiation factor 5B | 1220 | EIF5B | 1 | 0 | 1 | 0.7 | 0 | 0.9 |
| 440 | -0.77 | 0.00 | 0.00 | X6RDA4 | Paraspeckle component 1 (Fragment) | 248 | PSPC1 | 1 | 0 | 0 | 4.4 | 0 | 0 |

| | | | | | | | | | | | | | |
|---|---|---|---|---|---|---|---|---|---|---|---|---|---|
| 441 | -0.77 | 1.22 | 1.19 | O75369 | Filamin-B (FLN-B) (ABP-278) (ABP-280 homolog) (Actin-binding-like protein) (Beta-filamin) (Filamin homolog 1) (Fh1) (Filamin-3) (Thyroid autoantigen) (Truncated actin-binding protein) (Truncated ABP) | 2602 | FLNB | 19 | 43 | 34 | 11.3 | 23.1 | 19.4 |
| 442 | -0.76 | 0.48 | 0.84 | P34932 | Heat shock 70 kDa protein 4 (HSP70RY) (Heat shock 70-related protein APG-2) | 840 | HSPA4 | 10 | 10 | 10 | 15.6 | 15.4 | 15.4 |
| 443 | -0.76 | 0.08 | 0.74 | O76003 | Glutaredoxin-3 (PKC-interacting cousin of thioredoxin) (PICOT) (PKC-theta-interacting protein) (PKCq-interacting protein) (Thioredoxin-like protein 2) | 335 | GLRX3 | 6 | 2 | 5 | 20.6 | 6.6 | 17.6 |
| 444 | -0.76 | -0.17 | 0.92 | P61758 | Prefoldin subunit 3 (HIBBJ46) (Von Hippel-Lindau-binding protein 1) (VBP-1) (VHL-binding protein 1) | 197 | VBP1 | 4 | 3 | 4 | 23.9 | 18.8 | 23.9 |
| 445 | -0.76 | 0.05 | 0.89 | O43242 | 26S proteasome non-ATPase regulatory subunit 3 (26S proteasome regulatory subunit RPN3) (26S proteasome regulatory subunit S3) (Proteasome subunit p58) | 534 | PSMD3 | 2 | 1 | 3 | 6.7 | 3.4 | 10.7 |
| 446 | -0.76 | 0.57 | -3.51 | P31937 | 3-hydroxyisobutyrate dehydrogenase, mitochondrial (HIBADH) (EC 1.1.1.31) | 336 | HIBADH | 2 | 4 | 2 | 8 | 14.6 | 8.9 |
| 447 | -0.76 | 1.22 | 0.14 | O75323 | Protein NipSnap homolog 2 (NipSnap2) (Glioblastoma-amplified sequence) | 286 | GBAS | 1 | 2 | 2 | 5.6 | 8.7 | 8 |
| 448 | -0.76 | 0.47 | 1.01 | H0Y3Y4 | Septin-7 (Fragment) | 373 | sept7 | 1 | 1 | 2 | 2.4 | 2.4 | 7 |
| 449 | -0.76 | 0.00 | 0.00 | Q6P1M0 | Long-chain fatty acid transport protein 4 (FATP-4) (Fatty acid transport protein 4) (EC 6.2.1.-) (Solute carrier family 27 member 4) | 643 | SLC27A4 | 1 | 0 | 0 | 1.7 | 0 | 0 |

| | | | | | | | | | | | | | |
|---|---|---|---|---|---|---|---|---|---|---|---|---|---|
| 450 | -0.75 | 0.00 | 0.00 | Q12904 | Aminoacyl tRNA synthase complex-interacting multifunctional protein 1 (Multisynthase complex auxiliary component p43) [Cleaved into: Endothelial monocyte-activating polypeptide 2 (EMAP-2) (Endothelial monocyte-activating polypeptide II) (EMAP-II) (Small inducible cytokine subfamily E member 1)] | 312 | AIMP1 | 1 | 0 | 0 | 5.8 | 0 | 0 |
| 451 | -0.75 | 0.00 | 0.68 | P50995 | Annexin A11 (56 kDa autoantigen) (Annexin XI) (Annexin-11) (Calcyclin-associated annexin 50) (CAP-50) | 505 | ANXA11 | 1 | 0 | 2 | 2.3 | 0 | 5.5 |
| 452 | -0.75 | 1.45 | -0.07 | P21964 | Catechol O-methyltransferase (EC 2.1.1.6) | 271 | COMT | 1 | 2 | 1 | 7.2 | 17.2 | 7.2 |
| 453 | -0.75 | -0.35 | -0.10 | Q01469 | Fatty acid-binding protein, epidermal (Epidermal-type fatty acid-binding protein) (E-FABP) (Fatty acid-binding protein 5) (Psoriasis-associated fatty acid-binding protein homolog) (PA-FABP) | 135 | FABP5 | 1 | 4 | 1 | 8.1 | 31.1 | 8.1 |
| 454 | -0.75 | 1.18 | 1.26 | Q9H7Z7 | Prostaglandin E synthase 2 (Membrane-associated prostaglandin E synthase-2) (mPGE synthase-2) (Microsomal prostaglandin E synthase 2) (mPGES-2) (Prostaglandin-H(2) E-isomerase) (EC 5.3.99.3) [Cleaved into: Prostaglandin E synthase 2 truncated form] | 377 | PTGES2 | 3 | 2 | 5 | 11.1 | 10.9 | 24.1 |
| 455 | -0.75 | 0.31 | 0.83 | P55010 | Eukaryotic translation initiation factor 5 (eIF-5) | 431 | EIF5 | 2 | 1 | 3 | 5.6 | 2.1 | 8.6 |
| 456 | -0.75 | 0.00 | 2.51 | X1WI28 | 60S ribosomal protein L10 (Fragment) | 200 | RPL10 | 1 | 0 | 3 | 6.5 | 0 | 21 |
| 457 | -0.75 | 0.73 | 2.36 | P10412 | Histone H1.4 (Histone H1b) (Histone H1s-4) | 219 | HIST1H1E | 5 | 5 | 5 | 15.5 | 22.4 | 15.1 |

| | | | | | | | | | | | | | |
|---|---|---|---|---|---|---|---|---|---|---|---|---|---|
| 458 | -0.74 | 0.97 | 1.11 | P62316 | Small nuclear ribonucleoprotein Sm D2 (Sm-D2) (snRNP core protein D2) | 118 | SNRPD2 | 4 | 4 | 4 | 28.8 | 36.4 | 28.8 |
| 459 | -0.74 | 1.45 | 0.95 | Q9P0L0 | Vesicle-associated membrane protein-associated protein A (VAMP-A) (VAMP-associated protein A) (VAP-A) (33 kDa VAMP-associated protein) (VAP-33) | 249 | VAPA | 2 | 4 | 2 | 5.6 | 17.7 | 5.6 |
| 460 | -0.74 | 0.00 | 1.32 | O95831 | Apoptosis-inducing factor 1, mitochondrial (EC 1.1.1.-) (Programmed cell death protein 8) | 613 | AIFM1 | 1 | 0 | 1 | 2.6 | 0 | 2.6 |
| 461 | -0.74 | -0.28 | 0.24 | Q99471 | Prefoldin subunit 5 (C-Myc-binding protein Mm-1) (Myc modulator 1) | 154 | PFDN5 | 3 | 3 | 2 | 31.2 | 21.4 | 21.4 |
| 462 | -0.74 | 3.08 | 1.04 | P53597 | Succinyl-CoA ligase [ADP/GDP-forming] subunit alpha, mitochondrial (EC 6.2.1.4) (EC 6.2.1.5) (Succinyl-CoA synthetase subunit alpha) (SCS-alpha) | 346 | SUCLG1 | 1 | 1 | 1 | 2.9 | 4.6 | 2.9 |
| 463 | -0.74 | 1.61 | 3.10 | Q99878 | Histone H2A type 1-J (Histone H2A/e) | 128 | HIST1H2AJ | 1 | 3 | 2 | 8.6 | 27.3 | 23.4 |
| 464 | -0.73 | 0.59 | 0.53 | Q9HAV4 | Exportin-5 (Exp5) (Ran-binding protein 21) | 1204 | XPO5 | 2 | 2 | 2 | 2.1 | 2.1 | 2.1 |
| 465 | -0.73 | 0.61 | 0.46 | P23919 | Thymidylate kinase (EC 2.7.4.9) (dTMP kinase) | 212 | DTYMK | 2 | 4 | 2 | 9.4 | 26.9 | 9.4 |
| 466 | -0.72 | -0.04 | 0.61 | P07195 | L-lactate dehydrogenase B chain (LDH-B) (EC 1.1.1.27) (LDH heart subunit) (LDH-H) (Renal carcinoma antigen NY-REN-46) | 334 | LDHB | 13 | 13 | 12 | 40.7 | 40.7 | 39.5 |
| 467 | -0.72 | 1.47 | 1.25 | E5RJU9 | Protein LYRIC | 526 | MTDH | 1 | 2 | 1 | 2.3 | 6.1 | 2.3 |
| 468 | -0.72 | 0.38 | 0.74 | O75396 | Vesicle-trafficking protein SEC22b (ER-Golgi SNARE of 24 kDa) (ERS-24) (ERS24) (SEC22 vesicle-trafficking protein homolog B) (SEC22 vesicle-trafficking protein-like 1) | 215 | SEC22B | 2 | 3 | 2 | 13.5 | 18.1 | 13.5 |

| | | | | | | | | | | | | | |
|---|---|---|---|---|---|---|---|---|---|---|---|---|---|
| 469 | -0.72 | 1.05 | 1.37 | O75533 | Splicing factor 3B subunit 1 (Pre-mRNA-splicing factor SF3b 155 kDa subunit) (SF3b155) (Spliceosome-associated protein 155) (SAP 155) | 1304 | SF3B1 | 4 | 5 | 5 | 3.9 | 5.1 | 5.7 |
| 470 | -0.71 | 0.27 | 0.81 | Q06830 | Peroxiredoxin-1 (EC 1.11.1.15) (Natural killer cell-enhancing factor A) (NKEF-A) (Proliferation-associated gene protein) (PAG) (Thioredoxin peroxidase 2) (Thioredoxin-dependent peroxide reductase 2) | 199 | PRDX1 | 10 | 12 | 11 | 55.3 | 68.8 | 51.3 |
| 471 | -0.71 | -0.21 | 0.06 | O94903 | Proline synthase co-transcribed bacterial homolog protein | 275 | PROSC | 3 | 1 | 1 | 17.5 | 4.7 | 4.7 |
| 472 | -0.70 | 0.61 | 0.76 | O60506 | Heterogeneous nuclear ribonucleoprotein Q (hnRNP Q) (Glycine- and tyrosine-rich RNA-binding protein) (GRY-RBP) (NS1-associated protein 1) (Synaptotagmin-binding, cytoplasmic RNA-interacting protein) | 623 | SYNCRIP | 5 | 2 | 6 | 13.1 | 6.5 | 13.9 |
| 473 | -0.70 | 0.00 | -0.01 | E9PHI6 | Cytoplasmic dynein 1 light intermediate chain 1 | 407 | DYNC1LI1 | 1 | 0 | 1 | 4.7 | 0 | 4.7 |
| 474 | -0.70 | 0.00 | 0.87 | Q9NZN4 | EH domain-containing protein 2 (PAST homolog 2) | 543 | EHD2 | 1 | 0 | 2 | 2.8 | 0 | 5 |
| 475 | -0.70 | 1.27 | 1.55 | Q15393 | Splicing factor 3B subunit 3 (Pre-mRNA-splicing factor SF3b 130 kDa subunit) (SF3b130) (STAF130) (Spliceosome-associated protein 130) (SAP 130) | 1217 | SF3B3 | 2 | 3 | 3 | 1.7 | 3.5 | 4.1 |
| 476 | -0.70 | 0.66 | -0.13 | P61006 | Ras-related protein Rab-8A (Oncogene c-mel) | 207 | RAB8A | 3 | 4 | 3 | 14.5 | 21.3 | 14.5 |
| 477 | -0.70 | -0.76 | 0.16 | Q14116 | Interleukin-18 (IL-18) (Iboctadekin) (Interferon gamma-inducing factor) (IFN-gamma-inducing factor) (Interleukin-1 gamma) (IL-1 gamma) | 193 | IL18 | 2 | 2 | 2 | 13.2 | 9.5 | 10.1 |
| 478 | -0.70 | 0.85 | 0.58 | P61026 | Ras-related protein Rab-10 | 200 | RAB10 | 2 | 7 | 3 | 9.5 | 32 | 17 |

| | | | | | | | | | | | | | |
|---|---|---|---|---|---|---|---|---|---|---|---|---|---|
| 479 | -0.70 | 1.11 | 0.91 | P40926 | Malate dehydrogenase, mitochondrial (EC 1.1.1.37) | 338 | MDH2 | 4 | 15 | 8 | 15.4 | 56.5 | 25.7 |
| 480 | -0.69 | 0.76 | 0.36 | Q8TED1 | Probable glutathione peroxidase 8 (GPx-8) (GSHPx-8) (EC 1.11.1.9) | 209 | GPX8 | 1 | 2 | 2 | 5.3 | 10 | 10 |
| 481 | -0.69 | -0.08 | 0.71 | P49773 | Histidine triad nucleotide-binding protein 1 (EC 3.-.-.-) (Adenosine 5'-monophosphoramidase) (Protein kinase C inhibitor 1) (Protein kinase C-interacting protein 1) (PKCI-1) | 126 | HINT1 | 2 | 4 | 2 | 35.7 | 31 | 35.7 |
| 482 | -0.69 | -0.48 | -0.23 | P22102 | Trifunctional purine biosynthetic protein adenosine-3 [Includes: Phosphoribosylamine--glycine ligase (EC 6.3.4.13) (Glycinamide ribonucleotide synthetase) (GARS) (Phosphoribosylglycinamide synthetase); Phosphoribosylformylglycinamidine cyclo-ligase (EC 6.3.3.1) (AIR synthase) (AIRS) (Phosphoribosyl-aminoimidazole synthetase); Phosphoribosylglycinamide formyltransferase (EC 2.1.2.2) (5'-phosphoribosylglycinamide transformylase) (GAR transformylase) (GART)] | 1010 | GART | 3 | 1 | 1 | 4.6 | 1.3 | 1.3 |
| 483 | -0.69 | -0.04 | 0.76 | P62826 | GTP-binding nuclear protein Ran (Androgen receptor-associated protein 24) (GTPase Ran) (Ras-like protein TC4) (Ras-related nuclear protein) | 216 | RAN | 8 | 7 | 6 | 38.4 | 38.9 | 27.3 |
| 484 | -0.69 | 0.00 | 0.85 | Q5T4S7 | E3 ubiquitin-protein ligase UBR4 (EC 6.3.2.-) (600 kDa retinoblastoma protein-associated factor) (N-recognin-4) (Retinoblastoma-associated factor of 600 kDa) | 5183 | UBR4 | 1 | 0 | 2 | 0.2 | 0 | 0.4 |

| | | | | | | | | | | | | | |
|---|---|---|---|---|---|---|---|---|---|---|---|---|---|
| | | | | | (RBAF600) (p600) (Zinc finger UBR1-type protein 1) | | | | | | | | |
| 485 | -0.69 | -0.04 | 0.93 | O14980 | Exportin-1 (Exp1) (Chromosome region maintenance 1 protein homolog) | 1071 | XPO1 | 10 | 7 | 7 | 11.4 | 8.3 | 8.3 |
| 486 | -0.68 | 0.00 | 1.37 | P62195 | 26S protease regulatory subunit 8 (26S proteasome AAA-ATPase subunit RPT6) (Proteasome 26S subunit ATPase 5) (Proteasome subunit p45) (Thyroid hormone receptor-interacting protein 1) (TRIP1) (p45/SUG) | 406 | PSMC5 | 4 | 0 | 6 | 15.8 | 0 | 22.1 |
| 487 | -0.68 | 0.33 | 0.40 | O14818 | Proteasome subunit alpha type-7 (EC 3.4.25.1) (Proteasome subunit RC6-1) (Proteasome subunit XAPC7) | 248 | PSMA7 | 6 | 8 | 5 | 32.7 | 40.3 | 23.4 |
| 488 | -0.68 | 1.03 | 0.40 | P61106 | Ras-related protein Rab-14 | 215 | RAB14 | 6 | 9 | 5 | 39.1 | 58.1 | 23.7 |
| 489 | -0.68 | 0.29 | 1.03 | P20618 | Proteasome subunit beta type-1 (EC 3.4.25.1) (Macropain subunit C5) (Multicatalytic endopeptidase complex subunit C5) (Proteasome component C5) (Proteasome gamma chain) | 241 | PSMB1 | 3 | 7 | 4 | 20.3 | 35.7 | 26.6 |
| 490 | -0.67 | 0.35 | 0.46 | O15145 | Actin-related protein 2/3 complex subunit 3 (Arp2/3 complex 21 kDa subunit) (p21-ARC) | 178 | ARPC3 | 3 | 3 | 1 | 16.3 | 20.8 | 7.3 |
| 491 | -0.67 | 0.00 | 0.30 | E9PI86 | Nuclear autoantigenic sperm protein (Fragment) | 256 | NASP | 2 | 0 | 1 | 13.7 | 0 | 4.7 |

| | | | | | | | | | | | | | |
|---|---|---|---|---|---|---|---|---|---|---|---|---|---|
| 492 | -0.67 | 0.00 | 0.00 | Q9NR28 | Diablo homolog, mitochondrial (Direct IAP-binding protein with low pI) (Second mitochondria-derived activator of caspase) (Smac) | 239 | DIABLO | 2 | 0 | 0 | 12.4 | 0 | 0 |
| 493 | -0.67 | 0.00 | 0.84 | B7ZBQ0 | Serine/threonine-protein phosphatase 2A activator (EC 5.2.1.8) (Phosphotyrosyl phosphatase activator) | 61 | PPP2R4 | 1 | 0 | 1 | 13.1 | 0 | 13.1 |
| 494 | -0.67 | 0.10 | 0.11 | O14737 | Programmed cell death protein 5 (TF-1 cell apoptosis-related protein 19) (Protein TFAR19) | 125 | PDCD5 | 3 | 2 | 3 | 18.4 | 17.6 | 18.4 |
| 495 | -0.66 | 0.12 | 0.58 | P33316 | Deoxyuridine 5'-triphosphate nucleotidohydrolase, mitochondrial (dUTPase) (EC 3.6.1.23) (dUTP pyrophosphatase) | 252 | DUT | 2 | 2 | 2 | 18.9 | 15.9 | 17.7 |
| 496 | -0.66 | 0.18 | 0.60 | P13639 | Elongation factor 2 (EF-2) | 858 | EEF2 | 15 | 20 | 24 | 21.9 | 33.4 | 34.5 |
| 497 | -0.66 | 0.64 | 0.47 | P0CW22 | 40S ribosomal protein S17-like | 135 | RPS17L | 4 | 5 | 1 | 51.9 | 39.3 | 7.4 |
| 498 | -0.65 | 0.06 | 0.75 | P62851 | 40S ribosomal protein S25 | 125 | RPS25 | 3 | 3 | 2 | 24 | 16 | 16 |
| 499 | -0.65 | -0.32 | 0.37 | H0YH58 | Oligoribonuclease, mitochondrial (Fragment) | 111 | REXO2 | 1 | 1 | 1 | 9 | 9 | 9 |
| 500 | -0.65 | 0.62 | 0.00 | A0A0A0MSK5 | Torsin-1A-interacting protein 1 | 462 | TOR1AIP1 | 1 | 1 | 0 | 2.6 | 2.8 | 0 |
| 501 | -0.65 | 0.75 | 1.08 | P04792 | Heat shock protein beta-1 (HspB1) (28 kDa heat shock protein) (Estrogen-regulated 24 kDa protein) (Heat shock 27 kDa protein) (HSP 27) (Stress-responsive protein 27) (SRP27) | 205 | HSPB1 | 4 | 10 | 5 | 21.5 | 69.8 | 24.9 |
| 502 | -0.65 | -0.76 | 0.70 | P00352 | Retinal dehydrogenase 1 (RALDH 1) (RalDH1) (EC 1.2.1.36) (ALDH-E1) (ALHDII) (Aldehyde dehydrogenase family 1 member A1) (Aldehyde dehydrogenase, cytosolic) | 501 | ALDH1A1 | 24 | 21 | 22 | 58.7 | 47.5 | 53.5 |

| | | | | | | | | | | | | | |
|---|---|---|---|---|---|---|---|---|---|---|---|---|---|
| 503 | -0.64 | 0.00 | 1.10 | Q6NUK1 | Calcium-binding mitochondrial carrier protein SCaMC-1 (Mitochondrial ATP-Mg/Pi carrier protein 1) (Mitochondrial Ca(2+)-dependent solute carrier protein 1) (Small calcium-binding mitochondrial carrier protein 1) (Solute carrier family 25 member 24) | 477 | SLC25A24 | 1 | 0 | 1 | 3.5 | 0 | 3.5 |
| 504 | -0.64 | -0.25 | 0.60 | Q13200 | 26S proteasome non-ATPase regulatory subunit 2 (26S proteasome regulatory subunit RPN1) (26S proteasome regulatory subunit S2) (26S proteasome subunit p97) (Protein 55.11) (Tumor necrosis factor type 1 receptor-associated protein 2) | 908 | PSMD2 | 5 | 1 | 4 | 8.5 | 1.2 | 7.1 |
| 505 | -0.64 | 1.02 | 1.48 | G3V576 | Heterogeneous nuclear ribonucleoproteins C1/C2 | 231 | HNRNPC | 4 | 5 | 5 | 15.2 | 22.9 | 19.5 |
| 506 | -0.64 | 1.04 | 1.05 | Q7KZF4 | Staphylococcal nuclease domain-containing protein 1 (100 kDa coactivator) (EBNA2 coactivator p100) (Tudor domain-containing protein 11) (p100 co-activator) | 910 | SND1 | 6 | 3 | 7 | 9.7 | 5.5 | 12.7 |
| 507 | -0.64 | 0.00 | 0.00 | Q12974 | Protein tyrosine phosphatase type IVA 2 (EC 3.1.3.48) (HU-PP-1) (OV-1) (PTP(CAAXII)) (Protein-tyrosine phosphatase 4a2) (Protein-tyrosine phosphatase of regenerating liver 2) (PRL-2) | 167 | PTP4A2 | 1 | 0 | 0 | 11 | 0 | 0 |
| 508 | -0.64 | 1.26 | 0.63 | Q9NZM1 | Myoferlin (Fer-1-like protein 3) | 2061 | MYOF | 11 | 22 | 14 | 5.7 | 13.3 | 8.9 |
| 509 | -0.63 | 0.51 | 0.03 | Q13509 | Tubulin beta-3 chain (Tubulin beta-4 chain) (Tubulin beta-III) | 450 | TUBB3 | 4 | 5 | 7 | 11.8 | 14.4 | 17.1 |
| 510 | -0.63 | -0.23 | 0.21 | P31947 | 14-3-3 protein sigma (Epithelial cell marker protein 1) (Stratifin) | 248 | SFN | 6 | 8 | 8 | 27 | 36.7 | 36.7 |

| | | | | | | | | | | | | | |
|---|---|---|---|---|---|---|---|---|---|---|---|---|---|
| 511 | -0.63 | 0.55 | 0.22 | C9J9K3 | 40S ribosomal protein SA (Fragment) | 263 | RPSA | 9 | 7 | 5 | 36.5 | 38.8 | 23.2 |
| 512 | -0.63 | 0.12 | 0.37 | P68371 | Tubulin beta-4B chain (Tubulin beta-2 chain) (Tubulin beta-2C chain) | 445 | TUBB4B | 8 | 11 | 9 | 26.7 | 36.2 | 26.5 |
| 513 | -0.62 | 0.75 | 0.83 | P01111 | GTPase NRas (Transforming protein N-Ras) | 189 | NRAS | 3 | 1 | 2 | 20.1 | 6.3 | 12.2 |
| 514 | -0.62 | 0.90 | 0.71 | J3QR09 | Ribosomal protein L19 | 193 | RPL19 | 2 | 1 | 2 | 8.8 | 8.8 | 8.8 |
| 515 | -0.62 | 0.00 | 0.17 | O00429 | Dynamin-1-like protein (EC 3.6.5.5) (Dnm1p/Vps1p-like protein) (DVLP) (Dynamin family member proline-rich carboxyl-terminal domain less) (Dymple) (Dynamin-like protein) (Dynamin-like protein 4) (Dynamin-like protein IV) (HdynIV) (Dynamin-related protein 1) | 736 | DNM1L | 2 | 0 | 1 | 2.9 | 0 | 1.6 |
| 516 | -0.62 | 0.72 | 0.31 | P07355 | Annexin A2 (Annexin II) (Annexin-2) (Calpactin I heavy chain) (Calpactin-1 heavy chain) (Chromobindin-8) (Lipocortin II) (Placental anticoagulant protein IV) (PAP-IV) (Protein I) (p36) | 339 | ANXA2 | 17 | 17 | 11 | 51 | 52.8 | 29.5 |
| 517 | -0.62 | 0.26 | 0.34 | P04406 | Glyceraldehyde-3-phosphate dehydrogenase (GAPDH) (EC 1.2.1.12) (Peptidyl-cysteine S-nitrosylase GAPDH) (EC 2.6.99.-) | 335 | GAPDH | 8 | 9 | 8 | 31.9 | 46 | 28.7 |
| 518 | -0.62 | 1.25 | 1.21 | H7C2U6 | Protein NipSnap homolog 1 (Fragment) | 221 | NIPSNAP1 | 2 | 1 | 2 | 10.9 | 6.3 | 10.9 |
| 519 | -0.62 | 0.35 | 0.40 | F8VZJ2 | Nascent polypeptide-associated complex subunit alpha | 136 | NACA | 3 | 3 | 2 | 30.1 | 30.9 | 20.6 |
| 520 | -0.61 | 0.18 | 0.69 | P08238 | Heat shock protein HSP 90-beta (HSP 90) (Heat shock 84 kDa) (HSP 84) (HSP84) | 724 | HSP90AB1 | 28 | 14 | 29 | 35.8 | 21.3 | 35.6 |
| 521 | -0.61 | 2.78 | 0.63 | A0A087WTV6 | Pyrroline-5-carboxylate reductase 2 | 246 | PYCR2 | 2 | 1 | 1 | 13.4 | 7.3 | 6.1 |

| | | | | | | | | | | | | | |
|---|---|---|---|---|---|---|---|---|---|---|---|---|---|
| 522 | -0.61 | 0.37 | 0.64 | P31948 | Stress-induced-phosphoprotein 1 (STI1) (Hsc70/Hsp90-organizing protein) (Hop) (Renal carcinoma antigen NY-REN-11) (Transformation-sensitive protein IEF SSP 3521) | 543 | STIP1 | 11 | 5 | 15 | 20.3 | 11.2 | 30.9 |
| 523 | -0.61 | 0.58 | -0.03 | P26639 | Threonine--tRNA ligase, cytoplasmic (EC 6.1.1.3) (Threonyl-tRNA synthetase) (ThrRS) | 723 | TARS | 3 | 1 | 2 | 4 | 1.4 | 2.6 |
| 524 | -0.61 | -1.22 | 0.29 | Q9Y617 | Phosphoserine aminotransferase (EC 2.6.1.52) (Phosphohydroxythreonine aminotransferase) (PSAT) | 370 | PSAT1 | 7 | 1 | 6 | 19.7 | 2.7 | 16.8 |
| 525 | -0.61 | 0.00 | 0.59 | P68366 | Tubulin alpha-4A chain (Alpha-tubulin 1) (Testis-specific alpha-tubulin) (Tubulin H2-alpha) (Tubulin alpha-1 chain) | 448 | TUBA4A | 8 | 8 | 11 | 25.4 | 26.6 | 37.9 |
| 526 | -0.61 | 0.47 | 0.87 | P53396 | ATP-citrate synthase (EC 2.3.3.8) (ATP-citrate (pro-S-)-lyase) (ACL) (Citrate cleavage enzyme) | 1101 | ACLY | 9 | 12 | 10 | 11.2 | 14.8 | 14.2 |
| 527 | -0.60 | -0.50 | -0.02 | P60981 | Destrin (Actin-depolymerizing factor) (ADF) | 165 | DSTN | 6 | 6 | 3 | 34.5 | 41.9 | 14.9 |
| 528 | -0.60 | 0.03 | 0.71 | O60218 | Aldo-keto reductase family 1 member B10 (EC 1.1.1.-) (ARL-1) (Aldose reductase-like) (Aldose reductase-related protein) (ARP) (hARP) (Small intestine reductase) (SI reductase) | 316 | AKR1B10 | 15 | 16 | 13 | 50.9 | 64.6 | 49.1 |
| 529 | -0.59 | 0.29 | 0.84 | A0A024QZP7 | Cell division cycle 2, G1 to S and G2 to M, isoform CRA_a (Cyclin-dependent kinase 1) | 297 | CDC2 | 1 | 2 | 3 | 4 | 7.7 | 11.8 |
| 530 | -0.59 | 0.02 | -0.14 | O75223 | Gamma-glutamylcyclotransferase (EC 2.3.2.4) (Cytochrome c-releasing factor 21) | 188 | GGCT | 2 | 3 | 1 | 14.4 | 17 | 5.3 |
| 531 | -0.59 | 2.57 | 1.07 | M0QXS5 | Heterogeneous nuclear ribonucleoprotein L (Fragment) | 530 | HNRNPL | 2 | 2 | 2 | 4.9 | 4 | 9.6 |

| | | | | | | | | | | | | | |
|---|---|---|---|---|---|---|---|---|---|---|---|---|---|
| 532 | -0.59 | -0.53 | 0.12 | P43490 | Nicotinamide phosphoribosyltransferase (NAmPRTase) (Nampt) (EC 2.4.2.12) (Pre-B-cell colony-enhancing factor 1) (Pre-B cell-enhancing factor) (Visfatin) | 491 | NAMPT | 10 | 6 | 10 | 28.9 | 16.9 | 25.1 |
| 533 | -0.59 | 0.60 | 0.72 | Q9Y266 | Nuclear migration protein nudC (Nuclear distribution protein C homolog) | 331 | NUDC | 4 | 2 | 2 | 13.9 | 6.9 | 7.9 |
| 534 | -0.58 | 0.88 | 1.99 | Q08211 | ATP-dependent RNA helicase A (RHA) (EC 3.6.4.13) (DEAH box protein 9) (Leukophysin) (LKP) (Nuclear DNA helicase II) (NDH II) | 1270 | DHX9 | 5 | 3 | 7 | 4.4 | 2.8 | 7.2 |
| 535 | -0.58 | 1.13 | 1.17 | P05556 | Integrin beta-1 (Fibronectin receptor subunit beta) (Glycoprotein IIa) (GPIIA) (VLA-4 subunit beta) (CD antigen CD29) | 798 | ITGB1 | 6 | 9 | 6 | 11 | 14.2 | 9.5 |
| 536 | -0.58 | 1.25 | 0.88 | P30048 | Thioredoxin-dependent peroxide reductase, mitochondrial (EC 1.11.1.15) (Antioxidant protein 1) (AOP-1) (HBC189) (Peroxiredoxin III) (Prx-III) (Peroxiredoxin-3) (Protein MER5 homolog) | 256 | PRDX3 | 7 | 7 | 6 | 34 | 43.3 | 21 |
| 537 | -0.58 | 0.00 | 0.49 | P14735 | Insulin-degrading enzyme (EC 3.4.24.56) (Abeta-degrading protease) (Insulin protease) (Insulinase) (Insulysin) | 1019 | IDE | 3 | 0 | 3 | 3.5 | 0 | 3.5 |
| 538 | -0.58 | 1.02 | 0.60 | P48047 | ATP synthase subunit O, mitochondrial (Oligomycin sensitivity conferral protein) (OSCP) | 213 | ATP5O | 5 | 7 | 5 | 25.8 | 39 | 33.3 |
| 539 | -0.58 | 0.03 | 0.36 | B8ZZQ6 | Prothymosin alpha | 107 | PTMA | 1 | 1 | 1 | 13.1 | 13.1 | 13.1 |
| 540 | -0.58 | 1.77 | 0.28 | P09601 | Heme oxygenase 1 (HO-1) (EC 1.14.99.3) | 288 | HMOX1 | 4 | 6 | 2 | 15.3 | 26 | 4.5 |

| | | | | | | | | | | | | | |
|---|---|---|---|---|---|---|---|---|---|---|---|---|---|
| 541 | -0.58 | 0.01 | 1.10 | P62906 | 60S ribosomal protein L10a (CSA-19) (Neural precursor cell expressed developmentally down-regulated protein 6) (NEDD-6) | 217 | RPL10A | 9 | 5 | 8 | 36.9 | 23.5 | 36.9 |
| 542 | -0.57 | 0.00 | 1.00 | P04181 | Ornithine aminotransferase, mitochondrial (EC 2.6.1.13) (Ornithine delta-aminotransferase) (Ornithine--oxo-acid aminotransferase) [Cleaved into: Ornithine aminotransferase, hepatic form; Ornithine aminotransferase, renal form] | 439 | OAT | 3 | 0 | 3 | 11.6 | 0 | 11.6 |
| 543 | -0.57 | 0.21 | 1.18 | P52272 | Heterogeneous nuclear ribonucleoprotein M (hnRNP M) | 730 | HNRNPM | 3 | 1 | 5 | 6.2 | 2.3 | 11 |
| 544 | -0.57 | 1.02 | 0.53 | Q13162 | Peroxiredoxin-4 (EC 1.11.1.15) (Antioxidant enzyme AOE372) (AOE37-2) (Peroxiredoxin IV) (Prx-IV) (Thioredoxin peroxidase AO372) (Thioredoxin-dependent peroxide reductase A0372) | 271 | PRDX4 | 5 | 7 | 4 | 23.6 | 35.4 | 15.9 |
| 545 | -0.57 | 0.81 | 0.00 | P02792 | Ferritin light chain (Ferritin L subunit) | 175 | FTL | 1 | 1 | 0 | 8.6 | 8.6 | 0 |
| 546 | -0.57 | 0.76 | 0.69 | P30101 | Protein disulfide-isomerase A3 (EC 5.3.4.1) (58 kDa glucose-regulated protein) (58 kDa microsomal protein) (p58) (Disulfide isomerase ER-60) (Endoplasmic reticulum resident protein 57) (ER protein 57) (ERp57) (Endoplasmic reticulum resident protein 60) (ER protein 60) (ERp60) | 505 | PDIA3 | 10 | 9 | 10 | 22.8 | 18.4 | 26.7 |
| 547 | -0.57 | 0.54 | 0.20 | K7ERF1 | Eukaryotic translation initiation factor 3 subunit K (eIF3k) (Eukaryotic translation initiation factor 3 subunit 12) (eIF-3 p25) | 192 | EIF3K | 2 | 2 | 1 | 14.6 | 13 | 7.3 |

| | | | | | | | | | | | | | |
|---|---|---|---|---|---|---|---|---|---|---|---|---|---|
| 548 | -0.57 | 0.25 | 0.86 | Q02218 | 2-oxoglutarate dehydrogenase, mitochondrial (EC 1.2.4.2) (2-oxoglutarate dehydrogenase complex component E1) (OGDC-E1) (Alpha-ketoglutarate dehydrogenase) | 1023 | OGDH | 2 | 2 | 2 | 2.1 | 2.5 | 2.1 |
| 549 | -0.56 | 0.49 | 0.65 | Q5VTE0 | Putative elongation factor 1-alpha-like 3 (EF-1-alpha-like 3) (Eukaryotic elongation factor 1 A-like 3) (eEF1A-like 3) (Eukaryotic translation elongation factor 1 alpha-1 pseudogene 5) | 462 | EEF1A1P5 | 11 | 9 | 13 | 30.5 | 29.7 | 33.3 |
| 550 | -0.56 | -0.10 | 0.56 | P27348 | 14-3-3 protein theta (14-3-3 protein T-cell) (14-3-3 protein tau) (Protein HS1) | 245 | YWHAQ | 6 | 9 | 7 | 31.8 | 36.7 | 29 |
| 551 | -0.56 | 1.28 | 0.97 | P35998 | 26S protease regulatory subunit 7 (26S proteasome AAA-ATPase subunit RPT1) (Proteasome 26S subunit ATPase 2) (Protein MSS1) | 433 | PSMC2 | 4 | 2 | 4 | 11.3 | 6.9 | 12.5 |
| 552 | -0.56 | 0.00 | 0.71 | O43837 | Isocitrate dehydrogenase [NAD] subunit beta, mitochondrial (EC 1.1.1.41) (Isocitric dehydrogenase subunit beta) (NAD(+)-specific ICDH subunit beta) | 385 | IDH3B | 3 | 0 | 3 | 9.9 | 0 | 9.9 |
| 553 | -0.56 | -0.04 | 0.23 | P31689 | DnaJ homolog subfamily A member 1 (DnaJ protein homolog 2) (HSDJ) (Heat shock 40 kDa protein 4) (Heat shock protein J2) (HSJ-2) (Human DnaJ protein 2) (hDj-2) | 397 | DNAJA1 | 3 | 1 | 3 | 10 | 3.9 | 10 |
| 554 | -0.56 | -0.29 | 0.97 | Q9UBQ7 | Glyoxylate reductase/hydroxypyruvate reductase (EC 1.1.1.79) (EC 1.1.1.81) | 328 | GRHPR | 3 | 1 | 5 | 18 | 3 | 24.1 |
| 555 | -0.56 | 1.59 | 0.86 | P61604 | 10 kDa heat shock protein, mitochondrial (Hsp10) (10 kDa chaperonin) (Chaperonin 10) (CPN10) (Early-pregnancy factor) (EPF) | 102 | HSPE1 | 1 | 7 | 4 | 13.7 | 57.8 | 35.3 |

| | | | | | | | | | | | | | |
|---|---|---|---|---|---|---|---|---|---|---|---|---|---|
| 556 | -0.55 | 1.44 | 1.14 | E9PKG1 | Protein arginine N-methyltransferase 1 | 325 | PRMT1 | 1 | 2 | 2 | 3.7 | 8.3 | 8.3 |
| 557 | -0.55 | 0.15 | 0.52 | P26038 | Moesin (Membrane-organizing extension spike protein) | 577 | MSN | 10 | 5 | 9 | 15.3 | 8.7 | 13.5 |
| 558 | -0.55 | 0.00 | 1.09 | Q6PI48 | Aspartate--tRNA ligase, mitochondrial (EC 6.1.1.12) (Aspartyl-tRNA synthetase) (AspRS) | 645 | DARS2 | 1 | 0 | 1 | 1.9 | 0 | 1.1 |
| 559 | -0.55 | 0.43 | 0.52 | P23284 | Peptidyl-prolyl cis-trans isomerase B (PPIase B) (EC 5.2.1.8) (CYP-S1) (Cyclophilin B) (Rotamase B) (S-cyclophilin) (SCYLP) | 216 | PPIB | 6 | 10 | 6 | 21.3 | 48.1 | 22.2 |
| 560 | -0.55 | 0.91 | 0.93 | O14880 | Microsomal glutathione S-transferase 3 (Microsomal GST-3) (EC 2.5.1.18) (Microsomal GST-III) | 152 | MGST3 | 1 | 1 | 1 | 9.2 | 9.2 | 9.9 |
| 561 | -0.54 | 1.63 | 0.88 | E9PPQ5 | Cysteine and histidine-rich domain-containing protein 1 | 184 | CHORDC1 | 2 | 1 | 3 | 12.5 | 7.6 | 25 |
| 562 | -0.54 | 0.00 | 2.38 | J3KQV6 | COP9 signalosome complex subunit 7b | 54 | COPS7B | 1 | 0 | 1 | 35.2 | 0 | 35.2 |
| 563 | -0.54 | -0.25 | -0.11 | P13693 | Translationally-controlled tumor protein (TCTP) (Fortilin) (Histamine-releasing factor) (HRF) (p23) | 172 | TPT1 | 4 | 4 | 2 | 30.2 | 30.2 | 15.7 |
| 564 | -0.54 | 0.00 | 0.97 | O00116 | Alkyldihydroxyacetonephosphate synthase, peroxisomal (Alkyl-DHAP synthase) (EC 2.5.1.26) (Aging-associated gene 5 protein) (Alkylglycerone-phosphate synthase) | 658 | AGPS | 2 | 0 | 3 | 4 | 0 | 6.5 |
| 565 | -0.54 | 0.81 | 0.49 | Q05193 | Dynamin-1 (EC 3.6.5.5) | 864 | DNM1 | 2 | 1 | 2 | 2.1 | 0.8 | 2.5 |

| | | | | | | | | | | | | | |
|---|---|---|---|---|---|---|---|---|---|---|---|---|---|
| 566 | -0.54 | 0.32 | 0.54 | Q06323 | Proteasome activator complex subunit 1 (11S regulator complex subunit alpha) (REG-alpha) (Activator of multicatalytic protease subunit 1) (Interferon gamma up-regulated I-5111 protein) (IGUP I-5111) (Proteasome activator 28 subunit alpha) (PA28a) (PA28alpha) | 249 | PSME1 | 7 | 6 | 9 | 32.5 | 24.5 | 36.5 |
| 567 | -0.54 | 0.00 | 1.10 | Q9H814 | Phosphorylated adapter RNA export protein (RNA U small nuclear RNA export adapter protein) | 394 | PHAX | 1 | 0 | 1 | 2.8 | 0 | 2.8 |
| 568 | -0.53 | -0.16 | 0.44 | Q9Y237 | Peptidyl-prolyl cis-trans isomerase NIMA-interacting 4 (EC 5.2.1.8) (Parvulin-14) (Par14) (hPar14) (Parvulin-17) (Par17) (hPar17) (Peptidyl-prolyl cis-trans isomerase Pin4) (PPIase Pin4) (Peptidyl-prolyl cis/trans isomerase EPVH) (hEPVH) (Rotamase Pin4) | 131 | PIN4 | 1 | 1 | 2 | 9.2 | 9.2 | 17.6 |
| 569 | -0.53 | 0.18 | 0.64 | C9J2Q4 | Septin-2 (Fragment) | 184 | sept2 | 2 | 2 | 2 | 10.9 | 10.9 | 10.9 |
| 570 | -0.53 | 1.87 | 1.19 | Q07021 | Complement component 1 Q subcomponent-binding protein, mitochondrial (ASF/SF2-associated protein p32) (Glycoprotein gC1qBP) (C1qBP) (Hyaluronan-binding protein 1) (Mitochondrial matrix protein p32) (gC1q-R protein) (p33) | 282 | C1QBP | 1 | 2 | 3 | 3.9 | 11 | 24.1 |
| 571 | -0.53 | 0.22 | 1.19 | P55884 | Eukaryotic translation initiation factor 3 subunit B (eIF3b) (Eukaryotic translation initiation factor 3 subunit 9) (Prt1 homolog) (hPrt1) (eIF-3-eta) (eIF3 p110) (eIF3 p116) | 814 | EIF3B | 3 | 3 | 3 | 4.4 | 4.2 | 4.4 |

| | | | | | | | | | | | | | |
|---|---|---|---|---|---|---|---|---|---|---|---|---|---|
| 572 | -0.53 | 1.54 | 1.06 | P52597 | Heterogeneous nuclear ribonucleoprotein F (hnRNP F) (Nucleolin-like protein mcs94-1) [Cleaved into: Heterogeneous nuclear ribonucleoprotein F, N-terminally processed] | 415 | HNRNPF | 2 | 1 | 2 | 6.5 | 4.1 | 7.2 |
| 573 | -0.53 | 0.00 | 0.00 | A0A087X 1D8 | Farnesyl pyrophosphate synthase (Fragment) | 174 | FDPS | 2 | 0 | 0 | 17.8 | 0 | 0 |
| 574 | -0.53 | 0.02 | 0.05 | O00410 | Importin-5 (Imp5) (Importin subunit beta-3) (Karyopherin beta-3) (Ran-binding protein 5) (RanBP5) | 1097 | IPO5 | 6 | 5 | 4 | 8.4 | 6.8 | 5.8 |
| 575 | -0.53 | 1.13 | 0.45 | Q92973 | Transportin-1 (Importin beta-2) (Karyopherin beta-2) (M9 region interaction protein) (MIP) | 898 | TNPO1 | 6 | 3 | 6 | 8 | 3.6 | 6.9 |
| 576 | -0.52 | 0.95 | 0.42 | P38646 | Stress-70 protein, mitochondrial (75 kDa glucose-regulated protein) (GRP-75) (Heat shock 70 kDa protein 9) (Mortalin) (MOT) (Peptide-binding protein 74) (PBP74) | 679 | HSPA9 | 11 | 6 | 14 | 22.7 | 12.4 | 27.4 |
| 577 | -0.52 | 0.22 | 0.71 | D6RAN4 | 60S ribosomal protein L9 (Fragment) | 181 | RPL9 | 4 | 3 | 6 | 19.9 | 23.8 | 45.3 |
| 578 | -0.52 | 0.00 | 0.74 | P25205 | DNA replication licensing factor MCM3 (EC 3.6.4.12) (DNA polymerase alpha holoenzyme-associated protein P1) (P1-MCM3) (RLF subunit beta) (p102) | 808 | MCM3 | 3 | 0 | 3 | 4.5 | 0 | 4.2 |
| 579 | -0.52 | 0.00 | 0.47 | Q9Y3E7 | Charged multivesicular body protein 3 (Chromatin-modifying protein 3) (Neuroendocrine differentiation factor) (Vacuolar protein sorting-associated protein 24) (hVps24) | 222 | CHMP3 | 1 | 0 | 1 | 7.7 | 0 | 7.7 |

| | | | | | | | | | | | | | |
|---|---|---|---|---|---|---|---|---|---|---|---|---|---|
| 580 | -0.52 | 0.36 | 0.58 | O00299 | Chloride intracellular channel protein 1 (Chloride channel ABP) (Nuclear chloride ion channel 27) (NCC27) (Regulatory nuclear chloride ion channel protein) (hRNCC) | 241 | CLIC1 | 3 | 6 | 5 | 13.7 | 28.6 | 26.1 |
| 581 | -0.52 | 1.15 | 1.73 | P27824 | Calnexin (IP90) (Major histocompatibility complex class I antigen-binding protein p88) (p90) | 592 | CANX | 4 | 5 | 4 | 9.6 | 11.8 | 8.3 |
| 582 | -0.52 | 1.68 | 1.81 | P26599 | Polypyrimidine tract-binding protein 1 (PTB) (57 kDa RNA-binding protein PPTB-1) (Heterogeneous nuclear ribonucleoprotein I) (hnRNP I) | 531 | PTBP1 | 3 | 2 | 4 | 6.2 | 6.6 | 11.9 |
| 583 | -0.52 | 0.00 | 0.69 | H7C235 | Sphingomyelin phosphodiesterase 4 (Fragment) | 211 | SMPD4 | 1 | 0 | 1 | 4.3 | 0 | 4.3 |
| 584 | -0.52 | 0.00 | 0.02 | C9J363 | Programmed cell death 10, isoform CRA_b (Programmed cell death protein 10) | 149 | PDCD10 | 1 | 0 | 1 | 8.7 | 0 | 8.7 |
| 585 | -0.51 | 0.00 | 0.45 | M0QX85 | Nitric oxide synthase-interacting protein (Fragment) | 96 | NOSIP | 1 | 0 | 1 | 11.5 | 0 | 11.5 |
| 586 | -0.51 | 0.33 | 0.94 | Q9Y6C9 | Mitochondrial carrier homolog 2 (Met-induced mitochondrial protein) | 303 | MTCH2 | 5 | 4 | 4 | 27.1 | 21.1 | 18.5 |
| 587 | -0.51 | 1.05 | 0.42 | Q9Y6N5 | Sulfide:quinone oxidoreductase, mitochondrial (SQOR) (EC 1.8.5.-) | 450 | SQRDL | 4 | 3 | 3 | 9.1 | 7.3 | 7.1 |
| 588 | -0.51 | 0.69 | 0.37 | Q01082 | Spectrin beta chain, non-erythrocytic 1 (Beta-II spectrin) (Fodrin beta chain) (Spectrin, non-erythroid beta chain 1) | 2364 | SPTBN1 | 11 | 17 | 10 | 6.6 | 9.8 | 6.2 |
| 589 | -0.51 | -0.26 | 1.14 | C9JXB8 | 60S ribosomal protein L24 | 121 | RPL24 | 3 | 4 | 2 | 25.6 | 32.2 | 17.4 |
| 590 | -0.50 | 0.53 | 0.64 | P17987 | T-complex protein 1 subunit alpha (TCP-1-alpha) (CCT-alpha) | 556 | TCP1 | 5 | 3 | 6 | 11.5 | 5.8 | 13.1 |

| | | | | | | | | | | | | | |
|---|---|---|---|---|---|---|---|---|---|---|---|---|---|
| 591 | -0.50 | 0.98 | 0.99 | P07602 | Prosaposin (Proactivator polypeptide) [Cleaved into: Saposin-A (Protein A); Saposin-B-Val; Saposin-B (Cerebroside sulfate activator) (CSAct) (Dispersin) (Sphingolipid activator protein 1) (SAP-1) (Sulfatide/GM1 activator); Saposin-C (A1 activator) (Co-beta-glucosidase) (Glucosylceramidase activator) (Sphingolipid activator protein 2) (SAP-2); Saposin-D (Component C) (Protein C)] | 524 | PSAP | 1 | 6 | 2 | 2.5 | 12.6 | 5.3 |
| 592 | -0.50 | 0.27 | 0.31 | Q92688 | Acidic leucine-rich nuclear phosphoprotein 32 family member B (Acidic protein rich in leucines) (Putative HLA-DR-associated protein I-2) (PHAPI2) (Silver-stainable protein SSP29) | 251 | ANP32B | 4 | 5 | 3 | 22.6 | 22.6 | 16.4 |
| 593 | -0.50 | 0.09 | -0.05 | P16435 | NADPH--cytochrome P450 reductase (CPR) (P450R) (EC 1.6.2.4) | 677 | POR | 4 | 1 | 2 | 8.4 | 1.3 | 3.4 |
| 594 | -0.50 | 0.15 | 0.76 | Q86VP6 | Cullin-associated NEDD8-dissociated protein 1 (Cullin-associated and neddylation-dissociated protein 1) (TBP-interacting protein of 120 kDa A) (TBP-interacting protein 120A) (p120 CAND1) | 1230 | CAND1 | 9 | 10 | 13 | 9.6 | 11.1 | 14.2 |
| 595 | -0.50 | 0.00 | 0.94 | G5EA06 | 28S ribosomal protein S27, mitochondrial (Mitochondrial ribosomal protein S27, isoform CRA_b) | 358 | MRPS27 | 2 | 0 | 1 | 7.5 | 0 | 3.4 |
| 596 | -0.50 | 0.00 | 0.08 | J3KTI3 | NAD kinase (Fragment) | 74 | NADK | 1 | 0 | 1 | 13.5 | 0 | 13.5 |
| 597 | -0.50 | 0.21 | 1.49 | O00151 | PDZ and LIM domain protein 1 (C-terminal LIM domain protein 1) (Elfin) (LIM domain protein CLP-36) | 329 | PDLIM1 | 2 | 1 | 3 | 8.8 | 3 | 17 |

| | | | | | | | | | | | | | |
|---|---|---|---|---|---|---|---|---|---|---|---|---|---|
| 598 | -0.50 | 0.70 | 0.47 | P62877 | E3 ubiquitin-protein ligase RBX1 (EC 6.3.2.-) (Protein ZYP) (RING finger protein 75) (RING-box protein 1) (Rbx1) (Regulator of cullins 1) [Cleaved into: E3 ubiquitin-protein ligase RBX1, N-terminally processed] | 108 | RBX1 | 1 | 1 | 1 | 17.6 | 16.7 | 16.7 |
| 599 | -0.49 | -0.27 | 0.62 | I3L3P7 | 40S ribosomal protein S15a | 100 | RPS15A | 4 | 2 | 3 | 26 | 15 | 19 |
| 600 | -0.49 | -0.26 | 0.45 | P00338 | L-lactate dehydrogenase A chain (LDH-A) (EC 1.1.1.27) (Cell proliferation-inducing gene 19 protein) (LDH muscle subunit) (LDH-M) (Renal carcinoma antigen NY-REN-59) | 332 | LDHA | 8 | 15 | 8 | 21.1 | 46.4 | 20.2 |
| 601 | -0.49 | 0.00 | 0.50 | P30419 | Glycylpeptide N-tetradecanoyltransferase 1 (EC 2.3.1.97) (Myristoyl-CoA:protein N-myristoyltransferase 1) (NMT 1) (Type I N-myristoyltransferase) (Peptide N-myristoyltransferase 1) | 496 | NMT1 | 2 | 0 | 2 | 6.5 | 0 | 6.5 |
| 602 | -0.49 | 0.88 | 0.64 | Q96AG4 | Leucine-rich repeat-containing protein 59 (Ribosome-binding protein p34) (p34) | 307 | LRRC59 | 4 | 5 | 6 | 17.6 | 21.5 | 25.1 |
| 603 | -0.49 | 0.01 | 0.43 | M0QZ52 | Calmodulin (Calmodulin 3 (Phosphorylase kinase, delta), isoform CRA_d) | 83 | CALM3 | 4 | 2 | 3 | 62.7 | 20.5 | 62.7 |
| 604 | -0.49 | 0.00 | -0.63 | P51452 | Dual specificity protein phosphatase 3 (EC 3.1.3.16) (EC 3.1.3.48) (Dual specificity protein phosphatase VHR) (Vaccinia H1-related phosphatase) (VHR) | 185 | DUSP3 | 2 | 1 | 1 | 14.1 | 7.6 | 6.5 |
| 605 | -0.48 | 0.00 | 0.00 | Q96F63 | Coiled-coil domain-containing protein 97 | 343 | CCDC97 | 1 | 0 | 0 | 3.2 | 0 | 0 |
| 606 | -0.48 | 0.35 | 0.12 | P07737 | Profilin-1 (Epididymis tissue protein Li 184a) (Profilin I) | 140 | PFN1 | 8 | 7 | 6 | 48.6 | 61.4 | 37.1 |

| | | | | | | | | | | | | | |
|---|---|---|---|---|---|---|---|---|---|---|---|---|---|
| 607 | -0.48 | -0.07 | 0.64 | P49207 | 60S ribosomal protein L34 | 117 | RPL34 | 1 | 2 | 1 | 6.8 | 14.5 | 6.8 |
| 608 | -0.48 | -0.71 | 0.00 | Q00535 | Cyclin-dependent-like kinase 5 (EC 2.7.11.1) (Cell division protein kinase 5) (Serine/threonine-protein kinase PSSALRE) (Tau protein kinase II catalytic subunit) (TPKII catalytic subunit) | 292 | CDK5 | 1 | 1 | 0 | 3.1 | 3.1 | 0 |
| 609 | -0.47 | 0.44 | 0.98 | P49721 | Proteasome subunit beta type-2 (EC 3.4.25.1) (Macropain subunit C7-I) (Multicatalytic endopeptidase complex subunit C7-I) (Proteasome component C7-I) | 201 | PSMB2 | 2 | 6 | 5 | 14.4 | 28.4 | 28.9 |
| 610 | -0.47 | -0.17 | 0.28 | P43487 | Ran-specific GTPase-activating protein (Ran-binding protein 1) (RanBP1) | 201 | RANBP1 | 1 | 2 | 1 | 5.5 | 10 | 5.5 |
| 611 | -0.47 | 0.36 | 0.49 | P40227 | T-complex protein 1 subunit zeta (TCP-1-zeta) (Acute morphine dependence-related protein 2) (CCT-zeta-1) (HTR3) (Tcp20) | 531 | CCT6A | 9 | 3 | 8 | 24.5 | 10 | 24.1 |
| 612 | -0.47 | 0.21 | 0.75 | P49914 | 5-formyltetrahydrofolate cyclo-ligase (EC 6.3.3.2) (5,10-methenyl-tetrahydrofolate synthetase) (MTHFS) (Methenyl-THF synthetase) | 203 | MTHFS | 1 | 2 | 2 | 4.4 | 10.3 | 10.3 |
| 613 | -0.47 | 0.72 | 1.08 | Q15366 | Poly(rC)-binding protein 2 (Alpha-CP2) (Heterogeneous nuclear ribonucleoprotein E2) (hnRNP E2) | 365 | PCBP2 | 1 | 5 | 3 | 3.9 | 18.8 | 13.6 |
| 614 | -0.47 | 0.73 | 0.61 | Q14152 | Eukaryotic translation initiation factor 3 subunit A (eIF3a) (Eukaryotic translation initiation factor 3 subunit 10) (eIF-3-theta) (eIF3 p167) (eIF3 p180) (eIF3 p185) | 1382 | EIF3A | 8 | 8 | 8 | 6 | 6.7 | 6.4 |
| 615 | -0.46 | 0.00 | 0.75 | E7ESP4 | Integrin alpha-2 | 942 | ITGA2 | 1 | 0 | 1 | 1.4 | 0 | 1.1 |

| | | | | | | | | | | | | | |
|---|---|---|---|---|---|---|---|---|---|---|---|---|---|
| 616 | -0.46 | 0.05 | 0.36 | P22234 | Multifunctional protein ADE2 [Includes: Phosphoribosylaminoimidazole-succinocarboxamide synthase (EC 6.3.2.6) (SAICAR synthetase); Phosphoribosylaminoimidazole carboxylase (EC 4.1.1.21) (AIR carboxylase) (AIRC)] | 425 | PAICS | 6 | 6 | 5 | 15.5 | 15.5 | 12.5 |
| 617 | -0.46 | 0.11 | 0.36 | P12004 | Proliferating cell nuclear antigen (PCNA) (Cyclin) | 261 | PCNA | 9 | 5 | 8 | 51.3 | 25.7 | 36 |
| 618 | -0.46 | -0.26 | 0.73 | P22392 | Nucleoside diphosphate kinase B (NDK B) (NDP kinase B) (EC 2.7.4.6) (C-myc purine-binding transcription factor PUF) (Histidine protein kinase NDKB) (EC 2.7.13.3) (nm23-H2) | 152 | NME2 | 5 | 10 | 6 | 28.8 | 55.4 | 34.5 |
| 619 | -0.46 | -0.35 | 1.44 | O15067 | Phosphoribosylformylglycinamidine synthase (FGAM synthase) (FGAMS) (EC 6.3.5.3) (Formylglycinamide ribonucleotide amidotransferase) (FGAR amidotransferase) (FGAR-AT) (Formylglycinamide ribotide amidotransferase) | 1338 | PFAS | 2 | 1 | 3 | 2.2 | 0.7 | 3.2 |
| 620 | -0.46 | 1.26 | 0.00 | Q5T4L4 | 40S ribosomal protein S27 | 66 | RPS27 | 1 | 2 | 0 | 19.7 | 36.4 | 0 |
| 621 | -0.46 | 0.47 | 0.72 | Q01813 | ATP-dependent 6-phosphofructokinase, platelet type (ATP-PFK) (PFK-P) (EC 2.7.1.11) (6-phosphofructokinase type C) (Phosphofructo-1-kinase isozyme C) (PFK-C) (Phosphohexokinase) | 784 | PFKP | 4 | 4 | 4 | 5.5 | 4.3 | 5.5 |
| 622 | -0.46 | -0.69 | 0.06 | P10599 | Thioredoxin (Trx) (ATL-derived factor) (ADF) (Surface-associated sulphydryl protein) (SASP) | 105 | TXN | 5 | 6 | 3 | 42.9 | 51.4 | 32.4 |

| | | | | | | | | | | | | | |
|---|---|---|---|---|---|---|---|---|---|---|---|---|---|
| 623 | -0.46 | 0.87 | 0.57 | P32322 | Pyrroline-5-carboxylate reductase 1, mitochondrial (P5C reductase 1) (P5CR 1) (EC 1.5.1.2) | 319 | PYCR1 | 1 | 1 | 2 | 3.4 | 3.4 | 7.2 |
| 624 | -0.45 | 0.00 | 0.00 | H0YLC0 | Tumor necrosis factor alpha-induced protein 2 (Fragment) | 347 | TNFAIP2 | 1 | 0 | 0 | 3.2 | 0 | 0 |
| 625 | -0.45 | -0.32 | 0.38 | Q99598 | Translin-associated protein X (Translin-associated factor X) | 290 | TSNAX | 1 | 1 | 2 | 3.4 | 6.9 | 10.3 |
| 626 | -0.45 | 0.92 | 0.28 | F5H7F6 | Microsomal glutathione S-transferase 1 (Fragment) | 77 | MGST1 | 1 | 1 | 1 | 18.2 | 16.9 | 16.9 |
| 627 | -0.45 | -0.43 | 0.03 | D6RFZ2 | TBC1 domain family member 8B | 419 | TBC1D8B | 1 | 1 | 1 | 2.6 | 2.6 | 2.6 |
| 628 | -0.45 | 0.68 | 1.74 | P49327 | Fatty acid synthase (EC 2.3.1.85) [Includes: [Acyl-carrier-protein] S-acetyltransferase (EC 2.3.1.38); [Acyl-carrier-protein] S-malonyltransferase (EC 2.3.1.39); 3-oxoacyl-[acyl-carrier-protein] synthase (EC 2.3.1.41); 3-oxoacyl-[acyl-carrier-protein] reductase (EC 1.1.1.100); 3-hydroxyacyl-[acyl-carrier-protein] dehydratase (EC 4.2.1.59); Enoyl-[acyl-carrier-protein] reductase (EC 1.3.1.39); Oleoyl-[acyl-carrier-protein] hydrolase (EC 3.1.2.14)] | 2511 | FASN | 10 | 17 | 20 | 5.5 | 10.4 | 11.6 |
| 629 | -0.45 | 0.19 | 0.18 | O95373 | Importin-7 (Imp7) (Ran-binding protein 7) (RanBP7) | 1038 | IPO7 | 6 | 5 | 5 | 6.9 | 6.3 | 6.4 |
| 630 | -0.45 | 0.00 | -0.56 | D6RA82 | Annexin | 284 | ANXA3 | 3 | 0 | 3 | 14.4 | 0 | 12.3 |
| 631 | -0.45 | 0.48 | 0.71 | K9J957 | Proteasome activator complex subunit 3 (REG gamma-3 variant) | 231 | PSME3 | 4 | 3 | 4 | 20.8 | 15.2 | 15.6 |

| | | | | | | | | | | | | | |
|---|---|---|---|---|---|---|---|---|---|---|---|---|---|
| 632 | -0.45 | 1.04 | 0.52 | Q8NBS9 | Thioredoxin domain-containing protein 5 (Endoplasmic reticulum resident protein 46) (ER protein 46) (ERp46) (Thioredoxin-like protein p46) | 432 | TXNDC5 | 3 | 2 | 4 | 9 | 5.9 | 11.7 |
| 633 | -0.44 | 0.00 | 1.36 | Q9Y224 | UPF0568 protein C14orf166 (CLE7 homolog) (CLE) | 244 | C14orf166 | 2 | 0 | 3 | 10.2 | 0 | 16.8 |
| 634 | -0.44 | -0.30 | 0.36 | P49588 | Alanine--tRNA ligase, cytoplasmic (EC 6.1.1.7) (Alanyl-tRNA synthetase) (AlaRS) (Renal carcinoma antigen NY-REN-42) | 968 | AARS | 7 | 4 | 7 | 8 | 4.3 | 8 |
| 635 | -0.44 | 0.85 | 0.75 | Q12931 | Heat shock protein 75 kDa, mitochondrial (HSP 75) (TNFR-associated protein 1) (Tumor necrosis factor type 1 receptor-associated protein) (TRAP-1) | 704 | TRAP1 | 4 | 3 | 3 | 9.4 | 6.1 | 5.7 |
| 636 | -0.44 | 1.14 | 0.72 | P13667 | Protein disulfide-isomerase A4 (EC 5.3.4.1) (Endoplasmic reticulum resident protein 70) (ER protein 70) (ERp70) (Endoplasmic reticulum resident protein 72) (ER protein 72) (ERp-72) (ERp72) | 645 | PDIA4 | 7 | 7 | 11 | 14.9 | 13.5 | 23.4 |
| 637 | -0.44 | -0.52 | 0.56 | P52209 | 6-phosphogluconate dehydrogenase, decarboxylating (EC 1.1.1.44) | 483 | PGD | 9 | 10 | 13 | 25.5 | 23.6 | 33.6 |
| 638 | -0.44 | -0.02 | 0.25 | B8ZZU8 | Transcription elongation factor B (SIII), polypeptide 2 (18kDa, elongin B), isoform CRA_b (Transcription elongation factor B polypeptide 2) | 113 | TCEB2 | 2 | 2 | 2 | 16.8 | 17.7 | 16.8 |

| | | | | | | | | | | | | | |
|---|---|---|---|---|---|---|---|---|---|---|---|---|---|
| 639 | -0.44 | 0.00 | 0.43 | Q2TAY7 | WD40 repeat-containing protein SMU1 (Smu-1 suppressor of mec-8 and unc-52 protein homolog) [Cleaved into: WD40 repeat-containing protein SMU1, N-terminally processed] | 513 | SMU1 | 1 | 0 | 1 | 1.9 | 0 | 1.9 |
| 640 | -0.44 | 0.00 | 0.05 | Q7L2H7 | Eukaryotic translation initiation factor 3 subunit M (eIF3m) (Fetal lung protein B5) (hFL-B5) (PCI domain-containing protein 1) | 374 | EIF3M | 3 | 0 | 3 | 12 | 0 | 9.1 |
| 641 | -0.44 | -0.72 | 0.00 | Q9BRG1 | Vacuolar protein-sorting-associated protein 25 (hVps25) (Dermal papilla-derived protein 9) (ELL-associated protein of 20 kDa) (ESCRT-II complex subunit VPS25) | 176 | VPS25 | 1 | 2 | 0 | 5.7 | 10.2 | 0 |
| 642 | -0.43 | 0.35 | 0.39 | Q14974 | Importin subunit beta-1 (Importin-90) (Karyopherin subunit beta-1) (Nuclear factor p97) (Pore targeting complex 97 kDa subunit) (PTAC97) | 876 | KPNB1 | 12 | 6 | 9 | 20 | 9.2 | 13.4 |
| 643 | -0.43 | 0.25 | 0.50 | P29401 | Transketolase (TK) (EC 2.2.1.1) | 623 | TKT | 20 | 13 | 14 | 40.6 | 31 | 23.1 |
| 644 | -0.43 | -0.05 | 0.39 | I3L397 | Eukaryotic translation initiation factor 5A (eIF-5A) (Fragment) | 146 | EIF5A | 7 | 5 | 8 | 34.9 | 37 | 37.7 |
| 645 | -0.43 | -0.26 | 0.68 | Q04760 | Lactoylglutathione lyase (EC 4.4.1.5) (Aldoketomutase) (Glyoxalase I) (Glx I) (Ketone-aldehyde mutase) (Methylglyoxalase) (S-D-lactoylglutathione methylglyoxal lyase) | 184 | GLO1 | 3 | 4 | 4 | 19.5 | 26 | 29 |
| 646 | -0.42 | 1.22 | -0.22 | H3BN98 | Uncharacterized protein (Fragment) | 237 | | 3 | 2 | 2 | 11 | 8 | 8 |
| 647 | -0.42 | 0.00 | 0.16 | P46109 | Crk-like protein | 303 | CRKL | 1 | 0 | 1 | 4 | 0 | 4 |
| 648 | -0.42 | 0.00 | 0.29 | A6NDG6 | Phosphoglycolate phosphatase (PGP) (PGPase) (EC 3.1.3.18) (EC 3.1.3.48) | 321 | PGP | 2 | 0 | 1 | 9 | 0 | 4.7 |

| | | | | | | | | | | | | | |
|---|---|---|---|---|---|---|---|---|---|---|---|---|---|
| 649 | -0.42 | 0.42 | 1.36 | O75643 | U5 small nuclear ribonucleoprotein 200 kDa helicase (EC 3.6.4.13) (Activating signal cointegrator 1 complex subunit 3-like 1) (BRR2 homolog) (U5 snRNP-specific 200 kDa protein) (U5-200KD) | 2136 | SNRNP200 | 1 | 3 | 3 | 0.5 | 1.7 | 1.9 |
| 650 | -0.42 | 0.49 | 0.14 | F8W914 | Reticulon | 345 | RTN4 | 5 | 5 | 4 | 18.8 | 18.8 | 15.1 |
| 651 | -0.42 | 0.00 | 0.00 | H7C131 | 3-ketoacyl-CoA thiolase, peroxisomal (Fragment) | 290 | ACAA1 | 1 | 0 | 0 | 4.5 | 0 | 0 |
| 652 | -0.42 | -0.27 | 0.40 | P00492 | Hypoxanthine-guanine phosphoribosyltransferase (HGPRT) (HGPRTase) (EC 2.4.2.8) | 218 | HPRT1 | 5 | 8 | 6 | 25.7 | 43.1 | 32.1 |
| 653 | -0.42 | 1.42 | 1.12 | P55084 | Trifunctional enzyme subunit beta, mitochondrial (TP-beta) [Includes: 3-ketoacyl-CoA thiolase (EC 2.3.1.16) (Acetyl-CoA acyltransferase) (Beta-ketothiolase)] | 474 | HADHB | 2 | 1 | 5 | 4.2 | 1.7 | 12 |
| 654 | -0.42 | -0.07 | 0.54 | P23526 | Adenosylhomocysteinase (AdoHcyase) (EC 3.3.1.1) (S-adenosyl-L-homocysteine hydrolase) | 432 | AHCY | 4 | 4 | 6 | 12 | 12 | 19.7 |
| 655 | -0.42 | 0.56 | 0.61 | H0YAM7 | Guanine nucleotide-binding protein subunit beta-2-like 1 (Fragment) | 247 | GNB2L1 | 3 | 7 | 3 | 25.5 | 43.7 | 25.5 |
| 656 | -0.41 | 0.50 | 0.89 | P09651 | Heterogeneous nuclear ribonucleoprotein A1 (hnRNP A1) (Helix-destabilizing protein) (Single-strand RNA-binding protein) (hnRNP core protein A1) [Cleaved into: Heterogeneous nuclear ribonucleoprotein A1, N-terminally processed] | 372 | HNRNPA1 | 2 | 6 | 3 | 14.2 | 29.6 | 13.9 |
| 657 | -0.41 | 0.38 | 0.51 | P47813 | Eukaryotic translation initiation factor 1A, X-chromosomal (eIF-1A X isoform) (Eukaryotic translation initiation factor 4C) (eIF-4C) | 144 | EIF1AX | 3 | 4 | 5 | 26.4 | 25 | 33.3 |

| | | | | | | | | | | | | | |
|---|---|---|---|---|---|---|---|---|---|---|---|---|---|
| 658 | -0.41 | 0.00 | 0.52 | Q92600 | Cell differentiation protein RCD1 homolog (Rcd-1) (CCR4-NOT transcription complex subunit 9) | 299 | RQCD1 | 1 | 1 | 2 | 3.5 | 4.7 | 8.1 |
| 659 | -0.41 | -0.20 | 0.29 | H0YKT8 | Proteasome subunit beta type (EC 3.4.25.1) (Fragment) | 181 | PSMA4 | 4 | 3 | 3 | 30.9 | 14.9 | 18.8 |
| 660 | -0.40 | -0.48 | 0.19 | P55327 | Tumor protein D52 (Protein N8) | 224 | TPD52 | 2 | 3 | 2 | 15.8 | 20.7 | 15.8 |
| 661 | -0.40 | -0.07 | 0.59 | P37837 | Transaldolase (EC 2.2.1.2) | 337 | TALDO1 | 6 | 8 | 5 | 16.9 | 26.1 | 11.6 |
| 662 | -0.40 | 0.76 | 1.25 | O15355 | Protein phosphatase 1G (EC 3.1.3.16) (Protein phosphatase 1C) (Protein phosphatase 2C isoform gamma) (PP2C-gamma) (Protein phosphatase magnesium-dependent 1 gamma) | 546 | PPM1G | 1 | 1 | 3 | 2.2 | 2.2 | 6.6 |
| 663 | -0.40 | 0.37 | 1.67 | P18124 | 60S ribosomal protein L7 | 248 | RPL7 | 3 | 4 | 4 | 15.7 | 19.8 | 16.5 |
| 664 | -0.40 | -0.36 | -0.81 | Q13501 | Sequestosome-1 (EBI3-associated protein of 60 kDa) (EBIAP) (p60) (Phosphotyrosine-independent ligand for the Lck SH2 domain of 62 kDa) (Ubiquitin-binding protein p62) | 440 | SQSTM1 | 2 | 1 | 1 | 4.8 | 1.4 | 3.4 |
| 665 | -0.39 | 1.11 | 0.00 | G3V5X4 | Nesprin-2 | 6818 | SYNE2 | 1 | 3 | 1 | 0.1 | 0.4 | 0.1 |
| 666 | -0.39 | 0.00 | 0.51 | Q9Y305 | Acyl-coenzyme A thioesterase 9, mitochondrial (Acyl-CoA thioesterase 9) (EC 3.1.2.-) (Acyl-CoA thioester hydrolase 9) | 439 | ACOT9 | 2 | 0 | 3 | 5.8 | 0 | 7.1 |
| 667 | -0.39 | 0.62 | 0.45 | E9PJK1 | Tetraspanin | 165 | CD81 | 3 | 2 | 3 | 35.8 | 23.6 | 35.8 |
| 668 | -0.39 | 0.14 | 0.60 | O15144 | Actin-related protein 2/3 complex subunit 2 (Arp2/3 complex 34 kDa subunit) (p34-ARC) | 300 | ARPC2 | 3 | 3 | 4 | 9.3 | 8.7 | 12.3 |
| 669 | -0.38 | 0.94 | 1.09 | Q9NSE4 | Isoleucine--tRNA ligase, mitochondrial (EC 6.1.1.5) (Isoleucyl-tRNA synthetase) (IleRS) | 1012 | IARS2 | 6 | 6 | 10 | 6.9 | 6.9 | 12.2 |
| 670 | -0.38 | 1.06 | 1.14 | P31327 | Carbamoyl-phosphate synthase [ammonia], mitochondrial (EC 6.3.4.16) (Carbamoyl-phosphate synthetase I) (CPSase I) | 1500 | CPS1 | 2 | 5 | 4 | 1.7 | 4.1 | 3.3 |

| | | | | | | | | | | | | | |
|---|---|---|---|---|---|---|---|---|---|---|---|---|---|
| 671 | -0.38 | 1.13 | 0.65 | O95881 | Thioredoxin domain-containing protein 12 (EC 1.8.4.2) (Endoplasmic reticulum resident protein 18) (ER protein 18) (ERp18) (Endoplasmic reticulum resident protein 19) (ER protein 19) (ERp19) (Thioredoxin-like protein p19) (hTLP19) | 172 | TXNDC12 | 2 | 3 | 3 | 13.4 | 22.1 | 22.1 |
| 672 | -0.38 | -0.29 | 0.62 | H0YN26 | Acidic leucine-rich nuclear phosphoprotein 32 family member A | 177 | ANP32A | 3 | 3 | 3 | 18.6 | 18.6 | 11.9 |
| 673 | -0.38 | -0.43 | 0.29 | P15531 | Nucleoside diphosphate kinase A (NDK A) (NDP kinase A) (EC 2.7.4.6) (Granzyme A-activated DNase) (GAAD) (Metastasis inhibition factor nm23) (NM23-H1) (Tumor metastatic process-associated protein) | 152 | NME1 | 5 | 6 | 5 | 36.8 | 42.8 | 36.8 |
| 674 | -0.38 | 0.07 | 0.31 | A0A087WZB3 | Proline and serine-rich protein 3 | 324 | PROSER3 | 1 | 1 | 1 | 2.8 | 2.8 | 2.8 |
| 675 | -0.38 | 2.00 | 1.11 | Q07955 | Serine/arginine-rich splicing factor 1 (Alternative-splicing factor 1) (ASF-1) (Splicing factor, arginine/serine-rich 1) (pre-mRNA-splicing factor SF2, P33 subunit) | 248 | SRSF1 | 1 | 1 | 2 | 4 | 5.6 | 8.5 |
| 676 | -0.37 | 0.09 | 0.08 | P29692 | Elongation factor 1-delta (EF-1-delta) (Antigen NY-CO-4) | 281 | EEF1D | 4 | 6 | 3 | 20.3 | 30.2 | 11.7 |
| 677 | -0.37 | 0.33 | 0.91 | I3L1L3 | Myb-binding protein 1A (Fragment) | 1252 | MYBBP1A | 3 | 4 | 4 | 2.7 | 3.7 | 3.7 |
| 678 | -0.37 | -0.21 | -0.12 | Q96A49 | Synapse-associated protein 1 | 352 | SYAP1 | 1 | 1 | 1 | 4.3 | 4.3 | 4.3 |
| 679 | -0.37 | 0.20 | 0.63 | P11216 | Glycogen phosphorylase, brain form (EC 2.4.1.1) | 843 | PYGB | 3 | 3 | 6 | 3.1 | 3.1 | 8.4 |
| 680 | -0.37 | 0.25 | 1.33 | P55060 | Exportin-2 (Exp2) (Cellular apoptosis susceptibility protein) (Chromosome segregation 1-like protein) (Importin-alpha re-exporter) | 971 | CSE1L | 10 | 7 | 9 | 11.5 | 11.1 | 11.3 |

| | | | | | | | | | | | | | |
|---|---|---|---|---|---|---|---|---|---|---|---|---|---|
| 681 | -0.37 | -0.75 | 0.56 | D6RE83 | Ubiquitin carboxyl-terminal hydrolase (EC 3.4.19.12) | 207 | UCHL1 | 5 | 6 | 6 | 34.3 | 36.7 | 41.5 |
| 682 | -0.37 | -0.45 | 0.98 | F8WBS5 | 60S ribosomal protein L35a | 55 | RPL35A | 1 | 1 | 1 | 16.4 | 16.4 | 16.4 |
| 683 | -0.36 | -0.06 | 0.35 | Q9BTT0 | Acidic leucine-rich nuclear phosphoprotein 32 family member E (LANP-like protein) (LANP-L) | 268 | ANP32E | 2 | 4 | 2 | 15 | 20.9 | 15 |
| 684 | -0.36 | 1.07 | 0.60 | Q9GZT3 | SRA stem-loop-interacting RNA-binding protein, mitochondrial | 109 | SLIRP | 1 | 4 | 2 | 13.1 | 48.6 | 24.3 |
| 685 | -0.36 | 1.65 | 0.69 | P35232 | Prohibitin | 272 | PHB | 3 | 7 | 2 | 9.9 | 37.9 | 6.6 |
| 686 | -0.36 | 0.63 | 0.47 | Q13813 | Spectrin alpha chain, non-erythrocytic 1 (Alpha-II spectrin) (Fodrin alpha chain) (Spectrin, non-erythroid alpha subunit) | 2472 | SPTAN1 | 21 | 29 | 21 | 13.5 | 15.7 | 12.7 |
| 687 | -0.36 | -0.25 | 0.37 | Q15185 | Prostaglandin E synthase 3 (EC 5.3.99.3) (Cytosolic prostaglandin E2 synthase) (cPGES) (Hsp90 co-chaperone) (Progesterone receptor complex p23) (Telomerase-binding protein p23) | 160 | PTGES3 | 3 | 4 | 4 | 24.6 | 36.9 | 30.8 |
| 688 | -0.36 | 0.51 | 0.54 | Q9Y265 | RuvB-like 1 (EC 3.6.4.12) (49 kDa TATA box-binding protein-interacting protein) (49 kDa TBP-interacting protein) (54 kDa erythrocyte cytosolic protein) (ECP-54) (INO80 complex subunit H) (Nuclear matrix protein 238) (NMP 238) (Pontin 52) (TIP49a) (TIP60-associated protein 54-alpha) (TAP54-alpha) | 456 | RUVBL1 | 6 | 5 | 8 | 17.3 | 16 | 23 |
| 689 | -0.36 | 0.00 | 0.21 | Q15526 | Surfeit locus protein 1 | 300 | SURF1 | 1 | 0 | 1 | 5.6 | 0 | 5.6 |
| 690 | -0.36 | 1.91 | 0.00 | Q6P1J9 | Parafibromin (Cell division cycle protein 73 homolog) (Hyperparathyroidism 2 protein) | 531 | CDC73 | 2 | 1 | 0 | 5.6 | 3.8 | 0 |

| | | | | | | | | | | | | | |
|---|---|---|---|---|---|---|---|---|---|---|---|---|---|
| 691 | -0.36 | 0.21 | 0.20 | P25786 | Proteasome subunit alpha type-1 (EC 3.4.25.1) (30 kDa prosomal protein) (PROS-30) (Macropain subunit C2) (Multicatalytic endopeptidase complex subunit C2) (Proteasome component C2) (Proteasome nu chain) | 263 | PSMA1 | 4 | 6 | 5 | 17.9 | 26.2 | 20.2 |
| 692 | -0.35 | 0.15 | 0.66 | O94925 | Glutaminase kidney isoform, mitochondrial (GLS) (EC 3.5.1.2) (K-glutaminase) (L-glutamine amidohydrolase) | 669 | GLS | 3 | 1 | 4 | 6.2 | 1.7 | 10.5 |
| 693 | -0.35 | -0.36 | 0.48 | B1AK87 | Capping protein (Actin filament) muscle Z-line, beta, isoform CRA_a (F-actin-capping protein subunit beta) | 260 | CAPZB | 1 | 1 | 1 | 3.8 | 3.8 | 3.8 |
| 694 | -0.35 | -0.54 | -0.12 | C9JD53 | Isopentenyl-diphosphate Delta-isomerase 1 (Fragment) | 120 | IDI1 | 1 | 1 | 1 | 10 | 10 | 10 |
| 695 | -0.35 | 0.71 | 0.43 | P50454 | Serpin H1 (47 kDa heat shock protein) (Arsenic-transactivated protein 3) (AsTP3) (Cell proliferation-inducing gene 14 protein) (Collagen-binding protein) (Colligin) (Rheumatoid arthritis-related antigen RA-A47) | 418 | SERPINH1 | 4 | 7 | 4 | 12.7 | 27.3 | 12.2 |
| 696 | -0.35 | 0.50 | 1.34 | P52758 | Ribonuclease UK114 (EC 3.1.-.-) (14.5 kDa translational inhibitor protein) (p14.5) (Heat-responsive protein 12) (UK114 antigen homolog) | 137 | HRSP12 | 1 | 3 | 1 | 7.3 | 27 | 7.3 |
| 697 | -0.35 | 0.23 | 0.54 | P49755 | Transmembrane emp24 domain-containing protein 10 (21 kDa transmembrane-trafficking protein) (S31III125) (S31I125) (Tmp-21-I) (Transmembrane protein Tmp21) (p23) (p24 family protein delta-1) (p24delta1) (p24delta) | 219 | TMED10 | 3 | 4 | 3 | 17.8 | 21 | 17.8 |

| | | | | | | | | | | | | | |
|---|---|---|---|---|---|---|---|---|---|---|---|---|---|
| 698 | -0.35 | -0.11 | 0.63 | P26447 | Protein S100-A4 (Calvasculin) (Metastasin) (Placental calcium-binding protein) (Protein Mts1) (S100 calcium-binding protein A4) | 101 | S100A4 | 2 | 5 | 2 | 18.8 | 33.7 | 18.8 |
| 699 | -0.35 | 0.00 | 1.34 | Q9UJS0 | Calcium-binding mitochondrial carrier protein Aralar2 (Citrin) (Mitochondrial aspartate glutamate carrier 2) (Solute carrier family 25 member 13) | 675 | SLC25A13 | 1 | 0 | 1 | 1.8 | 0 | 3.3 |
| 700 | -0.35 | 0.80 | -0.31 | P05783 | Keratin, type I cytoskeletal 18 (Cell proliferation-inducing gene 46 protein) (Cytokeratin-18) (CK-18) (Keratin-18) (K18) | 430 | KRT18 | 6 | 7 | 5 | 24.7 | 21.4 | 20.5 |
| 701 | -0.35 | 0.88 | -0.08 | D6R967 | Inorganic pyrophosphatase 2, mitochondrial (Fragment) | 191 | PPA2 | 3 | 1 | 1 | 17.3 | 6.8 | 5.8 |
| 702 | -0.34 | -0.56 | 0.09 | H0YI98 | Dynactin subunit 2 (Fragment) | 268 | DCTN2 | 1 | 1 | 1 | 3.4 | 3.4 | 3.4 |
| 703 | -0.34 | 0.80 | 0.40 | P24539 | ATP synthase F(0) complex subunit B1, mitochondrial (ATP synthase proton-transporting mitochondrial F(0) complex subunit B1) (ATP synthase subunit b) (ATPase subunit b) | 256 | ATP5F1 | 5 | 6 | 4 | 21.5 | 27 | 15.6 |
| 704 | -0.34 | 0.54 | 0.24 | Q9NP72 | Ras-related protein Rab-18 | 206 | RAB18 | 4 | 4 | 2 | 22.8 | 22.8 | 10.2 |
| 705 | -0.34 | -0.03 | -0.04 | K7EM62 | GPI ethanolamine phosphate transferase 1 (Fragment) | 257 | PIGN | 1 | 1 | 1 | 6.2 | 6.2 | 6.2 |
| 706 | -0.34 | 0.00 | 0.36 | O43172 | U4/U6 small nuclear ribonucleoprotein Prp4 (PRP4 homolog) (hPrp4) (U4/U6 snRNP 60 kDa protein) (WD splicing factor Prp4) | 522 | PRPF4 | 1 | 0 | 1 | 2.5 | 0 | 2.5 |
| 707 | -0.33 | 0.27 | 0.65 | P51572 | B-cell receptor-associated protein 31 (BCR-associated protein 31) (Bap31) (6C6-AG tumor-associated antigen) (Protein CDM) (p28) | 246 | BCAP31 | 6 | 5 | 4 | 26.8 | 22.4 | 18.7 |

| | | | | | | | | | | | | | |
|---|---|---|---|---|---|---|---|---|---|---|---|---|---|
| 708 | -0.33 | -0.31 | 0.68 | P35241 | Radixin | 583 | RDX | 12 | 5 | 8 | 19.2 | 8.2 | 11.8 |
| 709 | -0.33 | 0.00 | 0.29 | P42166 | Lamina-associated polypeptide 2, isoform alpha (Thymopoietin isoform alpha) (TP alpha) (Thymopoietin-related peptide isoform alpha) (TPRP isoform alpha) [Cleaved into: Thymopoietin (TP) (Splenin); Thymopentin (TP5)] | 694 | TMPO | 1 | 0 | 2 | 2.9 | 0 | 5.2 |
| 710 | -0.33 | 0.13 | 0.23 | Q9Y3A6 | Transmembrane emp24 domain-containing protein 5 (p24 family protein gamma-2) (p24gamma2) (p28) | 229 | TMED5 | 1 | 2 | 1 | 5.2 | 10 | 5.2 |
| 711 | -0.33 | 0.89 | 1.13 | P25787 | Proteasome subunit alpha type-2 (EC 3.4.25.1) (Macropain subunit C3) (Multicatalytic endopeptidase complex subunit C3) (Proteasome component C3) | 234 | PSMA2 | 4 | 8 | 8 | 27.4 | 50 | 47.4 |
| 712 | -0.33 | 1.72 | 1.33 | P40939 | Trifunctional enzyme subunit alpha, mitochondrial (78 kDa gastrin-binding protein) (TP-alpha) [Includes: Long-chain enoyl-CoA hydratase (EC 4.2.1.17); Long chain 3-hydroxyacyl-CoA dehydrogenase (EC 1.1.1.211)] | 763 | HADHA | 8 | 3 | 9 | 15.6 | 5.8 | 15.5 |
| 713 | -0.32 | 0.32 | 0.84 | P46940 | Ras GTPase-activating-like protein IQGAP1 (p195) | 1657 | IQGAP1 | 10 | 16 | 12 | 7.8 | 13.5 | 10.9 |
| 714 | -0.31 | 0.69 | 0.00 | Q86SX6 | Glutaredoxin-related protein 5, mitochondrial (Monothiol glutaredoxin-5) | 157 | GLRX5 | 1 | 2 | 0 | 7 | 15.9 | 0 |
| 715 | -0.31 | 0.00 | 0.00 | F8WDI0 | Ubiquitin-like-conjugating enzyme ATG3 | 74 | ATG3 | 1 | 0 | 1 | 12.2 | 0 | 12.2 |

| | | | | | | | | | | | | | |
|---|---|---|---|---|---|---|---|---|---|---|---|---|---|
| 716 | -0.31 | 0.15 | 0.90 | Q9Y3B4 | Splicing factor 3B subunit 6 (Pre-mRNA branch site protein p14) (SF3b 14 kDa subunit) (SF3B14a) (Spliceosome-associated protein, 14-kDa) (Splicing factor 3b, subunit 6, 14kDa) | 125 | SF3B6 | 3 | 2 | 2 | 25.6 | 20.8 | 20.8 |
| 717 | -0.31 | 0.39 | 0.41 | Q15417 | Calponin-3 (Calponin, acidic isoform) | 329 | CNN3 | 2 | 1 | 2 | 7.8 | 3.9 | 7.8 |
| 718 | -0.31 | 1.47 | 1.53 | Q8IWJ2 | GRIP and coiled-coil domain-containing protein 2 (185 kDa Golgi coiled-coil protein) (GCC185) (CLL-associated antigen KW-11) (CTCL tumor antigen se1-1) (Ran-binding protein 2-like 4) (RanBP2L4) (Renal carcinoma antigen NY-REN-53) | 1684 | GCC2 | 1 | 2 | 1 | 0.5 | 1.1 | 0.5 |
| 719 | -0.31 | 0.00 | 0.16 | O43933 | Peroxisome biogenesis factor 1 (Peroxin-1) (Peroxisome biogenesis disorder protein 1) | 1283 | PEX1 | 1 | 0 | 1 | 0.9 | 0 | 0.9 |
| 720 | -0.31 | 0.00 | 0.07 | P41227 | N-alpha-acetyltransferase 10 (EC 2.3.1.-) (EC 2.3.1.88) (N-terminal acetyltransferase complex ARD1 subunit homolog A) (NatA catalytic subunit Naa10) | 235 | NAA10 | 2 | 0 | 2 | 6.4 | 0 | 6.4 |
| 721 | -0.31 | 0.00 | 0.96 | Q9P287 | BRCA2 and CDKN1A-interacting protein (P21- and CDK-associated protein 1) (Protein TOK-1) | 314 | BCCIP | 1 | 0 | 2 | 2.7 | 0 | 8.2 |
| 722 | -0.31 | 0.00 | 0.41 | P43686 | 26S protease regulatory subunit 6B (26S proteasome AAA-ATPase subunit RPT3) (MB67-interacting protein) (MIP224) (Proteasome 26S subunit ATPase 4) (Tat-binding protein 7) (TBP-7) | 418 | PSMC4 | 4 | 0 | 6 | 14.5 | 0 | 17.3 |
| 723 | -0.30 | -0.01 | 0.43 | P60842 | Eukaryotic initiation factor 4A-I (eIF-4A-I) (eIF4A-I) (EC 3.6.4.13) (ATP-dependent RNA helicase eIF4A-1) | 406 | EIF4A1 | 7 | 8 | 9 | 21.2 | 20.2 | 21.4 |

| | | | | | | | | | | | | | |
|---|---|---|---|---|---|---|---|---|---|---|---|---|---|
| 724 | -0.30 | 0.00 | 0.90 | D6RD67 | Methylcrotonoyl-CoA carboxylase beta chain, mitochondrial (Fragment) | 286 | MCCC2 | 1 | 0 | 1 | 4.2 | 0 | 4.2 |
| 725 | -0.30 | 0.12 | 0.02 | F8WBH7 | Proteasome assembly chaperone 1 | 63 | PSMG1 | 1 | 1 | 1 | 15.9 | 15.9 | 15.9 |
| 726 | -0.30 | 0.43 | 0.70 | Q9NPJ3 | Acyl-coenzyme A thioesterase 13 (Acyl-CoA thioesterase 13) (EC 3.1.2.-) (Thioesterase superfamily member 2) [Cleaved into: Acyl-coenzyme A thioesterase 13, N-terminally processed] | 140 | ACOT13 | 4 | 3 | 4 | 45.3 | 29.9 | 45.3 |
| 727 | -0.30 | 0.00 | 0.55 | E7EMC6 | Annexin | 330 | ANXA6 | 2 | 0 | 1 | 7.9 | 0 | 3.3 |
| 728 | -0.30 | 0.58 | 0.36 | P62820 | Ras-related protein Rab-1A (YPT1-related protein) | 205 | RAB1A | 4 | 8 | 5 | 22.9 | 42.4 | 36.6 |
| 729 | -0.29 | -0.15 | 0.35 | P28066 | Proteasome subunit alpha type-5 (EC 3.4.25.1) (Macropain zeta chain) (Multicatalytic endopeptidase complex zeta chain) (Proteasome zeta chain) | 241 | PSMA5 | 5 | 5 | 6 | 30.7 | 27 | 33.6 |
| 730 | -0.29 | 0.59 | 0.11 | P40925 | Malate dehydrogenase, cytoplasmic (EC 1.1.1.37) (Cytosolic malate dehydrogenase) (Diiodophenylpyruvate reductase) (EC 1.1.1.96) | 334 | MDH1 | 6 | 2 | 3 | 19.5 | 5.7 | 10.5 |
| 731 | -0.29 | 0.01 | 0.53 | Q9Y490 | Talin-1 | 2541 | TLN1 | 22 | 14 | 19 | 13.2 | 9.1 | 11.6 |
| 732 | -0.29 | 0.02 | 0.85 | P00558 | Phosphoglycerate kinase 1 (EC 2.7.2.3) (Cell migration-inducing gene 10 protein) (Primer recognition protein 2) (PRP 2) | 417 | PGK1 | 15 | 12 | 19 | 37.2 | 33.1 | 49.4 |
| 733 | -0.29 | 0.11 | 0.00 | F5GWX2 | Heme-binding protein 1 | 133 | HEBP1 | 2 | 1 | 0 | 15.8 | 12.8 | 0 |

| | | | | | | | | | | | | | |
|---|---|---|---|---|---|---|---|---|---|---|---|---|---|
| 734 | -0.29 | 0.04 | 0.36 | P63010 | AP-2 complex subunit beta (AP105B) (Adaptor protein complex AP-2 subunit beta) (Adaptor-related protein complex 2 subunit beta) (Beta-2-adaptin) (Beta-adaptin) (Clathrin assembly protein complex 2 beta large chain) (Plasma membrane adaptor HA2/AP2 adaptin beta subunit) | 937 | AP2B1 | 5 | 2 | 3 | 7.7 | 3.6 | 4.4 |
| 735 | -0.29 | 0.23 | 0.20 | P06753 | Tropomyosin alpha-3 chain (Gamma-tropomyosin) (Tropomyosin-3) (Tropomyosin-5) (hTM5) | 285 | TPM3 | 6 | 5 | 7 | 25.4 | 20.6 | 27 |
| 736 | -0.29 | 0.34 | 0.71 | P61224 | Ras-related protein Rap-1b (GTP-binding protein smg p21B) | 184 | RAP1B | 2 | 3 | 2 | 15.8 | 20.6 | 15.8 |
| 737 | -0.28 | 0.00 | 1.76 | P17844 | Probable ATP-dependent RNA helicase DDX5 (EC 3.6.4.13) (DEAD box protein 5) (RNA helicase p68) | 614 | DDX5 | 2 | 0 | 1 | 3.4 | 0 | 2.6 |
| 738 | -0.28 | 1.67 | 0.37 | Q9H9B4 | Sideroflexin-1 (Tricarboxylate carrier protein) (TCC) | 322 | SFXN1 | 3 | 3 | 1 | 14.9 | 13.7 | 4.7 |
| 739 | -0.28 | -0.30 | 0.04 | P63104 | 14-3-3 protein zeta/delta (Protein kinase C inhibitor protein 1) (KCIP-1) | 245 | YWHAZ | 11 | 11 | 11 | 40.8 | 48.2 | 38.4 |
| 740 | -0.28 | 0.14 | 0.68 | O60701 | UDP-glucose 6-dehydrogenase (UDP-Glc dehydrogenase) (UDP-GlcDH) (UDPGDH) (EC 1.1.1.22) | 494 | UGDH | 10 | 12 | 14 | 27.7 | 34.6 | 31 |
| 741 | -0.28 | -0.49 | -0.24 | Q9BRA2 | Thioredoxin domain-containing protein 17 (14 kDa thioredoxin-related protein) (TRP14) (Protein 42-9-9) (Thioredoxin-like protein 5) | 123 | TXNDC17 | 2 | 2 | 2 | 15.4 | 15.4 | 15.4 |
| 742 | -0.28 | 0.69 | 0.41 | Q99613 | Eukaryotic translation initiation factor 3 subunit C (eIF3c) (Eukaryotic translation initiation factor 3 subunit 8) (eIF3 p110) | 913 | EIF3C | 2 | 2 | 1 | 2.3 | 2.2 | 1.2 |

| | | | | | | | | | | | | | |
|---|---|---|---|---|---|---|---|---|---|---|---|---|---|
| 743 | -0.28 | -0.31 | 0.35 | B1ALC0 | Actin-related protein 2/3 complex subunit 5 | 135 | ARPC5 | 1 | 1 | 1 | 9.6 | 9.6 | 9.6 |
| 744 | -0.28 | -0.35 | 0.53 | Q96KP4 | Cytosolic non-specific dipeptidase (EC 3.4.13.18) (CNDP dipeptidase 2) (Glutamate carboxypeptidase-like protein 1) (Peptidase A) | 475 | CNDP2 | 3 | 1 | 5 | 11 | 2 | 18.9 |
| 745 | -0.27 | 0.00 | 0.00 | Q9NY33 | Dipeptidyl peptidase 3 (EC 3.4.14.4) (Dipeptidyl aminopeptidase III) (Dipeptidyl arylamidase III) (Dipeptidyl peptidase III) (DPP III) (Enkephalinase B) | 737 | DPP3 | 1 | 0 | 0 | 1.4 | 0 | 0 |
| 746 | -0.27 | -0.73 | 0.09 | A0A0A0MR52 | Eukaryotic translation initiation factor 4 gamma 1 | 1395 | EIF4G1 | 2 | 2 | 1 | 1.5 | 1.5 | 0.8 |
| 747 | -0.27 | 0.00 | -0.34 | B4DIH5 | COP9 signalosome complex subunit 2 (cDNA FLJ52928, highly similar to COP9 signalosome complex subunit 2) | 379 | COPS2 | 1 | 0 | 1 | 2.6 | 0 | 2.6 |
| 748 | -0.27 | 0.33 | 0.21 | P46779 | 60S ribosomal protein L28 | 137 | RPL28 | 2 | 5 | 4 | 13.9 | 29.9 | 22.6 |
| 749 | -0.27 | 0.46 | 0.69 | P78527 | DNA-dependent protein kinase catalytic subunit (DNA-PK catalytic subunit) (DNA-PKcs) (EC 2.7.11.1) (DNPK1) (p460) | 4128 | PRKDC | 19 | 35 | 30 | 6.3 | 9.8 | 9 |
| 750 | -0.27 | -0.34 | 0.25 | P62258 | 14-3-3 protein epsilon (14-3-3E) | 255 | YWHAE | 11 | 9 | 13 | 35.3 | 33.3 | 44.7 |
| 751 | -0.27 | 0.92 | 0.76 | P25705 | ATP synthase subunit alpha, mitochondrial | 553 | ATP5A1 | 9 | 6 | 12 | 18.5 | 14.7 | 28 |
| 752 | -0.26 | 0.32 | 0.15 | A6NNI4 | Tetraspanin | 159 | CD9 | 1 | 1 | 1 | 6.3 | 6.3 | 6.3 |
| 753 | -0.26 | 0.71 | 0.37 | P21796 | Voltage-dependent anion-selective channel protein 1 (VDAC-1) (hVDAC1) (Outer mitochondrial membrane protein porin 1) (Plasmalemmal porin) (Porin 31HL) (Porin 31HM) | 283 | VDAC1 | 6 | 9 | 8 | 25.8 | 33.6 | 34.3 |

| | | | | | | | | | | | | | |
|---|---|---|---|---|---|---|---|---|---|---|---|---|---|
| 754 | -0.26 | 0.00 | 1.05 | P30085 | UMP-CMP kinase (EC 2.7.4.14) (Deoxycytidylate kinase) (CK) (dCMP kinase) (Nucleoside-diphosphate kinase) (EC 2.7.4.6) (Uridine monophosphate/cytidine monophosphate kinase) (UMP/CMP kinase) (UMP/CMPK) | 196 | CMPK1 | 1 | 3 | 2 | 8.7 | 18.9 | 14.8 |
| 755 | -0.26 | 0.21 | -0.23 | P60953 | Cell division control protein 42 homolog (G25K GTP-binding protein) | 191 | CDC42 | 4 | 5 | 2 | 27.7 | 32.5 | 11 |
| 756 | -0.26 | -0.18 | 0.34 | P28074 | Proteasome subunit beta type-5 (EC 3.4.25.1) (Macropain epsilon chain) (Multicatalytic endopeptidase complex epsilon chain) (Proteasome chain 6) (Proteasome epsilon chain) (Proteasome subunit MB1) (Proteasome subunit X) | 263 | PSMB5 | 4 | 3 | 3 | 13.7 | 13.3 | 13.3 |
| 757 | -0.25 | 0.00 | -0.42 | F8VS02 | Alpha-aminoadipic semialdehyde dehydrogenase | 475 | ALDH7A1 | 3 | 0 | 2 | 8.8 | 0 | 5.9 |
| 758 | -0.25 | 1.12 | 0.49 | P27105 | Erythrocyte band 7 integral membrane protein (Protein 7.2b) (Stomatin) | 288 | STOM | 2 | 3 | 2 | 9 | 16.7 | 9 |
| 759 | -0.25 | 0.00 | 0.00 | Q9NX02 | NACHT, LRR and PYD domains-containing protein 2 (Nucleotide-binding site protein 1) (PYRIN domain and NACHT domain-containing protein 1) (PYRIN-containing APAF1-like protein 2) | 1062 | NLRP2 | 1 | 0 | 0 | 1 | 0 | 0 |

| | | | | | | | | | | | | | |
|---|---|---|---|---|---|---|---|---|---|---|---|---|---|
| 760 | -0.25 | 0.99 | 0.00 | Q99643 | Succinate dehydrogenase cytochrome b560 subunit, mitochondrial (Integral membrane protein CII-3) (QPs-1) (QPs1) (Succinate dehydrogenase complex subunit C) (Succinate-ubiquinone oxidoreductase cytochrome B large subunit) (CYBL) | 169 | SDHC | 1 | 1 | 0 | 11.2 | 11.2 | 0 |
| 761 | -0.25 | -0.59 | -0.23 | P18206 | Vinculin (Metavinculin) (MV) | 1134 | VCL | 9 | 11 | 12 | 11.7 | 12.8 | 13.9 |
| 762 | -0.25 | 0.04 | 0.44 | E9PLK3 | Puromycin-sensitive aminopeptidase | 915 | NPEPPS | 6 | 4 | 7 | 7.8 | 4.7 | 8.9 |
| 763 | -0.25 | 1.08 | 0.61 | Q14697 | Neutral alpha-glucosidase AB (EC 3.2.1.84) (Alpha-glucosidase 2) (Glucosidase II subunit alpha) | 944 | GANAB | 4 | 9 | 11 | 5.7 | 12.3 | 12.4 |
| 764 | -0.24 | -0.48 | 0.60 | P54652 | Heat shock-related 70 kDa protein 2 (Heat shock 70 kDa protein 2) | 639 | HSPA2 | 5 | 6 | 5 | 8.8 | 10.3 | 9.5 |
| 765 | -0.24 | 0.50 | -0.09 | P35579 | Myosin-9 (Cellular myosin heavy chain, type A) (Myosin heavy chain 9) (Myosin heavy chain, non-muscle IIa) (Non-muscle myosin heavy chain A) (NMMHC-A) (Non-muscle myosin heavy chain IIa) (NMMHC II-a) (NMMHC-IIA) | 1960 | MYH9 | 15 | 10 | 16 | 9.3 | 6.9 | 9.9 |
| 766 | -0.24 | -0.68 | -0.10 | P08758 | Annexin A5 (Anchorin CII) (Annexin V) (Annexin-5) (Calphobindin I) (CBP-I) (Endonexin II) (Lipocortin V) (Placental anticoagulant protein 4) (PP4) (Placental anticoagulant protein I) (PAP-I) (Thromboplastin inhibitor) (Vascular anticoagulant-alpha) (VAC-alpha) | 320 | ANXA5 | 7 | 7 | 5 | 24.1 | 25 | 15.6 |
| 767 | -0.24 | 0.00 | 1.03 | Q12874 | Splicing factor 3A subunit 3 (SF3a60) (Spliceosome-associated protein 61) (SAP 61) | 501 | SF3A3 | 3 | 0 | 2 | 6.2 | 0 | 4.4 |

| | | | | | | | | | | | | | |
|---|---|---|---|---|---|---|---|---|---|---|---|---|---|
| 768 | -0.24 | -0.57 | 0.53 | P15121 | Aldose reductase (AR) (EC 1.1.1.21) (Aldehyde reductase) (Aldo-keto reductase family 1 member B1) | 316 | AKR1B1 | 10 | 12 | 10 | 38.3 | 49.7 | 47.8 |
| 769 | -0.24 | 1.22 | 0.98 | Q12797 | Aspartyl/asparaginyl beta-hydroxylase (EC 1.14.11.16) (Aspartate beta-hydroxylase) (ASP beta-hydroxylase) (Peptide-aspartate beta-dioxygenase) | 758 | ASPH | 3 | 5 | 4 | 4.7 | 10.2 | 11.9 |
| 770 | -0.24 | 0.30 | 0.55 | P63092 | Guanine nucleotide-binding protein G(s) subunit alpha isoforms short (Adenylate cyclase-stimulating G alpha protein) | 394 | GNAS | 3 | 2 | 2 | 7.9 | 5.3 | 5.3 |
| 771 | -0.24 | 0.00 | 0.00 | P61086 | Ubiquitin-conjugating enzyme E2 K (EC 6.3.2.19) (Huntingtin-interacting protein 2) (HIP-2) (Ubiquitin carrier protein) (Ubiquitin-conjugating enzyme E2-25 kDa) (Ubiquitin-conjugating enzyme E2(25K)) (Ubiquitin-conjugating enzyme E2-25K) (Ubiquitin-protein ligase) | 200 | UBE2K | 3 | 0 | 0 | 26.5 | 0 | 0 |
| 772 | -0.24 | 0.01 | 0.20 | P31946 | 14-3-3 protein beta/alpha (Protein 1054) (Protein kinase C inhibitor protein 1) (KCIP-1) [Cleaved into: 14-3-3 protein beta/alpha, N-terminally processed] | 246 | YWHAB | 6 | 8 | 8 | 30.7 | 43.9 | 38.9 |
| 773 | -0.24 | 0.00 | 0.00 | Q96K76 | Ubiquitin carboxyl-terminal hydrolase 47 (EC 3.4.19.12) (Deubiquitinating enzyme 47) (Ubiquitin thioesterase 47) (Ubiquitin-specific-processing protease 47) | 1375 | USP47 | 1 | 0 | 0 | 0.7 | 0 | 0 |

| | | | | | | | | | | | | | |
|---|---|---|---|---|---|---|---|---|---|---|---|---|---|
| 774 | -0.24 | -0.25 | 0.00 | P30043 | Flavin reductase (NADPH) (FR) (EC 1.5.1.30) (Biliverdin reductase B) (BVR-B) (EC 1.3.1.24) (Biliverdin-IX beta-reductase) (Green heme-binding protein) (GHBP) (NADPH-dependent diaphorase) (NADPH-flavin reductase) (FLR) | 206 | BLVRB | 2 | 3 | 0 | 11.7 | 22.8 | 0 |
| 775 | -0.24 | -0.52 | 1.17 | Q8J015 | 60S ribosomal protein L13a (Ribosomal protein L13a) (Ribosomal protein L13a, isoform CRA_b) | 142 | RPL13a | 3 | 2 | 4 | 14.8 | 7 | 20.4 |
| 776 | -0.24 | 0.00 | 0.63 | Q15006 | ER membrane protein complex subunit 2 (Tetratricopeptide repeat protein 35) (TPR repeat protein 35) | 297 | EMC2 | 1 | 0 | 1 | 6.4 | 0 | 6.4 |
| 777 | -0.24 | 1.32 | 0.00 | E9PH64 | NADH dehydrogenase [ubiquinone] 1 beta subcomplex subunit 9 | 168 | NDUFB9 | 1 | 3 | 0 | 8.3 | 22 | 0 |
| 778 | -0.23 | 0.00 | 1.40 | Q92820 | Gamma-glutamyl hydrolase (EC 3.4.19.9) (Conjugase) (GH) (Gamma-Glu-X carboxypeptidase) | 318 | GGH | 1 | 0 | 2 | 4.1 | 0 | 7.2 |
| 779 | -0.23 | -0.22 | 0.20 | O00231 | 26S proteasome non-ATPase regulatory subunit 11 (26S proteasome regulatory subunit RPN6) (26S proteasome regulatory subunit S9) (26S proteasome regulatory subunit p44.5) | 422 | PSMD11 | 7 | 5 | 7 | 20.4 | 14 | 20.1 |
| 780 | -0.23 | -0.02 | 0.89 | E5RI99 | 60S ribosomal protein L30 (Fragment) | 114 | RPL30 | 4 | 3 | 4 | 32.5 | 35.1 | 32.5 |
| 781 | -0.23 | 0.84 | 0.64 | P52701 | DNA mismatch repair protein Msh6 (hMSH6) (G/T mismatch-binding protein) (GTBP) (GTMBP) (MutS-alpha 160 kDa subunit) (p160) | 1360 | MSH6 | 2 | 3 | 2 | 2.4 | 3.4 | 2.4 |
| 782 | -0.23 | -0.04 | -0.02 | M0R0F0 | 40S ribosomal protein S5 (Fragment) | 200 | RPS5 | 1 | 2 | 1 | 7.5 | 12 | 7.5 |
| 783 | -0.23 | 0.00 | 0.00 | E9PDQ8 | Succinyl-CoA ligase [GDP-forming] subunit beta, mitochondrial | 379 | SUCLG2 | 1 | 0 | 0 | 4 | 0 | 0 |

| | | | | | | | | | | | | | |
|---|---|---|---|---|---|---|---|---|---|---|---|---|---|
| 784 | -0.23 | 0.00 | 0.11 | P11766 | Alcohol dehydrogenase class-3 (EC 1.1.1.1) (Alcohol dehydrogenase 5) (Alcohol dehydrogenase class chi chain) (Alcohol dehydrogenase class-III) (Glutathione-dependent formaldehyde dehydrogenase) (FALDH) (FDH) (GSH-FDH) (EC 1.1.1.-) (S-(hydroxymethyl)glutathione dehydrogenase) (EC 1.1.1.284) | 374 | ADH5 | 4 | 0 | 3 | 11.5 | 0 | 9.9 |
| 785 | -0.23 | 0.33 | 0.66 | Q9Y230 | RuvB-like 2 (EC 3.6.4.12) (48 kDa TATA box-binding protein-interacting protein) (48 kDa TBP-interacting protein) (51 kDa erythrocyte cytosolic protein) (ECP-51) (INO80 complex subunit J) (Repressing pontin 52) (Reptin 52) (TIP49b) (TIP60-associated protein 54-beta) (TAP54-beta) | 463 | RUVBL2 | 6 | 4 | 8 | 15.8 | 10.2 | 22.9 |
| 786 | -0.23 | -0.05 | -0.60 | K7EL96 | Perilipin-3 (Fragment) | 172 | PLIN3 | 2 | 2 | 3 | 15.7 | 18 | 24.4 |
| 787 | -0.22 | -0.78 | 0.62 | Q16719 | Kynureninase (EC 3.7.1.3) (L-kynurenine hydrolase) | 465 | KYNU | 7 | 8 | 12 | 20.6 | 21.5 | 37 |
| 788 | -0.22 | -0.09 | -0.24 | P31949 | Protein S100-A11 (Calgizzarin) (Metastatic lymph node gene 70 protein) (MLN 70) (Protein S100-C) (S100 calcium-binding protein A11) [Cleaved into: Protein S100-A11, N-terminally processed] | 105 | S100A11 | 1 | 3 | 3 | 15.2 | 34.3 | 34.3 |
| 789 | -0.22 | -0.33 | 0.70 | P48163 | NADP-dependent malic enzyme (NADP-ME) (EC 1.1.1.40) (Malic enzyme 1) | 572 | ME1 | 1 | 1 | 1 | 3.6 | 4.2 | 3.6 |
| 790 | -0.22 | -0.72 | -0.26 | Q14019 | Coactosin-like protein | 142 | COTL1 | 4 | 3 | 4 | 15.5 | 19.7 | 15.5 |

| | | | | | | | | | | | | | |
|---|---|---|---|---|---|---|---|---|---|---|---|---|---|
| 791 | -0.22 | 0.00 | 0.62 | Q92905 | COP9 signalosome complex subunit 5 (SGN5) (Signalosome subunit 5) (EC 3.4.-.-) (Jun activation domain-binding protein 1) | 334 | COPS5 | 1 | 0 | 1 | 4.5 | 0 | 4.5 |
| 792 | -0.21 | -0.15 | 0.19 | G3V295 | Proteasome subunit alpha type (EC 3.4.25.1) | 203 | PSMA6 | 2 | 7 | 4 | 10.3 | 36.9 | 21.7 |
| 793 | -0.21 | 0.61 | 0.47 | E9PEX6 | Dihydrolipoyl dehydrogenase (EC 1.8.1.4) | 486 | DLD | 3 | 2 | 5 | 8.8 | 5.1 | 11.7 |
| 794 | -0.21 | 0.39 | 0.00 | P19525 | Interferon-induced, double-stranded RNA-activated protein kinase (EC 2.7.11.1) (Eukaryotic translation initiation factor 2-alpha kinase 2) (eIF-2A protein kinase 2) (Interferon-inducible RNA-dependent protein kinase) (P1/eIF-2A protein kinase) (Protein kinase RNA-activated) (PKR) (Protein kinase R) (Tyrosine-protein kinase EIF2AK2) (EC 2.7.10.2) (p68 kinase) | 551 | EIF2AK2 | 1 | 1 | 0 | 2.4 | 2.4 | 0 |
| 795 | -0.21 | 0.36 | 0.64 | P28070 | Proteasome subunit beta type-4 (EC 3.4.25.1) (26 kDa prosomal protein) (HsBPROS26) (PROS-26) (Macropain beta chain) (Multicatalytic endopeptidase complex beta chain) (Proteasome beta chain) (Proteasome chain 3) (HsN3) | 264 | PSMB4 | 3 | 4 | 2 | 20.8 | 25.4 | 15.9 |
| 796 | -0.21 | -0.14 | 1.26 | P09012 | U1 small nuclear ribonucleoprotein A (U1 snRNP A) (U1-A) (U1A) | 282 | SNRPA | 1 | 4 | 2 | 2.8 | 15.6 | 9.6 |
| 797 | -0.21 | 0.27 | 1.29 | P61586 | Transforming protein RhoA (Rho cDNA clone 12) (h12) | 193 | RHOA | 5 | 5 | 5 | 35.2 | 24.4 | 24.4 |

| | | | | | | | | | | | | | |
|---|---|---|---|---|---|---|---|---|---|---|---|---|---|
| 798 | -0.21 | 0.23 | 0.28 | P25788 | Proteasome subunit alpha type-3 (EC 3.4.25.1) (Macropain subunit C8) (Multicatalytic endopeptidase complex subunit C8) (Proteasome component C8) | 255 | PSMA3 | 3 | 2 | 3 | 8.5 | 8.1 | 8.5 |
| 799 | -0.21 | 1.04 | 0.46 | P12236 | ADP/ATP translocase 3 (ADP,ATP carrier protein 3) (ADP,ATP carrier protein, isoform T2) (ANT 2) (Adenine nucleotide translocator 3) (ANT 3) (Solute carrier family 25 member 6) [Cleaved into: ADP/ATP translocase 3, N-terminally processed] | 298 | SLC25A6 | 8 | 8 | 8 | 27.2 | 29.5 | 26.2 |
| 800 | -0.20 | 0.00 | 0.17 | O95202 | LETM1 and EF-hand domain-containing protein 1, mitochondrial (Leucine zipper-EF-hand-containing transmembrane protein 1) | 739 | LETM1 | 2 | 0 | 2 | 2.7 | 0 | 2.7 |
| 801 | -0.20 | -0.43 | 0.13 | P23528 | Cofilin-1 (18 kDa phosphoprotein) (p18) (Cofilin, non-muscle isoform) | 166 | CFL1 | 12 | 10 | 13 | 57.2 | 63.9 | 57.2 |
| 802 | -0.20 | 0.00 | -0.02 | P21283 | V-type proton ATPase subunit C 1 (V-ATPase subunit C 1) (Vacuolar proton pump subunit C 1) | 382 | ATP6V1C1 | 1 | 0 | 1 | 4.2 | 0 | 4.2 |
| 803 | -0.20 | -0.29 | 0.86 | P33176 | Kinesin-1 heavy chain (Conventional kinesin heavy chain) (Ubiquitous kinesin heavy chain) (UKHC) | 963 | KIF5B | 2 | 2 | 1 | 2.4 | 2.5 | 1 |
| 804 | -0.20 | -0.31 | 0.11 | R4GMY8 | Transcription elongation factor B polypeptide 1 | 65 | TCEB1 | 1 | 2 | 1 | 18.5 | 33.8 | 18.5 |
| 805 | -0.20 | -0.29 | 0.24 | H7C3I1 | Hsc70-interacting protein (Fragment) | 146 | ST13 | 1 | 2 | 3 | 6.8 | 18.5 | 25.3 |
| 806 | -0.20 | 0.00 | 0.24 | O14828 | Secretory carrier-associated membrane protein 3 (Secretory carrier membrane protein 3) | 347 | SCAMP3 | 1 | 0 | 1 | 6.2 | 0 | 6.2 |
| 807 | -0.20 | 0.44 | 0.15 | P20340 | Ras-related protein Rab-6A (Rab-6) | 208 | RAB6A | 4 | 3 | 4 | 23.6 | 17.8 | 22.6 |

| | | | | | | | | | | | | | |
|---|---|---|---|---|---|---|---|---|---|---|---|---|---|
| 808 | -0.20 | -0.41 | 0.10 | Q8WUM4 | Programmed cell death 6-interacting protein (PDCD6-interacting protein) (ALG-2-interacting protein 1) (ALG-2-interacting protein X) (Hp95) | 868 | PDCD6IP | 5 | 3 | 6 | 5.8 | 5.2 | 6.9 |
| 809 | -0.20 | 0.00 | -0.12 | P35237 | Serpin B6 (Cytoplasmic antiproteinase) (CAP) (Peptidase inhibitor 6) (PI-6) (Placental thrombin inhibitor) | 376 | SERPINB6 | 3 | 0 | 2 | 12.5 | 0 | 8.5 |
| 810 | -0.19 | 0.74 | 0.23 | P07814 | Bifunctional glutamate/proline--tRNA ligase (Bifunctional aminoacyl-tRNA synthetase) (Cell proliferation-inducing gene 32 protein) (Glutamatyl-prolyl-tRNA synthetase) [Includes: Glutamate--tRNA ligase (EC 6.1.1.17) (Glutamyl-tRNA synthetase) (GluRS); Proline--tRNA ligase (EC 6.1.1.15) (Prolyl-tRNA synthetase)] | 1512 | EPRS | 5 | 4 | 4 | 4.4 | 3.6 | 3.3 |
| 811 | -0.19 | 0.33 | 0.37 | Q9P2J5 | Leucine--tRNA ligase, cytoplasmic (EC 6.1.1.4) (Leucyl-tRNA synthetase) (LeuRS) | 1176 | LARS | 5 | 9 | 6 | 5.1 | 9.6 | 6 |
| 812 | -0.19 | 0.00 | -0.31 | Q7Z406 | Myosin-14 (Myosin heavy chain 14) (Myosin heavy chain, non-muscle IIc) (Non-muscle myosin heavy chain IIc) (NMHC II-C) | 1995 | MYH14 | 2 | 2 | 2 | 1.1 | 1.3 | 1.1 |
| 813 | -0.19 | -0.13 | 1.09 | Q13347 | Eukaryotic translation initiation factor 3 subunit I (eIF3i) (Eukaryotic translation initiation factor 3 subunit 2) (TGF-beta receptor-interacting protein 1) (TRIP-1) (eIF-3-beta) (eIF3 p36) | 325 | EIF3I | 1 | 1 | 4 | 3.4 | 2.8 | 21.5 |
| 814 | -0.19 | -0.39 | 0.16 | Q02809 | Procollagen-lysine,2-oxoglutarate 5-dioxygenase 1 (EC 1.14.11.4) (Lysyl hydroxylase 1) (LH1) | 727 | PLOD1 | 1 | 1 | 1 | 2.1 | 2.1 | 2.1 |

| | | | | | | | | | | | | | |
|---|---|---|---|---|---|---|---|---|---|---|---|---|---|
| 815 | -0.19 | 0.00 | 1.22 | B4DT28 | Heterogeneous nuclear ribonucleoprotein R (Heterogeneous nuclear ribonucleoprotein R, isoform CRA_a) (cDNA FLJ54544, highly similar to Heterogeneous nuclear ribonucleoprotein R) | 494 | HNRNPR | 2 | 1 | 3 | 4.9 | 2.6 | 7.1 |
| 816 | -0.18 | -0.50 | -0.48 | Q5VYK3 | Proteasome-associated protein ECM29 homolog (Ecm29) | 1845 | ECM29 | 2 | 1 | 1 | 1.4 | 0.7 | 0.7 |
| 817 | -0.18 | 0.00 | 0.11 | Q58FF6 | Putative heat shock protein HSP 90-beta 4 | 505 | HSP90AB4P | 5 | 1 | 5 | 7.7 | 2.4 | 7.7 |
| 818 | -0.18 | -0.21 | 0.33 | Q5TA02 | Glutathione S-transferase omega-1 (Fragment) | 200 | GSTO1 | 2 | 4 | 4 | 11 | 20 | 20.5 |
| 819 | -0.18 | 0.27 | 0.31 | P39019 | 40S ribosomal protein S19 | 145 | RPS19 | 3 | 6 | 3 | 18.6 | 34.5 | 18.6 |
| 820 | -0.18 | 0.00 | 0.04 | C9J0K5 | Cytoplasmic protein NCK1 (Fragment) | 75 | NCK1 | 1 | 0 | 1 | 12 | 0 | 12 |
| 821 | -0.17 | -0.72 | -0.01 | Q15819 | Ubiquitin-conjugating enzyme E2 variant 2 (DDVit 1) (Enterocyte differentiation-associated factor 1) (EDAF-1) (Enterocyte differentiation-promoting factor 1) (EDPF-1) (MMS2 homolog) (Vitamin D3-inducible protein) | 145 | UBE2V2 | 2 | 2 | 1 | 11.7 | 11.7 | 6.9 |
| 822 | -0.17 | -0.90 | 0.72 | O43143 | Pre-mRNA-splicing factor ATP-dependent RNA helicase DHX15 (EC 3.6.4.13) (ATP-dependent RNA helicase #46) (DEAH box protein 15) | 795 | DHX15 | 2 | 1 | 2 | 2.9 | 1.5 | 2.9 |
| 823 | -0.17 | -0.10 | 0.32 | Q9Y5S9 | RNA-binding protein 8A (Binder of OVCA1-1) (BOV-1) (RNA-binding motif protein 8A) (RNA-binding protein Y14) (Ribonucleoprotein RBM8A) | 174 | RBM8A | 1 | 2 | 1 | 6.4 | 11 | 6.4 |

| | | | | | | | | | | | | | |
|---|---|---|---|---|---|---|---|---|---|---|---|---|---|
| 824 | -0.17 | -0.07 | 0.33 | Q13838 | Spliceosome RNA helicase DDX39B (EC 3.6.4.13) (56 kDa U2AF65-associated protein) (ATP-dependent RNA helicase p47) (DEAD box protein UAP56) (HLA-B-associated transcript 1 protein) | 428 | DDX39B | 3 | 2 | 5 | 8.2 | 5.1 | 16.6 |
| 825 | -0.17 | 0.89 | 0.38 | E9PLD0 | Ras-related protein Rab-1B | 169 | RAB1B | 3 | 5 | 5 | 20.7 | 30.8 | 37.9 |
| 826 | -0.17 | 0.00 | -0.06 | C9JIJ1 | RAC-beta serine/threonine-protein kinase (Fragment) | 119 | AKT2 | 1 | 0 | 1 | 8.4 | 0 | 8.4 |
| 827 | -0.17 | 2.06 | 0.37 | O95573 | Long-chain-fatty-acid--CoA ligase 3 (EC 6.2.1.3) (Long-chain acyl-CoA synthetase 3) (LACS 3) | 720 | ACSL3 | 2 | 2 | 2 | 3.5 | 4.6 | 3.5 |
| 828 | -0.17 | 0.01 | 0.07 | A0A0A6YYA0 | Protein TMED7-TICAM2 | 188 | TMED7-TICAM2 | 1 | 1 | 2 | 10.1 | 6.4 | 16.5 |
| 829 | -0.16 | -0.63 | -1.88 | B1AJY5 | 26S proteasome non-ATPase regulatory subunit 10 | 185 | PSMD10 | 2 | 1 | 3 | 10.8 | 4.3 | 21.1 |
| 830 | -0.16 | 0.26 | 0.86 | Q01081 | Splicing factor U2AF 35 kDa subunit (U2 auxiliary factor 35 kDa subunit) (U2 small nuclear RNA auxiliary factor 1) (U2 snRNP auxiliary factor small subunit) | 240 | U2AF1 | 2 | 2 | 2 | 5.8 | 10.4 | 5.8 |
| 831 | -0.16 | 0.42 | 0.23 | Q9Y6M7 | Sodium bicarbonate cotransporter 3 (Electroneutral Na/HCO(3) cotransporter) (Sodium bicarbonate cotransporter 2) (Sodium bicarbonate cotransporter 2b) (Bicarbonate transporter) (Solute carrier family 4 member 7) | 1214 | SLC4A7 | 3 | 2 | 1 | 4.8 | 3.5 | 2.1 |

| | | | | | | | | | | | | | |
|---|---|---|---|---|---|---|---|---|---|---|---|---|---|
| 832 | -0.16 | 0.74 | 0.66 | P16615 | Sarcoplasmic/endoplasmic reticulum calcium ATPase 2 (SERCA2) (SR Ca(2+)-ATPase 2) (EC 3.6.3.8) (Calcium pump 2) (Calcium-transporting ATPase sarcoplasmic reticulum type, slow twitch skeletal muscle isoform) (Endoplasmic reticulum class 1/2 Ca(2+) ATPase) | 1042 | ATP2A2 | 8 | 10 | 12 | 10.9 | 13.2 | 17.7 |
| 833 | -0.16 | -0.29 | -0.46 | P61956 | Small ubiquitin-related modifier 2 (SUMO-2) (HSMT3) (SMT3 homolog 2) (SUMO-3) (Sentrin-2) (Ubiquitin-like protein SMT3B) (Smt3B) | 95 | SUMO2 | 1 | 2 | 2 | 16.9 | 31 | 31 |
| 834 | -0.15 | -0.04 | 0.43 | Q9UBT2 | SUMO-activating enzyme subunit 2 (EC 6.3.2.-) (Anthracycline-associated resistance ARX) (Ubiquitin-like 1-activating enzyme E1B) (Ubiquitin-like modifier-activating enzyme 2) | 640 | UBA2 | 3 | 1 | 1 | 7 | 2.3 | 2.3 |
| 835 | -0.15 | 0.31 | 0.61 | Q15181 | Inorganic pyrophosphatase (EC 3.6.1.1) (Pyrophosphate phospho-hydrolase) (PPase) | 289 | PPA1 | 1 | 1 | 3 | 3.5 | 3.1 | 11.8 |
| 836 | -0.15 | 2.08 | 0.44 | H0YDT8 | ER membrane protein complex subunit 7 (Fragment) | 192 | EMC7 | 1 | 1 | 2 | 6.8 | 8.3 | 10.9 |
| 837 | -0.15 | 0.00 | 0.00 | F8WCQ2 | Phosphatidylinositide phosphatase SAC1 | 71 | SACM1L | 1 | 0 | 0 | 12.7 | 0 | 0 |
| 838 | -0.15 | -0.41 | 0.29 | Q86Y56 | Dynein assembly factor 5, axonemal (HEAT repeat-containing protein 2) | 855 | DNAAF5 | 1 | 1 | 1 | 4.7 | 4.7 | 4.7 |
| 839 | -0.15 | 0.46 | 0.01 | P30049 | ATP synthase subunit delta, mitochondrial (F-ATPase delta subunit) | 168 | ATP5D | 1 | 1 | 1 | 5.4 | 5.4 | 5.4 |
| 840 | -0.15 | 0.00 | 0.00 | Q9UBK9 | Protein UXT (Androgen receptor trapped clone 27 protein) (ART-27) (Ubiquitously expressed transcript protein) | 157 | UXT | 1 | 0 | 0 | 8.3 | 0 | 0 |

| | | | | | | | | | | | | | |
|---|---|---|---|---|---|---|---|---|---|---|---|---|---|
| 841 | -0.15 | 2.44 | 0.00 | Q96HC4 | PDZ and LIM domain protein 5 (Enigma homolog) (Enigma-like PDZ and LIM domains protein) | 596 | PDLIM5 | 1 | 1 | 0 | 1.9 | 2.9 | 0 |
| 842 | -0.15 | 0.00 | -0.08 | Q92621 | Nuclear pore complex protein Nup205 (205 kDa nucleoporin) (Nucleoporin Nup205) | 2012 | NUP205 | 1 | 0 | 2 | 0.6 | 0 | 1.2 |
| 843 | -0.14 | 0.00 | 0.33 | Q16878 | Cysteine dioxygenase type 1 (EC 1.13.11.20) (Cysteine dioxygenase type I) (CDO) (CDO-I) | 200 | CDO1 | 1 | 0 | 1 | 4 | 0 | 4 |
| 844 | -0.14 | 0.00 | 2.49 | O94826 | Mitochondrial import receptor subunit TOM70 (Mitochondrial precursor proteins import receptor) (Translocase of outer membrane 70 kDa subunit) | 608 | TOMM70A | 1 | 0 | 2 | 1.5 | 0 | 7.6 |
| 845 | -0.14 | -0.14 | 0.13 | P61981 | 14-3-3 protein gamma (Protein kinase C inhibitor protein 1) (KCIP-1) [Cleaved into: 14-3-3 protein gamma, N-terminally processed] | 247 | YWHAG | 6 | 8 | 7 | 33.2 | 47 | 37.2 |
| 846 | -0.14 | 0.27 | 0.61 | O43707 | Alpha-actinin-4 (Non-muscle alpha-actinin 4) | 911 | ACTN4 | 18 | 28 | 22 | 24.3 | 38.5 | 30.7 |
| 847 | -0.14 | -0.10 | 0.19 | P62333 | 26S protease regulatory subunit 10B (26S proteasome AAA-ATPase subunit RPT4) (Proteasome 26S subunit ATPase 6) (Proteasome subunit p42) | 389 | PSMC6 | 4 | 3 | 5 | 11.8 | 9.5 | 14.9 |
| 848 | -0.14 | 1.04 | -0.24 | P00167 | Cytochrome b5 (Microsomal cytochrome b5 type A) (MCB5) | 134 | CYB5A | 1 | 3 | 1 | 9.2 | 36.7 | 9.2 |
| 849 | -0.14 | 1.07 | 0.64 | Q8N5K1 | CDGSH iron-sulfur domain-containing protein 2 (Endoplasmic reticulum intermembrane small protein) (MitoNEET-related 1 protein) (Miner1) (Nutrient-deprivation autophagy factor-1) (NAF-1) | 135 | CISD2 | 3 | 7 | 3 | 21.5 | 47.4 | 21.5 |
| 850 | -0.14 | 0.37 | 0.31 | E9PF10 | Nuclear pore complex protein Nup155 | 1327 | NUP155 | 1 | 1 | 1 | 0.8 | 0.8 | 0.8 |

| | | | | | | | | | | | | | |
|---|---|---|---|---|---|---|---|---|---|---|---|---|---|
| 851 | -0.14 | -0.59 | 0.61 | Q08752 | Peptidyl-prolyl cis-trans isomerase D (PPIase D) (EC 5.2.1.8) (40 kDa peptidyl-prolyl cis-trans isomerase) (Cyclophilin-40) (CYP-40) (Cyclophilin-related protein) (Rotamase D) | 370 | PPID | 2 | 1 | 1 | 4.6 | 2.4 | 2.2 |
| 852 | -0.14 | 0.00 | 0.26 | H0Y4Q3 | Ran GTPase-activating protein 1 (Fragment) | 253 | RANGAP1 | 2 | 0 | 2 | 11.1 | 0 | 11.1 |
| 853 | -0.14 | -0.05 | -0.12 | A8MU58 | Aminoacyl tRNA synthase complex-interacting multifunctional protein 2 | 242 | AIMP2 | 2 | 1 | 2 | 8.7 | 3.7 | 8.7 |
| 854 | -0.13 | 1.31 | -0.18 | P47985 | Cytochrome b-c1 complex subunit Rieske, mitochondrial (EC 1.10.2.2) (Complex III subunit 5) (Cytochrome b-c1 complex subunit 5) (Rieske iron-sulfur protein) (RISP) (Ubiquinol-cytochrome c reductase iron-sulfur subunit) [Cleaved into: Cytochrome b-c1 complex subunit 11 (Complex III subunit IX) (Ubiquinol-cytochrome c reductase 8 kDa protein)] | 274 | UQCRFS1 | 2 | 3 | 1 | 10.9 | 15.3 | 7.7 |
| 855 | -0.13 | 1.40 | 0.23 | Q9P0I2 | ER membrane protein complex subunit 3 (Transmembrane protein 111) | 261 | EMC3 | 1 | 1 | 1 | 3.4 | 5 | 3.4 |
| 856 | -0.13 | 0.11 | 0.42 | P53621 | Coatomer subunit alpha (Alpha-coat protein) (Alpha-COP) (HEP-COP) (HEPCOP) [Cleaved into: Xenin (Xenopsin-related peptide); Proxenin] | 1224 | COPA | 4 | 3 | 4 | 4.3 | 3.2 | 4 |
| 857 | -0.13 | 0.00 | 0.07 | B4E1C5 | Histidine--tRNA ligase, cytoplasmic (cDNA FLJ58562, highly similar to Histidyl-tRNA synthetase (EC 6.1.1.21)) | 395 | HARS | 1 | 0 | 2 | 3.3 | 0 | 6.3 |
| 858 | -0.13 | 0.00 | -0.09 | P63096 | Guanine nucleotide-binding protein G(i) subunit alpha-1 (Adenylate cyclase-inhibiting G alpha protein) | 354 | GNAI1 | 2 | 1 | 2 | 6.8 | 3.1 | 6.8 |

| | | | | | | | | | | | | | |
|---|---|---|---|---|---|---|---|---|---|---|---|---|---|
| 859 | -0.13 | 0.04 | 0.53 | Q14914 | Prostaglandin reductase 1 (PRG-1) (EC 1.3.1.-) (15-oxoprostaglandin 13-reductase) (EC 1.3.1.48) (NADP-dependent leukotriene B4 12-hydroxydehydrogenase) (EC 1.3.1.74) | 329 | PTGR1 | 5 | 5 | 5 | 22.2 | 26.1 | 21.3 |
| 860 | -0.13 | 0.00 | 0.15 | Q5MIZ7 | Serine/threonine-protein phosphatase 4 regulatory subunit 3B (SMEK homolog 2) | 849 | SMEK2 | 1 | 0 | 1 | 1.6 | 0 | 1.6 |
| 861 | -0.12 | 0.46 | -0.44 | Q9NX63 | MICOS complex subunit MIC19 (Coiled-coil-helix-coiled-coil-helix domain-containing protein 3) | 227 | CHCHD3 | 2 | 2 | 3 | 11.5 | 11.5 | 11.9 |
| 862 | -0.12 | 0.00 | 0.67 | H0YA83 | Beta-hexosaminidase subunit beta (Fragment) | 170 | HEXB | 1 | 0 | 1 | 5.9 | 0 | 5.9 |
| 863 | -0.12 | 1.26 | 0.70 | Q9NYU2 | UDP-glucose:glycoprotein glucosyltransferase 1 (UGT1) (hUGT1) (EC 2.4.1.-) (UDP--Glc:glycoprotein glucosyltransferase) (UDP-glucose ceramide glucosyltransferase-like 1) | 1555 | UGGT1 | 1 | 3 | 3 | 0.8 | 3.4 | 2.9 |
| 864 | -0.12 | 0.36 | 0.93 | Q9UKD2 | mRNA turnover protein 4 homolog (Ribosome assembly factor MRTO4) | 239 | MRTO4 | 1 | 1 | 1 | 5.4 | 5.4 | 5.4 |
| 865 | -0.12 | 0.27 | 0.11 | P61019 | Ras-related protein Rab-2A | 212 | RAB2A | 4 | 5 | 5 | 20.8 | 25.5 | 28.3 |
| 866 | -0.12 | 0.00 | -0.01 | Q8N0U8 | Vitamin K epoxide reductase complex subunit 1-like protein 1 (VKORC1-like protein 1) (EC 1.17.4.4) | 176 | VKORC1L1 | 1 | 0 | 1 | 6.2 | 0 | 6.2 |
| 867 | -0.12 | -0.49 | 0.38 | O00232 | 26S proteasome non-ATPase regulatory subunit 12 (26S proteasome regulatory subunit RPN5) (26S proteasome regulatory subunit p55) | 456 | PSMD12 | 2 | 1 | 2 | 5 | 2.2 | 5 |
| 868 | -0.12 | 0.35 | 0.22 | P51149 | Ras-related protein Rab-7a | 207 | RAB7A | 5 | 8 | 6 | 29 | 47.8 | 34.8 |

| | | | | | | | | | | | | | |
|---|---|---|---|---|---|---|---|---|---|---|---|---|---|
| 869 | -0.12 | 0.59 | 1.31 | K7EJE8 | Lon protease homolog, mitochondrial (EC 3.4.21.-) (Lon protease-like protein) (Mitochondrial ATP-dependent protease Lon) (Serine protease 15) | 829 | LONP1 | 1 | 3 | 3 | 1.3 | 4.2 | 4.5 |
| 870 | -0.11 | 0.94 | 0.44 | P02786 | Transferrin receptor protein 1 (TR) (TfR) (TfR1) (Trfr) (T9) (p90) (CD antigen CD71) [Cleaved into: Transferrin receptor protein 1, serum form (sTfR)] | 760 | TFRC | 5 | 5 | 9 | 7.6 | 8.4 | 14.5 |
| 871 | -0.11 | 0.00 | 0.22 | Q9H0A8 | COMM domain-containing protein 4 | 199 | COMMD4 | 1 | 0 | 1 | 13.3 | 0 | 13.3 |
| 872 | -0.11 | -0.44 | -0.15 | P21291 | Cysteine and glycine-rich protein 1 (Cysteine-rich protein 1) (CRP) (CRP1) (Epididymis luminal protein 141) (HEL-141) | 193 | CSRP1 | 6 | 8 | 7 | 50.3 | 55.4 | 59.1 |
| 873 | -0.11 | -0.52 | 0.40 | P04083 | Annexin A1 (Annexin I) (Annexin-1) (Calpactin II) (Calpactin-2) (Chromobindin-9) (Lipocortin I) (Phospholipase A2 inhibitory protein) (p35) | 346 | ANXA1 | 11 | 17 | 12 | 32.7 | 55.5 | 30.3 |
| 874 | -0.11 | 0.00 | 0.00 | K7EP09 | Bifunctional coenzyme A synthase (Fragment) | 90 | COASY | 1 | 0 | 0 | 11.1 | 0 | 0 |
| 875 | -0.11 | 0.49 | 0.77 | C9JQ41 | Coiled-coil domain-containing protein 58 | 130 | CCDC58 | 1 | 1 | 2 | 6.9 | 8.5 | 20 |
| 876 | -0.11 | 0.62 | 0.32 | I3L1P8 | Mitochondrial 2-oxoglutarate/malate carrier protein (Fragment) | 296 | SLC25A11 | 3 | 1 | 1 | 14.5 | 3.7 | 5.4 |
| 877 | -0.11 | 1.23 | 0.23 | F8VVM2 | Phosphate carrier protein, mitochondrial | 324 | SLC25A3 | 2 | 4 | 2 | 5.9 | 17.9 | 5.9 |
| 878 | -0.11 | 0.07 | 0.09 | M0QZS6 | SUMO-activating enzyme subunit 1 | 265 | SAE1 | 3 | 1 | 2 | 12.1 | 4.9 | 7.9 |
| 879 | -0.11 | 0.75 | 1.02 | P49720 | Proteasome subunit beta type-3 (EC 3.4.25.1) (Proteasome chain 13) (Proteasome component C10-II) (Proteasome theta chain) | 205 | PSMB3 | 5 | 4 | 6 | 25.4 | 25.4 | 33.2 |

| | | | | | | | | | | | | | |
|---|---|---|---|---|---|---|---|---|---|---|---|---|---|
| 880 | -0.11 | 0.00 | 0.09 | Q9Y262 | Eukaryotic translation initiation factor 3 subunit L (eIF3l) (Eukaryotic translation initiation factor 3 subunit 6-interacting protein) (Eukaryotic translation initiation factor 3 subunit E-interacting protein) | 564 | EIF3L | 1 | 0 | 1 | 1.6 | 0 | 1.6 |
| 881 | -0.10 | 0.61 | -0.23 | P04632 | Calpain small subunit 1 (CSS1) (Calcium-activated neutral proteinase small subunit) (CANP small subunit) (Calcium-dependent protease small subunit) (CDPS) (Calcium-dependent protease small subunit 1) (Calpain regulatory subunit) | 268 | CAPNS1 | 3 | 3 | 2 | 10.8 | 14.9 | 6 |
| 882 | -0.10 | -0.06 | 0.28 | J3KTJ1 | Myosin regulatory light chain 12A (Fragment) | 114 | MYL12A | 1 | 1 | 1 | 15.8 | 15.8 | 15.8 |
| 883 | -0.10 | -0.33 | 0.10 | P56192 | Methionine--tRNA ligase, cytoplasmic (EC 6.1.1.10) (Methionyl-tRNA synthetase) (MetRS) | 900 | MARS | 3 | 1 | 2 | 4.1 | 1.2 | 2.6 |
| 884 | -0.10 | 1.04 | 0.28 | P00505 | Aspartate aminotransferase, mitochondrial (mAspAT) (EC 2.6.1.1) (EC 2.6.1.7) (Fatty acid-binding protein) (FABP-1) (Glutamate oxaloacetate transaminase 2) (Kynurenine aminotransferase 4) (Kynurenine aminotransferase IV) (Kynurenine--oxoglutarate transaminase 4) (Kynurenine--oxoglutarate transaminase IV) (Plasma membrane-associated fatty acid-binding protein) (FABPpm) (Transaminase A) | 430 | GOT2 | 4 | 7 | 8 | 13 | 16.5 | 21.4 |

| | | | | | | | | | | | | | |
|---|---|---|---|---|---|---|---|---|---|---|---|---|---|
| 885 | -0.10 | 0.67 | 0.35 | P30040 | Endoplasmic reticulum resident protein 29 (ERp29) (Endoplasmic reticulum resident protein 28) (ERp28) (Endoplasmic reticulum resident protein 31) (ERp31) | 261 | ERP29 | 2 | 5 | 3 | 7.7 | 25.3 | 21.8 |
| 886 | -0.09 | 0.07 | 0.22 | P46108 | Adapter molecule crk (Proto-oncogene c-Crk) (p38) | 304 | CRK | 2 | 1 | 2 | 9.5 | 5.6 | 9.5 |
| 887 | -0.09 | 0.82 | 0.76 | P14625 | Endoplasmin (94 kDa glucose-regulated protein) (GRP-94) (Heat shock protein 90 kDa beta member 1) (Tumor rejection antigen 1) (gp96 homolog) | 803 | HSP90B1 | 14 | 14 | 18 | 16.9 | 20.7 | 23.8 |
| 888 | -0.09 | 0.54 | 0.53 | P30044 | Peroxiredoxin-5, mitochondrial (EC 1.11.1.15) (Alu corepressor 1) (Antioxidant enzyme B166) (AOEB166) (Liver tissue 2D-page spot 71B) (PLP) (Peroxiredoxin V) (Prx-V) (Peroxisomal antioxidant enzyme) (TPx type VI) (Thioredoxin peroxidase PMP20) (Thioredoxin reductase) | 214 | PRDX5 | 5 | 7 | 6 | 37 | 62.3 | 48.8 |
| 889 | -0.09 | 0.86 | 0.64 | H7C0V0 | Uncharacterized protein C2orf47, mitochondrial (Fragment) | 225 | C2orf47 | 1 | 1 | 1 | 4.4 | 4.4 | 4.4 |
| 890 | -0.09 | 0.00 | 0.00 | H0YLW7 | SAFB-like transcription modulator | 62 | SLTM | 1 | 0 | 0 | 30.6 | 0 | 0 |
| 891 | -0.09 | 0.14 | 0.09 | Q9H488 | GDP-fucose protein O-fucosyltransferase 1 (EC 2.4.1.221) (Peptide-O-fucosyltransferase 1) (O-FucT-1) | 388 | POFUT1 | 1 | 1 | 1 | 5.2 | 5.2 | 5.2 |
| 892 | -0.09 | 0.65 | -0.40 | O43674 | NADH dehydrogenase [ubiquinone] 1 beta subcomplex subunit 5, mitochondrial (Complex I-SGDH) (CI-SGDH) (NADH-ubiquinone oxidoreductase SGDH subunit) | 189 | NDUFB5 | 1 | 1 | 1 | 5.8 | 4.2 | 5.8 |
| 893 | -0.09 | 0.30 | 0.63 | A6PVX3 | 26S proteasome non-ATPase regulatory subunit 4 (Fragment) | 203 | PSMD4 | 1 | 1 | 3 | 7.4 | 7.4 | 19.7 |

| | | | | | | | | | | | | | |
|---|---|---|---|---|---|---|---|---|---|---|---|---|---|
| 894 | -0.09 | 0.41 | 0.18 | P07237 | Protein disulfide-isomerase (PDI) (EC 5.3.4.1) (Cellular thyroid hormone-binding protein) (Prolyl 4-hydroxylase subunit beta) (p55) | 508 | P4HB | 15 | 13 | 17 | 26 | 33.1 | 30.9 |
| 895 | -0.08 | 0.00 | 0.09 | H3BP42 | Apoptosis-associated speck-like protein-containing a CARD (Fragment) | 125 | PYCARD | 1 | 0 | 1 | 13.6 | 0 | 13.6 |
| 896 | -0.08 | 0.00 | 0.21 | O15143 | Actin-related protein 2/3 complex subunit 1B (Arp2/3 complex 41 kDa subunit) (p41-ARC) | 372 | ARPC1B | 2 | 0 | 2 | 8.3 | 0 | 8.3 |
| 897 | -0.08 | 0.00 | 0.22 | Q9UHN6 | Transmembrane protein 2 | 1383 | TMEM2 | 1 | 0 | 1 | 0.9 | 0 | 0.9 |
| 898 | -0.08 | 0.00 | 0.22 | A0A087 WXS7 | ATPase ASNA1 (EC 3.6.-.-) (Arsenical pump-driving ATPase) (Arsenite-stimulated ATPase) | 331 | ASNA1 | 1 | 0 | 1 | 4.8 | 0 | 4.8 |
| 899 | -0.08 | -0.49 | 0.13 | Q9UNM6 | 26S proteasome non-ATPase regulatory subunit 13 (26S proteasome regulatory subunit RPN9) (26S proteasome regulatory subunit S11) (26S proteasome regulatory subunit p40.5) | 376 | PSMD13 | 3 | 1 | 5 | 9 | 2.4 | 15.7 |
| 900 | -0.08 | -0.17 | -0.06 | Q9UBQ0 | Vacuolar protein sorting-associated protein 29 (hVPS29) (PEP11 homolog) (Vesicle protein sorting 29) | 182 | VPS29 | 1 | 3 | 1 | 7.1 | 19.2 | 7.1 |
| 901 | -0.08 | -0.16 | -0.26 | R4GMT0 | Alpha-centractin | 332 | ACTR1A | 1 | 2 | 1 | 4.8 | 7.2 | 4.8 |
| 902 | -0.08 | 0.00 | 0.15 | O75083 | WD repeat-containing protein 1 (Actin-interacting protein 1) (AIP1) (NORI-1) | 606 | WDR1 | 1 | 0 | 2 | 2.3 | 0 | 5 |
| 903 | -0.08 | 0.00 | 0.00 | M0R0Q7 | DNA ligase (EC 6.5.1.1) | 800 | LIG1 | 1 | 0 | 1 | 1.8 | 0 | 1.5 |
| 904 | -0.08 | 0.00 | 0.25 | Q96FX7 | tRNA (adenine(58)-N(1))-methyltransferase catalytic subunit TRMT61A (EC 2.1.1.220) (tRNA(m1A58)-methyltransferase subunit TRMT61A) (tRNA(m1A58)MTase subunit TRMT61A) | 289 | TRMT61A | 1 | 0 | 1 | 3.5 | 0 | 3.5 |

| | | | | | | | | | | | | | |
|---|---|---|---|---|---|---|---|---|---|---|---|---|---|
| 905 | -0.08 | -0.25 | -0.11 | Q9P000 | COMM domain-containing protein 9 | 198 | COMMD9 | 1 | 1 | 1 | 10.3 | 10.3 | 10.3 |
| 906 | -0.08 | 0.00 | 0.00 | E9PNC7 | Dr1-associated corepressor (Fragment) | 155 | DRAP1 | 1 | 0 | 0 | 7.1 | 0 | 0 |
| 907 | -0.07 | -0.07 | 0.37 | Q9NR31 | GTP-binding protein SAR1a (COPII-associated small GTPase) | 198 | SAR1A | 2 | 4 | 5 | 9.6 | 25.8 | 32.8 |
| 908 | -0.07 | 0.00 | 0.18 | M0R208 | ATP-dependent Clp protease proteolytic subunit (EC 3.4.21.92) | 190 | CLPP | 1 | 0 | 1 | 7.9 | 0 | 7.9 |
| 909 | -0.07 | -0.01 | 0.50 | Q00688 | Peptidyl-prolyl cis-trans isomerase FKBP3 (PPIase FKBP3) (EC 5.2.1.8) (25 kDa FK506-binding protein) (25 kDa FKBP) (FKBP-25) (FK506-binding protein 3) (FKBP-3) (Immunophilin FKBP25) (Rapamycin-selective 25 kDa immunophilin) (Rotamase) | 224 | FKBP3 | 2 | 6 | 3 | 14.7 | 28.1 | 18.3 |
| 910 | -0.07 | 0.00 | 0.00 | Q96P11 | Probable 28S rRNA (cytosine-C(5))-methyltransferase (EC 2.1.1.-) (NOL1-related protein) (NOL1R) (NOL1/NOP2/Sun domain family member 5) (Williams-Beuren syndrome chromosomal region 20A protein) | 429 | NSUN5 | 1 | 0 | 0 | 2.3 | 0 | 0 |
| 911 | -0.07 | -0.08 | 0.18 | P84077 | ADP-ribosylation factor 1 | 181 | ARF1 | 3 | 4 | 3 | 19.9 | 33.1 | 19.9 |
| 912 | -0.07 | -0.32 | -0.09 | P61020 | Ras-related protein Rab-5B | 215 | RAB5B | 3 | 2 | 2 | 20.1 | 13.2 | 14.4 |
| 913 | -0.07 | 0.51 | 0.08 | Q07812 | Apoptosis regulator BAX (Bcl-2-like protein 4) (Bcl2-L-4) | 192 | BAX | 1 | 1 | 1 | 6.1 | 7.3 | 6.1 |
| 914 | -0.06 | 0.26 | 0.31 | Q04917 | 14-3-3 protein eta (Protein AS1) | 246 | YWHAH | 3 | 5 | 3 | 12.2 | 26.8 | 13.4 |

| | | | | | | | | | | | | | |
|---|---|---|---|---|---|---|---|---|---|---|---|---|---|
| 915 | -0.06 | 0.00 | 0.46 | O96007 | Molybdopterin synthase catalytic subunit (EC 2.8.1.12) (MOCO1-B) (Molybdenum cofactor synthesis protein 2 large subunit) (Molybdenum cofactor synthesis protein 2B) (MOCS2B) (Molybdopterin-synthase large subunit) (MPT synthase large subunit) | 188 | MOCS2 | 1 | 0 | 1 | 8 | 0 | 8 |
| 916 | -0.06 | 0.00 | 0.97 | P82673 | 28S ribosomal protein S35, mitochondrial (MRP-S35) (S35mt) (28S ribosomal protein S28, mitochondrial) (MRP-S28) (S28mt) | 323 | MRPS35 | 1 | 0 | 1 | 3.1 | 0 | 3.1 |
| 917 | -0.06 | -0.27 | 0.61 | Q9Y678 | Coatomer subunit gamma-1 (Gamma-1-coat protein) (Gamma-1-COP) | 874 | COPG1 | 4 | 2 | 4 | 6.8 | 3.3 | 6.8 |
| 918 | -0.06 | 0.24 | -0.20 | Q9H4M9 | EH domain-containing protein 1 (PAST homolog 1) (hPAST1) (Testilin) | 534 | EHD1 | 1 | 1 | 2 | 1.7 | 2.2 | 6 |
| 919 | -0.06 | -0.38 | 0.00 | G5E9R3 | 60S ribosomal protein L37a (Ribosomal protein L37a, isoform CRA_b) | 58 | RPL37A | 1 | 2 | 0 | 15.5 | 29.3 | 0 |
| 920 | -0.05 | -0.01 | 0.28 | Q9NYP7 | Elongation of very long chain fatty acids protein 5 (EC 2.3.1.199) (3-keto acyl-CoA synthase ELOVL5) (ELOVL fatty acid elongase 5) (ELOVL FA elongase 5) (Fatty acid elongase 1) (hELO1) (Very-long-chain 3-oxoacyl-CoA synthase 5) | 299 | ELOVL5 | 1 | 1 | 1 | 14.8 | 14.8 | 14.8 |
| 921 | -0.05 | 0.00 | 0.30 | A0A0A0MSP6 | Serrate RNA effector molecule homolog (Fragment) | 148 | SRRT | 1 | 0 | 1 | 8.8 | 0 | 8.8 |
| 922 | -0.05 | 0.00 | 0.10 | P25398 | 40S ribosomal protein S12 | 132 | RPS12 | 6 | 7 | 4 | 63.6 | 50 | 42.4 |
| 923 | -0.05 | -0.34 | -0.16 | E9PQH6 | Rho-related GTP-binding protein RhoC (Fragment) | 169 | RHOC | 3 | 5 | 5 | 18.9 | 27.8 | 27.8 |
| 924 | -0.05 | 0.07 | 0.13 | P46783 | 40S ribosomal protein S10 | 165 | RPS10 | 4 | 4 | 2 | 24.2 | 24.2 | 9.7 |

| | | | | | | | | | | | | | |
|---|---|---|---|---|---|---|---|---|---|---|---|---|---|
| 925 | -0.05 | 0.05 | 0.53 | P42126 | Enoyl-CoA delta isomerase 1, mitochondrial (EC 5.3.3.8) (3,2-trans-enoyl-CoA isomerase) (Delta(3),Delta(2)-enoyl-CoA isomerase) (D3,D2-enoyl-CoA isomerase) (Dodecenoyl-CoA isomerase) | 302 | ECI1 | 2 | 2 | 2 | 9.3 | 10.6 | 9.3 |
| 926 | -0.05 | 0.39 | -0.06 | P35606 | Coatomer subunit beta' (Beta'-coat protein) (Beta'-COP) (p102) | 906 | COPB2 | 2 | 5 | 2 | 2.7 | 7.2 | 2.7 |
| 927 | -0.05 | 0.93 | 0.15 | P27797 | Calreticulin (CRP55) (Calregulin) (Endoplasmic reticulum resident protein 60) (ERp60) (HACBP) (grp60) | 417 | CALR | 7 | 5 | 7 | 15.1 | 21.3 | 14.4 |
| 928 | -0.04 | 0.00 | 0.19 | A0FGR8 | Extended synaptotagmin-2 (E-Syt2) (Chr2Syt) | 921 | ESYT2 | 1 | 0 | 1 | 3.7 | 0 | 3.7 |
| 929 | -0.04 | -0.09 | 0.24 | H0YJS4 | Eukaryotic translation initiation factor 2 subunit 1 (Fragment) | 252 | EIF2S1 | 1 | 1 | 1 | 6 | 6 | 6 |
| 930 | -0.04 | -0.13 | 0.30 | Q09161 | Nuclear cap-binding protein subunit 1 (80 kDa nuclear cap-binding protein) (CBP80) (NCBP 80 kDa subunit) | 790 | NCBP1 | 1 | 1 | 1 | 1.4 | 1.4 | 1.4 |
| 931 | -0.04 | 1.80 | 0.98 | P36957 | Dihydrolipoyllysine-residue succinyltransferase component of 2-oxoglutarate dehydrogenase complex, mitochondrial (EC 2.3.1.61) (2-oxoglutarate dehydrogenase complex component E2) (OGDC-E2) (Dihydrolipoamide succinyltransferase component of 2-oxoglutarate dehydrogenase complex) (E2K) | 453 | DLST | 1 | 2 | 3 | 2 | 6 | 7.9 |
| 932 | -0.03 | -0.49 | 0.12 | Q9BTW9 | Tubulin-specific chaperone D (Beta-tubulin cofactor D) (tfcD) (SSD-1) (Tubulin-folding cofactor D) | 1192 | TBCD | 2 | 2 | 1 | 2.6 | 2.6 | 1.3 |
| 933 | -0.03 | 0.08 | 0.11 | M0R0Y2 | Alpha-soluble NSF attachment protein | 256 | NAPA | 3 | 3 | 5 | 19.9 | 15.6 | 32.4 |

| | | | | | | | | | | | | | |
|---|---|---|---|---|---|---|---|---|---|---|---|---|---|
| 934 | -0.03 | 0.12 | 0.29 | P50213 | Isocitrate dehydrogenase [NAD] subunit alpha, mitochondrial (EC 1.1.1.41) (Isocitric dehydrogenase subunit alpha) (NAD(+)-specific ICDH subunit alpha) | 366 | IDH3A | 1 | 4 | 1 | 2.7 | 13.9 | 6.3 |
| 935 | -0.03 | 0.00 | 0.86 | Q9UIQ6 | Leucyl-cystinyl aminopeptidase (Cystinyl aminopeptidase) (EC 3.4.11.3) (Insulin-regulated membrane aminopeptidase) (Insulin-responsive aminopeptidase) (IRAP) (Oxytocinase) (OTase) (Placental leucine aminopeptidase) (P-LAP) [Cleaved into: Leucyl-cystinyl aminopeptidase, pregnancy serum form] | 1025 | LNPEP | 1 | 0 | 1 | 1.5 | 0 | 1.5 |
| 936 | -0.03 | 0.00 | 0.00 | E9PLB0 | RNA-binding protein 4B | 143 | RBM4B | 1 | 0 | 0 | 8.4 | 0 | 0 |
| 937 | -0.03 | 0.00 | 0.00 | Q9H173 | Nucleotide exchange factor SIL1 (BiP-associated protein) (BAP) | 461 | SIL1 | 1 | 0 | 0 | 2.6 | 0 | 0 |
| 938 | -0.03 | 0.00 | 1.02 | Q9UBB4 | Ataxin-10 (Brain protein E46 homolog) (Spinocerebellar ataxia type 10 protein) | 475 | ATXN10 | 1 | 0 | 3 | 2.4 | 0 | 10.2 |
| 939 | -0.03 | 0.37 | 0.24 | C9JIS1 | Guanine nucleotide-binding protein G(I)/G(S)/G(T) subunit beta-2 (Fragment) | 232 | GNB2 | 1 | 1 | 1 | 4.3 | 4.7 | 4.3 |
| 940 | -0.03 | 0.00 | 0.36 | Q16401 | 26S proteasome non-ATPase regulatory subunit 5 (26S protease subunit S5 basic) (26S proteasome subunit S5B) | 504 | PSMD5 | 2 | 0 | 1 | 6.7 | 0 | 4.1 |
| 941 | -0.02 | 0.06 | 0.62 | P36542 | ATP synthase subunit gamma, mitochondrial (F-ATPase gamma subunit) | 298 | ATP5C1 | 2 | 3 | 2 | 7.7 | 11.4 | 7.7 |
| 942 | -0.02 | 0.27 | 0.35 | Q86UP2 | Kinectin (CG-1 antigen) (Kinesin receptor) | 1357 | KTN1 | 10 | 15 | 11 | 9.8 | 14.2 | 10.2 |
| 943 | -0.02 | -0.71 | 0.16 | P36405 | ADP-ribosylation factor-like protein 3 | 182 | ARL3 | 1 | 2 | 1 | 6 | 12.6 | 6 |
| 944 | -0.02 | 0.04 | 0.26 | K7ENK9 | Vesicle-associated membrane protein 2 | 68 | VAMP2 | 1 | 1 | 1 | 25 | 25 | 25 |

| | | | | | | | | | | | | | |
|---|---|---|---|---|---|---|---|---|---|---|---|---|---|
| 945 | -0.02 | 0.00 | 1.05 | P11177 | Pyruvate dehydrogenase E1 component subunit beta, mitochondrial (PDHE1-B) (EC 1.2.4.1) | 359 | PDHB | 1 | 0 | 2 | 4.7 | 0 | 7.9 |
| 946 | -0.02 | 0.37 | 0.33 | Q04837 | Single-stranded DNA-binding protein, mitochondrial (Mt-SSB) (MtSSB) (PWP1-interacting protein 17) | 148 | SSBP1 | 3 | 3 | 4 | 28.4 | 27.7 | 33.8 |
| 947 | -0.02 | -0.29 | 0.07 | P15559 | NAD(P)H dehydrogenase [quinone] 1 (EC 1.6.5.2) (Azoreductase) (DT-diaphorase) (DTD) (Menadione reductase) (NAD(P)H:quinone oxidoreductase 1) (Phylloquinone reductase) (Quinone reductase 1) (QR1) | 274 | NQO1 | 8 | 10 | 7 | 25.8 | 32.1 | 22.5 |
| 948 | -0.01 | -0.36 | -0.07 | Q96EK6 | Glucosamine 6-phosphate N-acetyltransferase (EC 2.3.1.4) (Phosphoglucosamine acetylase) (Phosphoglucosamine transacetylase) | 184 | GNPNAT1 | 2 | 2 | 2 | 15.8 | 13.6 | 15.8 |
| 949 | -0.01 | 1.37 | 1.36 | O00264 | Membrane-associated progesterone receptor component 1 (mPR) | 195 | PGRMC1 | 2 | 4 | 2 | 9.2 | 22.6 | 12.3 |
| 950 | -0.01 | -0.08 | 0.11 | P51665 | 26S proteasome non-ATPase regulatory subunit 7 (26S proteasome regulatory subunit RPN8) (26S proteasome regulatory subunit S12) (Mov34 protein homolog) (Proteasome subunit p40) | 324 | PSMD7 | 2 | 2 | 2 | 6.5 | 5.9 | 6.5 |
| 951 | -0.01 | 0.00 | 0.16 | J9JIC5 | Protein Njmu-R1 | 396 | C17orf75 | 1 | 0 | 1 | 2.8 | 0 | 2.8 |
| 952 | -0.01 | 0.00 | 0.15 | Q9P032 | NADH dehydrogenase [ubiquinone] 1 alpha subcomplex assembly factor 4 (Hormone-regulated proliferation-associated protein of 20 kDa) | 175 | NDUFAF4 | 1 | 0 | 1 | 8 | 0 | 8 |
| 953 | -0.01 | 0.48 | 0.51 | Q96E11 | Ribosome-recycling factor, mitochondrial (RRF) (Ribosome-releasing factor, mitochondrial) | 262 | MRRF | 1 | 1 | 3 | 5.7 | 4.3 | 17.1 |

| | | | | | | | | | | | | | |
|---|---|---|---|---|---|---|---|---|---|---|---|---|---|
| 954 | -0.01 | 1.03 | 0.69 | P49411 | Elongation factor Tu, mitochondrial (EF-Tu) (P43) | 452 | TUFM | 7 | 8 | 8 | 21.2 | 26.3 | 25 |
| 955 | -0.01 | 0.00 | 0.32 | P06730 | Eukaryotic translation initiation factor 4E (eIF-4E) (eIF4E) (eIF-4F 25 kDa subunit) (mRNA cap-binding protein) | 217 | EIF4E | 1 | 0 | 1 | 6.5 | 0 | 6.5 |
| 956 | 0.00 | 0.40 | 0.44 | P10619 | Lysosomal protective protein (EC 3.4.16.5) (Carboxypeptidase C) (Carboxypeptidase L) (Cathepsin A) (Protective protein cathepsin A) (PPCA) (Protective protein for beta-galactosidase) [Cleaved into: Lysosomal protective protein 32 kDa chain; Lysosomal protective protein 20 kDa chain] | 480 | CTSA | 1 | 3 | 1 | 2.4 | 8 | 2.4 |
| 957 | 0.00 | 0.43 | 0.51 | Q71U36 | Tubulin alpha-1A chain (Alpha-tubulin 3) (Tubulin B-alpha-1) (Tubulin alpha-3 chain) | 451 | TUBA1A | 10 | 10 | 12 | 34.9 | 35.1 | 43 |
| 958 | 0.00 | 0.00 | 1.06 | Q9BQE3 | Tubulin alpha-1C chain (Alpha-tubulin 6) (Tubulin alpha-6 chain) | 449 | TUBA1C | 10 | 11 | 12 | 32.3 | 35.6 | 39.9 |
| 959 | 0.00 | 0.00 | 0.23 | V9GYG0 | ADP/ATP translocase 1 | 208 | SLC25A4 | 3 | 3 | 3 | 15.9 | 15.9 | 16.8 |
| 960 | 0.00 | 0.90 | 0.00 | Q9Y6H1 | Coiled-coil-helix-coiled-coil-helix domain-containing protein 2 (Aging-associated gene 10 protein) (HCV NS2 trans-regulated protein) (NS2TP) | 151 | CHCHD2 | 1 | 1 | 0 | 15.9 | 8.6 | 0 |
| 961 | 0.00 | 0.09 | 0.00 | Q71UM5 | 40S ribosomal protein S27-like | 84 | RPS27L | 1 | 2 | 0 | 15.5 | 28.6 | 0 |
| 962 | 0.00 | -0.41 | 0.00 | C9J1Z8 | ADP-ribosylation factor 5 (Fragment) | 150 | ARF5 | 2 | 3 | 2 | 14 | 27.3 | 14 |
| 963 | 0.00 | 1.46 | 0.00 | Q15388 | Mitochondrial import receptor subunit TOM20 homolog (Mitochondrial 20 kDa outer membrane protein) (Outer mitochondrial membrane receptor Tom20) | 145 | TOMM20 | 1 | 2 | 0 | 9 | 17.2 | 0 |

| | | | | | | | | | | | | | |
|---|---|---|---|---|---|---|---|---|---|---|---|---|---|
| 964 | 0.00 | 3.39 | 0.00 | P12273 | Prolactin-inducible protein (Gross cystic disease fluid protein 15) (GCDFP-15) (Prolactin-induced protein) (Secretory actin-binding protein) (SABP) (gp17) | 146 | PIP | 1 | 5 | 1 | 8.2 | 33.6 | 8.2 |
| 965 | 0.00 | 0.42 | 0.00 | A0A087X 1K9 | Acyl-protein thioesterase 1 | 166 | LYPLA1 | 1 | 4 | 0 | 7.8 | 27.7 | 0 |
| 966 | 0.00 | 0.00 | 1.47 | K7ESM5 | Tubulin beta-6 chain (Fragment) | 338 | TUBB6 | 2 | 4 | 5 | 7.7 | 11.5 | 18.3 |
| 967 | 0.00 | 2.32 | 0.00 | C9J1E7 | AP-1 complex subunit beta-1 (Fragment) | 578 | AP1B1 | 3 | 2 | 2 | 6.9 | 6.2 | 5 |
| 968 | 0.00 | 1.20 | 0.00 | Q6P1L8 | 39S ribosomal protein L14, mitochondrial (L14mt) (MRP-L14) (39S ribosomal protein L32, mitochondrial) (L32mt) (MRP-L32) | 145 | MRPL14 | 1 | 2 | 0 | 6.2 | 15.9 | 0 |
| 969 | 0.00 | 0.06 | 1.05 | Q5JR95 | 40S ribosomal protein S8 | 188 | RPS8 | 1 | 2 | 1 | 5.9 | 13.3 | 6.9 |
| 970 | 0.00 | - 0.70 | 0.00 | E5RGX5 | Stathmin | 168 | STMN2 | 1 | 2 | 1 | 5.4 | 11.3 | 5.4 |
| 971 | 0.00 | 1.29 | 0.00 | P00403 | Cytochrome c oxidase subunit 2 (Cytochrome c oxidase polypeptide II) | 227 | MT-CO2 | 1 | 3 | 1 | 4.4 | 20.3 | 4.4 |
| 972 | 0.00 | 1.68 | 1.63 | O15173 | Membrane-associated progesterone receptor component 2 (Progesterone membrane-binding protein) (Steroid receptor protein DG6) | 223 | PGRMC2 | 1 | 2 | 3 | 4 | 12.6 | 18.8 |
| 973 | 0.00 | 0.78 | 0.00 | P08579 | U2 small nuclear ribonucleoprotein B'' (U2 snRNP B'') | 225 | SNRPB2 | 1 | 4 | 1 | 3.6 | 16 | 3.6 |
| 974 | 0.00 | - 0.47 | 0.38 | A0A087 WZK9 | Eukaryotic translation initiation factor 3 subunit H (eIF3h) (Eukaryotic translation initiation factor 3 subunit 3) (eIF-3 gamma) (eIF3 p40 subunit) | 349 | EIF3H | 1 | 1 | 3 | 3.4 | 2.6 | 12 |
| 975 | 0.00 | 0.44 | 0.24 | E9PCY7 | Heterogeneous nuclear ribonucleoprotein H | 429 | HNRNPH1 | 1 | 3 | 1 | 2.3 | 11.7 | 4 |

| | | | | | | | | | | | | | |
|---|---|---|---|---|---|---|---|---|---|---|---|---|---|
| 976 | 0.00 | 0.66 | 1.64 | P06748 | Nucleophosmin (NPM) (Nucleolar phosphoprotein B23) (Nucleolar protein NO38) (Numatrin) | 294 | NPM1 | 0 | 6 | 5 | 0 | 26.4 | 18.9 |
| 977 | 0.00 | 0.97 | 0.89 | P26006 | Integrin alpha-3 (CD49 antigen-like family member C) (FRP-2) (Galactoprotein B3) (GAPB3) (VLA-3 subunit alpha) (CD antigen CD49c) [Cleaved into: Integrin alpha-3 heavy chain; Integrin alpha-3 light chain] | 1051 | ITGA3 | 0 | 4 | 3 | 0 | 4.8 | 3.8 |
| 978 | 0.00 | 0.60 | 0.98 | P61081 | NEDD8-conjugating enzyme Ubc12 (EC 6.3.2.-) (NEDD8 carrier protein) (NEDD8 protein ligase) (Ubiquitin-conjugating enzyme E2 M) | 183 | UBE2M | 0 | 3 | 3 | 0 | 16.9 | 19.7 |
| 979 | 0.00 | 0.40 | 0.66 | P31153 | S-adenosylmethionine synthase isoform type-2 (AdoMet synthase 2) (EC 2.5.1.6) (Methionine adenosyltransferase 2) (MAT 2) (Methionine adenosyltransferase II) (MAT-II) | 395 | MAT2A | 0 | 1 | 3 | 0 | 3.8 | 8.1 |
| 980 | 0.00 | 0.00 | 2.43 | E9PC52 | Histone-binding protein RBBP7 | 416 | RBBP7 | 0 | 0 | 3 | 0 | 0 | 13.2 |
| 981 | 0.00 | 0.00 | 0.84 | P14868 | Aspartate--tRNA ligase, cytoplasmic (EC 6.1.1.12) (Aspartyl-tRNA synthetase) (AspRS) (Cell proliferation-inducing gene 40 protein) | 501 | DARS | 0 | 0 | 3 | 0 | 0 | 7 |
| 982 | 0.00 | 0.65 | 1.84 | Q71UI9 | Histone H2A.V (H2A.F/Z) | 128 | H2AFV | 0 | 4 | 2 | 0 | 31.2 | 18.8 |
| 983 | 0.00 | -0.47 | 1.75 | A0A096LPI6 | Uncharacterized protein | 238 | | 0 | 3 | 2 | 0 | 14.7 | 13.4 |
| 984 | 0.00 | -0.55 | 0.21 | H3BPH4 | Phosphomannomutase (EC 5.4.2.8) (Fragment) | 142 | PMM2 | 0 | 2 | 2 | 0 | 16.2 | 14.1 |
| 985 | 0.00 | 0.93 | 0.48 | A0A087WX29 | TAR DNA-binding protein 43 (Fragment) | 243 | TARDBP | 0 | 1 | 2 | 0 | 4.9 | 12.3 |
| 986 | 0.00 | 2.15 | 2.68 | A0A087WZZ5 | Splicing factor 3B subunit 2 | 871 | SF3B2 | 0 | 1 | 2 | 0 | 1.6 | 3.1 |
| 987 | 0.00 | 0.83 | 1.84 | A0A0A0MSJ0 | ATP-dependent RNA helicase DDX42 | 708 | DDX42 | 0 | 1 | 2 | 0 | 1.6 | 5.8 |

| | | | | | | | | | | | | | |
|---|---|---|---|---|---|---|---|---|---|---|---|---|---|
| 988 | 0.00 | 0.50 | -0.31 | P11717 | Cation-independent mannose-6-phosphate receptor (CI Man-6-P receptor) (CI-MPR) (M6PR) (300 kDa mannose 6-phosphate receptor) (MPR 300) (Insulin-like growth factor 2 receptor) (Insulin-like growth factor II receptor) (IGF-II receptor) (M6P/IGF2 receptor) (M6P/IGF2R) (CD antigen CD222) | 2491 | IGF2R | 0 | 1 | 2 | 0 | 0.4 | 0.8 |
| 989 | 0.00 | 0.80 | 1.02 | P23634 | Plasma membrane calcium-transporting ATPase 4 (PMCA4) (EC 3.6.3.8) (Matrix-remodeling-associated protein 1) (Plasma membrane calcium ATPase isoform 4) (Plasma membrane calcium pump isoform 4) | 1241 | ATP2B4 | 0 | 1 | 2 | 0 | 1.6 | 3.1 |
| 990 | 0.00 | 0.17 | 0.63 | P30740 | Leukocyte elastase inhibitor (LEI) (Monocyte/neutrophil elastase inhibitor) (EI) (M/NEI) (Peptidase inhibitor 2) (PI-2) (Serpin B1) | 379 | SERPINB1 | 0 | 1 | 2 | 0 | 3.7 | 6.6 |
| 991 | 0.00 | 0.85 | 0.49 | Q6P587 | Acylpyruvase FAHD1, mitochondrial (EC 3.7.1.5) (Fumarylacetoacetate hydrolase domain-containing protein 1) (Oxaloacetate decarboxylase) (OAA decarboxylase) (EC 4.1.1.3) (YisK-like protein) | 224 | FAHD1 | 0 | 1 | 2 | 0 | 7.1 | 13.4 |
| 992 | 0.00 | 0.00 | 1.08 | A0A024R4E5 | High density lipoprotein binding protein (Vigilin), isoform CRA_a (Vigilin) | 1268 | HDLBP | 0 | 0 | 2 | 0 | 0 | 2.1 |
| 993 | 0.00 | 0.00 | 2.02 | A0A087WZU1 | Protein-tyrosine-phosphatase (EC 3.1.3.48) | 2299 | PTPRQ | 0 | 0 | 2 | 0 | 0 | 1.5 |
| 994 | 0.00 | 0.00 | -0.05 | E7EQL5 | Cytoplasmic dynein 1 intermediate chain 2 (Fragment) | 305 | DYNC1I2 | 0 | 0 | 2 | 0 | 0 | 9.2 |
| 995 | 0.00 | 0.00 | 0.65 | F8VR84 | UPF0160 protein MYG1, mitochondrial | 213 | C12orf10 | 0 | 0 | 2 | 0 | 0 | 12.2 |

| | | | | | | | | | | | | | |
|---|---|---|---|---|---|---|---|---|---|---|---|---|---|
| 996 | 0.00 | 0.00 | 1.06 | H7C3C5 | Polyribonucleotide nucleotidyltransferase 1, mitochondrial (Fragment) | 139 | PNPT1 | 0 | 0 | 2 | 0 | 0 | 20.1 |
| 997 | 0.00 | 0.00 | 1.17 | O95782 | AP-2 complex subunit alpha-1 (100 kDa coated vesicle protein A) (Adaptor protein complex AP-2 subunit alpha-1) (Adaptor-related protein complex 2 subunit alpha-1) (Alpha-adaptin A) (Alpha1-adaptin) (Clathrin assembly protein complex 2 alpha-A large chain) (Plasma membrane adaptor HA2/AP2 adaptin alpha A subunit) | 977 | AP2A1 | 0 | 0 | 2 | 0 | 0 | 2.2 |
| 998 | 0.00 | 0.00 | 0.35 | P07384 | Calpain-1 catalytic subunit (EC 3.4.22.52) (Calcium-activated neutral proteinase 1) (CANP 1) (Calpain mu-type) (Calpain-1 large subunit) (Cell proliferation-inducing gene 30 protein) (Micromolar-calpain) (muCANP) | 714 | CAPN1 | 0 | 0 | 2 | 0 | 0 | 2.8 |
| 999 | 0.00 | 0.00 | 0.14 | P12270 | Nucleoprotein TPR (Megator) (NPC-associated intranuclear protein) (Translocated promoter region protein) | 2363 | TPR | 0 | 0 | 2 | 0 | 0 | 2.9 |
| 1000 | 0.00 | 0.00 | 1.29 | P45974 | Ubiquitin carboxyl-terminal hydrolase 5 (EC 3.4.19.12) (Deubiquitinating enzyme 5) (Isopeptidase T) (Ubiquitin thioesterase 5) (Ubiquitin-specific-processing protease 5) | 858 | USP5 | 0 | 0 | 2 | 0 | 0 | 3.2 |
| 1001 | 0.00 | 0.00 | 1.72 | P46063 | ATP-dependent DNA helicase Q1 (EC 3.6.4.12) (DNA helicase, RecQ-like type 1) (RecQ1) (DNA-dependent ATPase Q1) (RecQ protein-like 1) | 649 | RECQL | 0 | 0 | 2 | 0 | 0 | 4.2 |

| | | | | | | | | | | | | | |
|---|---|---|---|---|---|---|---|---|---|---|---|---|---|
| 1002 | 0.00 | 0.00 | 0.92 | P67775 | Serine/threonine-protein phosphatase 2A catalytic subunit alpha isoform (PP2A-alpha) (EC 3.1.3.16) (Replication protein C) (RP-C) | 309 | PPP2CA | 0 | 0 | 2 | 0 | 0 | 11.4 |
| 1003 | 0.00 | 0.00 | -1.61 | Q02818 | Nucleobindin-1 (CALNUC) | 461 | NUCB1 | 0 | 0 | 2 | 0 | 0 | 5 |
| 1004 | 0.00 | 0.00 | 0.98 | Q5T3Q7 | HEAT repeat-containing protein 1 | 2063 | HEATR1 | 0 | 0 | 2 | 0 | 0 | 1.2 |
| 1005 | 0.00 | 0.00 | 1.23 | Q8TEM1 | Nuclear pore membrane glycoprotein 210 (Nuclear pore protein gp210) (Nuclear envelope pore membrane protein POM 210) (POM210) (Nucleoporin Nup210) (Pore membrane protein of 210 kDa) | 1887 | NUP210 | 0 | 0 | 2 | 0 | 0 | 1.4 |
| 1006 | 0.00 | 0.00 | 1.42 | Q99426 | Tubulin-folding cofactor B (Cytoskeleton-associated protein 1) (Cytoskeleton-associated protein CKAPI) (Tubulin-specific chaperone B) | 244 | TBCB | 0 | 0 | 2 | 0 | 0 | 22.3 |
| 1007 | 0.00 | 0.00 | -0.28 | Q9BTV4 | Transmembrane protein 43 (Protein LUMA) | 400 | TMEM43 | 0 | 0 | 2 | 0 | 0 | 6.2 |
| 1008 | 0.00 | 0.00 | 1.39 | Q9NR45 | Sialic acid synthase (N-acetylneuraminate synthase) (EC 2.5.1.56) (N-acetylneuraminate-9-phosphate synthase) (EC 2.5.1.57) (N-acetylneuraminic acid phosphate synthase) (N-acetylneuraminic acid synthase) | 359 | NANS | 0 | 0 | 2 | 0 | 0 | 6.7 |
| 1009 | 0.00 | 4.31 | 3.06 | P06702 | Protein S100-A9 (Calgranulin-B) (Calprotectin L1H subunit) (Leukocyte L1 complex heavy chain) (Migration inhibitory factor-related protein 14) (MRP-14) (p14) (S100 calcium-binding protein A9) | 114 | S100A9 | 0 | 5 | 1 | 0 | 44.7 | 13.2 |

| | | | | | | | | | | | | | |
|---|---|---|---|---|---|---|---|---|---|---|---|---|---|
| 1010 | 0.00 | 1.09 | 1.01 | F5GXX5 | Dolichyl-diphosphooligosaccharide--protein glycosyltransferase subunit DAD1 (Oligosaccharyl transferase subunit DAD1) (EC 2.4.99.18) | 85 | DAD1 | 0 | 4 | 1 | 0 | 47.1 | 14.1 |
| 1011 | 0.00 | 1.09 | 0.66 | H7C1U8 | Apolipoprotein O (Fragment) | 178 | APOO | 0 | 4 | 1 | 0 | 30.9 | 8.4 |
| 1012 | 0.00 | 1.08 | 0.15 | O00483 | Cytochrome c oxidase subunit NDUFA4 (Complex I-MLRQ) (CI-MLRQ) (NADH-ubiquinone oxidoreductase MLRQ subunit) | 81 | NDUFA4 | 0 | 4 | 1 | 0 | 46.9 | 12.3 |
| 1013 | 0.00 | 0.11 | 0.69 | P00568 | Adenylate kinase isoenzyme 1 (AK 1) (EC 2.7.4.3) (EC 2.7.4.6) (ATP-AMP transphosphorylase 1) (ATP:AMP phosphotransferase) (Adenylate monophosphate kinase) (Myokinase) | 194 | AK1 | 0 | 4 | 1 | 0 | 30.9 | 6.7 |
| 1014 | 0.00 | 3.75 | 0.00 | P05109 | Protein S100-A8 (Calgranulin-A) (Calprotectin L1L subunit) (Cystic fibrosis antigen) (CFAG) (Leukocyte L1 complex light chain) (Migration inhibitory factor-related protein 8) (MRP-8) (p8) (S100 calcium-binding protein A8) (Urinary stone protein band A) [Cleaved into: Protein S100-A8, N-terminally processed] | 93 | S100A8 | 0 | 4 | 1 | 0 | 39.8 | 11.8 |
| 1015 | 0.00 | 1.17 | 1.65 | P63279 | SUMO-conjugating enzyme UBC9 (EC 6.3.2.-) (SUMO-protein ligase) (Ubiquitin carrier protein 9) (Ubiquitin carrier protein I) (Ubiquitin-conjugating enzyme E2 I) (Ubiquitin-protein ligase I) (p18) | 158 | UBE2I | 0 | 4 | 1 | 0 | 34.2 | 7.6 |
| 1016 | 0.00 | 1.30 | 0.37 | Q8IVF2 | Protein AHNAK2 | 5795 | AHNAK2 | 0 | 4 | 1 | 0 | 2.1 | 2.4 |
| 1017 | 0.00 | 1.13 | 1.06 | B9A067 | MICOS complex subunit MIC60 | 711 | IMMT | 0 | 3 | 1 | 0 | 6.3 | 1.8 |
| 1018 | 0.00 | 0.58 | -0.22 | E9PN17 | ATP synthase subunit g, mitochondrial | 76 | ATP5L | 0 | 3 | 1 | 0 | 43.4 | 14.5 |

| | | | | | | | | | | | | | |
|---|---|---|---|---|---|---|---|---|---|---|---|---|---|
| 1019 | 0.00 | -0.28 | 2.29 | P00441 | Superoxide dismutase [Cu-Zn] (EC 1.15.1.1) (Superoxide dismutase 1) (hSod1) | 154 | SOD1 | 0 | 3 | 1 | 0 | 21.4 | 21.4 |
| 1020 | 0.00 | 1.13 | 1.07 | P24534 | Elongation factor 1-beta (EF-1-beta) | 225 | EEF1B2 | 0 | 3 | 1 | 0 | 26.7 | 4 |
| 1021 | 0.00 | 0.79 | 0.32 | P49458 | Signal recognition particle 9 kDa protein (SRP9) | 86 | SRP9 | 0 | 3 | 1 | 0 | 34.9 | 12.8 |
| 1022 | 0.00 | -0.23 | 0.61 | Q99584 | Protein S100-A13 (S100 calcium-binding protein A13) | 98 | S100A13 | 0 | 3 | 1 | 0 | 33.7 | 12.2 |
| 1023 | 0.00 | -0.53 | -0.13 | Q9Y3F4 | Serine-threonine kinase receptor-associated protein (MAP activator with WD repeats) (UNR-interacting protein) (WD-40 repeat protein PT-WD) | 350 | STRAP | 0 | 3 | 1 | 0 | 9.7 | 5.1 |
| 1024 | 0.00 | 1.28 | 1.22 | Q9Y5L4 | Mitochondrial import inner membrane translocase subunit Tim13 | 95 | TIMM13 | 0 | 3 | 1 | 0 | 44.2 | 11.6 |
| 1025 | 0.00 | -0.30 | -0.41 | C9J0K6 | Sorcin | 155 | SRI | 0 | 2 | 1 | 0 | 14.2 | 7.7 |
| 1026 | 0.00 | -0.21 | 0.55 | D6RCD0 | Estradiol 17-beta-dehydrogenase 11 | 256 | HSD17B11 | 0 | 2 | 1 | 0 | 10.2 | 3.9 |
| 1027 | 0.00 | 0.35 | 1.78 | E9PKZ0 | 60S ribosomal protein L8 (Fragment) | 205 | RPL8 | 0 | 2 | 1 | 0 | 11.2 | 7.8 |
| 1028 | 0.00 | 1.51 | 2.78 | E9PQR7 | Vacuolar protein sorting-associated protein 28 homolog (Fragment) | 161 | VPS28 | 0 | 2 | 1 | 0 | 21.7 | 15.5 |
| 1029 | 0.00 | 0.17 | 0.93 | F2Z2K0 | NSFL1 cofactor p47 | 274 | NSFL1C | 0 | 2 | 1 | 0 | 9.9 | 4 |
| 1030 | 0.00 | 1.98 | 1.58 | H0YLY7 | Calcineurin B homologous protein 1 (Fragment) | 91 | CHP1 | 0 | 2 | 1 | 0 | 30.8 | 17.6 |
| 1031 | 0.00 | 1.19 | 0.91 | H0YNK8 | ER membrane protein complex subunit 4 | 102 | EMC4 | 0 | 2 | 1 | 0 | 25.5 | 15.7 |
| 1032 | 0.00 | 2.70 | 2.16 | J3QQY2 | Transmembrane and coiled-coil domain-containing protein 1 (Transmembrane and coiled-coil domains 1, isoform CRA_c) | 104 | TMCO1 | 0 | 2 | 1 | 0 | 27.9 | 14.4 |
| 1033 | 0.00 | 2.10 | 1.33 | O14925 | Mitochondrial import inner membrane translocase subunit Tim23 | 209 | TIMM23 | 0 | 2 | 1 | 0 | 12 | 8.1 |

| | | | | | | | | | | | | | |
|---|---|---|---|---|---|---|---|---|---|---|---|---|---|
| 1034 | 0.00 | 0.91 | 0.91 | O43181 | NADH dehydrogenase [ubiquinone] iron-sulfur protein 4, mitochondrial (Complex I-18 kDa) (CI-18 kDa) (Complex I-AQDQ) (CI-AQDQ) (NADH-ubiquinone oxidoreductase 18 kDa subunit) | 175 | NDUFS4 | 0 | 2 | 1 | 0 | 14.3 | 8.6 |
| 1035 | 0.00 | -0.53 | -0.16 | O43399 | Tumor protein D54 (hD54) (Tumor protein D52-like 2) | 206 | TPD52L2 | 0 | 2 | 1 | 0 | 12.9 | 6.1 |
| 1036 | 0.00 | -0.93 | 1.31 | O75368 | SH3 domain-binding glutamic acid-rich-like protein | 114 | SH3BGRL | 0 | 2 | 1 | 0 | 21.9 | 15.8 |
| 1037 | 0.00 | 1.53 | 0.32 | P20674 | Cytochrome c oxidase subunit 5A, mitochondrial (Cytochrome c oxidase polypeptide Va) | 150 | COX5A | 0 | 2 | 1 | 0 | 15.3 | 12.7 |
| 1038 | 0.00 | 1.31 | 1.33 | P56134 | ATP synthase subunit f, mitochondrial | 94 | ATP5J2 | 0 | 2 | 1 | 0 | 49 | 14.3 |
| 1039 | 0.00 | 0.32 | 0.37 | P61009 | Signal peptidase complex subunit 3 (EC 3.4.-.-) (Microsomal signal peptidase 22/23 kDa subunit) (SPC22/23) (SPase 22/23 kDa subunit) | 180 | SPCS3 | 0 | 2 | 1 | 0 | 12.8 | 6.1 |
| 1040 | 0.00 | -0.60 | 0.19 | P62993 | Growth factor receptor-bound protein 2 (Adapter protein GRB2) (Protein Ash) (SH2/SH3 adapter GRB2) | 217 | GRB2 | 0 | 2 | 1 | 0 | 9.7 | 3.4 |
| 1041 | 0.00 | 1.04 | 1.61 | P84090 | Enhancer of rudimentary homolog | 104 | ERH | 0 | 2 | 1 | 0 | 21.2 | 10.6 |
| 1042 | 0.00 | 0.90 | 0.88 | Q03135 | Caveolin-1 | 178 | CAV1 | 0 | 2 | 1 | 0 | 10.7 | 7.9 |
| 1043 | 0.00 | 0.83 | 1.36 | Q16629 | Serine/arginine-rich splicing factor 7 (Splicing factor 9G8) (Splicing factor, arginine/serine-rich 7) | 238 | SRSF7 | 0 | 2 | 1 | 0 | 19.7 | 9.1 |
| 1044 | 0.00 | 0.19 | 0.76 | Q5JWB9 | Transmembrane protein 230 (Fragment) | 74 | TMEM230 | 0 | 2 | 1 | 0 | 33.8 | 18.9 |

| | | | | | | | | | | | | | |
|---|---|---|---|---|---|---|---|---|---|---|---|---|---|
| 1045 | 0.00 | 0.17 | 1.10 | Q5VTU3 | Dynein light chain Tctex-type 1 (Dynein, light chain, Tctex-type 1, isoform CRA_a) | 92 | DYNLT1 | 0 | 2 | 1 | 0 | 37 | 17.4 |
| 1046 | 0.00 | 1.36 | -1.05 | Q9C002 | Normal mucosa of esophagus-specific gene 1 protein (Protein FOAP-11) | 83 | NMES1 | 0 | 2 | 1 | 0 | 27.7 | 9.6 |
| 1047 | 0.00 | 1.19 | -0.05 | Q9NVJ2 | ADP-ribosylation factor-like protein 8B (ADP-ribosylation factor-like protein 10C) (Novel small G protein indispensable for equal chromosome segregation 1) | 186 | ARL8B | 0 | 2 | 1 | 0 | 21.5 | 8.6 |
| 1048 | 0.00 | 0.67 | 0.84 | Q9Y248 | DNA replication complex GINS protein PSF2 (GINS complex subunit 2) | 185 | GINS2 | 0 | 2 | 1 | 0 | 14.1 | 7 |
| 1049 | 0.00 | -0.10 | 0.03 | A0A087WU14 | Secretory carrier-associated membrane protein 1 (Fragment) | 206 | SCAMP1 | 0 | 1 | 1 | 0 | 11.2 | 11.2 |
| 1050 | 0.00 | 3.16 | 4.57 | C9JHF5 | Mitochondrial fission factor (Fragment) | 138 | MFF | 0 | 1 | 1 | 0 | 13 | 19.6 |
| 1051 | 0.00 | 1.28 | 0.47 | E7EU96 | Casein kinase II subunit alpha | 385 | CSNK2A1 | 0 | 1 | 1 | 0 | 3.9 | 3.9 |
| 1052 | 0.00 | 0.00 | 1.38 | E7EW33 | Cytoplasmic FMR1-interacting protein 2 | 1057 | CYFIP2 | 0 | 1 | 1 | 0 | 0.9 | 1 |
| 1053 | 0.00 | 0.87 | 0.96 | F5GYK7 | Glycerol-3-phosphate dehydrogenase (EC 1.1.5.3) | 379 | GPD2 | 0 | 1 | 1 | 0 | 2.4 | 4.2 |
| 1054 | 0.00 | 0.39 | 0.36 | F5H6P7 | Protein mago nashi homolog 2 | 102 | MAGOHB | 0 | 1 | 1 | 0 | 10.8 | 10.8 |
| 1055 | 0.00 | 1.67 | 2.60 | G3V1U5 | Golgi transport 1 homolog B (S. cerevisiae), isoform CRA_c (Vesicle transport protein GOT1B) | 74 | GOLT1B | 0 | 1 | 1 | 0 | 18.9 | 20.3 |
| 1056 | 0.00 | 1.11 | 2.91 | G3V2V8 | Epididymal secretory protein E1 (Fragment) | 122 | NPC2 | 0 | 1 | 1 | 0 | 7.4 | 18.9 |
| 1057 | 0.00 | 0.71 | 1.30 | G3V4F2 | Acyl-coenzyme A thioesterase 1 | 395 | ACOT1 | 0 | 1 | 1 | 0 | 2.8 | 2.8 |
| 1058 | 0.00 | 9.25 | 0.84 | H0YE25 | Parkinson disease 7 domain-containing protein 1 (Fragment) | 144 | PDDC1 | 0 | 1 | 1 | 0 | 6.2 | 6.9 |
| 1059 | 0.00 | 1.35 | 1.73 | H7BZ81 | All-trans-retinol 13,14-reductase (Fragment) | 320 | RETSAT | 0 | 1 | 1 | 0 | 3.8 | 5.6 |

| | | | | | | | | | | | | | |
|---|---|---|---|---|---|---|---|---|---|---|---|---|---|
| 1060 | 0.00 | 1.12 | 1.15 | H7C3P7 | Ras-related protein Ral-A (Fragment) | 164 | RALA | 0 | 1 | 1 | 0 | 9.1 | 9.1 |
| 1061 | 0.00 | 2.54 | 0.07 | J3KT68 | Transmembrane protein 97 | 92 | TMEM97 | 0 | 1 | 1 | 0 | 16.3 | 8.7 |
| 1062 | 0.00 | 1.46 | 1.62 | J3QLR8 | 28S ribosomal protein S23, mitochondrial | 152 | MRPS23 | 0 | 1 | 1 | 0 | 6.6 | 6.6 |
| 1063 | 0.00 | 1.39 | 1.87 | K7EM02 | Katanin p60 ATPase-containing subunit A-like 2 (Fragment) | 128 | KATNAL2 | 0 | 1 | 1 | 0 | 9.4 | 9.4 |
| 1064 | 0.00 | 1.19 | 1.17 | K7ENI6 | Uncharacterized protein | 41 | | 0 | 1 | 1 | 0 | 68.3 | 68.3 |
| 1065 | 0.00 | 0.51 | 1.33 | M0QY80 | Persulfide dioxygenase ETHE1, mitochondrial | 95 | ETHE1 | 0 | 1 | 1 | 0 | 14.7 | 14.7 |
| 1066 | 0.00 | 0.31 | -0.20 | O14672 | Disintegrin and metalloproteinase domain-containing protein 10 (ADAM 10) (EC 3.4.24.81) (CDw156) (Kuzbanian protein homolog) (Mammalian disintegrin-metalloprotease) (CD antigen CD156c) | 748 | ADAM10 | 0 | 1 | 1 | 0 | 1.6 | 1.6 |
| 1067 | 0.00 | 0.85 | -0.69 | O14975 | Very long-chain acyl-CoA synthetase (VLACS) (VLCS) (EC 6.2.1.-) (Fatty acid transport protein 2) (FATP-2) (Fatty-acid-coenzyme A ligase, very long-chain 1) (Long-chain-fatty-acid--CoA ligase) (EC 6.2.1.3) (Solute carrier family 27 member 2) (THCA-CoA ligase) (Very long-chain-fatty-acid-CoA ligase) | 620 | SLC27A2 | 0 | 1 | 1 | 0 | 3.5 | 1.9 |

| | | | | | | | | | | | | | |
|---|---|---|---|---|---|---|---|---|---|---|---|---|---|
| 1068 | 0.00 | 0.68 | 1.37 | O75915 | PRA1 family protein 3 (ADP-ribosylation factor-like protein 6-interacting protein 5) (ARL-6-interacting protein 5) (Aip-5) (Cytoskeleton-related vitamin A-responsive protein) (Dermal papilla-derived protein 11) (GTRAP3-18) (Glutamate transporter EAAC1-interacting protein) (JM5) (Prenylated Rab acceptor protein 2) (Protein JWa) (Putative MAPK-activating protein PM27) | 188 | ARL6IP5 | 0 | 1 | 1 | 0 | 10.1 | 10.1 |
| 1069 | 0.00 | 0.82 | 0.75 | P01034 | Cystatin-C (Cystatin-3) (Gamma-trace) (Neuroendocrine basic polypeptide) (Post-gamma-globulin) | 146 | CST3 | 0 | 1 | 1 | 0 | 7.5 | 7.5 |
| 1070 | 0.00 | -0.97 | 0.31 | P11908 | Ribose-phosphate pyrophosphokinase 2 (EC 2.7.6.1) (PPRibP) (Phosphoribosyl pyrophosphate synthase II) (PRS-II) | 318 | PRPS2 | 0 | 1 | 1 | 0 | 3.1 | 3.1 |
| 1071 | 0.00 | 1.98 | 1.09 | P14174 | Macrophage migration inhibitory factor (MIF) (EC 5.3.2.1) (Glycosylation-inhibiting factor) (GIF) (L-dopachrome isomerase) (L-dopachrome tautomerase) (EC 5.3.3.12) (Phenylpyruvate tautomerase) | 115 | MIF | 0 | 1 | 1 | 0 | 7.8 | 9.6 |
| 1072 | 0.00 | 2.17 | 2.22 | P14678 | Small nuclear ribonucleoprotein-associated proteins B and B' (snRNP-B) (Sm protein B/B') (Sm-B/B') (SmB/B') | 240 | SNRPB | 0 | 1 | 1 | 0 | 6.1 | 6.1 |
| 1073 | 0.00 | 0.55 | 2.01 | P15927 | Replication protein A 32 kDa subunit (RP-A p32) (Replication factor A protein 2) (RF-A protein 2) (Replication protein A 34 kDa subunit) (RP-A p34) | 270 | RPA2 | 0 | 1 | 1 | 0 | 3 | 5.2 |

| | | | | | | | | | | | | | |
|---|---|---|---|---|---|---|---|---|---|---|---|---|---|
| 1074 | 0.00 | 2.09 | 2.25 | P17152 | Transmembrane protein 11, mitochondrial (Protein PM1) (Protein PMI) | 192 | TMEM11 | 0 | 1 | 1 | 0 | 8.3 | 8.3 |
| 1075 | 0.00 | 1.26 | 0.33 | P17568 | NADH dehydrogenase [ubiquinone] 1 beta subcomplex subunit 7 (Cell adhesion protein SQM1) (Complex I-B18) (CI-B18) (NADH-ubiquinone oxidoreductase B18 subunit) | 137 | NDUFB7 | 0 | 1 | 1 | 0 | 7.3 | 7.3 |
| 1076 | 0.00 | -0.18 | 1.27 | P35754 | Glutaredoxin-1 (Thioltransferase-1) (TTase-1) | 106 | GLRX | 0 | 1 | 1 | 0 | 10.4 | 10.4 |
| 1077 | 0.00 | -0.11 | 0.60 | P52434 | DNA-directed RNA polymerases I, II, and III subunit RPABC3 (RNA polymerases I, II, and III subunit ABC3) (DNA-directed RNA polymerase II subunit H) (DNA-directed RNA polymerases I, II, and III 17.1 kDa polypeptide) (RPB17) (RPB8 homolog) (hRPB8) | 150 | POLR2H | 0 | 1 | 1 | 0 | 7.4 | 9.8 |
| 1078 | 0.00 | 2.34 | 1.27 | P60903 | Protein S100-A10 (Calpactin I light chain) (Calpactin-1 light chain) (Cellular ligand of annexin II) (S100 calcium-binding protein A10) (p10 protein) (p11) | 97 | S100A10 | 0 | 1 | 1 | 0 | 17.5 | 17.5 |
| 1079 | 0.00 | -0.57 | -0.15 | P61011 | Signal recognition particle 54 kDa protein (SRP54) | 504 | SRP54 | 0 | 1 | 1 | 0 | 2.4 | 2.4 |
| 1080 | 0.00 | 0.30 | 0.36 | P62273 | 40S ribosomal protein S29 | 56 | RPS29 | 0 | 1 | 1 | 0 | 19.6 | 14.3 |
| 1081 | 0.00 | 2.39 | 0.89 | P78347 | General transcription factor II-I (GTFII-I) (TFII-I) (Bruton tyrosine kinase-associated protein 135) (BAP-135) (BTK-associated protein 135) (SRF-Phox1-interacting protein) (SPIN) (Williams-Beuren syndrome chromosomal region 6 protein) | 998 | GTF2I | 0 | 1 | 1 | 0 | 0.9 | 1.3 |
| 1082 | 0.00 | 6.44 | 2.92 | Q15058 | Kinesin-like protein KIF14 | 1648 | KIF14 | 0 | 1 | 1 | 0 | 0.7 | 0.7 |

| | | | | | | | | | | | | | |
|---|---|---|---|---|---|---|---|---|---|---|---|---|---|
| 1083 | 0.00 | 1.00 | 2.57 | Q15149 | Plectin (PCN) (PLTN) (Hemidesmosomal protein 1) (HD1) (Plectin-1) | 4684 | PLEC | 0 | 1 | 1 | 0 | 0.3 | 0.3 |
| 1084 | 0.00 | 0.03 | 3.59 | Q6P2Q9 | Pre-mRNA-processing-splicing factor 8 (220 kDa U5 snRNP-specific protein) (PRP8 homolog) (Splicing factor Prp8) (p220) | 2335 | PRPF8 | 0 | 1 | 1 | 0 | 0.3 | 1.1 |
| 1085 | 0.00 | 1.89 | 1.54 | Q8N6L1 | Keratinocyte-associated protein 2 (KCP-2) | 136 | KRTCAP2 | 0 | 1 | 1 | 0 | 12.5 | 12.5 |
| 1086 | 0.00 | 0.42 | 1.32 | Q93009 | Ubiquitin carboxyl-terminal hydrolase 7 (EC 3.4.19.12) (Deubiquitinating enzyme 7) (Herpesvirus-associated ubiquitin-specific protease) (Ubiquitin thioesterase 7) (Ubiquitin-specific-processing protease 7) | 1102 | USP7 | 0 | 1 | 1 | 0 | 1.2 | 1.1 |
| 1087 | 0.00 | 0.29 | 1.00 | Q96TA1 | Niban-like protein 1 (Meg-3) (Melanoma invasion by ERK) (MINERVA) (Protein FAM129B) | 746 | FAM129B | 0 | 1 | 1 | 0 | 1.8 | 1.8 |
| 1088 | 0.00 | -0.31 | 1.90 | Q99436 | Proteasome subunit beta type-7 (EC 3.4.25.1) (Macropain chain Z) (Multicatalytic endopeptidase complex chain Z) (Proteasome subunit Z) | 277 | PSMB7 | 0 | 1 | 1 | 0 | 3.2 | 4.7 |
| 1089 | 0.00 | 0.00 | 1.24 | Q9NPA8 | Transcription and mRNA export factor ENY2 (Enhancer of yellow 2 transcription factor homolog) | 101 | ENY2 | 0 | 1 | 1 | 0 | 9.4 | 17.7 |
| 1090 | 0.00 | 2.65 | 2.47 | Q9UBI6 | Guanine nucleotide-binding protein G(I)/G(S)/G(O) subunit gamma-12 | 72 | GNG12 | 0 | 1 | 1 | 0 | 22.2 | 25 |
| 1091 | 0.00 | -0.20 | 0.30 | Q9UHV9 | Prefoldin subunit 2 | 154 | PFDN2 | 0 | 1 | 1 | 0 | 7.8 | 7.8 |
| 1092 | 0.00 | 1.26 | 0.97 | S4R3B5 | Protein transport protein Sec61 subunit beta | 42 | SEC61B | 0 | 1 | 1 | 0 | 23.8 | 23.8 |
| 1093 | 0.00 | -0.58 | -0.37 | U3KQT1 | S-formylglutathione hydrolase (Fragment) | 120 | ESD | 0 | 1 | 1 | 0 | 7.5 | 7.5 |

| | | | | | | | | | | | | | |
|---|---|---|---|---|---|---|---|---|---|---|---|---|---|
| 1094 | 0.00 | 0.00 | 0.12 | A0A087 WTA2 | Protein SYNJ2BP-COX16 | 40 | SYNJ2BP-COX16 | 0 | 0 | 1 | 0 | 0 | 35 |
| 1095 | 0.00 | 0.00 | 2.48 | C9JUG7 | F-actin-capping protein subunit alpha-2 | 146 | CAPZA2 | 0 | 0 | 1 | 0 | 0 | 12.3 |
| 1096 | 0.00 | 0.00 | 1.64 | D6RDY6 | Signal recognition particle subunit SRP72 (Fragment) | 357 | SRP72 | 0 | 0 | 1 | 0 | 0 | 5.3 |
| 1097 | 0.00 | 0.00 | 1.03 | D6RGI3 | Septin 11, isoform CRA_b (Septin-11) | 425 | sept11 | 0 | 0 | 1 | 0 | 0 | 3.1 |
| 1098 | 0.00 | 0.00 | -0.22 | D6RH17 | Alcohol dehydrogenase 6 (Fragment) | 257 | ADH6 | 0 | 0 | 1 | 0 | 0 | 3.9 |
| 1099 | 0.00 | 0.00 | 1.14 | D6RHZ5 | Protein transport protein Sec31A | 877 | SEC31A | 0 | 0 | 1 | 0 | 0 | 1.5 |
| 1100 | 0.00 | 0.00 | -0.15 | E5RIX8 | Tubulin-specific chaperone A | 79 | TBCA | 0 | 0 | 1 | 0 | 0 | 12.7 |
| 1101 | 0.00 | 0.00 | 2.51 | E7EMM4 | Acid ceramidase | 370 | ASAH1 | 0 | 0 | 1 | 0 | 0 | 4.6 |
| 1102 | 0.00 | 0.00 | 2.39 | F5GX77 | Multifunctional methyltransferase subunit TRM112-like protein | 106 | TRMT112 | 0 | 0 | 1 | 0 | 0 | 12.3 |
| 1103 | 0.00 | 0.00 | 0.60 | F8VVL1 | Density-regulated protein | 160 | DENR | 0 | 0 | 1 | 0 | 0 | 5 |
| 1104 | 0.00 | 0.00 | 0.43 | F8VVW8 | 3'(2'),5'-bisphosphate nucleotidase 1 (Fragment) | 130 | BPNT1 | 0 | 0 | 1 | 0 | 0 | 8.5 |
| 1105 | 0.00 | 0.00 | 0.54 | F8WJN3 | Cleavage and polyadenylation-specificity factor subunit 6 | 478 | CPSF6 | 0 | 0 | 1 | 0 | 0 | 2.9 |
| 1106 | 0.00 | 0.00 | 0.38 | H0Y9Q1 | Cytosol aminopeptidase (Fragment) | 208 | LAP3 | 0 | 0 | 1 | 0 | 0 | 8.7 |
| 1107 | 0.00 | 0.00 | 1.20 | H0YDS0 | Ubiquilin-1 (Fragment) | 157 | UBQLN1 | 0 | 0 | 1 | 0 | 0 | 10.2 |
| 1108 | 0.00 | 0.00 | 3.28 | H0YE46 | RNA-binding protein 25 (Fragment) | 73 | RBM25 | 0 | 0 | 1 | 0 | 0 | 20.5 |
| 1109 | 0.00 | 0.00 | 4.86 | H0YFA4 | Cysteine-rich protein 2 (Fragment) | 192 | CRIP2 | 0 | 0 | 1 | 0 | 0 | 16.7 |
| 1110 | 0.00 | 0.00 | 1.27 | I3L252 | Bifunctional ATP-dependent dihydroxyacetone kinase/FAD-AMP lyase (cyclizing) (Fragment) | 219 | DAK | 0 | 0 | 1 | 0 | 0 | 5.5 |
| 1111 | 0.00 | 0.00 | 0.96 | J3QLD9 | Flotillin-2 (HCG1998851, isoform CRA_h) | 428 | FLOT2 | 0 | 0 | 1 | 0 | 0 | 4.4 |
| 1112 | 0.00 | 0.00 | 1.63 | K7EMV3 | Histone H3 | 92 | H3F3B | 0 | 0 | 1 | 0 | 0 | 7.6 |
| 1113 | 0.00 | 0.00 | 1.23 | K7EP16 | Eukaryotic translation initiation factor 3 subunit G (Fragment) | 118 | EIF3G | 0 | 0 | 1 | 0 | 0 | 13.6 |

| | | | | | | | | | | | | | |
|---|---|---|---|---|---|---|---|---|---|---|---|---|---|
| 1114 | 0.00 | 0.00 | 1.70 | O00170 | AH receptor-interacting protein (AIP) (Aryl-hydrocarbon receptor-interacting protein) (HBV X-associated protein 2) (XAP-2) (Immunophilin homolog ARA9) | 330 | AIP | 0 | 0 | 1 | 0 | 0 | 4.2 |
| 1115 | 0.00 | 0.00 | 0.77 | O43852 | Calumenin (Crocalbin) (IEF SSP 9302) | 315 | CALU | 0 | 0 | 1 | 0 | 0 | 8.4 |
| 1116 | 0.00 | 0.00 | 3.17 | O43896 | Kinesin-like protein KIF1C | 1103 | KIF1C | 0 | 0 | 1 | 0 | 0 | 0.8 |
| 1117 | 0.00 | 0.00 | 1.24 | O60869 | Endothelial differentiation-related factor 1 (EDF-1) (Multiprotein-bridging factor 1) (MBF1) | 148 | EDF1 | 0 | 0 | 1 | 0 | 0 | 10.1 |
| 1118 | 0.00 | 0.00 | 0.95 | O75436 | Vacuolar protein sorting-associated protein 26A (Vesicle protein sorting 26A) (hVPS26) | 327 | VPS26A | 0 | 0 | 1 | 0 | 0 | 5.5 |
| 1119 | 0.00 | 0.00 | 4.01 | O75937 | DnaJ homolog subfamily C member 8 (Splicing protein spf31) | 253 | DNAJC8 | 0 | 0 | 1 | 0 | 0 | 11.5 |
| 1120 | 0.00 | 0.00 | 1.76 | P12074 | Cytochrome c oxidase subunit 6A1, mitochondrial (Cytochrome c oxidase polypeptide VIa-liver) (Cytochrome c oxidase subunit VIA-liver) (COX VIa-L) | 109 | COX6A1 | 0 | 0 | 1 | 0 | 0 | 16.5 |
| 1121 | 0.00 | 0.00 | 2.35 | P13995 | Bifunctional methylenetetrahydrofolate dehydrogenase/cyclohydrolase, mitochondrial [Includes: NAD-dependent methylenetetrahydrofolate dehydrogenase (EC 1.5.1.15); Methenyltetrahydrofolate cyclohydrolase (EC 3.5.4.9)] | 350 | MTHFD2 | 0 | 0 | 1 | 0 | 0 | 9.3 |
| 1122 | 0.00 | 0.00 | 1.94 | P17174 | Aspartate aminotransferase, cytoplasmic (cAspAT) (EC 2.6.1.1) (EC 2.6.1.3) (Cysteine aminotransferase, cytoplasmic) (Cysteine transaminase, cytoplasmic) (cCAT) (Glutamate oxaloacetate transaminase 1) (Transaminase A) | 413 | GOT1 | 0 | 0 | 1 | 0 | 0 | 3.6 |

| | | | | | | | | | | | | | |
|---|---|---|---|---|---|---|---|---|---|---|---|---|---|
| 1123 | 0.00 | 0.00 | 2.22 | P18084 | Integrin beta-5 | 799 | ITGB5 | 0 | 0 | 1 | 0 | 0 | 2.5 |
| 1124 | 0.00 | 0.00 | -0.01 | P20073 | Annexin A7 (Annexin VII) (Annexin-7) (Synexin) | 488 | ANXA7 | 0 | 0 | 1 | 0 | 0 | 1.9 |
| 1125 | 0.00 | 0.00 | 0.79 | P25685 | DnaJ homolog subfamily B member 1 (DnaJ protein homolog 1) (Heat shock 40 kDa protein 1) (HSP40) (Heat shock protein 40) (Human DnaJ protein 1) (hDj-1) | 340 | DNAJB1 | 0 | 0 | 1 | 0 | 0 | 2.9 |
| 1126 | 0.00 | 0.00 | 2.33 | P26196 | Probable ATP-dependent RNA helicase DDX6 (EC 3.6.4.13) (ATP-dependent RNA helicase p54) (DEAD box protein 6) (Oncogene RCK) | 483 | DDX6 | 0 | 0 | 1 | 0 | 0 | 3.5 |
| 1127 | 0.00 | 0.00 | 1.07 | P39023 | 60S ribosomal protein L3 (HIV-1 TAR RNA-binding protein B) (TARBP-B) | 403 | RPL3 | 0 | 0 | 1 | 0 | 0 | 2 |
| 1128 | 0.00 | 0.00 | 0.54 | P39748 | Flap endonuclease 1 (FEN-1) (EC 3.1.-.-) (DNase IV) (Flap structure-specific endonuclease 1) (Maturation factor 1) (MF1) (hFEN-1) | 380 | FEN1 | 0 | 0 | 1 | 0 | 0 | 4.2 |
| 1129 | 0.00 | 0.00 | 1.53 | Q13045 | Protein flightless-1 homolog | 1269 | FLII | 0 | 0 | 1 | 0 | 0 | 0.8 |
| 1130 | 0.00 | 0.00 | 0.51 | Q13085 | Acetyl-CoA carboxylase 1 (ACC1) (EC 6.4.1.2) (ACC-alpha) [Includes: Biotin carboxylase (EC 6.3.4.14)] | 2346 | ACACA | 0 | 0 | 1 | 0 | 0 | 0.5 |
| 1131 | 0.00 | 0.00 | 0.36 | Q14320 | Protein FAM50A (Protein HXC-26) (Protein XAP-5) | 339 | FAM50A | 0 | 0 | 1 | 0 | 0 | 4.7 |
| 1132 | 0.00 | 0.00 | -0.10 | Q5QPP3 | UDP-glucose 4-epimerase (Fragment) | 227 | GALE | 0 | 0 | 1 | 0 | 0 | 6.2 |
| 1133 | 0.00 | 0.00 | 0.18 | Q7KZ85 | Transcription elongation factor SPT6 (hSPT6) (Histone chaperone suppressor of Ty6) (Tat-cotransactivator 2 protein) (Tat-CT2 protein) | 1726 | SUPT6H | 0 | 0 | 1 | 0 | 0 | 1.5 |

| | | | | | | | | | | | | | |
|---|---|---|---|---|---|---|---|---|---|---|---|---|---|
| 1134 | 0.00 | 0.00 | 1.28 | Q8WVM8 | Sec1 family domain-containing protein 1 (SLY1 homolog) (Sly1p) (Syntaxin-binding protein 1-like 2) | 642 | SCFD1 | 0 | 0 | 1 | 0 | 0 | 2.5 |
| 1135 | 0.00 | 0.00 | -0.14 | Q8WZ42 | Titin (EC 2.7.11.1) (Connectin) (Rhabdomyosarcoma antigen MU-RMS-40.14) | 34350 | TTN | 0 | 0 | 1 | 0 | 0 | 0 |
| 1136 | 0.00 | 0.00 | 1.88 | Q92887 | Canalicular multispecific organic anion transporter 1 (ATP-binding cassette sub-family C member 2) (Canalicular multidrug resistance protein) (Multidrug resistance-associated protein 2) | 1545 | ABCC2 | 0 | 0 | 1 | 0 | 0 | 1 |
| 1137 | 0.00 | 0.00 | 0.76 | Q93008 | Probable ubiquitin carboxyl-terminal hydrolase FAF-X (EC 3.4.19.12) (Deubiquitinating enzyme FAF-X) (Fat facets in mammals) (hFAM) (Fat facets protein-related, X-linked) (Ubiquitin thioesterase FAF-X) (Ubiquitin-specific protease 9, X chromosome) (Ubiquitin-specific-processing protease FAF-X) | 2570 | USP9X | 0 | 0 | 1 | 0 | 0 | 0.5 |
| 1138 | 0.00 | 0.00 | 1.59 | Q96BS2 | Calcineurin B homologous protein 3 (Tescalcin) (TSC) | 214 | TESC | 0 | 0 | 1 | 0 | 0 | 8 |
| 1139 | 0.00 | 0.00 | 2.18 | Q96GC9 | Vacuole membrane protein 1 (Transmembrane protein 49) | 406 | VMP1 | 0 | 0 | 1 | 0 | 0 | 14 |
| 1140 | 0.00 | 0.00 | 0.54 | Q96HE7 | ERO1-like protein alpha (ERO1-L) (ERO1-L-alpha) (EC 1.8.4.-) (Endoplasmic oxidoreductin-1-like protein) (Oxidoreductin-1-L-alpha) | 468 | ERO1L | 0 | 0 | 1 | 0 | 0 | 3 |
| 1141 | 0.00 | 0.00 | 2.13 | Q96K17 | Transcription factor BTF3 homolog 4 (Basic transcription factor 3-like 4) | 158 | BTF3L4 | 0 | 0 | 1 | 0 | 0 | 26 |

| | | | | | | | | | | | | | |
|---|---|---|---|---|---|---|---|---|---|---|---|---|---|
| 1142 | 0.00 | 0.00 | 0.60 | Q9BSH4 | Translational activator of cytochrome c oxidase 1 (Coiled-coil domain-containing protein 44) (Translational activator of mitochondrially-encoded cytochrome c oxidase I) | 297 | TACO1 | 0 | 0 | 1 | 0 | 0 | 6.1 |
| 1143 | 0.00 | 0.00 | 0.99 | Q9BZX2 | Uridine-cytidine kinase 2 (UCK 2) (EC 2.7.1.48) (Cytidine monophosphokinase 2) (Testis-specific protein TSA903) (Uridine monophosphokinase 2) | 261 | UCK2 | 0 | 0 | 1 | 0 | 0 | 6.9 |
| 1144 | 0.00 | 0.00 | 0.28 | Q9H0U6 | 39S ribosomal protein L18, mitochondrial (L18mt) (MRP-L18) | 180 | MRPL18 | 0 | 0 | 1 | 0 | 0 | 5 |
| 1145 | 0.00 | 0.00 | 0.24 | Q9Y512 | Sorting and assembly machinery component 50 homolog (Transformation-related gene 3 protein) (TRG-3) | 469 | SAMM50 | 0 | 0 | 1 | 0 | 0 | 3.2 |
| 1146 | 0.00 | 0.00 | -0.11 | R4GN98 | Protein S100 (S100 calcium-binding protein) (Fragment) | 85 | S100A6 | 0 | 0 | 1 | 0 | 0 | 8.2 |
| 1147 | 0.00 | 0.00 | 1.29 | V9H019 | DNA mismatch repair protein Msh2 (MSH2 protein) | 810 | MSH2 | 0 | 0 | 1 | 0 | 0 | 1.4 |
| 1148 | 0.00 | 4.06 | 0.00 | P15924 | Desmoplakin (DP) (250/210 kDa paraneoplastic pemphigus antigen) | 2871 | DSP | 0 | 35 | 0 | 0 | 13.1 | 0 |
| 1149 | 0.00 | 4.04 | 0.00 | Q02413 | Desmoglein-1 (Cadherin family member 4) (Desmosomal glycoprotein 1) (DG1) (DGI) (Pemphigus foliaceus antigen) | 1049 | DSG1 | 0 | 12 | 0 | 0 | 16.2 | 0 |
| 1150 | 0.00 | 3.40 | 0.00 | P31944 | Caspase-14 (CASP-14) (EC 3.4.22.-) [Cleaved into: Caspase-14 subunit p17, mature form; Caspase-14 subunit p10, mature form; Caspase-14 subunit p20, intermediate form; Caspase-14 subunit p8, intermediate form] | 242 | CASP14 | 0 | 5 | 0 | 0 | 17.8 | 0 |
| 1151 | 0.00 | 4.93 | 0.00 | Q96P63 | Serpin B12 | 405 | SERPINB12 | 0 | 5 | 0 | 0 | 12.3 | 0 |

| | | | | | | | | | | | | | |
|---|---|---|---|---|---|---|---|---|---|---|---|---|---|
| 1152 | 0.00 | 0.39 | 0.00 | P09669 | Cytochrome c oxidase subunit 6C (Cytochrome c oxidase polypeptide VIc) | 75 | COX6C | 0 | 4 | 0 | 0 | 30.7 | 0 |
| 1153 | 0.00 | 4.10 | 0.00 | Q08554 | Desmocollin-1 (Cadherin family member 1) (Desmosomal glycoprotein 2/3) (DG2/DG3) | 894 | DSC1 | 0 | 4 | 0 | 0 | 6.2 | 0 |
| 1154 | 0.00 | 4.03 | 0.00 | Q5T749 | Keratinocyte proline-rich protein (hKPRP) | 579 | KPRP | 0 | 4 | 0 | 0 | 5 | 0 |
| 1155 | 0.00 | 0.13 | 0.00 | Q96FQ6 | Protein S100-A16 (Aging-associated gene 13 protein) (Protein S100-F) (S100 calcium-binding protein A16) | 103 | S100A16 | 0 | 4 | 0 | 0 | 51.5 | 0 |
| 1156 | 0.00 | 1.16 | 0.00 | K7ESP4 | Dephospho-CoA kinase domain-containing protein (Fragment) | 209 | DCAKD | 0 | 3 | 0 | 0 | 16.7 | 0 |
| 1157 | 0.00 | 1.27 | 0.00 | O95168 | NADH dehydrogenase [ubiquinone] 1 beta subcomplex subunit 4 (Complex I-B15) (CI-B15) (NADH-ubiquinone oxidoreductase B15 subunit) | 129 | NDUFB4 | 0 | 3 | 0 | 0 | 21.7 | 0 |
| 1158 | 0.00 | 4.07 | 0.00 | P04040 | Catalase (EC 1.11.1.6) | 527 | CAT | 0 | 3 | 0 | 0 | 7 | 0 |
| 1159 | 0.00 | 4.53 | 0.00 | P05089 | Arginase-1 (EC 3.5.3.1) (Liver-type arginase) (Type I arginase) | 322 | ARG1 | 0 | 3 | 0 | 0 | 9.6 | 0 |
| 1160 | 0.00 | -0.09 | 0.00 | Q6GMV3 | Putative peptidyl-tRNA hydrolase PTRHD1 (EC 3.1.1.29) (Peptidyl-tRNA hydrolase domain-containing protein 1) | 140 | PTRHD1 | 0 | 3 | 0 | 0 | 21.4 | 0 |
| 1161 | 0.00 | 0.69 | 0.00 | Q7RTV0 | PHD finger-like domain-containing protein 5A (PHD finger-like domain protein 5A) (Splicing factor 3B-associated 14 kDa protein) (SF3b14b) | 110 | PHF5A | 0 | 3 | 0 | 0 | 28.2 | 0 |

| | | | | | | | | | | | | | |
|---|---|---|---|---|---|---|---|---|---|---|---|---|---|
| 1162 | 0.00 | 0.87 | 0.00 | Q9Y2R0 | Cytochrome c oxidase assembly factor 3 homolog, mitochondrial (Coiled-coil domain-containing protein 56) (Mitochondrial translation regulation assembly intermediate of cytochrome c oxidase protein of 12 kDa) | 106 | COA3 | 0 | 3 | 0 | 0 | 29.2 | 0 |
| 1163 | 0.00 | 0.98 | 0.00 | C9JQD4 | Peptidyl-prolyl cis-trans isomerase (EC 5.2.1.8) (Fragment) | 144 | PPIH | 0 | 2 | 0 | 0 | 13.2 | 0 |
| 1164 | 0.00 | 0.20 | 0.00 | D6RAW0 | Ubiquitin-conjugating enzyme E2 D3 (Fragment) | 72 | UBE2D3 | 0 | 2 | 0 | 0 | 23.6 | 0 |
| 1165 | 0.00 | 0.27 | 0.00 | E7EQV9 | Ribosomal protein L15 (Fragment) | 174 | RPL15 | 0 | 2 | 0 | 0 | 14.9 | 0 |
| 1166 | 0.00 | 0.15 | 0.00 | E7ERH2 | S-phase kinase-associated protein 1 (Fragment) | 142 | SKP1 | 0 | 2 | 0 | 0 | 14.8 | 0 |
| 1167 | 0.00 | 1.63 | 0.00 | E9PPW7 | NADH dehydrogenase [ubiquinone] iron-sulfur protein 8, mitochondrial (Fragment) | 184 | NDUFS8 | 0 | 2 | 0 | 0 | 16.3 | 0 |
| 1168 | 0.00 | 0.75 | 0.00 | F5H702 | 39S ribosomal protein L48, mitochondrial | 113 | MRPL48 | 0 | 2 | 0 | 0 | 21.2 | 0 |
| 1169 | 0.00 | 0.43 | 0.00 | H7C0A3 | Protein ARPC4-TTLL3 (Fragment) | 167 | ARPC4-TTLL3 | 0 | 2 | 0 | 0 | 11.4 | 0 |
| 1170 | 0.00 | 0.76 | 0.00 | H7C585 | Frataxin, mitochondrial (Fragment) | 108 | FXN | 0 | 2 | 0 | 0 | 22.2 | 0 |
| 1171 | 0.00 | 1.05 | 0.00 | J3QL56 | Protein SCO1 homolog, mitochondrial | 270 | SCO1 | 0 | 2 | 0 | 0 | 10.4 | 0 |
| 1172 | 0.00 | 0.49 | 0.00 | O14907 | Tax1-binding protein 3 (Glutaminase-interacting protein 3) (Tax interaction protein 1) (TIP-1) (Tax-interacting protein 1) | 124 | TAX1BP3 | 0 | 2 | 0 | 0 | 25 | 0 |
| 1173 | 0.00 | 1.19 | 0.00 | O14949 | Cytochrome b-c1 complex subunit 8 (Complex III subunit 8) (Complex III subunit VIII) (Ubiquinol-cytochrome c reductase complex 9.5 kDa protein) (Ubiquinol-cytochrome c reductase complex ubiquinone-binding protein QP-C) | 82 | UQCRQ | 0 | 2 | 0 | 0 | 25.6 | 0 |

| | | | | | | | | | | | | | |
|---|---|---|---|---|---|---|---|---|---|---|---|---|---|
| 1174 | 0.00 | 1.27 | 0.00 | O43678 | NADH dehydrogenase [ubiquinone] 1 alpha subcomplex subunit 2 (Complex I-B8) (CI-B8) (NADH-ubiquinone oxidoreductase B8 subunit) | 99 | NDUFA2 | 0 | 2 | 0 | 0 | 30.3 | 0 |
| 1175 | 0.00 | 1.04 | 0.00 | O60830 | Mitochondrial import inner membrane translocase subunit Tim17-B | 172 | TIMM17B | 0 | 2 | 0 | 0 | 21.5 | 0 |
| 1176 | 0.00 | 0.53 | 0.00 | O75340 | Programmed cell death protein 6 (Apoptosis-linked gene 2 protein) (Probable calcium-binding protein ALG-2) | 191 | PDCD6 | 0 | 2 | 0 | 0 | 21.2 | 0 |
| 1177 | 0.00 | 1.25 | 0.00 | O95182 | NADH dehydrogenase [ubiquinone] 1 alpha subcomplex subunit 7 (Complex I-B14.5a) (CI-B14.5a) (NADH-ubiquinone oxidoreductase subunit B14.5a) | 113 | NDUFA7 | 0 | 2 | 0 | 0 | 23 | 0 |
| 1178 | 0.00 | 5.02 | 0.00 | P01040 | Cystatin-A (Cystatin-AS) (Stefin-A) [Cleaved into: Cystatin-A, N-terminally processed] | 98 | CSTA | 0 | 2 | 0 | 0 | 30.6 | 0 |
| 1179 | 0.00 | 2.08 | 0.00 | P10253 | Lysosomal alpha-glucosidase (EC 3.2.1.20) (Acid maltase) (Aglucosidase alfa) [Cleaved into: 76 kDa lysosomal alpha-glucosidase; 70 kDa lysosomal alpha-glucosidase] | 952 | GAA | 0 | 2 | 0 | 0 | 4.5 | 0 |
| 1180 | 0.00 | 1.09 | 0.00 | P14927 | Cytochrome b-c1 complex subunit 7 (Complex III subunit 7) (Complex III subunit VII) (QP-C) (Ubiquinol-cytochrome c reductase complex 14 kDa protein) | 111 | UQCRB | 0 | 2 | 0 | 0 | 21.6 | 0 |
| 1181 | 0.00 | 1.07 | 0.00 | P21912 | Succinate dehydrogenase [ubiquinone] iron-sulfur subunit, mitochondrial (EC 1.3.5.1) (Iron-sulfur subunit of complex II) (Ip) | 280 | SDHB | 0 | 2 | 0 | 0 | 7.5 | 0 |
| 1182 | 0.00 | 0.68 | 0.00 | P26373 | 60S ribosomal protein L13 (Breast basic conserved protein 1) | 211 | RPL13 | 0 | 2 | 0 | 0 | 14 | 0 |

| | | | | | | | | | | | | | |
|---|---|---|---|---|---|---|---|---|---|---|---|---|---|
| 1183 | 0.00 | 1.75 | 0.00 | P29508 | Serpin B3 (Protein T4-A) (Squamous cell carcinoma antigen 1) (SCCA-1) | 390 | SERPINB3 | 0 | 2 | 0 | 0 | 5.9 | 0 |
| 1184 | 0.00 | 0.58 | 0.00 | P53999 | Activated RNA polymerase II transcriptional coactivator p15 (Positive cofactor 4) (PC4) (SUB1 homolog) (p14) | 127 | SUB1 | 0 | 2 | 0 | 0 | 11 | 0 |
| 1185 | 0.00 | 0.10 | 0.00 | P55769 | NHP2-like protein 1 (High mobility group-like nuclear protein 2 homolog 1) (OTK27) (SNU13 homolog) (hSNU13) (U4/U6.U5 tri-snRNP 15.5 kDa protein) [Cleaved into: NHP2-like protein 1, N-terminally processed] | 128 | NHP2L1 | 0 | 2 | 0 | 0 | 18 | 0 |
| 1186 | 0.00 | 1.11 | 0.00 | P61769 | Beta-2-microglobulin [Cleaved into: Beta-2-microglobulin form pI 5.3] | 119 | B2M | 0 | 2 | 0 | 0 | 26.9 | 0 |
| 1187 | 0.00 | 1.04 | 0.00 | P62304 | Small nuclear ribonucleoprotein E (snRNP-E) (Sm protein E) (Sm-E) (SmE) | 92 | SNRPE | 0 | 2 | 0 | 0 | 25 | 0 |
| 1188 | 0.00 | -0.13 | 0.00 | P63167 | Dynein light chain 1, cytoplasmic (8 kDa dynein light chain) (DLC8) (Dynein light chain LC8-type 1) (Protein inhibitor of neuronal nitric oxide synthase) (PIN) | 89 | DYNLL1 | 0 | 2 | 0 | 0 | 20.2 | 0 |
| 1189 | 0.00 | 0.81 | 0.00 | P63173 | 60S ribosomal protein L38 | 70 | RPL38 | 0 | 2 | 0 | 0 | 28.6 | 0 |
| 1190 | 0.00 | 1.73 | 0.00 | P82664 | 28S ribosomal protein S10, mitochondrial (MRP-S10) (S10mt) | 201 | MRPS10 | 0 | 2 | 0 | 0 | 14.4 | 0 |
| 1191 | 0.00 | 0.74 | 0.00 | P82979 | SAP domain-containing ribonucleoprotein (Cytokine-induced protein of 29 kDa) (Nuclear protein Hcc-1) (Proliferation-associated cytokine-inducible protein CIP29) | 210 | SARNP | 0 | 2 | 0 | 0 | 9 | 0 |
| 1192 | 0.00 | -0.07 | 0.00 | Q13247 | Serine/arginine-rich splicing factor 6 (Pre-mRNA-splicing factor SRP55) (Splicing factor, arginine/serine-rich 6) | 344 | SRSF6 | 0 | 2 | 0 | 0 | 6.3 | 0 |

| | | | | | | | | | | | | | |
|---|---|---|---|---|---|---|---|---|---|---|---|---|---|
| 1193 | 0.00 | 1.99 | 0.00 | Q15125 | 3-beta-hydroxysteroid-Delta(8),Delta(7)-isomerase (EC 5.3.3.5) (Cholestenol Delta-isomerase) (Delta(8)-Delta(7) sterol isomerase) (D8-D7 sterol isomerase) (Emopamil-binding protein) | 230 | EBP | 0 | 2 | 0 | 0 | 12.2 | 0 |
| 1194 | 0.00 | 0.06 | 0.00 | Q3ZAQ7 | Vacuolar ATPase assembly integral membrane protein VMA21 (Myopathy with excessive autophagy protein) | 101 | VMA21 | 0 | 2 | 0 | 0 | 21.8 | 0 |
| 1195 | 0.00 | 1.30 | 0.00 | Q6P1X6 | UPF0598 protein C8orf82 | 216 | C8orf82 | 0 | 2 | 0 | 0 | 13.5 | 0 |
| 1196 | 0.00 | 1.37 | 0.00 | Q7Z7K0 | COX assembly mitochondrial protein homolog (Cmc1p) | 106 | CMC1 | 0 | 2 | 0 | 0 | 18.9 | 0 |
| 1197 | 0.00 | 0.17 | 0.00 | Q8N983 | 39S ribosomal protein L43, mitochondrial (L43mt) (MRP-L43) (Mitochondrial ribosomal protein bMRP36a) | 215 | MRPL43 | 0 | 2 | 0 | 0 | 12.6 | 0 |
| 1198 | 0.00 | 0.92 | 0.00 | Q8NI27 | THO complex subunit 2 (Tho2) (hTREX120) | 1593 | THOC2 | 0 | 2 | 0 | 0 | 1.4 | 0 |
| 1199 | 0.00 | 0.21 | 0.00 | Q8WVC2 | 40S ribosomal protein S21 | 81 | RPS21 | 0 | 2 | 0 | 0 | 35.8 | 0 |
| 1200 | 0.00 | 1.77 | 0.00 | Q9HB66 | Alternative protein MKKS (McKusick-Kaufman syndrome, isoform CRA_a) (McKusick-Kaufman/Bardet-Biedl syndromes putative chaperonin) (PNAS-117) | 63 | MKKS | 0 | 2 | 0 | 0 | 41.3 | 0 |
| 1201 | 0.00 | 0.69 | 0.00 | Q9NP97 | Dynein light chain roadblock-type 1 (Bithoraxoid-like protein) (BLP) (Dynein light chain 2A, cytoplasmic) (Dynein-associated protein Km23) (Roadblock domain-containing protein 1) | 96 | DYNLRB1 | 0 | 2 | 0 | 0 | 25 | 0 |

| | | | | | | | | | | | | | |
|---|---|---|---|---|---|---|---|---|---|---|---|---|---|
| 1202 | 0.00 | 0.86 | 0.00 | Q9Y333 | U6 snRNA-associated Sm-like protein LSm2 (Protein G7b) (Small nuclear ribonuclear protein D homolog) (snRNP core Sm-like protein Sm-x5) | 95 | LSM2 | 0 | 2 | 0 | 0 | 31.6 | 0 |
| 1203 | 0.00 | 1.06 | 0.00 | Q9Y3B7 | 39S ribosomal protein L11, mitochondrial (L11mt) (MRP-L11) | 192 | MRPL11 | 0 | 2 | 0 | 0 | 17.5 | 0 |
| 1204 | 0.00 | 0.83 | 0.00 | Q9Y3D6 | Mitochondrial fission 1 protein (FIS1 homolog) (hFis1) (Tetratricopeptide repeat protein 11) (TPR repeat protein 11) | 152 | FIS1 | 0 | 2 | 0 | 0 | 15.8 | 0 |
| 1205 | 0.00 | 0.35 | 0.00 | Q9Y5J7 | Mitochondrial import inner membrane translocase subunit Tim9 | 89 | TIMM9 | 0 | 2 | 0 | 0 | 29.2 | 0 |
| 1206 | 0.00 | 2.12 | 0.00 | A0A087 WUD3 | Oligosaccharyltransferase complex subunit OSTC | 83 | OSTC | 0 | 1 | 0 | 0 | 14.5 | 0 |
| 1207 | 0.00 | 2.31 | 0.00 | A0A087 WVT9 | Nucleoside diphosphate kinase (EC 2.7.4.6) | 153 | NME4 | 0 | 1 | 0 | 0 | 7.8 | 0 |
| 1208 | 0.00 | 0.22 | 0.00 | A0A087 WY88 | Protein jagunal homolog 1 | 181 | JAGN1 | 0 | 1 | 0 | 0 | 6.6 | 0 |
| 1209 | 0.00 | 6.49 | 0.00 | A0A087 WYF5 | Salivary acidic proline-rich phosphoprotein 1/2 (Fragment) | 140 | PRH1 | 0 | 1 | 0 | 0 | 12.1 | 0 |
| 1210 | 0.00 | -0.20 | 0.00 | B5MC59 | Replication protein A 14 kDa subunit (Replication protein A3, 14kDa, isoform CRA_a) | 82 | RPA3 | 0 | 1 | 0 | 0 | 14.6 | 0 |
| 1211 | 0.00 | 1.43 | 0.00 | B8ZZV5 | 39S ribosomal protein L30, mitochondrial | 101 | MRPL30 | 0 | 1 | 0 | 0 | 10.9 | 0 |
| 1212 | 0.00 | 0.32 | 0.00 | C9IZG4 | Protein CutA | 135 | CUTA | 0 | 1 | 0 | 0 | 13.3 | 0 |
| 1213 | 0.00 | 1.97 | 0.00 | C9JAW5 | HIG1 domain family member 1A, mitochondrial | 83 | HIGD1A | 0 | 1 | 0 | 0 | 21.7 | 0 |
| 1214 | 0.00 | 2.68 | 0.00 | C9JEV0 | Zinc-alpha-2-glycoprotein | 227 | AZGP1 | 0 | 1 | 0 | 0 | 5.3 | 0 |
| 1215 | 0.00 | 0.99 | 0.00 | C9JKQ2 | NADH dehydrogenase [ubiquinone] 1 beta subcomplex subunit 3 (Fragment) | 65 | NDUFB3 | 0 | 1 | 0 | 0 | 16.9 | 0 |

| | | | | | | | | | | | | | |
|---|---|---|---|---|---|---|---|---|---|---|---|---|---|
| 1216 | 0.00 | 0.18 | 0.00 | C9JL85 | Myotrophin | 52 | MTPN | 0 | 1 | 0 | 0 | 32.7 | 0 |
| 1217 | 0.00 | 0.90 | 0.00 | C9JY28 | LYR motif-containing protein 4 | 70 | LYRM4 | 0 | 1 | 0 | 0 | 20 | 0 |
| 1218 | 0.00 | 0.38 | 0.00 | C9JYN0 | Synaptophysin-like protein 1 | 224 | SYPL1 | 0 | 1 | 0 | 0 | 4.9 | 0 |
| 1219 | 0.00 | 5.73 | 0.00 | D3DRR9 | Chromosome 10 open reading frame 47, isoform CRA_b (Proline and serine-rich protein 2) | 239 | C10orf47 | 0 | 1 | 0 | 0 | 5.4 | 0 |
| 1220 | 0.00 | 1.32 | 0.00 | D6R9Z7 | Cytochrome c oxidase subunit 7C, mitochondrial | 56 | COX7C | 0 | 1 | 0 | 0 | 16.1 | 0 |
| 1221 | 0.00 | 1.32 | 0.00 | E5RGY0 | Derlin-1 | 151 | DERL1 | 0 | 1 | 0 | 0 | 7.3 | 0 |
| 1222 | 0.00 | 2.21 | 0.00 | E7EWF7 | Uncharacterized protein (Fragment) | 191 | | 0 | 1 | 0 | 0 | 13.1 | 0 |
| 1223 | 0.00 | 2.20 | 0.00 | E9PKY5 | Peptidyl-prolyl cis-trans isomerase (EC 5.2.1.8) (Fragment) | 213 | PPIE | 0 | 1 | 0 | 0 | 4.2 | 0 |
| 1224 | 0.00 | -0.04 | 0.00 | E9PNW4 | CD59 glycoprotein | 108 | CD59 | 0 | 1 | 0 | 0 | 7.4 | 0 |
| 1225 | 0.00 | -0.58 | 0.00 | E9PPQ4 | Ferritin (Fragment) | 58 | FTH1 | 0 | 1 | 0 | 0 | 12.1 | 0 |
| 1226 | 0.00 | 1.45 | 0.00 | F5H169 | 26S proteasome non-ATPase regulatory subunit 9 | 128 | PSMD9 | 0 | 1 | 0 | 0 | 9.4 | 0 |
| 1227 | 0.00 | -0.11 | 0.00 | F5H1S8 | Malectin (Fragment) | 146 | MLEC | 0 | 1 | 0 | 0 | 8.2 | 0 |
| 1228 | 0.00 | 1.60 | 0.00 | F6WST4 | ORM1-like protein 1 (Fragment) | 108 | ORMDL1 | 0 | 1 | 0 | 0 | 13.9 | 0 |
| 1229 | 0.00 | 0.17 | 0.00 | F8VSA6 | NEDD8 | 50 | NEDD8 | 0 | 1 | 0 | 0 | 22 | 0 |
| 1230 | 0.00 | 2.57 | 0.00 | F8VV32 | Lysozyme (EC 3.2.1.17) | 104 | LYZ | 0 | 1 | 0 | 0 | 11.5 | 0 |
| 1231 | 0.00 | 0.85 | 0.00 | F8WEX5 | Sulfatase-modifying factor 2 | 165 | SUMF2 | 0 | 1 | 0 | 0 | 6.1 | 0 |
| 1232 | 0.00 | 4.84 | 0.00 | G8JLG2 | Corneodesmosin | 529 | CDSN | 0 | 1 | 0 | 0 | 3.4 | 0 |
| 1233 | 0.00 | -0.45 | 0.00 | G8JLQ3 | Biogenesis of lysosome-related organelles complex 1 subunit 1 | 75 | BLOC1S1 | 0 | 1 | 0 | 0 | 14.7 | 0 |
| 1234 | 0.00 | 0.69 | 0.00 | H0Y993 | Protein DEK (Fragment) | 156 | DEK | 0 | 1 | 0 | 0 | 8.3 | 0 |
| 1235 | 0.00 | 1.42 | 0.00 | H0YDP7 | 39S ribosomal protein L49, mitochondrial (Fragment) | 127 | MRPL49 | 0 | 1 | 0 | 0 | 11 | 0 |

| | | | | | | | | | | | | | |
|---|---|---|---|---|---|---|---|---|---|---|---|---|---|
| 1236 | 0.00 | -0.02 | 0.00 | H0YER1 | Remodeling and spacing factor 1 (Fragment) | 702 | RSF1 | 0 | 1 | 0 | 0 | 1.7 | 0 |
| 1237 | 0.00 | 1.58 | 0.00 | H0YF90 | Cation-dependent mannose-6-phosphate receptor (Fragment) | 148 | M6PR | 0 | 1 | 0 | 0 | 18.2 | 0 |
| 1238 | 0.00 | 0.82 | 0.00 | H3BSA6 | dCTP pyrophosphatase 1 | 71 | DCTPP1 | 0 | 1 | 0 | 0 | 14.1 | 0 |
| 1239 | 0.00 | -0.12 | 0.00 | H7BY91 | 60S ribosomal protein L36a | 70 | RPL36A | 0 | 1 | 0 | 0 | 14.3 | 0 |
| 1240 | 0.00 | 1.56 | 0.00 | J3KS15 | Peptidyl-tRNA hydrolase ICT1, mitochondrial (Fragment) | 192 | ICT1 | 0 | 1 | 0 | 0 | 7.3 | 0 |
| 1241 | 0.00 | 0.74 | 0.00 | J3KTI5 | Receptor tyrosine-protein kinase erbB-2 (Fragment) | 251 | ERBB2 | 0 | 1 | 0 | 0 | 4.8 | 0 |
| 1242 | 0.00 | -9.65 | 0.00 | J3KTP2 | WD repeat-containing protein WRAP73 (Fragment) | 200 | WRAP73 | 0 | 1 | 0 | 0 | 4.5 | 0 |
| 1243 | 0.00 | -0.19 | 0.00 | J3QR72 | Bromodomain-containing protein 2 | 162 | BRD2 | 0 | 1 | 0 | 0 | 8 | 0 |
| 1244 | 0.00 | -0.97 | 0.00 | K4DIB9 | Cysteine-rich protein 1 (Fragment) | 46 | CRIP1 | 0 | 1 | 0 | 0 | 19.6 | 0 |
| 1245 | 0.00 | -0.04 | 0.00 | K7EM09 | Transmembrane protein 205 (Fragment) | 120 | TMEM205 | 0 | 1 | 0 | 0 | 10.8 | 0 |
| 1246 | 0.00 | -0.07 | 0.00 | K7ENF1 | Peroxisomal acyl-coenzyme A oxidase 1 (Fragment) | 132 | ACOX1 | 0 | 1 | 0 | 0 | 10.6 | 0 |
| 1247 | 0.00 | 1.70 | 0.00 | K7EQ77 | NADH dehydrogenase [ubiquinone] 1 alpha subcomplex subunit 11 | 120 | NDUFA11 | 0 | 1 | 0 | 0 | 19.2 | 0 |
| 1248 | 0.00 | 2.27 | 0.00 | K7EQQ1 | Cleft lip and palate transmembrane protein 1 (Fragment) | 171 | CLPTM1 | 0 | 1 | 0 | 0 | 8.8 | 0 |
| 1249 | 0.00 | 0.81 | 0.00 | L0R6Q1 | Alternative protein SLC35A4 (Probable UDP-sugar transporter protein SLC35A4) | 103 | SLC35A4 | 0 | 1 | 0 | 0 | 7.8 | 0 |
| 1250 | 0.00 | 0.29 | 0.00 | M0QXH0 | Thioredoxin, mitochondrial | 64 | TXN2 | 0 | 1 | 0 | 0 | 23.4 | 0 |
| 1251 | 0.00 | -2.71 | 0.00 | M0R050 | Exosome complex component RRP46 | 197 | EXOSC5 | 0 | 1 | 0 | 0 | 6.1 | 0 |

| | | | | | | | | | | | | | |
|---|---|---|---|---|---|---|---|---|---|---|---|---|---|
| 1252 | 0.00 | 0.52 | 0.00 | M0R203 | Heterogeneous nuclear ribonucleoprotein U-like protein 1 (Fragment) | 81 | HNRNPUL1 | 0 | 1 | 0 | 0 | 13.6 | 0 |
| 1253 | 0.00 | 0.08 | 0.00 | O15155 | BET1 homolog (hBET1) (Golgi vesicular membrane-trafficking protein p18) | 118 | BET1 | 0 | 1 | 0 | 0 | 13.3 | 0 |
| 1254 | 0.00 | 1.39 | 0.00 | O43402 | ER membrane protein complex subunit 8 (Neighbor of COX4) (Protein FAM158B) | 210 | EMC8 | 0 | 1 | 0 | 0 | 5.7 | 0 |
| 1255 | 0.00 | -0.58 | 0.00 | O43598 | 2'-deoxynucleoside 5'-phosphate N-hydrolase 1 (EC 3.2.2.-) (c-Myc-responsive protein RCL) | 174 | DNPH1 | 0 | 1 | 0 | 0 | 11.5 | 0 |
| 1256 | 0.00 | -0.30 | 0.00 | O60763 | General vesicular transport factor p115 (Protein USO1 homolog) (Transcytosis-associated protein) (TAP) (Vesicle-docking protein) | 962 | USO1 | 0 | 1 | 0 | 0 | 1.2 | 0 |
| 1257 | 0.00 | 1.42 | 0.00 | O60783 | 28S ribosomal protein S14, mitochondrial (MRP-S14) (S14mt) | 128 | MRPS14 | 0 | 1 | 0 | 0 | 11.7 | 0 |
| 1258 | 0.00 | 1.02 | 0.00 | O75380 | NADH dehydrogenase [ubiquinone] iron-sulfur protein 6, mitochondrial (Complex I-13kD-A) (CI-13kD-A) (NADH-ubiquinone oxidoreductase 13 kDa-A subunit) | 124 | NDUFS6 | 0 | 1 | 0 | 0 | 12.1 | 0 |
| 1259 | 0.00 | -0.23 | 0.00 | O75531 | Barrier-to-autointegration factor (Breakpoint cluster region protein 1) [Cleaved into: Barrier-to-autointegration factor, N-terminally processed] | 89 | BANF1 | 0 | 1 | 0 | 0 | 13.5 | 0 |
| 1260 | 0.00 | 0.75 | 0.00 | O75934 | Pre-mRNA-splicing factor SPF27 (Breast carcinoma-amplified sequence 2) (DNA amplified in mammary carcinoma 1 protein) (Spliceosome-associated protein SPF 27) | 225 | BCAS2 | 0 | 1 | 0 | 0 | 5.8 | 0 |

| | | | | | | | | | | | | | |
|---|---|---|---|---|---|---|---|---|---|---|---|---|---|
| 1261 | 0.00 | 1.57 | 0.00 | O95298 | NADH dehydrogenase [ubiquinone] 1 subunit C2 (Complex I-B14.5b) (CI-B14.5b) (Human lung cancer oncogene 1 protein) (HLC-1) (NADH-ubiquinone oxidoreductase subunit B14.5b) | 119 | NDUFC2 | 0 | 1 | 0 | 0 | 12.5 | 0 |
| 1262 | 0.00 | 0.50 | 0.00 | O95969 | Secretoglobin family 1D member 2 (Lipophilin-B) | 90 | SCGB1D2 | 0 | 1 | 0 | 0 | 10 | 0 |
| 1263 | 0.00 | 0.41 | 0.00 | P00846 | ATP synthase subunit a (F-ATPase protein 6) | 226 | MT-ATP6 | 0 | 1 | 0 | 0 | 4.4 | 0 |
| 1264 | 0.00 | 2.42 | 0.00 | P03928 | ATP synthase protein 8 (A6L) (F-ATPase subunit 8) | 68 | MT-ATP8 | 0 | 1 | 0 | 0 | 13.2 | 0 |
| 1265 | 0.00 | 1.75 | 0.00 | P05386 | 60S acidic ribosomal protein P1 | 114 | RPLP1 | 0 | 1 | 0 | 0 | 14 | 0 |
| 1266 | 0.00 | 0.31 | 0.00 | P07311 | Acylphosphatase-1 (EC 3.6.1.7) (Acylphosphatase, erythrocyte isozyme) (Acylphosphatase, organ-common type isozyme) (Acylphosphate phosphohydrolase 1) | 99 | ACYP1 | 0 | 1 | 0 | 0 | 13.1 | 0 |
| 1267 | 0.00 | 1.10 | 0.00 | P09234 | U1 small nuclear ribonucleoprotein C (U1 snRNP C) (U1-C) (U1C) | 159 | SNRPC | 0 | 1 | 0 | 0 | 11.3 | 0 |
| 1268 | 0.00 | 1.70 | 0.00 | P10301 | Ras-related protein R-Ras (p23) | 218 | RRAS | 0 | 1 | 0 | 0 | 5 | 0 |
| 1269 | 0.00 | 7.51 | 0.00 | P11532 | Dystrophin | 3685 | DMD | 0 | 1 | 0 | 0 | 0.5 | 0 |
| 1270 | 0.00 | 1.11 | 0.00 | P14406 | Cytochrome c oxidase subunit 7A2, mitochondrial (Cytochrome c oxidase subunit VIIa-liver/heart) (Cytochrome c oxidase subunit VIIa-L) (Cytochrome c oxidase subunit VIIaL) | 83 | COX7A2 | 0 | 1 | 0 | 0 | 15.7 | 0 |
| 1271 | 0.00 | 1.26 | 0.00 | P14854 | Cytochrome c oxidase subunit 6B1 (Cytochrome c oxidase subunit VIb isoform 1) (COX VIb-1) | 86 | COX6B1 | 0 | 1 | 0 | 0 | 20.9 | 0 |

| | | | | | | | | | | | | | |
|---|---|---|---|---|---|---|---|---|---|---|---|---|---|
| 1272 | 0.00 | -0.09 | 0.00 | P25815 | Protein S100-P (Migration-inducing gene 9 protein) (MIG9) (Protein S100-E) (S100 calcium-binding protein P) | 95 | S100P | 0 | 1 | 0 | 0 | 13.7 | 0 |
| 1273 | 0.00 | 0.70 | 0.00 | P28288 | ATP-binding cassette sub-family D member 3 (70 kDa peroxisomal membrane protein) (PMP70) | 659 | ABCD3 | 0 | 1 | 0 | 0 | 2.2 | 0 |
| 1274 | 0.00 | 0.13 | 0.00 | P36543 | V-type proton ATPase subunit E 1 (V-ATPase subunit E 1) (V-ATPase 31 kDa subunit) (p31) (Vacuolar proton pump subunit E 1) | 226 | ATP6V1E1 | 0 | 1 | 0 | 0 | 5.1 | 0 |
| 1275 | 0.00 | 4.09 | 0.00 | P47929 | Galectin-7 (Gal-7) (HKL-14) (PI7) (p53-induced gene 1 protein) | 136 | LGALS7 | 0 | 1 | 0 | 0 | 8.1 | 0 |
| 1276 | 0.00 | 0.79 | 0.00 | P51151 | Ras-related protein Rab-9A | 201 | RAB9A | 0 | 1 | 0 | 0 | 7 | 0 |
| 1277 | 0.00 | 0.43 | 0.00 | P56385 | ATP synthase subunit e, mitochondrial (ATPase subunit e) | 69 | ATP5I | 0 | 1 | 0 | 0 | 14.5 | 0 |
| 1278 | 0.00 | -0.15 | 0.00 | P61254 | 60S ribosomal protein L26 | 145 | RPL26 | 0 | 1 | 0 | 0 | 5.5 | 0 |
| 1279 | 0.00 | 0.43 | 0.00 | P61457 | Pterin-4-alpha-carbinolamine dehydratase (PHS) (EC 4.2.1.96) (4-alpha-hydroxy-tetrahydropterin dehydratase) (Dimerization cofactor of hepatocyte nuclear factor 1-alpha) (DCoH) (Dimerization cofactor of HNF1) (Phenylalanine hydroxylase-stimulating protein) (Pterin carbinolamine dehydratase) (PCD) | 104 | PCBD1 | 0 | 1 | 0 | 0 | 12.5 | 0 |
| 1280 | 0.00 | 2.05 | 0.00 | P62306 | Small nuclear ribonucleoprotein F (snRNP-F) (Sm protein F) (Sm-F) (SmF) | 86 | SNRPF | 0 | 1 | 0 | 0 | 8.1 | 0 |
| 1281 | 0.00 | 0.05 | 0.00 | P62312 | U6 snRNA-associated Sm-like protein LSm6 | 80 | LSM6 | 0 | 1 | 0 | 0 | 13.8 | 0 |
| 1282 | 0.00 | 0.40 | 0.00 | P63218 | Guanine nucleotide-binding protein G(I)/G(S)/G(O) subunit gamma-5 | 68 | GNG5 | 0 | 1 | 0 | 0 | 13.2 | 0 |

| | | | | | | | | | | | | | |
|---|---|---|---|---|---|---|---|---|---|---|---|---|---|
| 1283 | 0.00 | 2.08 | 0.00 | P80723 | Brain acid soluble protein 1 (22 kDa neuronal tissue-enriched acidic protein) (Neuronal axonal membrane protein NAP-22) | 227 | BASP1 | 0 | 1 | 0 | 0 | 6.2 | 0 |
| 1284 | 0.00 | 4.91 | 0.00 | Q08188 | Protein-glutamine gamma-glutamyltransferase E (EC 2.3.2.13) (Transglutaminase E) (TG(E)) (TGE) (TGase E) (Transglutaminase-3) (TGase-3) [Cleaved into: Protein-glutamine gamma-glutamyltransferase E 50 kDa catalytic chain; Protein-glutamine gamma-glutamyltransferase E 27 kDa non-catalytic chain] | 693 | TGM3 | 0 | 1 | 0 | 0 | 1.3 | 0 |
| 1285 | 0.00 | 5.57 | 0.00 | Q12873 | Chromodomain-helicase-DNA-binding protein 3 (CHD-3) (EC 3.6.4.12) (ATP-dependent helicase CHD3) (Mi-2 autoantigen 240 kDa protein) (Mi2-alpha) (Zinc finger helicase) (hZFH) | 2000 | CHD3 | 0 | 1 | 0 | 0 | 0.5 | 0 |
| 1286 | 0.00 | 0.01 | 0.00 | Q14061 | Cytochrome c oxidase copper chaperone | 63 | COX17 | 0 | 1 | 0 | 0 | 25.4 | 0 |
| 1287 | 0.00 | 1.22 | 0.00 | Q16864 | V-type proton ATPase subunit F (V-ATPase subunit F) (V-ATPase 14 kDa subunit) (Vacuolar proton pump subunit F) | 119 | ATP6V1F | 0 | 1 | 0 | 0 | 16 | 0 |
| 1288 | 0.00 | 0.51 | 0.00 | Q5HYK3 | 2-methoxy-6-polyprenyl-1,4-benzoquinol methylase, mitochondrial (EC 2.1.1.201) (Ubiquinone biosynthesis methyltransferase COQ5) | 327 | COQ5 | 0 | 1 | 0 | 0 | 4.1 | 0 |
| 1289 | 0.00 | 1.02 | 0.00 | Q5JTJ3 | Cytochrome c oxidase assembly factor 6 homolog | 125 | COA6 | 0 | 1 | 0 | 0 | 17.7 | 0 |
| 1290 | 0.00 | 0.64 | 0.00 | Q5QPM9 | Proteasome inhibitor PI31 subunit (Fragment) | 183 | PSMF1 | 0 | 1 | 0 | 0 | 9.3 | 0 |

| | | | | | | | | | | | | | |
|---|---|---|---|---|---|---|---|---|---|---|---|---|---|
| 1291 | 0.00 | 1.51 | 0.00 | Q5RI15 | Cytochrome c oxidase protein 20 homolog | 118 | COX20 | 0 | 1 | 0 | 0 | 11.9 | 0 |
| 1292 | 0.00 | 1.91 | 0.00 | Q5RKV6 | Exosome complex component MTR3 (Exosome component 6) (mRNA transport regulator 3 homolog) (hMtr3) (p11) | 272 | EXOSC6 | 0 | 1 | 0 | 0 | 5.9 | 0 |
| 1293 | 0.00 | -1.33 | 0.00 | Q5SXM8 | DNL-type zinc finger protein (Hsp70-escort protein 1) (HEP1) (mtHsp70-escort protein) | 178 | DNLZ | 0 | 1 | 0 | 0 | 5.1 | 0 |
| 1294 | 0.00 | 0.34 | 0.00 | Q5T123 | SH3 domain-binding glutamic acid-rich-like protein 3 | 88 | SH3BGRL3 | 0 | 1 | 0 | 0 | 17 | 0 |
| 1295 | 0.00 | 0.38 | 0.00 | Q5TGZ0 | MICOS complex subunit MIC10 (Mitochondrial inner membrane organizing system protein 1) | 78 | MINOS1 | 0 | 1 | 0 | 0 | 33.3 | 0 |
| 1296 | 0.00 | 5.34 | 0.00 | Q5VTQ0 | Tetratricopeptide repeat protein 39B (TPR repeat protein 39B) | 682 | TTC39B | 0 | 1 | 0 | 0 | 6 | 0 |
| 1297 | 0.00 | 0.56 | 0.00 | Q5VU10 | Ribonuclease P protein subunit p30 (Fragment) | 212 | RPP30 | 0 | 1 | 0 | 0 | 4.7 | 0 |
| 1298 | 0.00 | 6.60 | 0.00 | Q5VXJ5 | Synaptonemal complex protein 1 (Fragment) | 792 | SYCP1 | 0 | 1 | 0 | 0 | 1.6 | 0 |
| 1299 | 0.00 | 0.62 | 0.00 | Q6IAK0 | SELT protein (Selenoprotein T) (cDNA FLJ90525 fis, clone NT2RP4001001, highly similar to Selenoprotein T) | 137 | SELT | 0 | 1 | 0 | 0 | 9.5 | 0 |
| 1300 | 0.00 | 3.80 | 0.00 | Q6UWP8 | Suprabasin | 590 | SBSN | 0 | 1 | 0 | 0 | 9.2 | 0 |
| 1301 | 0.00 | 0.27 | 0.00 | Q7Z5G4 | acety | 137 | GOLGA7 | 0 | 1 | 0 | 0 | 8.2 | 0 |
| 1302 | 0.00 | -0.15 | 0.00 | Q7Z7H5 | Transmembrane emp24 domain-containing protein 4 (Endoplasmic reticulum stress-response protein 25) (ERS25) (GMP25iso) (Putative NF-kappa-B-activating protein 156) (p24 family protein alpha-3) (p24alpha3) | 227 | TMED4 | 0 | 1 | 0 | 0 | 5.9 | 0 |

| | | | | | | | | | | | | | |
|---|---|---|---|---|---|---|---|---|---|---|---|---|---|
| 1303 | 0.00 | 2.08 | 0.00 | Q7Z7H8 | 39S ribosomal protein L10, mitochondrial (L10mt) (MRP-L10) (39S ribosomal protein L8, mitochondrial) (L8mt) (MRP-L8) | 261 | MRPL10 | 0 | 1 | 0 | 0 | 7.3 | 0 |
| 1304 | 0.00 | 0.81 | 0.00 | Q86YN1 | Dolichyldiphosphatase 1 (EC 3.6.1.43) (Dolichyl pyrophosphate phosphatase 1) | 238 | DOLPP1 | 0 | 1 | 0 | 0 | 5.5 | 0 |
| 1305 | 0.00 | 1.46 | 0.00 | Q8N4V1 | Membrane magnesium transporter 1 (ER membrane protein complex subunit 5) (Transmembrane protein 32) | 131 | MMGT1 | 0 | 1 | 0 | 0 | 18.3 | 0 |
| 1306 | 0.00 | -0.32 | 0.00 | Q8WVF3 | RAB28 protein (Ras-related protein Rab-28) | 95 | RAB28 | 0 | 1 | 0 | 0 | 11.6 | 0 |
| 1307 | 0.00 | 1.53 | 0.00 | Q8WW12 | PEST proteolytic signal-containing nuclear protein (PCNP) (PEST-containing nuclear protein) | 178 | PCNP | 0 | 1 | 0 | 0 | 11.5 | 0 |
| 1308 | 0.00 | 3.57 | 0.00 | Q8WY22 | BRI3-binding protein (I3-binding protein) (Cervical cancer 1 proto-oncogene-binding protein KG19) (HCCRBP-1) | 251 | BRI3BP | 0 | 1 | 0 | 0 | 6.8 | 0 |
| 1309 | 0.00 | -0.98 | 0.00 | Q969X5 | Endoplasmic reticulum-Golgi intermediate compartment protein 1 (ER-Golgi intermediate compartment 32 kDa protein) (ERGIC-32) | 290 | ERGIC1 | 0 | 1 | 0 | 0 | 5.1 | 0 |
| 1310 | 0.00 | 1.22 | 0.00 | Q96DA6 | Mitochondrial import inner membrane translocase subunit TIM14 (DnaJ homolog subfamily C member 19) | 116 | DNAJC19 | 0 | 1 | 0 | 0 | 15.4 | 0 |
| 1311 | 0.00 | 0.55 | 0.00 | Q96EL3 | 39S ribosomal protein L53, mitochondrial (L53mt) (MRP-L53) | 112 | MRPL53 | 0 | 1 | 0 | 0 | 8.9 | 0 |
| 1312 | 0.00 | 1.01 | 0.00 | Q96EX1 | Small integral membrane protein 12 | 92 | SMIM12 | 0 | 1 | 0 | 0 | 9.8 | 0 |
| 1313 | 0.00 | 1.63 | 0.00 | Q96IX5 | Up-regulated during skeletal muscle growth protein 5 (Diabetes-associated protein in insulin-sensitive tissues) (HCV F-transactivated protein 2) | 58 | USMG5 | 0 | 1 | 0 | 0 | 25.9 | 0 |

| | | | | | | | | | | | | | |
|---|---|---|---|---|---|---|---|---|---|---|---|---|---|
| 1314 | 0.00 | 1.11 | 0.00 | Q96PD2 | Discoidin, CUB and LCCL domain-containing protein 2 (CUB, LCCL and coagulation factor V/VIII-homology domains protein 1) (Endothelial and smooth muscle cell-derived neuropilin-like protein) | 775 | DCBLD2 | 0 | 1 | 0 | 0 | 1.3 | 0 |
| 1315 | 0.00 | 5.08 | 0.00 | Q99595 | Mitochondrial import inner membrane translocase subunit Tim17-A (Inner membrane preprotein translocase Tim17a) | 171 | TIMM17A | 0 | 1 | 0 | 0 | 19.3 | 0 |
| 1316 | 0.00 | 0.31 | 0.00 | Q9BQC6 | Ribosomal protein 63, mitochondrial (hMRP63) (Mitochondrial ribosomal protein 63) (Mitochondrial ribosomal protein L57) | 102 | MRPL57 | 0 | 1 | 0 | 0 | 12.7 | 0 |
| 1317 | 0.00 | -0.29 | 0.00 | Q9BRF8 | Serine/threonine-protein phosphatase CPPED1 (EC 3.1.3.16) (Calcineurin-like phosphoesterase domain-containing protein 1) (Complete S-transactivated protein 1) | 314 | CPPED1 | 0 | 1 | 0 | 0 | 9 | 0 |
| 1318 | 0.00 | 0.45 | 0.00 | Q9BWJ5 | Splicing factor 3B subunit 5 (SF3b5) (Pre-mRNA-splicing factor SF3b 10 kDa subunit) | 86 | SF3B5 | 0 | 1 | 0 | 0 | 12.8 | 0 |
| 1319 | 0.00 | 3.54 | 0.00 | Q9BXW7 | Cat eye syndrome critical region protein 5 | 423 | CECR5 | 0 | 1 | 0 | 0 | 3.6 | 0 |
| 1320 | 0.00 | 0.55 | 0.00 | Q9BZG1 | Ras-related protein Rab-34 (Ras-related protein Rab-39) (Ras-related protein Rah) | 259 | RAB34 | 0 | 1 | 0 | 0 | 5.9 | 0 |
| 1321 | 0.00 | -0.16 | 0.00 | Q9H3Z4 | DnaJ homolog subfamily C member 5 (Cysteine string protein) (CSP) | 198 | DNAJC5 | 0 | 1 | 0 | 0 | 8.4 | 0 |
| 1322 | 0.00 | 6.18 | 0.00 | Q9H7L9 | Sin3 histone deacetylase corepressor complex component SDS3 (45 kDa Sin3-associated polypeptide) (Suppressor of defective silencing 3 protein homolog) | 328 | SUDS3 | 0 | 1 | 0 | 0 | 5.8 | 0 |

| | | | | | | | | | | | | | |
|---|---|---|---|---|---|---|---|---|---|---|---|---|---|
| 1323 | 0.00 | 1.60 | 0.00 | Q9HD45 | Transmembrane 9 superfamily member 3 (EP70-P-iso) (SM-11044-binding protein) | 589 | TM9SF3 | 0 | 1 | 0 | 0 | 1.9 | 0 |
| 1324 | 0.00 | 0.70 | 0.00 | Q9NQ50 | 39S ribosomal protein L40, mitochondrial (L40mt) (MRP-L40) (Nuclear localization signal-containing protein deleted in velocardiofacial syndrome) (Up-regulated in metastasis) | 206 | MRPL40 | 0 | 1 | 0 | 0 | 4.4 | 0 |
| 1325 | 0.00 | 1.13 | 0.00 | Q9NTX5 | Ethylmalonyl-CoA decarboxylase (EC 4.1.1.94) (Enoyl-CoA hydratase domain-containing protein 1) (Methylmalonyl-CoA decarboxylase) (MMCD) (EC 4.1.1.41) | 307 | ECHDC1 | 0 | 1 | 0 | 0 | 4.9 | 0 |
| 1326 | 0.00 | -0.11 | 0.00 | Q9NVS9 | Pyridoxine-5'-phosphate oxidase (EC 1.4.3.5) (Pyridoxamine-phosphate oxidase) | 261 | PNPO | 0 | 1 | 0 | 0 | 6.9 | 0 |
| 1327 | 0.00 | 0.02 | 0.00 | Q9NW13 | RNA-binding protein 28 (RNA-binding motif protein 28) | 759 | RBM28 | 0 | 1 | 0 | 0 | 1.5 | 0 |
| 1328 | 0.00 | 0.55 | 0.00 | Q9NZL9 | Methionine adenosyltransferase 2 subunit beta (Methionine adenosyltransferase II beta) (MAT II beta) (Putative dTDP-4-keto-6-deoxy-D-glucose 4-reductase) | 334 | MAT2B | 0 | 1 | 0 | 0 | 3.3 | 0 |
| 1329 | 0.00 | -1.44 | 0.00 | Q9P1F3 | Costars family protein ABRACL (ABRA C-terminal-like protein) | 81 | ABRACL | 0 | 1 | 0 | 0 | 16 | 0 |
| 1330 | 0.00 | 0.50 | 0.00 | Q9UDW1 | Cytochrome b-c1 complex subunit 9 (Complex III subunit 9) (Complex III subunit X) (Cytochrome c1 non-heme 7 kDa protein) (Ubiquinol-cytochrome c reductase complex 7.2 kDa protein) | 63 | UQCR10 | 0 | 1 | 0 | 0 | 27 | 0 |

| | | | | | | | | | | | | | |
|---|---|---|---|---|---|---|---|---|---|---|---|---|---|
| 1331 | 0.00 | 0.78 | 0.00 | Q9UHA4 | Ragulator complex protein LAMTOR3 (Late endosomal/lysosomal adaptor and MAPK and MTOR activator 3) (MEK-binding partner 1) (Mp1) (Mitogen-activated protein kinase kinase 1-interacting protein 1) (Mitogen-activated protein kinase scaffold protein 1) | 124 | LAMTOR3 | 0 | 1 | 0 | 0 | 13.7 | 0 |
| 1332 | 0.00 | 7.17 | 0.00 | Q9UKX3 | Myosin-13 (Myosin heavy chain 13) (Myosin heavy chain, skeletal muscle, extraocular) (MyHC-EO) (Myosin heavy chain, skeletal muscle, laryngeal) (MyHC-IIL) (Superfast myosin) | 1938 | MYH13 | 0 | 1 | 0 | 0 | 0.8 | 0 |
| 1333 | 0.00 | 1.24 | 0.00 | Q9Y5U9 | Immediate early response 3-interacting protein 1 | 82 | IER3IP1 | 0 | 1 | 0 | 0 | 24.4 | 0 |
| 1334 | 0.00 | 1.16 | 0.00 | R4GN18 | Membrane cofactor protein (Fragment) | 78 | CD46 | 0 | 1 | 0 | 0 | 16.7 | 0 |
| 1335 | 0.00 | 1.61 | 0.00 | R4GN83 | Basigin (Fragment) | 52 | BSG | 0 | 1 | 0 | 0 | 26.9 | 0 |
| 1336 | 0.00 | 0.91 | 0.00 | R4GN99 | Peptidyl-prolyl cis-trans isomerase (EC 5.2.1.8) | 145 | PPIF | 0 | 1 | 0 | 0 | 6.2 | 0 |
| 1337 | 0.00 | 1.15 | 0.00 | R4GNH9 | Exosome complex component CSL4 | 139 | EXOSC1 | 0 | 1 | 0 | 0 | 11.5 | 0 |
| 1338 | 0.00 | 2.84 | 0.00 | S4R3I5 | NADH dehydrogenase (Ubiquinone) 1 alpha subcomplex, 3, 9kDa, isoform CRA_b (NADH dehydrogenase [ubiquinone] 1 alpha subcomplex subunit 3) | 41 | NDUFA3 | 0 | 1 | 0 | 0 | 26.8 | 0 |
| 1339 | 0.00 | -0.76 | 0.00 | S4R402 | Nucleolar and coiled-body phosphoprotein 1 | 46 | NOLC1 | 0 | 1 | 0 | 0 | 21.7 | 0 |
| 1340 | 0.00 | -0.06 | 0.06 | O95670 | V-type proton ATPase subunit G 2 (V-ATPase subunit G 2) (V-ATPase 13 kDa subunit 2) (Vacuolar proton pump subunit G 2) | 118 | ATP6V1G2 | 1 | 1 | 1 | 19.2 | 19.2 | 19.2 |
| 1341 | 0.00 | 0.03 | 0.36 | Q3BDU5 | Prelamin-A/C (Rhabdomyosarcoma antigen MU-RMS-40.12) | 487 | LMNA | 5 | 3 | 6 | 11.7 | 7.2 | 14 |

| | | | | | | | | | | | | | |
|---|---|---|---|---|---|---|---|---|---|---|---|---|---|
| 1342 | 0.00 | -0.89 | -0.30 | H0YM70 | Proteasome activator complex subunit 2 | 228 | PSME2 | 4 | 3 | 4 | 28.9 | 19.3 | 28.1 |
| 1343 | 0.01 | -0.47 | 0.65 | P30046 | D-dopachrome decarboxylase (EC 4.1.1.84) (D-dopachrome tautomerase) (Phenylpyruvate tautomerase II) | 118 | DDT | 1 | 4 | 2 | 10.2 | 35.6 | 21.2 |
| 1344 | 0.01 | 0.00 | 0.00 | O00267 | Transcription elongation factor SPT5 (hSPT5) (DRB sensitivity-inducing factor 160 kDa subunit) (DSIF p160) (DRB sensitivity-inducing factor large subunit) (DSIF large subunit) (Tat-cotransactivator 1 protein) (Tat-CT1 protein) | 1087 | SUPT5H | 1 | 0 | 0 | 1.4 | 0 | 0 |
| 1345 | 0.01 | 0.95 | 0.88 | P19404 | NADH dehydrogenase [ubiquinone] flavoprotein 2, mitochondrial (EC 1.6.5.3) (EC 1.6.99.3) (NADH-ubiquinone oxidoreductase 24 kDa subunit) | 249 | NDUFV2 | 1 | 2 | 2 | 4 | 9.2 | 9.2 |
| 1346 | 0.01 | 0.22 | 0.66 | M0QZR9 | ELAV-like protein 1 | 153 | ELAVL1 | 1 | 1 | 1 | 8.5 | 8.5 | 8.5 |
| 1347 | 0.02 | 0.90 | 0.13 | P30084 | Enoyl-CoA hydratase, mitochondrial (EC 4.2.1.17) (Enoyl-CoA hydratase 1) (Short-chain enoyl-CoA hydratase) (SCEH) | 290 | ECHS1 | 1 | 5 | 2 | 6.2 | 27.6 | 9.3 |
| 1348 | 0.03 | 0.16 | -0.03 | Q00765 | Receptor expression-enhancing protein 5 (Polyposis locus protein 1) (Protein TB2) | 189 | REEP5 | 4 | 5 | 4 | 16.4 | 21.2 | 16.4 |
| 1349 | 0.03 | 0.25 | 1.21 | Q8N257 | Histone H2B type 3-B (H2B type 12) | 126 | HIST3H2BB | 2 | 3 | 3 | 15.1 | 23.8 | 27 |
| 1350 | 0.03 | -0.46 | 0.10 | Q3ZCM7 | Tubulin beta-8 chain | 444 | TUBB8 | 5 | 6 | 5 | 16 | 16.4 | 13.1 |
| 1351 | 0.03 | 0.79 | 0.10 | Q07065 | Cytoskeleton-associated protein 4 (63-kDa cytoskeleton-linking membrane protein) (Climp-63) (p63) | 602 | CKAP4 | 6 | 2 | 8 | 11.8 | 4.3 | 15.9 |

| | | | | | | | | | | | | | |
|---|---|---|---|---|---|---|---|---|---|---|---|---|---|
| 1352 | 0.04 | 0.63 | 1.04 | P53007 | Tricarboxylate transport protein, mitochondrial (Citrate transport protein) (CTP) (Solute carrier family 25 member 1) (Tricarboxylate carrier protein) | 311 | SLC25A1 | 2 | 2 | 2 | 7.7 | 7.4 | 7.7 |
| 1353 | 0.04 | 1.07 | 0.24 | Q5JPE7 | Nodal modulator 2 (pM5 protein 2) | 1267 | NOMO2 | 1 | 3 | 1 | 0.8 | 3.2 | 0.8 |
| 1354 | 0.04 | 0.36 | 0.41 | Q6Y1H2 | Very-long-chain (3R)-3-hydroxyacyl-CoA dehydratase 2 (EC 4.2.1.134) (3-hydroxyacyl-CoA dehydratase 2) (HACD2) (Protein-tyrosine phosphatase-like member B) | 254 | HACD2 | 1 | 1 | 1 | 3.9 | 3.5 | 3.9 |
| 1355 | 0.05 | -0.17 | 0.24 | H0YD18 | Nucleobindin-2 (Fragment) | 73 | NUCB2 | 1 | 1 | 1 | 12.3 | 12.3 | 12.3 |
| 1356 | 0.05 | 0.00 | 0.54 | P51610 | Host cell factor 1 (HCF) (HCF-1) (C1 factor) (CFF) (VCAF) (VP16 accessory protein) [Cleaved into: HCF N-terminal chain 1; HCF N-terminal chain 2; HCF N-terminal chain 3; HCF N-terminal chain 4; HCF N-terminal chain 5; HCF N-terminal chain 6; HCF C-terminal chain 1; HCF C-terminal chain 2; HCF C-terminal chain 3; HCF C-terminal chain 4; HCF C-terminal chain 5; HCF C-terminal chain 6] | 2035 | HCFC1 | 1 | 0 | 1 | 0.8 | 0 | 0.8 |
| 1357 | 0.05 | 0.25 | 0.88 | P62191 | 26S protease regulatory subunit 4 (P26s4) (26S proteasome AAA-ATPase subunit RPT2) (Proteasome 26S subunit ATPase 1) | 440 | PSMC1 | 1 | 1 | 3 | 2.5 | 5.7 | 10.9 |
| 1358 | 0.06 | 0.00 | 0.86 | A6NKB8 | Aminopeptidase B | 611 | RNPEP | 1 | 0 | 1 | 1.6 | 0 | 2 |
| 1359 | 0.06 | 0.50 | 0.65 | P05091 | Aldehyde dehydrogenase, mitochondrial (EC 1.2.1.3) (ALDH class 2) (ALDH-E2) (ALDHI) | 517 | ALDH2 | 7 | 7 | 8 | 16.2 | 16.8 | 19.1 |
| 1360 | 0.07 | 0.00 | 0.65 | P60891 | Ribose-phosphate pyrophosphokinase 1 (EC 2.7.6.1) (PPRibP) (Phosphoribosyl pyrophosphate synthase I) (PRS-I) | 318 | PRPS1 | 1 | 0 | 2 | 4.1 | 0 | 7.2 |

| | | | | | | | | | | | | | |
|---|---|---|---|---|---|---|---|---|---|---|---|---|---|
| 1361 | 0.07 | 0.10 | 1.13 | Q15005 | Signal peptidase complex subunit 2 (EC 3.4.-.-) (Microsomal signal peptidase 25 kDa subunit) (SPase 25 kDa subunit) | 226 | SPCS2 | 3 | 2 | 1 | 13.3 | 8 | 8.4 |
| 1362 | 0.08 | 0.40 | 0.52 | C9J0J7 | Profilin-2 | 91 | PFN2 | 1 | 2 | 1 | 8.8 | 29.7 | 8.8 |
| 1363 | 0.08 | 0.35 | 0.23 | P05141 | ADP/ATP translocase 2 (ADP,ATP carrier protein 2) (ADP,ATP carrier protein, fibroblast isoform) (Adenine nucleotide translocator 2) (ANT 2) (Solute carrier family 25 member 5) [Cleaved into: ADP/ATP translocase 2, N-terminally processed] | 298 | SLC25A5 | 7 | 7 | 8 | 21.8 | 24.2 | 24.5 |
| 1364 | 0.08 | 0.00 | -0.16 | D6REM1 | Golgi phosphoprotein 3 | 93 | GOLPH3 | 1 | 0 | 1 | 14 | 0 | 14 |
| 1365 | 0.08 | 0.39 | 0.49 | P34897 | Serine hydroxymethyltransferase, mitochondrial (SHMT) (EC 2.1.2.1) (Glycine hydroxymethyltransferase) (Serine methylase) | 504 | SHMT2 | 3 | 1 | 8 | 8.1 | 2.3 | 21.7 |
| 1366 | 0.09 | 0.00 | 0.28 | Q9Y3I0 | tRNA-splicing ligase RtcB homolog (EC 6.5.1.3) | 505 | RTCB | 1 | 0 | 1 | 3.2 | 0 | 3.2 |
| 1367 | 0.10 | 0.67 | 0.52 | A0A087X054 | Hypoxia up-regulated protein 1 | 937 | HYOU1 | 7 | 8 | 6 | 8 | 12.8 | 8.5 |
| 1368 | 0.10 | -0.09 | 0.33 | P38606 | V-type proton ATPase catalytic subunit A (V-ATPase subunit A) (EC 3.6.3.14) (V-ATPase 69 kDa subunit) (Vacuolar ATPase isoform VA68) (Vacuolar proton pump subunit alpha) | 617 | ATP6V1A | 1 | 1 | 2 | 3.1 | 3.1 | 3.4 |
| 1369 | 0.10 | 0.00 | 0.00 | Q15813 | Tubulin-specific chaperone E (Tubulin-folding cofactor E) | 527 | TBCE | 1 | 0 | 0 | 2.1 | 0 | 0 |
| 1370 | 0.10 | -0.03 | 0.44 | J3QT77 | Serum paraoxonase/arylesterase 2 | 342 | PON2 | 1 | 1 | 3 | 5.6 | 3.5 | 12.3 |
| 1371 | 0.10 | 0.03 | 0.44 | P12814 | Alpha-actinin-1 (Alpha-actinin cytoskeletal isoform) (F-actin cross-linking protein) (Non-muscle alpha-actinin-1) | 892 | ACTN1 | 13 | 20 | 13 | 20 | 27 | 20.3 |

| | | | | | | | | | | | | | |
|---|---|---|---|---|---|---|---|---|---|---|---|---|---|
| 1372 | 0.10 | -0.30 | 0.18 | O43324 | Eukaryotic translation elongation factor 1 epsilon-1 (Aminoacyl tRNA synthetase complex-interacting multifunctional protein 3) (Elongation factor p18) (Multisynthase complex auxiliary component p18) | 174 | EEF1E1 | 2 | 2 | 2 | 12.1 | 12.6 | 12.1 |
| 1373 | 0.11 | -0.60 | 0.30 | Q9NQ88 | Fructose-2,6-bisphosphatase TIGAR (EC 3.1.3.46) (TP53-induced glycolysis and apoptosis regulator) | 270 | TIGAR | 2 | 2 | 1 | 8.9 | 8.9 | 4.8 |
| 1374 | 0.11 | 0.00 | 0.83 | Q12792 | Twinfilin-1 (Protein A6) (Protein tyrosine kinase 9) | 350 | TWF1 | 1 | 0 | 2 | 2.8 | 0 | 10.3 |
| 1375 | 0.11 | -0.88 | 0.37 | Q9BY32 | Inosine triphosphate pyrophosphatase (ITPase) (Inosine triphosphatase) (EC 3.6.1.19) (Non-canonical purine NTP pyrophosphatase) (Non-standard purine NTP pyrophosphatase) (Nucleoside-triphosphate diphosphatase) (Nucleoside-triphosphate pyrophosphatase) (NTPase) (Putative oncogene protein hlc14-06-p) | 194 | ITPA | 5 | 3 | 3 | 24.2 | 16 | 19.1 |
| 1376 | 0.11 | 1.04 | -0.03 | H0YD13 | CD44 antigen | 206 | CD44 | 2 | 4 | 3 | 9.2 | 22.3 | 17 |
| 1377 | 0.12 | 0.00 | 0.25 | Q15046 | Lysine--tRNA ligase (EC 6.1.1.6) (Lysyl-tRNA synthetase) (LysRS) | 597 | KARS | 4 | 0 | 3 | 6.5 | 0 | 4.2 |
| 1378 | 0.13 | -0.95 | 0.07 | P30838 | Aldehyde dehydrogenase, dimeric NADP-preferring (EC 1.2.1.5) (ALDHIII) (Aldehyde dehydrogenase 3) (Aldehyde dehydrogenase family 3 member A1) | 453 | ALDH3A1 | 10 | 8 | 13 | 21.4 | 21.4 | 30 |
| 1379 | 0.13 | 0.36 | -0.22 | Q9UL25 | Ras-related protein Rab-21 | 225 | RAB21 | 2 | 2 | 1 | 12.4 | 12.4 | 7.6 |

| | | | | | | | | | | | | | |
|---|---|---|---|---|---|---|---|---|---|---|---|---|---|
| 1380 | 0.13 | 0.00 | 0.15 | Q14232 | Translation initiation factor eIF-2B subunit alpha (eIF-2B GDP-GTP exchange factor subunit alpha) | 305 | EIF2B1 | 2 | 0 | 2 | 6.6 | 0 | 8.2 |
| 1381 | 0.14 | -1.36 | -0.14 | P15428 | 15-hydroxyprostaglandin dehydrogenase [NAD(+)] (15-PGDH) (EC 1.1.1.141) (Prostaglandin dehydrogenase 1) (Short chain dehydrogenase/reductase family 36C member 1) | 266 | HPGD | 3 | 3 | 3 | 14.3 | 13.2 | 13.2 |
| 1382 | 0.14 | -0.07 | 0.28 | P20700 | Lamin-B1 | 586 | LMNB1 | 2 | 4 | 3 | 3.9 | 9.9 | 6 |
| 1383 | 0.14 | 0.00 | 0.00 | P49589 | Cysteine--tRNA ligase, cytoplasmic (EC 6.1.1.16) (Cysteinyl-tRNA synthetase) (CysRS) | 748 | CARS | 1 | 0 | 1 | 1.4 | 0 | 1.4 |
| 1384 | 0.15 | 0.00 | 0.06 | P62873 | Guanine nucleotide-binding protein G(I)/G(S)/G(T) subunit beta-1 (Transducin beta chain 1) | 340 | GNB1 | 2 | 1 | 2 | 6.6 | 3.3 | 6.6 |
| 1385 | 0.15 | -0.34 | 0.09 | O14579 | Coatomer subunit epsilon (Epsilon-coat protein) (Epsilon-COP) | 308 | COPE | 1 | 1 | 1 | 5.4 | 5.4 | 5.4 |
| 1386 | 0.16 | 1.44 | 0.25 | P51571 | Translocon-associated protein subunit delta (TRAP-delta) (Signal sequence receptor subunit delta) (SSR-delta) | 173 | SSR4 | 1 | 2 | 1 | 7.5 | 16.8 | 7.5 |
| 1387 | 0.16 | 0.83 | 0.68 | P62140 | Serine/threonine-protein phosphatase PP1-beta catalytic subunit (PP-1B) (PPP1CD) (EC 3.1.3.16) (EC 3.1.3.53) | 327 | PPP1CB | 1 | 2 | 3 | 4 | 9.5 | 13.8 |
| 1388 | 0.17 | 0.04 | 0.93 | Q9BVK6 | Transmembrane emp24 domain-containing protein 9 (GMP25) (Glycoprotein 25L2) (p24 family protein alpha-2) (p24alpha2) (p25) | 235 | TMED9 | 1 | 4 | 3 | 4.7 | 12.8 | 14.9 |
| 1389 | 0.17 | -0.28 | 0.00 | P36404 | ADP-ribosylation factor-like protein 2 | 184 | ARL2 | 1 | 2 | 0 | 6.5 | 12 | 0 |

| | | | | | | | | | | | | | |
|---|---|---|---|---|---|---|---|---|---|---|---|---|---|
| 1390 | 0.18 | 0.97 | 0.52 | B4DEZ3 | NADH dehydrogenase [ubiquinone] 1 alpha subcomplex subunit 13 (cDNA FLJ57958, highly similar to NADH dehydrogenase (ubiquinone) 1 alpha subcomplex subunit 13 (EC 1.6.5.3)) | 120 | NDUFA13 | 1 | 2 | 1 | 9.2 | 20.8 | 9.2 |
| 1391 | 0.18 | 0.00 | -0.34 | H7C4M9 | Ubiquitin-conjugating enzyme E2 H (Fragment) | 62 | UBE2H | 1 | 0 | 1 | 24.2 | 0 | 24.2 |
| 1392 | 0.18 | -0.45 | -0.11 | P18085 | ADP-ribosylation factor 4 | 180 | ARF4 | 3 | 3 | 3 | 17.2 | 22.8 | 17.2 |
| 1393 | 0.19 | 0.45 | 0.81 | P56556 | NADH dehydrogenase [ubiquinone] 1 alpha subcomplex subunit 6 (Complex I-B14) (CI-B14) (LYR motif-containing protein 6) (NADH-ubiquinone oxidoreductase B14 subunit) | 154 | NDUFA6 | 2 | 1 | 2 | 15.6 | 5.2 | 15.6 |
| 1394 | 0.19 | -1.06 | -0.17 | P04080 | Cystatin-B (CPI-B) (Liver thiol proteinase inhibitor) (Stefin-B) | 98 | CSTB | 2 | 1 | 2 | 21.4 | 12.2 | 21.4 |
| 1395 | 0.19 | 1.53 | 1.08 | Q9NZ45 | CDGSH iron-sulfur domain-containing protein 1 (MitoNEET) | 108 | CISD1 | 1 | 1 | 2 | 12 | 13.9 | 25.9 |
| 1396 | 0.20 | 0.51 | 0.33 | F5H6I7 | Atlastin-3 | 523 | ATL3 | 1 | 1 | 3 | 3.4 | 3.4 | 7.5 |
| 1397 | 0.20 | 0.00 | 0.58 | Q07973 | 1,25-dihydroxyvitamin D(3) 24-hydroxylase, mitochondrial (24-OHase) (Vitamin D(3) 24-hydroxylase) (EC 1.14.13.126) (Cytochrome P450 24A1) (Cytochrome P450-CC24) | 514 | CYP24A1 | 1 | 0 | 1 | 3.8 | 0 | 3.8 |
| 1398 | 0.20 | -0.17 | 0.00 | P29317 | Ephrin type-A receptor 2 (EC 2.7.10.1) (Epithelial cell kinase) (Tyrosine-protein kinase receptor ECK) | 976 | EPHA2 | 1 | 3 | 0 | 1 | 3.6 | 0 |
| 1399 | 0.20 | 1.05 | 1.02 | Q9BX68 | Histidine triad nucleotide-binding protein 2, mitochondrial (HINT-2) (EC 3.-.-.-) (HINT-3) (HIT-17kDa) (PKCI-1-related HIT protein) | 163 | HINT2 | 1 | 4 | 2 | 12.3 | 41.7 | 21.5 |

| | | | | | | | | | | | | | |
|---|---|---|---|---|---|---|---|---|---|---|---|---|---|
| 1400 | 0.21 | -0.29 | 0.17 | P48444 | Coatomer subunit delta (Archain) (Delta-coat protein) (Delta-COP) | 511 | ARCN1 | 2 | 2 | 2 | 4.1 | 4.3 | 4.1 |
| 1401 | 0.21 | -0.40 | -0.05 | P36406 | E3 ubiquitin-protein ligase TRIM23 (EC 6.3.2.-) (ADP-ribosylation factor domain-containing protein 1) (GTP-binding protein ARD-1) (RING finger protein 46) (Tripartite motif-containing protein 23) | 574 | TRIM23 | 1 | 1 | 1 | 1.8 | 1.8 | 1.8 |
| 1402 | 0.21 | -0.67 | -1.21 | Q01995 | Transgelin (22 kDa actin-binding protein) (Protein WS3-10) (Smooth muscle protein 22-alpha) (SM22-alpha) | 201 | TAGLN | 1 | 2 | 1 | 6 | 9.5 | 6 |
| 1403 | 0.22 | 0.00 | -0.06 | I3L1U8 | Active breakpoint cluster region-related protein (Fragment) | 165 | ABR | 1 | 0 | 1 | 6.1 | 0 | 6.1 |
| 1404 | 0.22 | 0.30 | 0.81 | O00469 | Procollagen-lysine,2-oxoglutarate 5-dioxygenase 2 (EC 1.14.11.4) (Lysyl hydroxylase 2) (LH2) | 737 | PLOD2 | 2 | 1 | 3 | 2.6 | 1.4 | 4.9 |
| 1405 | 0.22 | -0.71 | 0.07 | P12931 | Proto-oncogene tyrosine-protein kinase Src (EC 2.7.10.2) (Proto-oncogene c-Src) (pp60c-src) (p60-Src) | 536 | SRC | 2 | 1 | 2 | 5.2 | 2.8 | 5.2 |
| 1406 | 0.23 | 0.52 | 0.53 | B0YJC4 | Vimentin (Vimentin variant 3) | 431 | VIM | 3 | 2 | 2 | 8.6 | 3.9 | 5.6 |
| 1407 | 0.23 | 0.16 | 0.18 | Q15907 | Ras-related protein Rab-11B (GTP-binding protein YPT3) | 218 | RAB11B | 5 | 5 | 4 | 34.6 | 34.6 | 29.6 |
| 1408 | 0.23 | 0.76 | 0.30 | Q8NE86 | Calcium uniporter protein, mitochondrial (Coiled-coil domain-containing protein 109A) | 351 | MCU | 1 | 4 | 2 | 5.3 | 18.5 | 9.6 |
| 1409 | 0.23 | 0.00 | -0.11 | Q8NBU5 | ATPase family AAA domain-containing protein 1 (EC 3.6.1.3) (Thorase) | 361 | ATAD1 | 1 | 0 | 1 | 3.8 | 0 | 3.8 |
| 1410 | 0.23 | 0.00 | 0.10 | A0A087X1I3 | Succinate dehydrogenase [ubiquinone] flavoprotein subunit, mitochondrial | 519 | SDHA | 1 | 0 | 1 | 2.5 | 0 | 2.5 |

| | | | | | | | | | | | | | |
|---|---|---|---|---|---|---|---|---|---|---|---|---|---|
| 1411 | 0.23 | 0.00 | 1.43 | P55265 | Double-stranded RNA-specific adenosine deaminase (DRADA) (EC 3.5.4.37) (136 kDa double-stranded RNA-binding protein) (p136) (Interferon-inducible protein 4) (IFI-4) (K88DSRBP) | 1226 | ADAR | 1 | 0 | 1 | 1.1 | 0 | 1.1 |
| 1412 | 0.24 | 1.34 | 0.47 | Q6KC79 | Nipped-B-like protein (Delangin) (SCC2 homolog) | 2804 | NIPBL | 1 | 1 | 1 | 0.4 | 0.5 | 0.4 |
| 1413 | 0.24 | 0.03 | 0.61 | P49748 | Very long-chain specific acyl-CoA dehydrogenase, mitochondrial (VLCAD) (EC 1.3.8.9) | 655 | ACADVL | 2 | 2 | 7 | 3.2 | 3.5 | 15.3 |
| 1414 | 0.24 | 1.06 | 0.40 | Q16762 | Thiosulfate sulfurtransferase (EC 2.8.1.1) (Rhodanese) | 297 | TST | 1 | 3 | 1 | 4 | 14.5 | 4 |
| 1415 | 0.24 | 1.08 | 1.08 | C9JT21 | Elongation factor Ts (Fragment) | 168 | TSFM | 1 | 2 | 2 | 8.3 | 18.5 | 18.5 |
| 1416 | 0.24 | 0.01 | 0.76 | Q96QK1 | Vacuolar protein sorting-associated protein 35 (hVPS35) (Maternal-embryonic 3) (Vesicle protein sorting 35) | 796 | VPS35 | 2 | 1 | 1 | 3.5 | 1.8 | 1.8 |
| 1417 | 0.24 | 0.00 | 0.85 | Q96BP3 | Peptidylprolyl isomerase domain and WD repeat-containing protein 1 (EC 5.2.1.8) (Spliceosome-associated cyclophilin) | 646 | PPWD1 | 1 | 0 | 1 | 2.4 | 0 | 2.4 |
| 1418 | 0.24 | 0.23 | 0.09 | Q9BRX8 | Redox-regulatory protein FAM213A (Peroxiredoxin-like 2 activated in M-CSF stimulated monocytes) (Protein PAMM) | 229 | FAM213A | 3 | 3 | 4 | 13.8 | 17.4 | 20.6 |
| 1419 | 0.26 | -0.97 | 0.01 | C9JP16 | Cartilage-associated protein | 358 | CRTAP | 1 | 1 | 1 | 3.1 | 3.1 | 3.1 |
| 1420 | 0.26 | -0.08 | 0.20 | P51809 | Vesicle-associated membrane protein 7 (VAMP-7) (Synaptobrevin-like protein 1) (Tetanus-insensitive VAMP) (Ti-VAMP) | 220 | VAMP7 | 1 | 1 | 1 | 5 | 6.7 | 5 |
| 1421 | 0.26 | 0.00 | 0.01 | P67936 | Tropomyosin alpha-4 chain (TM30p1) (Tropomyosin-4) | 248 | TPM4 | 3 | 2 | 6 | 14.5 | 10.9 | 31 |

| | | | | | | | | | | | | | |
|---|---|---|---|---|---|---|---|---|---|---|---|---|---|
| 1422 | 0.26 | -0.49 | -0.04 | Q16851 | UTP--glucose-1-phosphate uridylyltransferase (EC 2.7.7.9) (UDP-glucose pyrophosphorylase) (UDPGP) (UGPase) | 508 | UGP2 | 2 | 1 | 2 | 7.6 | 3.6 | 6.4 |
| 1423 | 0.26 | 0.07 | 0.33 | P31930 | Cytochrome b-c1 complex subunit 1, mitochondrial (Complex III subunit 1) (Core protein I) (Ubiquinol-cytochrome-c reductase complex core protein 1) | 480 | UQCRC1 | 1 | 3 | 2 | 2.5 | 6.2 | 6.7 |
| 1424 | 0.27 | 0.50 | 0.53 | Q15363 | Transmembrane emp24 domain-containing protein 2 (Membrane protein p24A) (p24) (p24 family protein beta-1) (p24beta1) | 201 | TMED2 | 2 | 2 | 4 | 19.9 | 14.4 | 26.4 |
| 1425 | 0.27 | 0.12 | 0.36 | Q53H82 | Beta-lactamase-like protein 2 (EC 3.-.-.-) | 288 | LACTB2 | 1 | 4 | 3 | 2.8 | 16.7 | 12.2 |
| 1426 | 0.27 | -0.50 | -0.69 | P49902 | Cytosolic purine 5'-nucleotidase (EC 3.1.3.5) (Cytosolic 5'-nucleotidase II) | 561 | NT5C2 | 1 | 1 | 2 | 3.2 | 2.4 | 5.6 |
| 1427 | 0.28 | 0.64 | 0.43 | P26640 | Valine--tRNA ligase (EC 6.1.1.9) (Protein G7a) (Valyl-tRNA synthetase) (ValRS) | 1264 | VARS | 1 | 2 | 1 | 0.9 | 2.5 | 0.9 |
| 1428 | 0.28 | 0.61 | 0.16 | D6RAA6 | Transmembrane protein 33 (Fragment) | 222 | TMEM33 | 1 | 1 | 2 | 4.5 | 4.5 | 9.9 |
| 1429 | 0.28 | -0.74 | 0.14 | Q06210 | Glutamine--fructose-6-phosphate aminotransferase [isomerizing] 1 (EC 2.6.1.16) (D-fructose-6-phosphate amidotransferase 1) (Glutamine:fructose-6-phosphate amidotransferase 1) (GFAT 1) (GFAT1) (Hexosephosphate aminotransferase 1) | 699 | GFPT1 | 1 | 1 | 2 | 2.1 | 2.1 | 4.1 |
| 1430 | 0.28 | 0.00 | 0.33 | P53004 | Biliverdin reductase A (BVR A) (EC 1.3.1.24) (Biliverdin-IX alpha-reductase) | 296 | BLVRA | 3 | 0 | 2 | 11.5 | 0 | 6.8 |
| 1431 | 0.30 | 0.01 | 0.45 | P62330 | ADP-ribosylation factor 6 | 175 | ARF6 | 1 | 1 | 2 | 6.3 | 6.3 | 18.3 |

| | | | | | | | | | | | | | |
|---|---|---|---|---|---|---|---|---|---|---|---|---|---|
| 1432 | 0.30 | 0.36 | 0.19 | Q9BUP3 | Oxidoreductase HTATIP2 (EC 1.1.1.-) (30 kDa HIV-1 TAT-interacting protein) (HIV-1 TAT-interactive protein 2) | 242 | HTATIP2 | 1 | 3 | 4 | 4.1 | 12.4 | 16.1 |
| 1433 | 0.31 | 0.30 | 1.56 | D6RAT0 | 40S ribosomal protein S3a | 227 | RPS3A | 4 | 4 | 6 | 20.3 | 18.9 | 23.3 |
| 1434 | 0.31 | 0.00 | 0.39 | P46977 | Dolichyl-diphosphooligosaccharide--protein glycosyltransferase subunit STT3A (Oligosaccharyl transferase subunit STT3A) (STT3-A) (EC 2.4.99.18) (B5) (Integral membrane protein 1) (Transmembrane protein TMC) | 705 | STT3A | 1 | 0 | 1 | 1.3 | 0 | 1.3 |
| 1435 | 0.31 | 0.54 | 0.00 | E5RJZ1 | Cytochrome c oxidase subunit 7A-related protein, mitochondrial | 79 | COX7A2L | 1 | 2 | 0 | 16.5 | 26.6 | 0 |
| 1436 | 0.32 | 0.59 | 0.87 | Q9UM22 | Mammalian ependymin-related protein 1 (MERP-1) (Upregulated in colorectal cancer gene 1 protein) | 224 | EPDR1 | 1 | 1 | 1 | 5.5 | 5.5 | 5.5 |
| 1437 | 0.34 | 0.86 | 1.26 | B4DJ81 | NADH-ubiquinone oxidoreductase 75 kDa subunit, mitochondrial (cDNA FLJ60586, highly similar to NADH-ubiquinone oxidoreductase 75 kDa subunit, mitochondrial (EC 1.6.5.3)) | 611 | NDUFS1 | 1 | 1 | 2 | 2.6 | 2.6 | 4.3 |
| 1438 | 0.34 | 0.48 | 0.05 | F5GX19 | Ragulator complex protein LAMTOR1 | 142 | LAMTOR1 | 1 | 3 | 1 | 9.2 | 26.1 | 9.2 |
| 1439 | 0.34 | 0.00 | 1.08 | M0R261 | 6-phosphogluconolactonase (Fragment) | 216 | PGLS | 1 | 0 | 3 | 5.1 | 0 | 25 |
| 1440 | 0.35 | -0.04 | 0.08 | Q9NUJ1 | Mycophenolic acid acyl-glucuronide esterase, mitochondrial (EC 3.1.1.93) (Alpha/beta hydrolase domain-containing protein 10) (Abhydrolase domain-containing protein 10) | 306 | ABHD10 | 1 | 2 | 2 | 8.3 | 14.1 | 12.7 |

| | | | | | | | | | | | | | |
|---|---|---|---|---|---|---|---|---|---|---|---|---|---|
| 1441 | 0.35 | 0.00 | 1.04 | Q12906 | Interleukin enhancer-binding factor 3 (Double-stranded RNA-binding protein 76) (DRBP76) (M-phase phosphoprotein 4) (MPP4) (Nuclear factor associated with dsRNA) (NFAR) (Nuclear factor of activated T-cells 90 kDa) (NF-AT-90) (Translational control protein 80) (TCP80) | 894 | ILF3 | 2 | 0 | 2 | 2.8 | 0 | 2.8 |
| 1442 | 0.35 | 0.16 | 0.27 | Q13011 | Delta(3,5)-Delta(2,4)-dienoyl-CoA isomerase, mitochondrial (EC 5.3.3.-) | 328 | ECH1 | 1 | 3 | 1 | 3.4 | 12.8 | 3.4 |
| 1443 | 0.35 | 0.00 | 0.36 | E7ER27 | Peroxisomal multifunctional enzyme type 2 | 500 | HSD17B4 | 1 | 0 | 1 | 2.8 | 0 | 2.8 |
| 1444 | 0.36 | 0.00 | 0.00 | P10155 | 60 kDa SS-A/Ro ribonucleoprotein (60 kDa Ro protein) (60 kDa ribonucleoprotein Ro) (RoRNP) (Ro 60 kDa autoantigen) (Sjoegren syndrome antigen A2) (Sjoegren syndrome type A antigen) (SS-A) (TROVE domain family member 2) | 538 | TROVE2 | 1 | 0 | 0 | 2 | 0 | 0 |
| 1445 | 0.36 | 0.29 | 0.13 | Q969H8 | Myeloid-derived growth factor (MYDGF) (Interleukin-25) (IL-25) (Stromal cell-derived growth factor SF20) | 173 | MYDGF | 1 | 2 | 1 | 5.2 | 13.9 | 5.2 |
| 1446 | 0.37 | -1.26 | 0.40 | B0QZ43 | Erlin-1 (Fragment) | 275 | ERLIN1 | 1 | 2 | 1 | 4.4 | 8.7 | 4.4 |
| 1447 | 0.37 | -0.48 | -0.19 | P14550 | Alcohol dehydrogenase [NADP(+)] (EC 1.1.1.2) (Aldehyde reductase) (Aldo-keto reductase family 1 member A1) | 325 | AKR1A1 | 2 | 2 | 2 | 7.4 | 6.8 | 7.4 |
| 1448 | 0.37 | 0.00 | 0.32 | Q16658 | Fascin (55 kDa actin-bundling protein) (Singed-like protein) (p55) | 493 | FSCN1 | 1 | 0 | 2 | 2 | 0 | 5.3 |

| | | | | | | | | | | | | | |
|---|---|---|---|---|---|---|---|---|---|---|---|---|---|
| 1449 | 0.37 | 0.27 | 0.45 | O75874 | Isocitrate dehydrogenase [NADP] cytoplasmic (IDH) (EC 1.1.1.42) (Cytosolic NADP-isocitrate dehydrogenase) (IDP) (NADP(+)-specific ICDH) (Oxalosuccinate decarboxylase) | 414 | IDH1 | 5 | 3 | 5 | 13 | 8.2 | 14.7 |
| 1450 | 0.38 | 0.46 | 0.49 | F8W7Q4 | Protein FAM162A | 144 | FAM162A | 1 | 2 | 2 | 7.6 | 15.3 | 19.4 |
| 1451 | 0.40 | -0.20 | 0.44 | F5GYN4 | Ubiquitin thioesterase OTUB1 | 241 | OTUB1 | 3 | 3 | 5 | 14.5 | 17.4 | 27.4 |
| 1452 | 0.41 | 0.00 | 1.21 | P07305 | Histone H1.0 (Histone H1') (Histone H1(0)) [Cleaved into: Histone H1.0, N-terminally processed] | 194 | H1F0 | 1 | 0 | 1 | 7.3 | 0 | 7.3 |
| 1453 | 0.41 | 0.00 | 1.07 | Q16795 | NADH dehydrogenase [ubiquinone] 1 alpha subcomplex subunit 9, mitochondrial (Complex I-39kD) (CI-39kD) (NADH-ubiquinone oxidoreductase 39 kDa subunit) | 377 | NDUFA9 | 1 | 0 | 2 | 2.7 | 0 | 8 |
| 1454 | 0.41 | 0.00 | -0.02 | H0Y6T7 | Nicastrin (Fragment) | 275 | NCSTN | 2 | 0 | 1 | 7.3 | 0 | 4.4 |
| 1455 | 0.42 | 0.00 | 0.37 | G3V2G6 | Retinol dehydrogenase 11 (Fragment) | 178 | RDH11 | 1 | 1 | 1 | 7.3 | 7.3 | 7.3 |
| 1456 | 0.42 | 1.99 | 1.98 | P52292 | Importin subunit alpha-1 (Karyopherin subunit alpha-2) (RAG cohort protein 1) (SRP1-alpha) | 529 | KPNA2 | 3 | 2 | 2 | 7 | 7 | 5.5 |
| 1457 | 0.42 | 0.00 | 0.00 | Q6SZW1 | Sterile alpha and TIR motif-containing protein 1 (Sterile alpha and Armadillo repeat protein) (Sterile alpha motif domain-containing protein 2) (MyD88-5) (SAM domain-containing protein 2) (Tir-1 homolog) | 724 | SARM1 | 1 | 0 | 0 | 1.6 | 0 | 0 |
| 1458 | 0.43 | 0.70 | 1.27 | Q16698 | 2,4-dienoyl-CoA reductase, mitochondrial (EC 1.3.1.34) (2,4-dienoyl-CoA reductase [NADPH]) (4-enoyl-CoA reductase [NADPH]) (Short chain dehydrogenase/reductase family 18C member 1) | 335 | DECR1 | 1 | 4 | 2 | 4 | 17.8 | 10.7 |

| | | | | | | | | | | | | | |
|---|---|---|---|---|---|---|---|---|---|---|---|---|---|
| 1459 | 0.43 | 0.18 | -0.23 | Q56VL3 | OCIA domain-containing protein 2 (Ovarian carcinoma immunoreactive antigen-like protein) | 154 | OCIAD2 | 1 | 3 | 1 | 8.4 | 21.4 | 8.4 |
| 1460 | 0.44 | 0.16 | 0.49 | O95994 | Anterior gradient protein 2 homolog (AG-2) (hAG-2) (HPC8) (Secreted cement gland protein XAG-2 homolog) | 175 | AGR2 | 4 | 4 | 3 | 26.3 | 36 | 33.1 |
| 1461 | 0.44 | 1.96 | 0.43 | Q9UI09 | NADH dehydrogenase [ubiquinone] 1 alpha subcomplex subunit 12 (13 kDa differentiation-associated protein) (Complex I-B17.2) (CI-B17.2) (CIB17.2) (NADH-ubiquinone oxidoreductase subunit B17.2) | 145 | NDUFA12 | 1 | 1 | 1 | 5.5 | 11.7 | 5.5 |
| 1462 | 0.44 | 0.09 | -0.09 | Q16836 | Hydroxyacyl-coenzyme A dehydrogenase, mitochondrial (HCDH) (EC 1.1.1.35) (Medium and short-chain L-3-hydroxyacyl-coenzyme A dehydrogenase) (Short-chain 3-hydroxyacyl-CoA dehydrogenase) | 314 | HADH | 1 | 1 | 1 | 3.2 | 3.2 | 3.2 |
| 1463 | 0.44 | -0.16 | 0.06 | P40616 | ADP-ribosylation factor-like protein 1 | 181 | ARL1 | 1 | 2 | 1 | 5.5 | 12.2 | 5.5 |
| 1464 | 0.46 | 0.00 | -0.57 | B4DHN5 | Syntenin-1 (cDNA FLJ55055, moderately similar to Syntenin-1) | 239 | SDCBP | 1 | 0 | 2 | 4.6 | 0 | 8.8 |
| 1465 | 0.48 | 0.23 | 0.08 | P23434 | Glycine cleavage system H protein, mitochondrial (Lipoic acid-containing protein) | 173 | GCSH | 1 | 2 | 1 | 8.1 | 13.9 | 8.1 |
| 1466 | 0.50 | 0.54 | 0.79 | H3BPJ9 | NADH dehydrogenase [ubiquinone] 1 beta subcomplex subunit 10 | 161 | NDUFB10 | 1 | 2 | 2 | 6.2 | 14.9 | 14.3 |
| 1467 | 0.51 | 0.11 | -0.02 | P18564 | Integrin beta-6 | 788 | ITGB6 | 1 | 1 | 1 | 1.3 | 1.3 | 1.3 |

| | | | | | | | | | | | | | |
|---|---|---|---|---|---|---|---|---|---|---|---|---|---|
| 1468 | 0.52 | -0.60 | 0.01 | P21980 | Protein-glutamine gamma-glutamyltransferase 2 (EC 2.3.2.13) (Tissue transglutaminase) (Transglutaminase C) (TG(C)) (TGC) (TGase C) (Transglutaminase H) (TGase H) (Transglutaminase-2) (TGase-2) | 687 | TGM2 | 6 | 2 | 4 | 9.5 | 3.8 | 7.1 |
| 1469 | 0.53 | 0.00 | -0.46 | P48735 | Isocitrate dehydrogenase [NADP], mitochondrial (IDH) (EC 1.1.1.42) (ICD-M) (IDP) (NADP(+)-specific ICDH) (Oxalosuccinate decarboxylase) | 452 | IDH2 | 2 | 1 | 1 | 5 | 2.8 | 2.8 |
| 1470 | 0.54 | 0.52 | 0.77 | E9PFN5 | Glutathione S-transferase kappa 1 | 190 | GSTK1 | 1 | 4 | 4 | 7.4 | 26.8 | 25.8 |
| 1471 | 0.55 | 0.00 | 0.44 | A0A024R7W5 | YTH domain family, member 3, isoform CRA_a (YTH domain-containing family protein 3) | 534 | YTHDF3 | 1 | 0 | 1 | 2.6 | 0 | 2.6 |
| 1472 | 0.55 | 0.00 | -0.18 | D6RG15 | Twinfilin-2 | 254 | TWF2 | 2 | 0 | 2 | 9.8 | 0 | 9.8 |
| 1473 | 0.55 | 0.00 | -0.30 | Q9UNH7 | Sorting nexin-6 (TRAF4-associated factor 2) [Cleaved into: Sorting nexin-6, N-terminally processed] | 406 | SNX6 | 1 | 0 | 1 | 3.8 | 0 | 3.8 |
| 1474 | 0.58 | 0.40 | 0.26 | P14314 | Glucosidase 2 subunit beta (80K-H protein) (Glucosidase II subunit beta) (Protein kinase C substrate 60.1 kDa protein heavy chain) (PKCSH) | 528 | PRKCSH | 2 | 2 | 3 | 4.2 | 4.4 | 6.3 |
| 1475 | 0.59 | 0.25 | -0.11 | P22307 | Non-specific lipid-transfer protein (NSL-TP) (EC 2.3.1.176) (Propanoyl-CoA C-acyltransferase) (SCP-chi) (SCPX) (Sterol carrier protein 2) (SCP-2) (Sterol carrier protein X) (SCP-X) | 547 | SCP2 | 1 | 6 | 2 | 13.6 | 30 | 16.4 |

| | | | | | | | | | | | | | |
|---|---|---|---|---|---|---|---|---|---|---|---|---|---|
| 1476 | 0.61 | 0.26 | -0.05 | Q9BW60 | Elongation of very long chain fatty acids protein 1 (EC 2.3.1.199) (3-keto acyl-CoA synthase ELOVL1) (ELOVL fatty acid elongase 1) (ELOVL FA elongase 1) (Very-long-chain 3-oxoacyl-CoA synthase 1) | 279 | ELOVL1 | 1 | 1 | 1 | 5.2 | 5.2 | 5.2 |
| 1477 | 0.61 | 0.00 | -0.23 | O43175 | D-3-phosphoglycerate dehydrogenase (3-PGDH) (EC 1.1.1.95) | 533 | PHGDH | 1 | 0 | 2 | 2.4 | 0 | 4.7 |
| 1478 | 0.62 | -0.22 | 0.12 | Q6YN16 | Hydroxysteroid dehydrogenase-like protein 2 (EC 1.-.-.-) (Short chain dehydrogenase/reductase family 13C member 1) | 418 | HSDL2 | 2 | 1 | 2 | 9.6 | 5.2 | 9.6 |
| 1479 | 0.64 | 3.22 | 2.05 | P14923 | Junction plakoglobin (Catenin gamma) (Desmoplakin III) (Desmoplakin-3) | 745 | JUP | 1 | 10 | 1 | 1.3 | 15.8 | 1.6 |
| 1480 | 0.64 | -0.62 | -0.47 | G3XAM7 | Catenin (Cadherin-associated protein), alpha 1, 102kDa, isoform CRA_a (Catenin alpha-1) | 841 | CTNNA1 | 4 | 3 | 4 | 6.3 | 4.9 | 6.3 |
| 1481 | 0.66 | -1.33 | -0.10 | P40261 | Nicotinamide N-methyltransferase (EC 2.1.1.1) | 264 | NNMT | 3 | 3 | 4 | 17.4 | 17.4 | 17 |
| 1482 | 0.69 | -1.30 | -0.31 | M0R0M3 | Gamma-glutamylaminecyclotransferase (Fragment) | 26 | GGACT | 1 | 1 | 1 | 38.5 | 38.5 | 38.5 |
| 1483 | 0.72 | 0.00 | 0.48 | H0YM76 | WD repeat-containing protein 61 | 48 | WDR61 | 1 | 0 | 1 | 33.3 | 0 | 33.3 |
| 1484 | 0.73 | 0.31 | -0.19 | P19367 | Hexokinase-1 (EC 2.7.1.1) (Brain form hexokinase) (Hexokinase type I) (HK I) | 917 | HK1 | 2 | 2 | 2 | 2.5 | 2.5 | 2.5 |
| 1485 | 0.75 | -1.17 | -0.78 | O00194 | Ras-related protein Rab-27B (C25KG) | 218 | RAB27B | 2 | 2 | 2 | 10.1 | 10.1 | 10.1 |
| 1486 | 0.80 | 0.33 | 0.07 | G3XAL9 | Solute carrier family 12 (Sodium/potassium/chloride transporters), member 2, isoform CRA_a (Solute carrier family 12 member 2) | 1150 | SLC12A2 | 1 | 2 | 1 | 1.9 | 3 | 1.9 |

| | | | | | | | | | | | | | |
|---|---|---|---|---|---|---|---|---|---|---|---|---|---|
| 1487 | 0.90 | -0.83 | -0.47 | F5H3C5 | Superoxide dismutase [Mn], mitochondrial (Fragment) | 111 | SOD2 | 6 | 5 | 6 | 47.7 | 45.9 | 47.7 |
| 1488 | 0.94 | 0.00 | 0.00 | F8WAU7 | Vesicle-trafficking protein SEC22a | 71 | SEC22A | 1 | 0 | 0 | 14.1 | 0 | 0 |
| 1489 | 0.99 | 0.00 | -0.78 | Q2TB90 | Putative hexokinase HKDC1 (EC 2.7.1.1) (Hexokinase domain-containing protein 1) | 917 | HKDC1 | 1 | 1 | 2 | 1.1 | 1 | 2.1 |
| 1490 | 1.01 | -1.43 | -0.89 | Q6P452 | Annexin | 299 | ANXA4 | 5 | 10 | 4 | 18.7 | 30.1 | 17.1 |
| 1491 | 1.13 | -1.82 | -0.21 | Q9BS40 | Latexin (Endogenous carboxypeptidase inhibitor) (ECI) (Protein MUM) (Tissue carboxypeptidase inhibitor) (TCI) | 222 | LXN | 1 | 1 | 1 | 4.1 | 4.1 | 4.1 |
| 1492 | 1.39 | 0.00 | -0.22 | O14684 | Prostaglandin E synthase (EC 5.3.99.3) (Microsomal glutathione S-transferase 1-like 1) (MGST1-L1) (Microsomal prostaglandin E synthase 1) (MPGES-1) (p53-induced gene 12 protein) | 152 | PTGES | 1 | 0 | 1 | 6.6 | 0 | 6.6 |
