## Supplemental Table S2 for "Anti-inflammatory role of curcumin in Lipopolysaccharide treated A549 cells at global proteome level and on mycobacterial infection"

### Table S2A. List of all identified phosphoproteins from SILAC experiments in CURI

| S. No. | log2 CURI | Amino acid | Number of Phospho (STY) | Protein | Protein names | Gene names | Charge | Localization prob | PEP | Score | Delta score | Score for localization | Mass error [ppm] | Position | Positions within proteins | Sequence window |
|---|---|---|---|---|---|---|---|---|---|---|---|---|---|---|---|---|
| 1 | -0.19 | S | 1 | E7ENN3 | Nesprin-1 | SYNE1 | 2 | 0.90 | 0.01 | 63.18 | 63.18 | 63.18 | -0.92 | 1916 | 1916;1896;1903;1896 | SGILRQLRQTVEATNSMNKNESDLIEKDLND |
| 2 | 6.28 | S | 2 | O75691 | Small subunit processome component 20 homolog | UTP20 | 2 | 1 | 0.01 | 72.32 | 22.13 | 72.32 | 1.02 | 241 | 241 | QLLFEMCKGVRNMFHSCTGQAVKLILRKLGP |
| 3 | 5.10 | S | 2 | Q5VY30 | Retinol-binding protein 4;Plasma retinol-binding protein(1-182);Plasma retinol-binding protein(1-181);Plasma retinol-binding protein(1-179);Plasma retinol-binding protein(1-176) | RBP4 | 2 | 1 | 0.01 | 69.43 | 23.32 | 69.43 | -0.14 | 194 | 194;196 | RQYRLIVHNGYCDGRSERNLL__________ |
| 4 | -2.02 | S | 1 | P35558 | Phosphoenolpyruvate carboxykinase, cytosolic [GTP] | PCK1 | 2 | 0.97 | 0.01 | 91.31 | 9.68 | 91.31 | -0.97 | 286 | 286 | TNPEGEKKYLAAAFPSACGKTNLAMMNPSLP |
| 5 | -1.15 | S | 1 | P51858-2 | Hepatoma-derived growth factor | HDGF | 2 | 1 | 0.003 | 103.04 | 67.36 | 103.04 | -0.83 | 158 | 158;165;181 | EKGALKRRAGDLLEDSPKRPKEAENPEGEEK |
| 6 | -0.64 | S | 1 | Q5JR89 | Kinesin-like protein KIF2C | KIF2C | 2 | 0.99 | 0.008 | 72.2 | 27.58 | 72.2 | -1.01 | 273 | 273;241;228;282 | PLNKQELAKKEIDVISIPSKCLLLVHEPKL_ |
| 7 | 8.87 | S | 2 | Q7L513-15 | | | 2 | 1 | 0.02 | 75.10 | 26.55 | 74.40 | -1.20 | 6 | 6;6;6;6;6;6;6 | __________MLKKISVGVAGDLNTVTMKLG |
| 8 | -3.95 | S | 1 | Q8N0V4 | Leucine-rich repeat LGI family member 2 | LGI2 | 2 | 1 | 0.01 | 81.27 | 10.87 | 81.27 | -0.005 | 461 | 461 | MRWNSKQFVEIQALPSRGAMTLQPFSFKDNH |
| 9 | -1.71 | S | 2 | Q8NB66 | Protein unc-13 homolog C | UNC13C | 2 | 1 | 0.002 | 104.35 | 10.26 | 104.35 | -0.90 | 1534 | 1534 | LTSITFFRMKVLELQSPPKASMVVKDCVRAC |
| 10 | -1.71 | S | 2 | Q8NB66 | Protein unc-13 homolog C | UNC13C | 2 | 1 | 0.002 | 104.3 | 10.26 | 104.35 | -0.90 | 1539 | 1539 | FFRMKVLELQSPPKASMVVKDCVRACLDSTY |
| 11 | 0.06 | Y | 1 | C9JEV4 | Tectonin beta-propeller repeat-containing protein 1 | TECPR1 | 2 | 0.99 | 0.01 | 72.81 | 72.81 | 72.81 | -0.91 | 24 | 24;24;24;24;24;24 | VDLFGRVYTLSTAGQYWEMCKDSQLEFKRVS |
| 12 | 0.05 | Y | 1 | D6REQ7 | Protein ZGRF1 | ZGRF1 | 2 | 0.84 | 0.01 | 78.50 | 41.51 | 78.50 | -0.92 | 10 | 10;10;10;10;10;10;10;10 | ______MESQEFIVLYTHQKMKKSKVWQDGI |
| 13 | 8.11 | Y | 1 | G3V500 | Echinoderm microtubule-associated protein-like 1 | EML1 | 3 | 1 | 0.004 | 93.10 | 49.26 | 93.10 | -0.42 | 155 | 155;173;174;186;205 | SESKPKEPVFSAEEGYVKMFLRGRPVTMYMP |
| 14 | 5.10 | Y | 2 | Q5VY30 | Retinol-binding protein 4;Plasma retinol-binding protein(1-182);Plasma retinol-binding protein(1-181);Plasma retinol-binding protein(1-179);Plasma retinol-binding protein(1-176) | RBP4 | 2 | 1 | 0.01 | 69.43 | 23.32 | 69.43 | -0.14 | 189 | 189;191 | ELCLARQYRLIVHNGYCDGRSERNLL_____ |
| 15 | 0.09 | Y | 0 | Q5VWW1-3 | Complement C1q-like protein 3 | C1QL3 | 2 | 1 | 0.007 | 76.67 | 76.67 | 76.67 | -2.2 | 98 | 98;116;140 | PKIAFYAGLKRQHEGYEVLKFDDVVTNLGNH |

### Table S2B. List of all identified phosphoproteins from SILAC experiments in LPSI

| S. No. | log2 LPSI | Amino acid | Number of Phospho (STY) | Protein | Protein names | Gene names | Charge | Localization prob | PEP | Score | Delta score | Score for localization | Mass error [ppm] | Position | Positions within proteins | Sequence window |
|---|---|---|---|---|---|---|---|---|---|---|---|---|---|---|---|---|
| 1 | 0.86 | S | 1 | A0A0A0MRP6 | Probable global transcription activator SNF2L1 | SMARCA1 | 2 | 0.99 | 0.02 | 68.66 | 14.90 | 68.66 | 1.17 | 420 | 420;420;420;420 | KSLPPKKEIKIYLGLSKMQREWYTKILMKDI |
| 2 | -0.90 | S | 1 | C9J8H5 | Zinc finger matrin-type protein 3 | ZMAT3 | 3 | 1.00 | 0.03 | 53.26 | 53.26 | 53.26 | -0.44 | 44 | 44;44;44 | GTLQLPPQKPFGQEASLPLAGEEELSKGGEQ |
| 3 | 8.00 | S | 1 | C9JZY6 | Ubiquitin-conjugating enzyme E2 H | UBE2H | 2 | 1.00 | 0.01 | 66.06 | 39.27 | 66.06 | 1.45 | 2 | 2;2;2 | ______________MSSPSPGKRRMDTDVVK |
| 4 | -8.63 | S | 2 | D6RFK7 | | THAP6 | 3 | 1.00 | 0.02 | 67.42 | 63.31 | 67.42 | -0.25 | 50 | 50 | PNIKLKPGVIPSIFDSPYHLQKHRKRKQEQE |
| 5 | -8.63 | S | 2 | D6RFK7 | | THAP6 | 3 | 1.00 | 0.02 | 67.42 | 63.31 | 67.42 | -0.25 | 46 | 46 | DRSAPNIKLKPGVIPSIFDSPYHLQKHRKRK |
| 6 | 4.28 | S | 1 | E5RJ66 | Leucine-rich repeat-containing protein 69 | LRRC69 | 2 | 1.00 | 0.03 | 65.57 | 16.51 | 65.57 | -0.27 | 31 | 31;31;31;31 | TKIITLNGKKMTKMPSALGKLPGLKTLVLQN |
| 7 | 0.78 | T | | E7EPV1 | Speedy protein A | SPDYA | 2 | 1.00 | 0.01 | 96.33 | 35.30 | 96.33 | -1.44 | 104 | 104;104;104 | MDCCCKIADKYLLAMTFVYFKRAKFTISEHT |
| 8 | 5.86 | S | 1 | E9PMS6 | LIM domain only protein 7 | LMO7 | 2 | 1.00 | 0.00 | 118.24 | 40.58 | 92.94 | -0.17 | 1053 | 1053;1176;1176;1409;1446;1127;1176;1461;1461;1461 | NMTSSQRRSKKEQVPSGAELERQQILQEMRK |
| 9 | 5.19 | S | 1 | F5H0J3 | | NDUFA9 | 3 | 1.00 | 0.01 | 81.81 | 16.63 | 81.81 | -0.98 | 9 | 9 | _______MDQKAEVASQVEVVIFLKKKNQFR |
| 10 | -9.75 | Y | 1 | G3V500 | Echinoderm microtubule-associated protein-like 1 | EML1 | 3 | 1.00 | 0.01 | 85.99 | 57.36 | 77.73 | -1.09 | 155 | 155;173;174;186;205 | SESKPKEPVFSAEEGYVKMFLRGRPVTMYMP |
| 11 | 6.01 | S | 1 | P28329-3 | Choline O-acetyltransferase | CHAT | 3 | 0.85 | 0.01 | 91.59 | 56.49 | 81.53 | 0.96 | 77 | 77;113;195 | QQKLLERQEKTANWVSEYWLNDMYLNNRLAL |
| 12 | 0.35 | T | 2 | Q05193-5 | Dynamin-1 | DNM1 | 2 | 1.00 | 0.01 | 96.05 | 96.05 | 96.05 | -0.31 | 461 | 461;461;461;461 | LQQYPRLREEMERIVTTHIREREGRTKEQVM |
| 13 | 0.35 | T | 2 | Q05193-5 | Dynamin-1 | DNM1 | 2 | 1.00 | 0.01 | 96.05 | 96.05 | 96.05 | -0.31 | 462 | 462;462;462;462 | QQYPRLREEMERIVTTHIREREGRTKEQVML |
| 14 | 0.18 | S | 1 | Q5JR89 | Kinesin-like protein KIF2C | KIF2C | 2 | 1.00 | 0.01 | 67.39 | 30.97 | 67.39 | -0.32 | 273 | 273;241;228;282 | PLNKQELAKKEIDVISIPSKCLLLVHEPKL_ |
| 15 | 0.41 | S | | Q5T1R4-2 | Transcription factor HIVEP3 | HIVEP3 | 2 | 0.91 | 0.02 | 68.26 | 31.22 | 68.26 | 0.89 | 2254 | 2254;2255 | ERGRWSPTESSSASVSPVAKVSKFTLSSELE |
| 16 | 6.50 | S | 1 | Q7L311 | Armadillo repeat-containing X-linked protein 2 | ARMCX2 | 3 | 1.00 | 0.02 | 58.48 | 28.44 | 58.48 | -0.99 | 533 | 533 | SQGGGKIKVEILKILSNFAENPDMLKKLLST |
| 17 | -2.34 | S | 1 | Q8ND07 | Basal body-orientation factor 1 | CCDC176 | 3 | 1.00 | 0.02 | 82.59 | 34.65 | 82.59 | -1.46 | 37 | 37 | IKTDESVVDRAKANASLWEARLEVTELSRIK |
| 18 | 6.59 | S | 2 | Q8NDT2-2 | Putative RNA-binding protein 15B | RBM15B | 2 | 1.00 | 0.00 | 112.30 | 74.75 | 112.30 | -0.43 | 301 | 301;628 | ERSRTKGSGQQSERGSDRTPERSRKENHSSE |
| 19 | 6.59 | T | 2 | Q8NDT2-2 | Putative RNA-binding protein 15B | RBM15B | 2 | 1.00 | 0.00 | 112.30 | 74.75 | 112.30 | -0.43 | 304 | 304;631 | RTKGSGQQSERGSDRTPERSRKENHSSEGTK |
| 20 | 2.51 | S | 1 | Q96JS4 | | hucep-3 | 2 | 0.87 | 0.03 | 65.24 | 23.05 | 65.24 | 0.75 | 93 | 93 | GKGMGLWGRGGMGFRSICTIRKVLRSFFLEG |
| 21 | 9.57 | S | 2 | Q9BXJ3 | Complement C1q tumor necrosis factor-related protein 4 | C1QTNF4 | 2 | 1.00 | 0.02 | 63.18 | 63.18 | 63.18 | 0.39 | 244 | 244 | YFFSFTLGKLPRKTLSVKLMKNRDEVQAMIY |

#### Table S2C. List of all identified phosphoproteins from SILAC experiments in LPSCUR

| S. No. | log2 LPSCUR | Amino acid | Number of Phospho (STY) | Protein | Protein names | Gene names | Charge | Localization prob | PEP | Score | Delta score | Score for localization | Mass error [ppm] | Position | Positions within proteins | Sequence window |
|---|---|---|---|---|---|---|---|---|---|---|---|---|---|---|---|---|
| 1 | 7.70 | Y | 1 | G3V500 | Echinoderm microtubule-associated protein-like 1 | EML1 | 3 | 1.00 | 0.01 | 70.45 | 18.92 | 70.45 | -0.94 | 155 | 155;173;174;186;205 | SESKPKEPVFSAEEGYVKMFLRGRPVTMYMP |
| 2 | 3.39 | T | 1 | A6NHA9 | Olfactory receptor 4C46 | OR4C46 | 3 | 1.00 | 0.00 | 98.18 | 49.58 | 98.18 | -3.77 | 16 | 16 | MENRNNMTEFVLLGLTENPKMQKIIFVVFFV |
| 3 | 2.52 | T | 2 | O75691 | Small subunit processome component 20 homolog | UTP20 | 2 | 1.00 | 0.00 | 92.94 | 36.37 | 92.94 | 0.92 | 243 | 243 | LFEMCKGVRNMFHSCTGQAVKLILRKLGPVT |
| 4 | -0.94 | Y | 1 | Q658L1-2 | Protein FAM154B | FAM154B | 2 | 1.00 | 0.01 | 79.20 | 27.13 | 79.20 | -3.97 | 140 | 140;155 | RSTAPFNGITSHRLDYIPHQLELKFERPKEV |
