## Supplemental Table S3 for "Anti-inflammatory role of curcumin in Lipopolysaccharide treated A549 cells at global proteome level and on mycobacterial infection"

**Table S3. Number of phosphosites, charges in peptides and number of phosphogroups per peptide identified in CURI, LPSI and LPSCUR experiments.**

**Table S3A. Number of phosphosites**

| Experiments | Serine (S) | Threonine (T) | Tyrosine (Y) |
|---|---|---|---|
| **CURI** | 15 | 0 | 5 |
| **LPSI** | 19 | 6 | 1 |
| **LPSCUR** | 0 | 3 | 2 |

**Table S3B. Charges in peptides**

| Experiments | Doubly charge | Triply charge | Multiply charge |
|---|---|---|---|
| **CURI** | 14 | 1 | 0 |
| **LPSI** | 13 | 8 | 0 |
| **LPSCUR** | 2 | 2 | 0 |

**Table S3C. phosphogroups per peptide**

| Experiments | Singly phosphorylated | Doubly phosphorylated | Multiply phosphorylated |
|---|---|---|---|
| **CURI** | 9 | 6 | 0 |
| **LPSI** | 14 | 7 | 0 |
| **LPSCUR** | 3 | 1 | 0 |
